# Genomic and transcriptomic analysis of camptothecin producing novel fungal endophyte - *Alternaria burnsii* NCIM 1409

**DOI:** 10.1101/2023.05.14.540672

**Authors:** Shakunthala Natarajan, Boas Pucker, Smita Srivastava

## Abstract

Camptothecin is an important anticancer alkaloid produced by particular plant species. No suitable synthetic route has been established for camptothecin production yet, imposing a stress on plant-based production systems. Endophytes associated with these camptothecin-producing plants have been reported to also produce camptothecin and other high-value phytochemicals. A previous study identified a fungal endophyte *Alternaria burnsii* NCIM 1409, isolated from *Nothapodytes nimmoniana*, to be a sustainable producer of camptothecin. Our study provides key insights on camptothecin biosynthesis in this recently discovered endophyte. The whole genome sequence of *Alternaria burnsii* NCIM 1409 was assembled and screened for biosynthetic gene clusters. Comparative studies with related fungi supported the identification of candidate genes involved in camptothecin synthesis and also helped to understand some aspects of the endophyte’s defense against the toxic effects of camptothecin. No evidence for horizontal gene transfer of the camptothecin biosynthetic genes from the host plant to the endophyte was detected suggesting an independent evolution of the camptothecin biosynthesis in this fungus.

## Introduction

Humans are dependent on plants for a wide variety of natural products. Plants produce the active molecules for a majority of the drugs available in the market or at least inspired the drug design^1^. Anticancer drugs derived from plants occupy the top ladder in this list with active compounds like taxol and camptothecin leading in the front^2^. Camptothecin is the third most in-demand alkaloid mainly produced by the medicinal plants, *Camptotheca acuminata* and *Nothapodytes nimmoniana*^3^ (Fig. 1).

**Figure 1:**
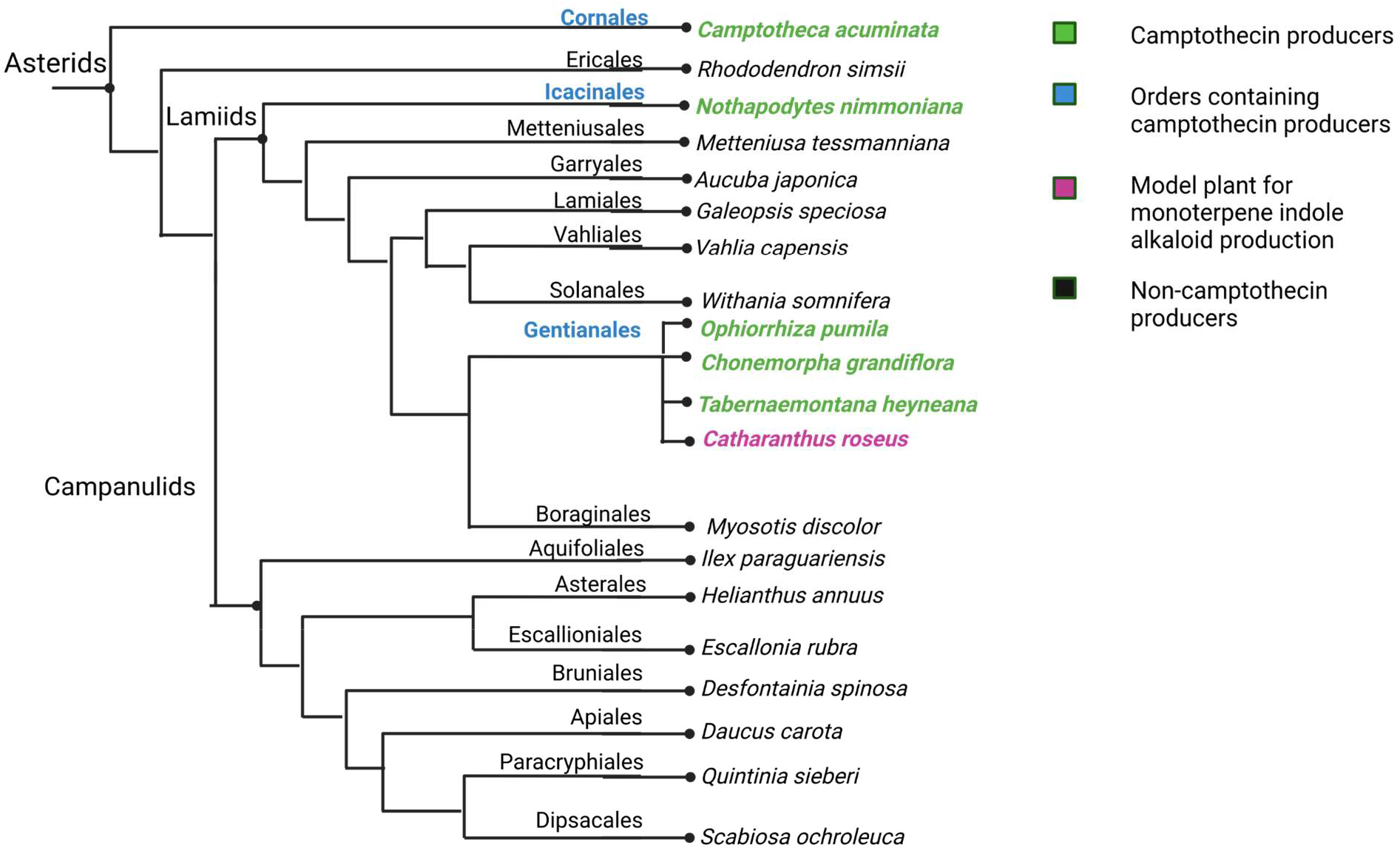
Phylogenetic tree of major camptothecin producing plants and some closely related species. Camptothecin-producing plant species are highlighted in green.

Camptothecin binds to the DNA topoisomerase-I, stalls the replication process, and causes cell death. This pharmacological effect makes camptothecin a valuable anticancer molecule. The mechanism of action of camptothecin takes place only during the S-phase of the cell cycle. Since cancer cells spend a greater proportion of time in S-phase compared to non-cancerous cells, the probability of camptothecin binding to cancer cells is greater^4^. The various drugs derived from camptothecin like topotecan and irinotecan are included in the World Health Organization’s model list of essential medicines^5^. The increasing demand for camptothecin has led to an irrational exploitation of the producing plant species, pushing them towards extinction^6^. Slow growth of these plants makes it difficult to achieve a balance of utilization and regeneration by planting activities. Highly variable amounts of camptothecin in different plants pose another issue that makes it difficult to standardize and produce plant-based extracts with consistent levels of active ingredient^7^. There is also a high degree of heterogeneity in the metabolites produced by the same plant species located in different geographical regions^8^. Moreover, chemical synthesis of products synthesized by plants is difficult and often not eco-friendly due to the use of toxic solvents and harsh reagents^9^. Complex chemical structures like quinolone rings, which are a part of the phytochemical molecules, are extremely difficult to synthesize artificially, making chemical synthesis of such high value phytochemicals unfeasible. Plant cell cultures could be harnessed for the production of these molecules. But such methods are not cost-effective and a tad time consuming when compared to microbial production platforms^10^. The challenges associated with plant-based extraction, culturing and chemical synthesis routes, make microbial synthesis of phytochemicals a suitable method for achieving the goal of sustainable production of these high value products^3,11^. Plants and their resident microbes called endophytes have established intriguing partnerships through co-evolution. Endophytes are microbes (bacteria or fungi) that reside within the tissues of a plant without harming the host plant^7^. Some of these endophytes are able to synthesize the specialized metabolites produced by the host plants. The endophytes could get this ability to produce metabolites from the host plant through horizontal gene transfer (HGT). However, evolutionary and environmental constraints placed on the endophyte for mutual co-existence along with the host, could also cause an independent evolution of the capacity to synthesize the host-produced metabolites^12^. Tolerance towards host-produced metabolites is necessary for the endophyte to ensure a mutual coexistence. This tolerance might be a prerequisite for the evolution of the corresponding biosynthesis pathway. For example, a tolerance for camptothecin might have evolved first in the endophyte which paved the way for a later evolution of the camptothecin biosynthesis pathway^13^.

It is important to delineate the roles of both partners (plant, endophyte) in the secondary metabolite production: (A) the plant and the endophyte could be equal contributors in the metabolite production i.e. plant and endophyte would catalyze complementary reactions and rely on each other for a complete biosynthesis pathway (B) the metabolite production could take place independently in the plant as well as the endophyte. If it is the latter scenario, a sustainable microbial production route for the metabolite could be feasible^12^. Moreover, such a microbe must also be a sustainable producer of the metabolite under industrial conditions to become a suitable production host. Unfortunately, attenuation of the product is frequently observed in the endophyte during subsequent sub-culture cycles^14,15^. A probable reason that has been proposed is the absence of inducing or silencing genes in the axenic cultures of the endophyte^14^. In summary, microbial production of plant specialized metabolites in an endophyte requires the microorganism to produce the molecule independently of the plant and to show its sustained production over many subcultures.

*Alternaria burnsii* NCIM 1409 is a sustainable producer of camptothecin^11^ and does not exhibit a decline in yield over subsequent sub-cultures like other camptothecin-producing endophytes (Table 1). Therefore, *Alternaria burnsii* NCIM 1409 has a huge potential to be harnessed as an industrial production source of camptothecin.

**Table 1:** Camptothecin-producing fungal endophytes from two major camptothecin-producing plant species.

| **Fungal endophytes producing camptothecin** | **Host plant** | **Degree of attenuation of camptothecin** | **References** |
| --- | --- | --- | --- |
| *Fusarium solani* | *Camptotheca acuminata* | Gradual attenuation observed from 1<sup>st</sup> to 7<sup>th</sup> subculture | 15 |
| *Trichoderma atroviride* LY357 | *Camptotheca acuminata* | Stable up to 8 subculture cycles | 16 |
| *Diaporthe* sp. F18 | *Nothapodytes nimmoniana* | Stable up to 6 subculture cycles | 17 |
| *Alternaria burnsii* NCIM 1409 * | *Nothapodytes nimmoniana* | Stable even after 12 continuous subculture cycles | 3,11 |

In this study, we obtained important details about the aspects of camptothecin biosynthesis in the endophyte *A. burnsii* NCIM 1409 through genomic analysis. Biosynthetic gene cluster mining and comparative studies with related fungi and host plants revealed insights into the ability of *A. burnsii* NCIM 1409 to produce camptothecin as well as defend itself against it.

## Materials and methods

### Fungal culture, genomic DNA extraction and quality check

The spores of *A. burnsii* NCIM 1409 were inoculated in potato dextrose broth (Himedia) and cultured at 28°C, 120 rpm for 8 days. Eight-day old fungal liquid suspension was harvested by centrifugation at 10,000 rpm, 4 °C for 15 minutes. The separated fungal mycelia was ground into powder using liquid nitrogen. The fungal genomic DNA was isolated using the potassium acetate DNA extraction protocol^18^. The isolated DNA was checked for its quality using agarose gel electrophoresis and quantified by NanoDrop measurements. The DNA sample was sent to Eurofins Genomics India Pvt. Ltd. for sequencing.

### Library preparation and quality check for whole genome sequencing

The paired-end sequencing library was prepared from the QC passed genomic DNA sample using Illumina TruSeq Nano DNA Library Prep Kit. Briefly, approximately 200 ng of DNA was fragmented by Covaris M220 to generate a mean fragment distribution of 350bp. Covaris shearing generates dsDNA fragments with 3’ or 5’ overhangs. The fragments were then subjected to end-repair. This process converts the overhangs resulting from fragmentation into blunt ends using End Repair Mix. The 3’ to 5’ exonuclease activity of this mix removes the 3’ overhangs and the 5’ to 3’ polymerase activity fills in the 5’ overhangs followed by adapter ligation to the fragments. This strategy ensures a low rate of chimera (concatenated template) formation. The ligated products were size selected using AMPure XP beads. The size-selected products were PCR amplified with the index primer as described in the kit protocol indexing adapters were ligated to the ends of the DNA fragments, preparing them for hybridization onto a flow cell.

After obtaining the Qubit concentration for the library and the mean peak size from Agilent TapeStation profile, the PE Illumina library was loaded onto NextSeq500 for cluster generation and sequencing using 2x150 bp chemistry. Paired-end sequencing allows the template fragments to be sequenced in both the forward and reverse directions on NextSeq500. The adapters were designed to allow selective cleavage of the forward strands after re-synthesis of the reverse strand during sequencing. The copied reverse strand was then used to sequence from the opposite end of the fragment.

### Genome sequence generation, and assembly

The genome of *A. burnsii* NCIM 1409 was sequenced on an Illumina NextSeq500 at Eurofins Genomics India Pvt. Ltd. FASTQ files obtained through sequencing were quality checked using FastQC (v-0.11.9)^19^ and low quality reads were trimmed using Trimmomatic (v-0.39)^20^ (see Supplementary Methods for details). The genome assembly of the endophyte was generated using SPAdes (v-3.15.5)^21^ (see Supplementary Methods for details). The assembly quality was checked using QUAST (v-5.2.0)^22^ (see Supplementary Methods for details) and assembly statistics were obtained using a custom python script (contig_stats.py^23^).

### Fungal culture for RNA isolation

The spores of *A. burnsii* NCIM 1409 were inoculated in potato dextrose broth (Himedia) and cultured at 28°C, 120 rpm for 8 days. One day old fungal culture was harvested by centrifugation at 13,500 x g,4°C, for 10 minutes, washed in phosphate-buffered saline. Three one-day old fungal samples were harvested thus followed by snap-freezing using liquid nitrogen, sealing and storage at -80°C. Similarly, eight-day old fungal culture was harvested by centrifugation at 13,500 x g, 4°C, for 10 minutes, washed in phosphate-buffered saline. Three eight-day old fungal samples were harvested thus followed by snap-freezing using liquid nitrogen, sealing and storage at -80°C. All the six samples (three replicates on day 1 and three replicates on day 8) were dispatched on dry ice to Eurofins Genomics India Pvt. Ltd. for RNA isolation and RNA-seq.

### RNA isolation and quality check

Total RNA was isolated from the received fungal pellet using conventional TRIzol method followed by column purification using Quick RNA Plant MiniPrep Kit (Zymo Research). The qualities and quantities of the isolated RNA were checked on NanoDrop followed by Agilent TapeStation using High Sensitivity RNA ScreenTape.

### RNA-seq library preparation and quality check

The RNA-Seq paired end sequencing libraries were prepared from the QC passed RNA samples using NEBNext® Ultra™ II Directional RNA Library Prep Kit for Illumina (NEB) as per manufacturer’s instruction. Briefly, mRNA was enriched from the total RNA using Poly-T attached magnetic beads, followed by enzymatic fragmentation, first strand cDNA conversion using NEBNext First Strand Synthesis Enzyme Mix to facilitate RNA dependent synthesis. The 1st strand cDNA served as a template to synthesize the second strand using the second strand mix. The dscDNA was then purified using AMPure XP beads followed by A-tailing, adapter ligation and then enriched by limited no of PCR cycles.

### Transcriptome analysis with RNA-seq

After obtaining the Qubit concentration for the libraries and the mean peak sizes from Agilent TapeStation profile, the PE Illumina libraries were loaded onto NovaSeq6000 for cluster generation and sequencing. Paired-end sequencing allows the template fragments to be sequenced in both the forward and reverse directions on NovaSeq6000. The adapters were designed to allow selective cleavage of the forward strands after re-synthesis of the reverse strand during sequencing. The copied reverse strand was used to sequence from the opposite end of the fragment.

### Genome annotation

The RNA-seq reads of the endophyte were used as reference for the structural annotation process. Protein hints from a closely related *Alternaria* strain^24–26^ were also integrated with the RNA-Seq hints. BRAKER2^27^ and TSEBRA (v-1.0.3)^28^ (see Supplementary Methods for details) were used to produce the structural genome sequence annotation. The completeness of the resulting annotation was assessed using BUSCO v5.4.2^29^ (see Supplementary Methods for details). The lineage dataset used for BUSCO assessment was pleosporales_odb10^29^, as *Alternaria* fungi belong to the *Pleosporales* order. The CDS and peptide FASTA files were obtained from the genomic FASTA and GFF3 file using a custom python script (get_peps_from_gff3.py). The functional annotation of the predicted genes was obtained using InterProScan5^30^ (Supplementary Table S1) (see Supplementary Methods for details on parameters used). The gene prediction was cleaned to adhere to ENA specifications for data submission. During that process, two genes with duplicated feature location identifiers were removed and sequences shorter than 100bp were removed.

### Identification of biosynthetic gene clusters in the endophyte

In fungi, specialized metabolism genes encoding proteins that participate in the same biosynthetic pathway are often genomically clustered. Based on the possibility of genes involved in camptothecin synthesis being clustered, the specialized metabolite gene clusters in the fungal endophyte were predicted via antiSMASH 6.1.1^31^ (see Supplementary Methods for details).

### Investigation of horizontal gene transfer (HGT) between endophyte and host plant(s)

Peptide sequences of three camptothecin producing plants - *C. acuminata*^32^, *N. nimmoniana*^33^, and *O. pumila*^34^, and peptide sequences of a monoterpene indole alkaloid producing plant - *C. roseus*^35^, were subjected to a BLASTp analysis against the endophyte’s peptide sequences to investigate the occurrence of horizontal gene transfer from the plants to the endophyte. Additionally, peptide sequences of key enzymes involved in camptothecin and MIA synthesis in these plants, were also separately retrieved from the NCBI protein database and compared against the *A. burnsii* peptide sequences using BLASTp to avoid missing potential hits indicating horizontal gene transfer (Fig. 2). Sequences of common enzymes involved in the mevalonate pathway like acetyl-CoA acetyltransferase were not found in the CPT producing plants’ taxonomy or MIA producing plants’ taxonomy in the NCBI repository (Supplementary Table S2). Hence, few such enzyme candidates belonging to the mevalonate pathway common to most plants were taken from related plants like *W. somnifera* belonging to the *Asterids* clade, same as the MIA and CPT producing plants. The BLASTp results (Supplementary Table S3) were processed using a custom python scripts (blast2best.py, process_blast_results_v2.py) that applied the following filters to select a gene from the endophyte as a potential candidate, (1) the protein sequence must be longer than 50 amino acids and (2) show a similarity of greater than 55% to the target plant sequence, and (3) have a normalized bit score greater than 0.2. The normalized bit score is calculated by dividing the bit score of a hit against a hit of the query sequence against itself.

**Figure 2:**
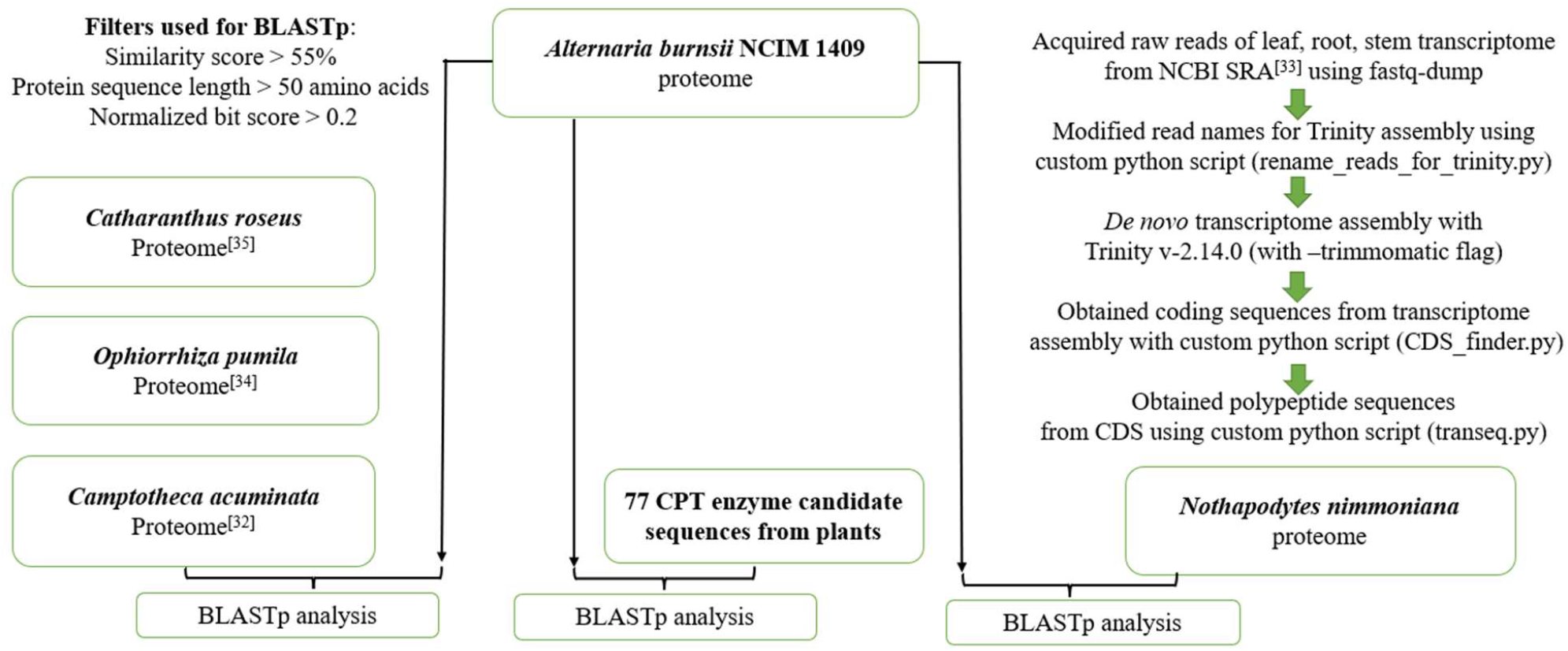
Schematic representation of horizontal gene transfer investigation

### Comparative genomics studies with related fungal organisms

Comparative studies of *A. burnsii* NCIM 1409 against closely related *Alternaria* fungi and other specialized metabolite producing fungi (Supplementary Table S4) were carried out using OrthoFinder2^36^. This analysis helped identify the genes unique to *A. burnsii* NCIM 1409 and lacking in the other fungi, by parsing the OrthoFinder2 results (Supplementary Table S5) using a custom python script (Process_orthofinder_for_unique_genes.py). Since gene duplications are important drivers of evolution and help organisms to acquire new functions^37^, it was speculated that the gene duplicates in the endophyte could reveal candidates involved in camptothecin production. Synteny analysis with JCVI/MCscan^38^ using a custom python script (jcvi_pairwise_synteny.py) was used to obtain pairwise syntenic blocks files connecting the annotations of *A. burnsii* NCIM 1409 and the *Alternaria* fungi. The blocks files were processed using a custom python script (Gene_duplications_synteny_v3.py) to obtain gene duplications in the fungal endophyte. Thus the comparative analyses with fungi helped consolidate some camptothecin candidate genes in *A. burnsii* NCIM 1409.

### Investigation of defense mechanism in the endophyte against the toxic effects of camptothecin

To understand the defense mechanism against camptothecin, the DNA topoisomerase I protein sequence of *A. burnsii* NCIM 1409 was retrieved and aligned with the DNA topoisomerase I sequences of *Homo sapiens*, other CPT-producing plants, non-CPT-producing fungi, and a CPT-producing fungus, using MAFFT (v-7.511)^39^. The aligned sequences were examined for specific CPT-resistance conferring mutations with the DNA topoisomerase I sequence from *Homo sapiens* serving as the reference for the amino acid residue positions.

## Results

### Assembly and annotation

The genome assembly of the endophyte comprises 104 contigs, with a total size of 33.2 Mb (Table 2). The N50 value of 832,062 bp indicates high assembly continuity.

**Table 2:** Statistics of the *A. burnsii* NCIM 1409 genome assembly

| Parameters | Values |
| --- | --- |
| N50 | 832,062 bp |
| Maximal contig length | 3,333,410 bp |
| Minimal contig length | 502 bp |
| Average contig length | 319,594 bp |
| Number of contigs | 104 |
| Assembly size | 33.2 Mb |

The structural annotation of the endophyte harbors 13,351 protein-encoding genes with an average gene size of 1434 bp. The longest gene has a size of 29413 bp. The BUSCO completeness score of this structural annotation was 98.6 % (C:98.6%[S:92.1%,D:6.5%],F:0.3%,M:1.1%,n:6641).

Mevalonate and shikimate pathways eventually lead to camptothecin synthesis and hence enzymes in these pathways are important for camptothecin biosynthesis^40^. The functional annotation of all predicted genes in *A. burnsii* NCIM 1409 identified 37 candidate genes (Supplementary Table S6) in the endophyte that coded for some enzymes in the mevalonate and shikimate pathways (Supplementary Figure).

### Biosynthetic gene clusters in *Alternaria burnsii* NCIM 1409

antiSMASH predicted 25 gene clusters in the fungus. The cluster details and the similarity of some clusters to known clusters are given in table 3 shown below.

**Table 3:** List of biosynthetic gene clusters predicted in the *A. burnsii* NCIM 1409 using antiSMASH 6.1.1

| Cluster | Genes in each cluster | Type | Similarity to known clusters |
| --- | --- | --- | --- |
| Cluster 1 | g3165-g3173 | Terpene |  |
| Cluster 2 | g3944-g3958 | T1PKS |  |
| Cluster 3 | g5907-g5926 | TIPKS, NRPS |  |
| Cluster 4 | g8313-g8329 | NRPS-like |  |
| Cluster 5 | g9283-g9298 | NRPS-like |  |
| Cluster 6 | g9546-g9566 | NRPS-like |  |
| Cluster 7 | g10734-g10752 | T1PKS |  |
| Cluster 8 | g11032-g11040 | Terpene |  |
| Cluster 9 | g11572-g11587 | NRPS | Dimethylcoprogen - NRP; 100% similarity |
| Cluster 10 | g11705-g11721 | NRPS |  |
| Cluster 11 | g12-g25 | NRPS-like, Indole |  |
| Cluster 12 | g1114-g1122 | Terpene |  |
| Cluster 13 | g1372-g1388 | T1PKS | Alternapyrone - Polyketide; 100 % similarity |
| Cluster 14 | g2119-g2127 | Terpene |  |
| Cluster 15 | g5532-g5540 | Terpene | PR-toxin - Terpene; 50% similarity |
| Cluster 16 | g6652-g6667 | NRPS-like |  |
| Cluster 17 | g7285-g7299 | T1PKS | Absciscic acid - Polyketide; 25% similarity |
| Cluster 18 | g7433-g7443 | T1PKS | Melanin - Polyketide; 100% similarity |
| Cluster 19 | g7661-g7674 | T1PKS | Depudecin - Polyketide iterative type I; 33 % similarity |
| Cluster 20 | g7683-g7689 | Terpene | Squalestatin S1 - Terpene; 40% similarity |
| Cluster 21 | g7842-g7856 | NRPS-like |  |
| Cluster 22 | g8594-g8603 | NRPS-like |  |
| Cluster 23 | g9129-g9140 | T1PKS |  |
| Cluster 24 | g9932-g9947 | T1PKS | Betaenone A/ betaenone B/ betaenone C - Polyketide; 62% similarity |
| Cluster 25 | g10060-g10068 | NRPS-like |  |

### Comparison of the *A. burnsii* NCIM 1409 gene set against genes of host plants

As endophytes reside in the host plant and acquire the ability to produce secondary metabolites, horizontal gene transfer of biosynthetic genes between them could be a possible occurrence^12,15,41^. If horizontal gene transfer of genes is detected, then this could reveal more genes involved in camptothecin synthesis in *A. burnsii* NCIM 1409 with a greater reliability. That would in turn provide a deeper understanding of camptothecin production in the endophyte. Camptothecin production has been found to be reported in 43 plant species belonging to different orders^42^. For example, *Camptotheca acuminata* belonging to *Cornales, Nothapodytes nimmoniana* belonging to *Icacinales*, and *Ophiorrhiza pumila* belonging to *Gentianales*, are all producers of camptothecin^43^. This phylogenetically scattered occurrence of camptothecin production capacity across different plant species is an intriguing phenomenon. Another medicinal plant called *Catharanthus roseus*, despite being a non-producer of camptothecin, is widely regarded as the model plant for monoterpene indole alkaloid synthesis and shares the same enzymes found in camptothecin producers, till a certain stage in its pathway leading to a wide variety of MIAs like vinblastine^35,43^. It is observed through comparative analyses of these medicinal plants that, the enzymes catalyzing the upper parts of the pathway leading to camptothecin or other MIAs in case of *C. roseus* share a high degree of similarity, whereas, towards the downstream steps of the pathway, the enzyme similarity, even among camptothecin producers becomes very low^43^. Due to such a divergence exhibited in the pathways leading to camptothecin and other MIAs in plants, the peptide sequences of all the four aforementioned plants were included in this investigation for horizontal gene transfer occurrence.

Despite extensive searches, no evidence was detected for horizontal gene transfer of the camptothecin biosynthesis from the host plant to the endophyte. A BLASTp screen of peptide sequences of *C. acuminata* and *C. roseus* against the endophyte revealed no significant hits. The BLASTp of peptide sequences of *N. nimmoniana* and *O. pumila* against the endophyte identified one gene (g616.t1) that showed a similarity greater than 55% and 59%, respectively, and a good normalized bit score. The BLASTp of candidates retrieved from the National Center for Biotechnology Information (NCBI) vs. the endophyte also identified g616.t1 as the best candidate. A similarity of 60% was observed in a comparison against the acetyl-CoA acetyltransferase sequence from *W. somnifera*. The functional annotation of this particular gene in the endophyte also indicated a function as acetyl-CoA acetyltransferase. This gene encodes for the initial enzyme in the mevalonate pathway, found in most plants and fungi. Also, the similarity values of around 60% do not indicate an evolutionarily recent HGT event. Further, those endophyte sequences that show high similarity to the plant sequences are involved in core functions like central metabolism, transcription, translation, which are highly conserved across organisms (Supplementary Table S7). Hence, there was no evidence to suggest horizontal gene transfer from the host plant to the endophyte.

### Comparison of the *A. burnsii* NCIM 1409 gene set against related fungi

Orthogroups were identified between a selection of fungal species to identify shared and private genes. The OrthoFinder2 analysis revealed 26 genes in the *A. burnsii* NCIM 1409 that were present in orthogroups not shared with genes from other investigated fungi. There were also some genes in the endophyte that were not present in the orthogroups file. There were also some species-specific single copy genes that could be unique to *A. burnsii* NCIM 1409, that were found by processing the OrthoFinder2 results using a custom python script (Process_orthofinder_for_unique_genes.py). In total, 233 unique genes were identified in the endophyte that did not have orthologs in other fungi. Since the comparison also included another fungus *Xylaria* sp. M71 that produces a camptothecin derivative (10-hydroxycamptothecin)^44^, genes that were only shared between *A. burnsii* NCIM 1409 and X. sp M71 were also searched. There was only one gene (g3550.t1) that was present only in these two organisms and it encoded a SAM-dependent methyltransferase. Hence, it was also included as a camptothecin candidate gene in *A. burnsii* NCIM 1409 along with the other genes mined from the OrthoFinder2 analysis. Next, the search for gene duplicates based on synteny analysis yielded 215 gene duplicates in *A. burnsii*, while all the other *Alternaria* fungi showed only one corresponding gene. Some of these candidate genes were identified by both the OrthoFinder and synteny analyses, making them more important ones for further investigation (Supplementary Table S6). After obtaining the peptide sequences of the identified candidates, those sequences that were too short (less than 30 amino acids in length) and those that appeared to be artifacts were removed from the candidate gene list. Thus, after cleaning the candidate genes using the above mentioned criteria, the comparative analyses with other fungi helped obtain a total of 449 CPT candidate genes in the endophyte (Supplementary Table S6, Supplementary Results).

### Analysis of DNA Topoisomerase I in *Alternaria burnsii* NCIM 1409

Since camptothecin is a toxic molecule that inhibits replication by binding to DNA topoisomerase I, it is quite interesting to see how camptothecin producers avoid detrimental impacts of their end product. Knowledge about this defense mechanism is important to understand camptothecin biosynthesis in the endophyte. A multiple sequence alignment (MSA) showed various critical amino acid residues of DNA topoisomerase I sequence of *A. burnsii* NCIM 1409. It also helped look out for well-known mutations modulating camptothecin and DNA topoisomerase binding, in the DNA topoisomerase I sequence of *A. burnsii* NCIM 1409 (Supplementary Tables S8-S12).

The camptothecin-producing endophytes, as well as non-camptothecin producing close relatives and distant fungi did not show the three camptothecin-resistance conferring mutations - N421K, L530I, N722S found in camptothecin producing plants^45^. Sequences of all species displayed N in the position of interest, which matches the amino acids in the corresponding sequences of camptothecin-producing plant *O. pumila* and non-camptothecin plant *C. roseus*. All the fungi included in this analysis have L corresponding to L530 and share this with camptothecin-producing *C. acuminata*, and non-camptothecin plant *C. roseus*. All of them have N corresponding to N722 and share this with N in non-camptothecin plant *C. roseus* and camptothecin producer *N. nimmoniana*. The catalytic amino acid residues and some camptothecin resistance conferring wild-type amino acid residues are highly conserved in camptothecin producers and non-camptothecin producers across all organisms. For example, N352 and F361, two residues in the binding region, are important for camptothecin resistance^46^. They are found in all the sequences used in this study and are not exclusive to camptothecin producers. Among other camptothecin resistance conferring mutations in the binding region, only M370T^47^ mutation is observed in the fungal sequences (except *N. aurantialba*) and in *N. nimmoniana*. Again, this mutation is not exclusive to camptothecin-producing endophytes. The other camptothecin resistance conferring mutations were not detected in the binding regions in the sequences considered here. The remaining residues in the binding region (E356, H367, V502, Y619, D725)^13^ were found to be intact without any variation across all the sequences. Critical residues in DNA topoisomerase I sequences from two fungi – *A. burnsii* NCIM 1409 producing camptothecin, and *Xylaria sp. M71* producing 10-Hydroxy-camptothecin show no variation and are highly similar to each other as well as to other compared sequences of fungi that are not reported producers of camptothecin.

## Discussion

The genomic and transcriptomic analysis of the novel fungal endophyte *Alternaria burnsii* NCIM 1409 was carried out for the first time in this study. The assembled genome sequence provides a basis for future studies. RNA-seq data sets support the predicted gene models. Potential functions have been assigned to most genes.

The specialized metabolite genes in fungi can be under control of a shared regulatory network resulting in simultaneous activity of all genes in the gene cluster. Among such clusters, terpene biosynthesis gene clusters seem to play a major role in facilitating signal transfer between the host and endophyte during plant-microbe interactions^48^. The 25 gene clusters identified in *A. burnsii* NCIM 1409 could provide potential clues to understand specialized metabolite synthesis in the endophyte. With camptothecin being a monoterpene indole alkaloid, the terpene clusters and the hybrid NRPS-like, indole cluster 11 (Table 2), could harbor potential genes involved in camptothecin biosynthesis. The novel endophyte in focus was isolated from *Nothapodytes nimmoniana*. The camptothecin biosynthesis pathway in the host plant is complex, starting from the mevalonate (MVA) pathway or the methylerythritol phosphate (MEP) pathway and the shikimate pathway. The MVA or the MEP pathway produce isoprenoid molecules that are modified further to produce secologanin. The shikimate pathway produces tryptophan that condenses with secologanin to produce ‘strictosidine synthase’ (STR) - the parent molecule for a wide variety of MIAs. The steps of the pathway leading to camptothecin, after the Pictet Spengler reaction catalyzed by STR, have eluded researchers so far^33,49^. Since the endophyte isolated from the host, retained its ability to produce camptothecin for over 12 sub-culture cycles independent of the host^3,11^, it was expected that during its life cycle in the host tissues, some of the key genes encoding crucial enzymes in camptothecin synthesis might have been transferred from the host plant to the endophyte. This prompted the investigation to assess for the occurrence of horizontal gene transfer between the host and the fungus. If horizontal gene transfer would have been detected, then that could have revealed more genes involved in camptothecin synthesis in *A. burnsii* NCIM 1409. However, the present study does not reveal any evidence for HGT from the host to the endophyte. This observation is also corroborated by a few other studies that investigated HGT from host plants to endophytes producing similar specialized metabolites. Extremely low similarity was found between fungal sequences of taxol producing fungi isolated from the yew tree and taxane specific genes in the yew tree^50^. HGT was also found to be absent between the paclitaxel producing fungus *Penicillium aurantiogriseum* NRRL 62431 isolated from the hazel plant (*Corylus avellana)*^51^. The researchers hypothesize that the taxol-producing genes in the endophyte possess a completely different evolutionary pattern. These reports and the inferences from the present study indicate the possibility for independent evolution of camptothecin biosynthesis in *A. burnsii* NCIM 1409. Although the HGT hypothesis was not quite helpful in finding camptothecin candidate genes in the endophyte, comparative studies with other fungi revealed a number of candidate genes. The fungi included in the comparative analysis were classified into three bins - ‘camptothecin producers’, ‘camptothecin non-producers’ and ‘Related product producers’ (fungi producing specialized metabolites like taxol and other specialized metabolites similar to camptothecin, but do not produce camptothecin itself) (Supplementary Table S4). This classification developed for the comparison was used as a basis to identify the candidate genes in the novel endophytes with respect to the other fungi. The endophyte genes that are not shared with the other (all the three classes) fungi can be considered to be specifically unique to itself. Camptothecin synthesis being a unique trait found in this endophyte, these unique genes could possibly hold the key for unraveling more aspects of camptothecin biosynthesis in the fungus and in fungi in general. Fungi could form a reservoir of camptothecin biosynthesis enzymes with properties desirable for biotechnological applications. Genes that are exclusively shared between camptothecin producers and are absent from other fungi, could be promising targets of future studies.

Synteny analysis was crucial for the identification of gene duplications in the endophyte with respect to closely related *Alternaria* fungi. Gene duplications are major drivers of evolutionary innovations as they enable neo- and subfunctionalization^37^. Duplications lower the constraints on natural selection processes, and play a significant role in causing new functions to appear in organisms^52^. Since camptothecin production is a novel trait in the endophyte, such duplications could play an important role in camptothecin synthesis. While focusing on the novel endophyte as a source organism for camptothecin production, it would be important to understand the mechanism being used by the endophyte to protect itself against the toxic effects of camptothecin. This tolerance mechanism could be harnessed to enhance the camptothecin yield from the producing organism. It was reported that the DNA Topoisomerase I in plants possess certain mutations that make them self-resistant to the camptothecin molecule they produce^45^. There have not been many reports investigating the resistance mechanism in camptothecin-producing endophytes, except for one by Kusari and co-workers (2011)^13^. However, no specific resistance-endowing mutations were detected in another camptothecin-producing endophyte isolated from *C. acuminata*. It was proposed that the fungus could be using some other mechanism to protect itself from the deleterious effects of camptothecin^13^. Based on the DNA topoisomerase I analysis of the endophyte in this work, it is evident that, camptothecin resistance and camptothecin production in fungal endophytes need not have co-evolved like the co-evolution of camptothecin resistance and camptothecin production in camptothecin-producing plants, as proposed by^45^. This can be further explained by the observation that non-camptothecin producing fungi too have some of the camptothecin resistance conferring amino acid residues. These inferences agree with the conclusions put forth by Kusari and co-workers^13^, which convey that fungi that colonize a toxic metabolite producing plant like a camptothecin producing one, must possess innate resistance to overcome the toxicity, and invade the plant. Of the invaders some may prove to be camptothecin producers while, some may not, as seen in the isolation of several camptothecin producing and non-producing endophytes from different camptothecin producing plants like *C. acuminata*^15^ and *N. nimmoniana*^11^. Although previously studied and expected mutations in DNA topoisomerase I were not present in the fungal sequences, it remains possible that these fungi could have evolved completely different mutations to combat camptothecin as hinted by^13^. But it is interesting to note the fact that *A. burnsii* NCIM 1409 exhibits resistance to camptothecin in a dose dependent manner^3^. This reduces the chances of mutations appearing in the DNA topoisomerase I sequence to create a completely foolproof camptothecin resistant sequence, and this view is also buttressed by dose-dependent resistance to camptothecin exhibited by camptothecin-producing fungus *Phomopsis sp*. isolated from *N. nimmoniana*^53^. This could mean that *A. burnsii* NCIM 1409 could have developed entirely new ways of resisting camptothecin apart from variation in its DNA topoisomerase I sequence. Or rather than resisting camptothecin, we speculate that the fungus might tolerate camptothecin up to a particular level, after which the fate of camptothecin within the producer needs to be investigated further.

## Supporting information

Supplementary Results

## Conclusion

Our work provided insights into the camptothecin biosynthesis ability of the fungal endophyte *A. burnsii* NCIM 1409, an camptothecin producer. A lack of evidence for horizontal transfer of camptothecin biosynthesis genes from the host plant to the endophyte suggests an independent evolution of the camptothecin biosynthesis pathway in the fungus. Comparative studies with other fungi narrowed down camptothecin candidate genes in the endophyte that could be validated in further analyses. The fungal endophyte does not possess unique and specific camptothecin-resistance conferring variations in its DNA topoisomerase I sequence. It seems to use a distinct mechanism to protect itself from the deleterious effects of the camptothecin molecule it produces. This study paved the way for further exploration of the genetic mechanisms underlying the camptothecin production in the endophyte. In the future, this could help to establish *Alternaria burnsii* NCIM 1409, as a sustainable production platform for camptothecin on a large scale.

### Abbreviations

PCR: Polymerase chain reaction;
PE: Paired-end;
RNA-seq: RNA sequencing;
cDNA: Complementary deoxyribonucleic acid;
HGT: Horizontal gene transfer;
MIA: Monoterpene indole alkaloids;
STR: Strictosidine synthase;
MSA: Multiple sequence alignment;
MVA: Mevalonate pathway;
MEP: Methylerythritol pathway;
T1PKS: Type I polyketide synthase;
NRPS: Non ribosomal peptide synthetase;
SAM-dependent methyltransferase: S-adenosyl-methionine-dependent methyltransferase

## Data availability

The Whole Genome Shotgun project has been deposited at ENA under the accession ERZ18273747. The ENA project accession number is PRJEB61631.

The plant and fungal annotations generated as a part of the study can be accessed at: https://github.com/ShakunthalaNatarajan/GenomeAssembly_AburnsiiNCIM1409/tree/main/Annotations

The Multiple sequence alignment FASTA file can be accessed at: https://github.com/ShakunthalaNatarajan/GenomeAssembly_AburnsiiNCIM1409/tree/main/MSA_file

## Code availability

The scripts and codes used in this study can be accessed at :

https://github.com/ShakunthalaNatarajan/GenomeAssembly_AburnsiiNCIM1409

## Acknowledgements

Authors would like to thank Karthigeyan S.P. for his assistance and support with the experimental aspects of the work. This work was supported by the BMBF-funded de.NBI Cloud within the German Network for Bioinformatics Infrastructure (de.NBI) (031A532B, 031A533A, 031A533B, 031A534A, 031A535A, 031A537A, 031A537B, 031A537C, 031A537D, 031A538A). This work was facilitated through the Combined Study and Practice Stays for Engineers from Developing Countries (KOSPIE) program of the German Academic Exchange Service (DAAD). bioRender.com was used to construct some of the figures.

## Ethics approval and consent to participate

Not applicable

## Consent for publication

Not applicable

## Author contributions

SN, BP, and SS designed the study. SN performed the bioinformatic analyses and wrote the manuscript. BP and SS supervised the work and revised the manuscript. All authors read and approved the final version of the manuscript.

## References

1. Atanasov, A. G. et al. Discovery and resupply of pharmacologically active plant-derived natural products: A review. Biotechnol. Adv.33, 1582–1614 (2015).

2. Swamy, M. K. et al. Biotechnology of camptothecin production in Nothapodytes nimmoniana, Ophiorrhiza sp. and Camptotheca acuminata. Appl. Microbiol. Biotechnol.105, 9089–9102 (2021).

3. Mohinudeen, I. A. H. K., Pandey, S., Kanniyappan, H., Muthuvijayan, V. & Srivastava, S. Screening and selection of camptothecin producing endophytes from Nothapodytes nimmoniana. Sci. Rep.11, 11205 (2021).

4. Almeida, A., Fernandes, E., Sarmento, B. & Lúcio, M. A Biophysical Insight of Camptothecin Biodistribution: Towards a Molecular Understanding of Its Pharmacokinetic Issues. Pharmaceutics13, 869 (2021).

5. Nguyen, T.-A. M. et al. Discovering and harnessing oxidative enzymes for chemoenzymatic synthesis and diversification of anticancer camptothecin analogues. Commun. Chem.4, 1–7 (2021).

6. Shrivastava, V., Sharma, N., Shrivastava, V. & Sharma, A. Review on Camptothecin Producing Medicinal Plant: Nothapodytes Nimmoniana. Biomed. Pharmacol. J.14, 1799–1813 (2021).

7. Venugopalan, A. & Srivastava, S. Endophytes as in vitro production platforms of high value plant secondary metabolites. Biotechnol. Adv.33, 873–887 (2015).

8. Narayani, M., Chadha, A. & Srivastava, S. Cyclotides from the Indian Medicinal Plant Viola odorata (Banafsha): Identification and Characterization. J. Nat. Prod.80, 1972–1980 (2017).

9. Kharissova, O. V., Kharisov, B. I., Oliva González, C. M., Méndez, Y. P. & López, I. Greener synthesis of chemical compounds and materials. R. Soc. Open Sci.6, 191378 (2019).

10. Li, S., Li, Y. & Smolke, C. D. Strategies for microbial synthesis of high-value phytochemicals. Nat. Chem.10, 395–404 (2018).

11. Mohinudeen, I. A. H. K. et al. Sustainable production of camptothecin from an Alternaria sp. isolated from Nothapodytes nimmoniana. Sci. Rep.11, 1478 (2021).

12. Ludwig-Müller, J. Plants and endophytes: equal partners in secondary metabolite production? Biotechnol. Lett.37, 1325–1334 (2015).

13. Kusari, S., Košuth, J., Čellárová, E. & Spiteller, M. Survival-strategies of endophytic Fusarium solani against indigenous camptothecin biosynthesis. Fungal Ecol.4, 219–223 (2011).

14. Bielecka, M., Pencakowski, B. & Nicoletti, R. Using Next-Generation Sequencing Technology to Explore Genetic Pathways in Endophytic Fungi in the Syntheses of Plant Bioactive Metabolites. Agriculture12, 187 (2022).

15. Kusari, S., Zühlke, S. & Spiteller, M. An Endophytic Fungus from Camptotheca acuminata That Produces Camptothecin and Analogues. J. Nat. Prod.72, 2–7 (2009).

16. Xiang, P. et al. Camptothecin-producing endophytic fungus Trichoderma atroviride LY357: Isolation, identification, and fermentation conditions optimization for camptothecin production. Appl. Microbiol. Biotechnol.97, (2013).

17. Degambada, K. D., Kumara, P. A. A. S. P., Salim, N., Abeysekera, A. M. & Chandrika, U.G. Diaporthe sp. F18; a new source of camptothecin-producing endophytic fungus from Nothapodytes nimmoniana growing in Sri Lanka. Nat. Prod. Res.0, 1–6 (2021).

18. Feng, J. et al. An inexpensive method for extraction of genomic DNA from fungal mycelia. Can. J. Plant Pathol.32, 396–401 (2010).

19. Babraham Bioinformatics - FastQC A Quality Control tool for High Throughput Sequence Data. https://www.bioinformatics.babraham.ac.uk/projects/fastqc/.

20. Bolger, A. M., Lohse, M. & Usadel, B. Trimmomatic: a flexible trimmer for Illumina sequence data. Bioinforma. Oxf. Engl.30, 2114–2120 (2014).

21. Prjibelski, A., Antipov, D., Meleshko, D., Lapidus, A. & Korobeynikov, A. Using SPAdes De Novo Assembler. Curr. Protoc. Bioinforma.70, e102 (2020).

22. Gurevich, A., Saveliev, V., Vyahhi, N. & Tesler, G. QUAST: quality assessment tool for genome assemblies. Bioinforma. Oxf. Engl.29, 1072–1075 (2013).

23. Pucker, B. et al. A De Novo Genome Sequence Assembly of the Arabidopsis thaliana Accession Niederzenz-1 Displays Presence/Absence Variation and Strong Synteny. PLOS ONE11, e0164321 (2016).

24. Dang, H., Pryor, B., Peever, T. & Lawrence, C. The Alternaria genomes database: A comprehensive resource for a fungal genus comprised of saprophytes, plant pathogens, and allergenic species. BMC Genomics16, (2015).

25. Mesny, F. et al. Genetic determinants of endophytism in the Arabidopsis root mycobiome. Nat. Commun.12, 7227 (2021).

26. Zeiner, C. A. et al. Comparative Analysis of Secretome Profiles of Manganese(II)-Oxidizing Ascomycete Fungi. PloS One11, e0157844 (2016).

27. Brůna, T., Hoff, K. J., Lomsadze, A., Stanke, M. & Borodovsky, M. BRAKER2: automatic eukaryotic genome annotation with GeneMark-EP+ and AUGUSTUS supported by a protein database. NAR Genomics Bioinforma.3, lqaa108 (2021).

28. Gabriel, L., Hoff, K. J., Brůna, T., Borodovsky, M. & Stanke, M. TSEBRA: transcript selector for BRAKER. BMC Bioinformatics22, 566 (2021).

29. Manni, M., Berkeley, M. R., Seppey, M., Simão, F. A. & Zdobnov, E. M. BUSCO Update: Novel and Streamlined Workflows along with Broader and Deeper Phylogenetic Coverage for Scoring of Eukaryotic, Prokaryotic, and Viral Genomes. Mol. Biol. Evol.38, 4647–4654 (2021).

30. Jones, P. et al. InterProScan 5: genome-scale protein function classification. Bioinforma. Oxf. Engl.30, 1236–1240 (2014).

31. Blin, K. et al. antiSMASH 6.0: improving cluster detection and comparison capabilities. Nucleic Acids Res.49, W29–W35 (2021).

32. Zhao, D. et al. De novo genome assembly of Camptotheca acuminata, a natural source of the anti-cancer compound camptothecin. GigaScience6, gix065 (2017).

33. Rather, G. A. et al. De novo transcriptome analyses reveals putative pathway genes involved in biosynthesis and regulation of camptothecin in Nothapodytes nimmoniana (Graham) Mabb. Plant Mol. Biol.96, 197–215 (2018).

34. Rai, A. et al. Chromosome-level genome assembly of Ophiorrhiza pumila reveals the evolution of camptothecin biosynthesis. Nat. Commun.12, 405 (2021).

35. Kellner, F. et al. Genome-guided investigation of plant natural product biosynthesis. Plant J. Cell Mol. Biol.82, 680–692 (2015).

36. Emms, D. & Kelly, S. OrthoFinder2: fast and accurate phylogenomic orthology analysis from gene sequences. (2018). doi:10.1101/466201.

37. Lallemand, T., Leduc, M., Landès, C., Rizzon, C. & Lerat, E. An Overview of Duplicated Gene Detection Methods: Why the Duplication Mechanism Has to Be Accounted for in Their Choice. Genes11, 1046 (2020).

38. Tang, H. et al. tanghaibao/jcvi: JCVI v0.7.5. (2017) doi:10.5281/zenodo.846919.

39. Katoh, K. & Standley, D. M. MAFFT multiple sequence alignment software version 7: improvements in performance and usability. Mol. Biol. Evol.30, 772–780 (2013).

40. Sadre, R. et al. Metabolite Diversity in Alkaloid Biosynthesis: A Multilane (Diastereomer) Highway for Camptothecin Synthesis in Camptotheca acuminata. Plant Cell28, 1926–1944 (2016).

41. Kusari, S., Hertweck, C. & Spiteller, M. Chemical Ecology of Endophytic Fungi: Origins of Secondary Metabolites. Chem. Biol.19, 792–798 (2012).

42. Pu, X. et al. Possible clues for camptothecin biosynthesis from the metabolites in camptothecin-producing plants. Fitoterapia134, 113–128 (2019).

43. Yang, M. et al. Divergent camptothecin biosynthetic pathway in Ophiorrhiza pumila. BMC Biol.19, 1–16 (2021).

44. Ding, X., Liu, K., Zhang, Y. & Liu, F. De novo transcriptome assembly and characterization of the 10-hydroxycamptothecin-producing Xylaria sp. M71 following salicylic acid treatment. J. Microbiol.55, 871–876 (2017).

45. Sirikantaramas, S., Yamazaki, M. & Saito, K. Mutations in topoisomerase I as a self-resistance mechanism coevolved with the production of the anticancer alkaloid camptothecin in plants. Proc. Natl. Acad. Sci.105, 6782–6786 (2008).

46. Staker, B. L. et al. Structures of three classes of anticancer agents bound to the human topoisomerase I-DNA covalent complex. J. Med. Chem.48, 2336–2345 (2005).

47. Fujimori, A., Harker, W. G., Kohlhagen, G., Hoki, Y. & Pommier, Y. Mutation at the catalytic site of topoisomerase I in CEM/C2, a human leukemia cell line resistant to camptothecin. Cancer Res.55, 1339–1346 (1995).

48. Collemare, J. & Lebrun, M.-H. Fungal Secondary Metabolites: Ancient Toxins and Novel Effectors in Plant–Microbe Interactions. in Effectors in Plant–Microbe Interactions 377–400 (John Wiley & Sons, Ltd, 2011). doi:10.1002/9781119949138.ch15.

49. Manjunatha, B. L. et al. Transcriptome analysis of stem wood of Nothapodytes nimmoniana (Graham) Mabb. identifies genes associated with biosynthesis of camptothecin, an anti-carcinogenic molecule. J. Biosci.41, 119–131 (2016).

50. Heinig, U., Scholz, S. & Jennewein, S. Getting to the bottom of Taxol biosynthesis by fungi. Fungal Divers.60, 161–170 (2013).

51. Yang, Y. et al. Genome sequencing and analysis of the paclitaxel-producing endophytic fungus Penicillium aurantiogriseum NRRL 62431. BMC Genomics15, 69 (2014).

52. Xu, Z. et al. Tandem gene duplications drive divergent evolution of caffeine and crocin biosynthetic pathways in plants. BMC Biol.18, 63 (2020).

53. Shweta, S. et al. Inhibition of fungal endophytes by camptothecine produced by their host plant, Nothapodytes nimmoniana (Grahm) Mabb. (Icacinaceae). Curr. Sci.107, 994–1000 (2014).

54. Miao, L. Y. et al. Transcriptome analysis of a Taxol-producing endophytic fungus Cladosporium cladosporioides MD2. AMB Express8, 41 (2018).

55. Che, J., Shi, J., Gao, Z. & Zhang, Y. Transcriptome analysis reveals the genetic basis of the resveratrol biosynthesis pathway in an endophytic fungus (Alternaria sp. MG1) isolated from Vitis vinifera. Front. Microbiol.7, 1257 (2016).

56. Tao, J. et al. Whole-genome sequence analysis of an endophytic fungus Alternaria sp. SPS-2 and its biosynthetic potential of bioactive secondary metabolites. Microorganisms10, 1789 (2022).

57. Sun, T. et al. Whole Genome Sequencing and Annotation of Naematelia aurantialba (Basidiomycota, Edible-Medicinal Fungi). J. Fungi (Basel)8, article 1 (2021).

58. Chang, J. Y., Liu, J. F., Juang, S. H., Liu, T. W. & Chen, L. T. Novel mutation of topoisomerase I in rendering cells resistant to camptothecin. Cancer Res.62, 3716–3721 (2002).

59. Losasso, C. et al. A single mutation in the 729 residue modulates human DNA topoisomerase IB DNA binding and drug resistance. Nucleic Acids Res.36, 5635–5644 (2008).

