## Supplementary Figure for "Genomic and transcriptomic analysis of camptothecin producing novel fungal endophyte - *Alternaria burnsii* NCIM 1409"

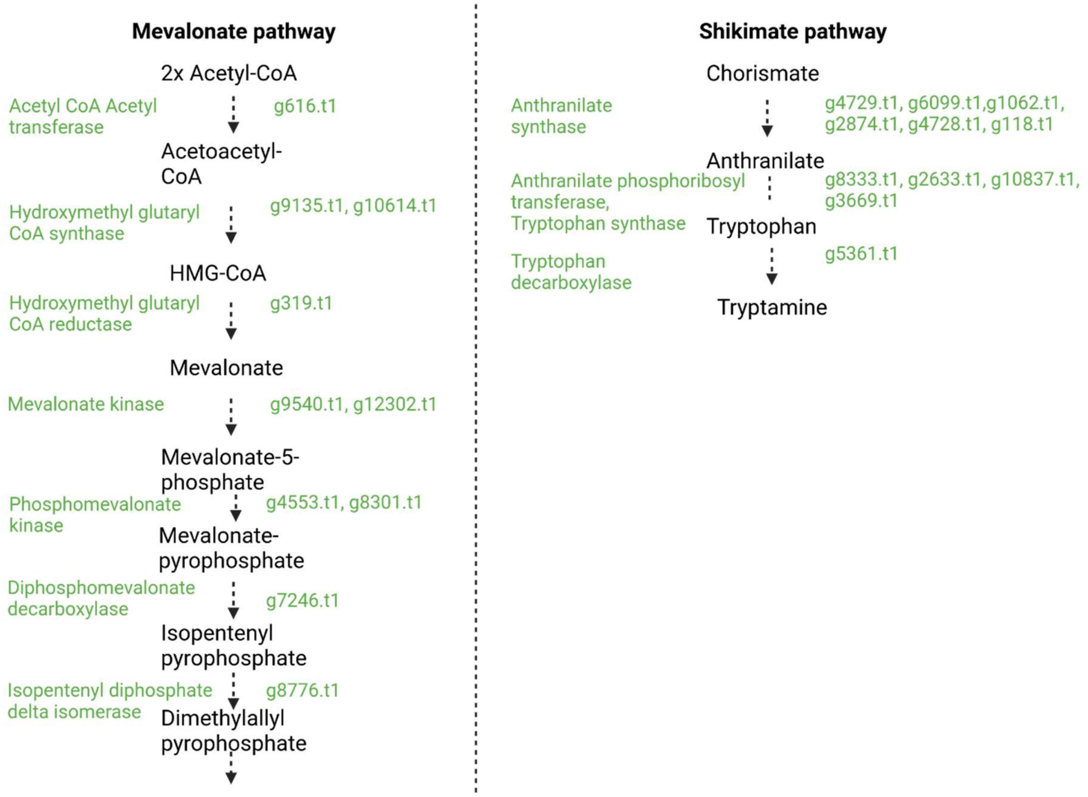


**Supplementary Figure:** Candidate genes of the endophyte involved in mevalonate and shikimate pathways
