## Supplementary Methods for "Genomic and transcriptomic analysis of camptothecin producing novel fungal endophyte - *Alternaria burnsii* NCIM 1409"

**1. Trimmomatic(v-0.39) parameters -**

adapters:TruSeq3-PE.fa:2:30:10:2:True

LEADING:20

TRAILING:20

SLIDINGWINDOW:10:20

MINLEN:50

**2. SPAdes(v-3.15.5)-**

default parameters were used along with the --careful flag

**3. QUAST(v-5.2.0)-**

default parameters with --fungus flag

**4. BRAKER2 parameters-**

a. integrating RNA-Seq hints-

default parameters with --fungus --bam flags

b. integrating protein hints-

default parameters with--fungus --prot_seq --prg=gth --trainFromGth flags

**5. TSEBRA(v-1.0.3)-**

-g augustus.hints.gtf (from RNA-Seq hints), augustus.hints.gtf (from protein hints), -c default config file -e hintsfile.gff (from RNA-seq hints), hintsfile.gff (from protein hints)

#TSEBRA output was a GTF file

#steps to convert GTF to GFF3 file

rename_gtf.py within TSEBRA bin folder was used with --translation_tab flag to rename the GTF file to process it using gtf2gff.pl script provided in AUGUSTUS

gtf2gff.pl script in AUGUSTUS folder was used to convert the GTF file to GFF3 file; get_peps_from_gff3.py was used to obtain peptide and CDS FASTA files from the GFF3 file

**6. BUSCO (v-5.4.2) parameters-**

default parameters with -m prot -l pleosporales_odb10 flags

**7. InterProScan5 parameters-**

default parameters with the following optional flags: -dp --goterms --pathways

**8. antiSMASH (v-6.1.1) parameters-**

default parameters with the following additional flags: --taxon fungi --pfam2go --cb-knownclusters --cassis
