## Supplementary Results for "Genomic and transcriptomic analysis of camptothecin producing novel fungal endophyte - *Alternaria burnsii* NCIM 1409"

**Sequences of candidate genes identified**

>g2175.t1

MSQIKTMANDKDSFLGKVAGFAAPNYRLSPNLEADAPTTPDPAASAKHPDHLQDVIYAIKHLYKQYDISQSMGYILV
GPSCGATLAFQVATYSLNRHQSVIPSPLGVIGLNGVYDLAGWLRVASSSVYDDIVETAFGSDREIWEKASPMHQVQ
SDWTVDHHTAKPISLTFLAYSLEDTVVADSQTLEMLEKLKHWSQPHQATVNTGSILSPPSAPDEQQQRIVLSKVING
GHDEVWEKPEQMLACIAEAVKKVCGMD*

>g2176.t1

MTFECLKSVSYPAPPDQPQTRLHTIDIWLPQQQAAFPLNERVWIVFVHGGAWRNPFQSSECFEPAMSQIKTMAN
DKDSFLGKVAGFAAPNYRLSPNLEADAPTTPDPAASAKHPDHLQDVIYAIKHLYKQYDISQSMGYILVGPSCGATLAF
QVATYSLNRHQSVIPSPLGVIGLNGVYDLAGWLRVASSSVYDDIVETAFGSDREIWEKASPMHQVQSDWTVDHHT
AKPISLTFLAYSLEDTVVADSQTLEMLEKLKHWSQPHQATVNTGSILSPPSAPDEQQQRIVLSKVINGGHDEVWEKP
EQMLACIAEAVKKVCGMD*

>g3996.t1

MHIIWCFVAALLSGGALADCPAGPYKMGSLCYKGCLGVARCGEDNHITICREAVTDIFVWQVGAQCNHCKGGKCV
*

>g3997.t1

MYISWCFIAALLSSSALADCPNGPYSTNSECPGKCYGFQRCGDYNQVIRCETSVGGGTRWIGKQWCKHCKWGGC
DS*

>g7667.t1

MAKSIHVVSKLDNAKHAVFHIDEDPPSLAASSVRVQTWLVSLTYNNLTYARSGTPLHWWDTYPVPEASPTPFNSWE
KWGIVPAWGYGRVLESTNDAIAPGSLLWGMWPTSEHTVDLQLEAIEPSGHWLERSAQRSKLMTVYNSYEQVSESD
AQTMRMTALCKPLWQGPHLLNTSVFSARRIHPFGFGAPWSQEDADLSSAVVVSLSASSKTGRSFGWEMARNRDV
SKHGPRALIQMTSVPNTLSEYDSSLPMRAAAYDDTSAMAWAEHFKPSRVVIVDFGASDAVLESVRATASKLAPKVT
VVAVGYEAKVYTQEEIAARMATASTKVPVNTSGMRDRVIESQGALEFSKELDGTWNKCLKEEAFSSLQVKVLKTVQ
GEQGIEGAWSALCNRKVIADVGIVVDFSVRYT*

>g7667.t2

MRTGPPAQRANQQAPSRGLVRPTKVRWSKTSTPSGLSVSINLLDTFVSVERVVTLHHVRHANRRNARIKALYSPRL
DPTIAQHKKLSAPACAAAKRPHLSALPAFAPRAEACRTHWRIMAKSIHVVSKLDNAKHAVFHIDEDPPSLAASSVRV
QTWLVSLTYNNLTYARSGTPLHWWDTYPVPEASPTPFNSWEKWGIVPAWGYGRVLESTNDAIAPGSLLWGMWP
TSEHTVDLQLEAIEPSGHWLERSAQRSKLMTVYNSYEQVSESDAQTMRMTALCKPLWQGPHLLNTSVFSARRIHPF
GFGAPWSQEDADLSSAVVVSLSASSKTGRSFGWEMARNRDVSKHGPRALIQMTSVPNTLSEYDSSLPMRAAAYD
DTSAMAWAEHFKPSRVVIVDFGASDAVLESVRATASKLAPKVTVVAVGYEAKVYTQEEIAARMATASTKVPVNTSG
MRDRVIESQGALEFSKELDGTWNKCLKEEAFSSLQVKVLKTVQGEQGIEGAWSALCNRKVIADVGIVVDFSVRYT*

>g6735.t1

GVSDRRPGSQRGRHRGLRAEQAGELQGSQALHPERRASPAADRQDRQARLAGGRARGRLWI

>g6736.t1

VSDRRPGSQRGRHRGLRAEQAGELQGSQALHPERRASPAADRQDRQARLAGGRARGRLWI

>g4475.t1

GVQRQQRRPQIPRTDPAVAGRQQRRAECGTGRVLQERHGFPDQPDLYQHEIPRPETGHRGAL

>g4476.t1

VQRQQRRPQIPRTDPAVAGRQQRRAECGTGRVLQERHGFPDQPDLYQHEIPRPETGHRGAL

>g2059.t1

ATGSALAAPDHPALIHDKPGFFVPLDVDRIISQTDLNSQSGIVPAHLDYLDHAGREHELNYLVFGNGCYNQN

>g2060.t1

ATGSALAAPDHPALIHDKPGFFVPLDVDRIISQTDLNSQSGIVPAHLDYLDHAGREHELNYLVFGNGCYNQN

>g7536.t1

MAEQIDKPYTLQLSTKICDRLPRELRDRIYSLLDLDQETKRSKPIDDFIDDQCCEQFSQECLLWYYENVPQMLYRPFAY
QDIEHFMQLYPKVKKIPGLIIILDAGQPDFEFEALTDAMDKLKATGHFDDLNKDFKLRVYIDIAHHAMWRSMNPNV
EQAILSSQRILQYFSTRVEDALCFVRLMEAEDTESRDVGMDITESMEEPMDEILNSFGFLVIAGMECSPPRWIEQPYR
MCRRKKQLLQL*

>g7537.t1

MLYRPFAYQDIEHFMQLYPKVKKIPGLIIILDAGQPDFEFEALTDAMDKLKATGHFDDLNKDFKLRVYIDIAHHAMW
RSMNPNVEQAILSSQRILQYFSTRVEDALCFVRLMEAEDTESRDVGMDITESMEEPMDEILNSFGFLVIAGMECSPP
RYVEVEI*

>g2330.t1

NAGNPTVFVRAETLGLSGTETQAQVNGDGPLLARLEALRAAGAVAMGLAATAAQAKAERPHTPKLCLIAPPQTYRV
AGGKQVQAEELDVI

>g2331.t1

NAGNPTVFVRAETLGLSGTETQAQVNGDGPLLARLEALRAAGAVAMGLAATAAQAKAERPHTPKLCLIAPPQTYRV
AGGKQVQAEELDVI

>g9057.t1

MSSQDTIPDNTLYQEYRDNTVHQSSVPEGPTITDESFSGQKENDQLDESASCKTKEAVPRKATRQSEQHDAGIKRL
WDELNDLREQKKQLYIQEDNVLSRLYEALGYMAKTSQSRPSTSHKTLRWANNVDDHDPDEDIRRSNKRARNSS*

>g9862.t1

MSSQDTIPDNTLYQEYRDNSVHQSSVPEGPTITDKSFSGKKEIDQLDESASCKTKEAVPRKAARQSEQHDAGIKRLW
DELKDLREQKKRLSIQEDIVLSRLYEAVGYMGKTSQSSPPKSRKTLRWANSVDDRDSDEDIRRSNKRARNSS*

>g1425.t1

MYLCHLILLPSVLALPLLDPILQPITDSVWSTQGGLVETLLGSLTGTLGAKQSYDYVVVGGGTGGNTIAYRLAEAGFTV
AIVEAGGSYELGKPLVGPAPLGDIIGVGSNPADSIPTVDYGLRTVPQVGAGNREMHYAQGKCLGGSSGVNFMIHHR
PNRGALDAWAEAVGDESYSFDQFLPYFKKSFTFTPPNLETRLANATTAYVESDFTSSPSSPIQVTYPNWTPVWSTWA
AKGLEALGMNLTDKFNEGVLNGYHYAQTTIDPHAQVRSSSAEFVYAARDANMSDKLTVYLKSRVDKVRFNENKTAT
GVEVTGAGLLKYTISANKEVILSAGAVHTPQLLMLSGIGPAEHLAEHGIDVLADRPGVGQNLTDHALFGPAYEMTLD
TLNRITGDPIALTEAVAEYALTQTGPLTTNVAEFLAWERMPSSANLSQSTWEKLLEFPQDWPHIEYFPAAAYIGNFNIP
WLDQPKDGRMYASILAALAAPLSRGNISLASASPAVSPLINPNWLTHQGDVEVAVAMYKRTRDIFNTEVIRSIRAND
GEYWPGSEVETDSQILQNIRTSVMAVMHASCTARMGRVDDPNAVTDNLARVIGVQGLRVVDGSSLALLPPGHPQ
ALIYALAEKIADAIIKKLE*

>g1425.t2

LLPSVLALPLLDPILQPITDSVWSTQGGLVETLLGSLTGTLGAKQSYDYVVVGGGTGGNTIAYRLAEAGFTVAIVEAGG
SYELGKPLVGPAPLGDIIGVGSNPADSIPTVDYGLRTVPQVGAGNREMHYAQGKCLGGSSGVNFMIHHRPNRGALD
AWAEAVGDESYSFDQFLPYFKKSFTFTPPNLETRLANATTAYVESDFTSSPSSPIQVTYPNWTPVWSTWAAKGLEAL

GMNLTDKFNEGVLNGYHYAQTTIDPHAQVRSSSAEFVYAARDANMSDKLTVYLKSRVDKVRFNENKTATGVEVTG
AGLLKYTISANKEVILSAGAVHTPQLLMLSGIGPAEHLAEHGIDVLADRPGVGQNLTDHALFGPAYEMTLDTLNRITG
DPIALTEAVAEYALTQTGPLTTNVAEFLAWERMPSSANLSQSTWEKLLEFPQDWPHIEYFPAAAYIGNFNIPWLDQP
KDGRMYASILAALAAPLSRGNISLASASPAVSPLINPNWLTHQGDVEVAVAMYKRTRDIFNTEVIRSIRANDGEYWP
GSEVETDSQILQNIRTSVMAVMHASCTARMGRVDDPNAVTDNLARVIGVQGLRVVDGSSLALLPPGHPQALIYAL
AEKIADAIIKKLE

>g17.t1

MLTWDPSSKLRLQHHHCWFSAGPHLRRSQKEYQSQPQSHERKEVAHLVFSAYFRSCITELEMERLPHTLKDADELL
RLSDLLQEAALTIKDEWAKEDFSYQDESKDTARILPSHRLWNAERTIEAVTGAIVELVAEPHQRIQQILAEFMESRALFI
AAERKIPDLLHGAGPNGLDIQTISERTGIERRKLARILRTLCSIHIFREVADNRFANNRISGALVDNPGLRAYVQLFGLHI
YSSSDHFPRYLLGSTGHSYKVDETAFHRAMGTNKPLWEWMTENLLPSQVVSDGPGYPGVPDLSACSDPRDAPGV
VINRPELENFALAMVAGGKTSGAAHAFDFPWLELGEGIVVDVGGGVGGFPLQLLNVYPKLKFVVQDRPENVERGE
HKVYPKEAPDAVSSGRVTFQAHDFFQPNPVKNADVYWLRGILHDWSDDYCVAILKAIKTAMGPKSRILICDPVMNT
TFGCAEIPPAPSPLPANYGYHVRYCHTRDLALMATLNGIERTPTEFKALLESAGLRLRKFWEVRSMVGITEAGLNDAS
D*

>g17.t2

MERLPHTLKDADELLRLSDLLQEAALTIKDEWAKEDFSYQDESKDTARILPSHRLWNAERTIEAVTGAIVELVAEPHQ
RIQQILAEFMESRALFIAAERKIPDLLHGAGPNGLDIQTISERTGIERRKLARILRTLCSIHIFREVADNRFANNRISGALV
DNPGLRAYVQLFGLHIYSSSDHFPRYLLGSTGHSYKVDETAFHRAMGTNKPLWEWMTENLLPSQVVSDGPGYPGV
PDLSACSDPRDAPGVVINRPELENFALAMVAGGKTSGAAHAFDFPWLELGEGIVVDVGGGVGGFPLQLLNVYPKL
KFVVQDRPENVERGEHKVYPKEAPDAVSSGRVTFQAHDFFQPNPVKNADVYWLRGILHDWSDDYCVAILKAIKTA
MGPKSRILICDPVMNTTFGCAEIPPAPSPLPANYGYHVRYCHTRDLALMATLNGIERTPTEFKALLESAGLRLRKFWE
VRSMVGITEAGLNDASD*

>g4267.t1

PEQRDVVLRLRDASLGFGDRTLWSGLDLDVHAGEFVAVLGPNGSGKTSLLRTILGQQRLDAGEIAFEGHPVRRGDR
RIGYIPQQKLIP

>g4267.t2

LRLRDASLGFGDRTLWSGLDLDVHAGEFVAVLGPNGSGKTSLLRTILGQQRLDAGEIAFEGHPVRRGDRRIGYIPQQ
KLIP

>g10344.t1

MGHPKRARTTQPQSAHTTIIVKAQSAMQELPSLPPTLKKQPADTHWRYRGLLLDVYLQQFVEIFVRGSLQLPANYR
LKPFVVAG*

>g11734.t1

MTMCNQVYRFFDEAAASDIGISHGEILGGSSSYGASSSSVATLCFVLYSVSGGLTRAGGPWNRSTDE*

>g8860.t1

MGKDARRVALAALIVAGVKFGANPLLQNALATERVDLVDKMNMTIGMADRIPEGLLAVRTGGGELDAIQRAFRTS
*

>g11687.t1

MAENSPQSKKSQLSGDKPLTKHYDWADEDPDPLPVVREHITDRWDDIVGDKHDDLTDVLPYLREERIRTLRDSSPQ
EVSAESDNGLPEHGYLRIIDAMHRTFQRDHALHDDY*

>g7686.t1

MVPKPPPPRPILPEFHEERNEARLEPRGGQDEADENWYKERVQGTVSKKDIVRGFNTTYAAYEEAEFFAINWMIKN
NTYSLDFGKSENLGVRGELIDALEISSPFFQQRVYRAWRDHATRGFITTIRNQFKENPALSDLSSARKRQRYDLVKTN
GSPRKRSGPAQKTPLAFCKIVVQHHWGSLEKDVIFLRDLIDDGQEILNLDMNLNIEAVTYEKLVDRLEQSDIIRYVSGR
DKLYGIVNKVQEDVEDHSDFHWVLYDAHPTDGVYTFLVHRVEGYQRT*

>g7177.t1

MVPALASLSIDTTLAHPSYRYFRIELRRVVAADLALFLYFRYRRKSRQAMIKHRMASVPTSRRPRLRTTPSQYWNACL
REPPACAVITRPCLWTPDHDMLRGINRRCCNNASQTSSPTCSVQDWEL*

>g2358.t1

MPTSARIICKLAMMVQCLYPNCYEVERMSGQFRTPLRPANLRSIVSRHRAASSVPSVYPIVAWGSVVYTSGDLGDLF
THSIPLQAHYQYKPSLNEPYLIIALGLGTMKMFFI*

>g7059.t1

MIPYLHIHPIPPEQRRREFVGSSSAIYELPAPWQVCSRVASQSGGIHAGWLCKDPVLVLLRRPSGDAFDGRDRLW*

>g6488.t1

MNHRNPASSTITGRGFGVVAVSPSRESSSGQASKFEDLEHEAPWAFSSRCDRDIVSRMRKYRKRGRGRANNVVHL
ERSLMSFDWV*

>g1086.t1

MKIPVPMLAYEHVYRDVHLPQDHKRQSEKFQQRRMTVYKRFGCADRFYGAMGTPRLSFPINLPLAWPVIARCIFE
TARRQLSLVTHNPGSESQ*

>g1235.t1

MKVTSTARRIAFTSSLIKTLVEAEARAKAETITPKTAKKNELYCQVNEMAALTSPYLTGGDPVDAGRDARWSET*

>g9190.t1

MKDEMNLVVSGVVVEFVRPDTASLASIPLANPKTARSCSRRWKKSVGGPHTRKSYRPLTDNVRKNAIAFHNRMQR
FGHIAVNKG*

>g1531.t1

MFQSTGSYEGSTAKSSVTAGEAGKWLLPSAHMLDSMAQQVLSIFCVEHCPRLDDFPVNVQGVVIQFFQLRWHED
*

>g680.t1

MASITVNGNTIVPIGSQGEGWIVNANDQEKVESAPNAKDSNFILVQVDHILTVAEKGVLAHHHVEIQEYVAENTFLC
RYEPDDLQALRLLPFVVTADVYLPQLKTTISLKEMVESEHDQEDYEVDLILHETPNLTSEQLASYVAQAAEVALVDLEIL
ERKIRVTVHQDKLAALAALDSVNRIEEVRQYTTFNDQARAVLFDVGAIQNASIPYQGTGQIVCVADSGFDQGIATDT
GDIKVHPAFSDRVVQVIGMVPDTATPNDPVGHGTHVCASICGNGVYKNTADMVDVPIKGTAPNAKIIVQAMSQW
YPDLRAWGLKPPADTSILYSSAYQLGARIHNNSWGLKWSSTARQFGYAGGATAIDRFMCDNLDFCILVAAGNDARA
KNAGASQIGDNGAAKNCITVGATGTTRDNDGQRYTRGFKHGSDVTSVAIFSSRGPTLPARNANNETTVGRIKPDVV
APGVAILSAASRALSPKDHRRVANGESADPDWMFQSGTSQATPLVSGCVALLREALQGVGKDKISAALIKALLVNG
AVLHSSKDESGKTMIYDYAQGFGRVNVSTSLKMVRQLSFVDGWKGNIKCESEHPHAQDAPMLRVTSEKDKTWQS
PPLTIPSTGARMRLTVTMTYPDPHGALLQNDMNLIVRAGAGDEPIERHGNMADGDQGFDCENTVEKIIWDDVPG
PTAVIIVKAQAFAMTSVEQTFAVAWDLQSI*

>g1484.t1

MGAGDYPTREGRQQLTCDDIGTIPLPQPISATLHISELNNEIAFRIRAVESLSSSATKRADRVSAIHVSAQDRKLLWPD
VLLHATSVEKVLEREALNAASSSATVICSAHIQPPSPLVRYRYHPGTHIQSHTAIGGYTNLALSVAHTTSLTAAGSILVV*

>g1379.t1

MAGSWESQFQRYLNERSHSNERQWGYYVLFKYHLTKSGVETTGFASRPNNCHIEENSPSFDSAEAI*

>g3178.t1

MLCSEPPTVTCRERAIIELLRISRGLHPPSGRPWSMYILNLYGDTHLDNFPTPIVGLSSSIICVYLHVHVDPGQTGVTV
QQVSNIICSCWRQRSEWVVLIDSSRDIVA*

>g9375.t1

MEVFLVPVRARWQLKMELRQKTAVLVVFEWVLCGSLDWLERRACARSIQCDGGWLLWIVLSSHAYGTMGLYLAG
EPNDTGLYGGLGRTDDAPRVPRWVRSCPRLTNVWPRVVSSKEKKRQRGGGTEAGRQRGVDGVVHLRWSLIHGYS
SSLGPPERRRQPQLPKERT*

>g2003.t1

MFQQDIPRSLERKRLLAPASYRSLCLKARNGLLKTTITNQSGLMQIAKRNSLQFSLLKLPGKIRNKIYGYAVEYHPVGIH
GYFPIRR*

>g2436.t1

MRLTHRRQPTSILTNIEFQVYMPPEFTSYPTLPNHLHPEQEQRKSQQIIATKTILTLLLPFLLLFLLNLLLPIVFHLLLPIPP
LLPPPTLPNRPPQLE*

>g9250.t1

MSPATPPKTQSRHDFDRLWVAPAHSLRLSLLHRVMNFECGCREATESPEHLYPTIHHRVYLVYRWLAQSGPRTCRVP
DRLKVRARRLLRHESLLL*

>g4790.t1

MSSSVTQDSLGVDGDVRDSNSPSDPDSSRVDLDTDLVGISRITKDLNIVNRYRPKWTCKEAFRESYQNWRDGIFRSF
SLDHSQFRPKYTEELNDKQGFILIEVYHPEDTNKLLGFIQFTSNQDGYCVEDHNQIRRNPS*

>g6603.t1

MNDDLVYILGRTIHDSEGPTYAKNVASLSETDRMLTTEGLFEGHDEPMVVHKRKAKEDLKDEIDGTNIVAENKSAD
NKAKVMVRRGED*

>g1058.t1

MPWLADLQKRENIVAGCEAELEERQWTLRRQEDEVKDRESTVFRREIEVDEKERNVERTTASLVRKRRELKILEKSIK
RREEALREKMT*

>g2071.t1

MSHHTLNNFLHHPSISNLVELYEPIRYFSRTPRTAQEKPLLSDGHFRRSVYHETSFSSDFELRETSEAYFLEGEFPGISGG
TAIKVHWLDERTLRVKGIIHKTDLKTEWDAGPAENRSQDQPESPQYRRDVDDED*

>g510.t1

MSFAQACGACRRPKQSIIRPFSDIVCRSYPQRIGTVEAFFNGDGARYHRSRRSASVDLDEVGLVWYSNR*

>g5523.t1

MSIASKDDSTRYMQKSSARPTSAGTDPGNDVEVAHGALIVSPGWLGAASSRKILVQVQRGFAGPGPRRLANSIFH
MVGCRQCQAPVDCKSSAFESTLGEHRSMSPCTMWRGQASTQGRFTTAACNDKHESQDRNRDQASRTK*

>g3055.t1

MTCSLSLLTNTTRQNIAIATRPALYLIPSNAICLDTRAISIRQIREISSNKMLVVTLKTEEILAARILNS*

>g9068.t1

MRFLHVSVPSALFANAFAQTTIVSLQNNRVEQAGCAPKLHFCSNTSWRECSAPITSPDRCVATDRLNDVGSIAIELGL
CCRFYSDDNCNTLLMDQGGEETWYPGLEQVSDAFKADVGSYMCNNGTLSSDCPGAGAANVSISGGVATPTASQ*

>g662.t1

VGGGLARLLAAEGAKLTLADVTADRAQKLADELGAETVSADRIMQVEADVFSPNALGAILDSESIEKLNVGIVAGGA
NNQLARPHHGDMLIDRGILYAPDYV

>g11566.t1

MIVFYFLRLAALAVAVAINDPSLNFSPFCGTPSPVYDNGTQATGWAEYFEELEKTSGQNGQGVTDAWDAKPGPIP
WPRDSENKVVIPYCFTQEWDRKKHPPYH*

>g4043.t1

MLTVPVDTTWLTKDLLDYYTYSELRDVAKILNRSVSIIQNPQRLAVLRFLVQGLKAYYRSYRSYLLGEVLPISK*

>g3052.t1

MSEAPIATKNATRKVAKKADTRTNEEKAQAAGLKRAKWKQDMKVDEIKWHDSLAIHDGPGIAMYVGDYEEDVED
GGKTTGKLYLSFTGPADNVIFKYYEHDGKRIPGNIFDKSSPLEIRWAFPWEHTAENETDQRKAVRTLGKWLFLEKKPE
FFPT*

>g2691.t1

MLSNNTVVLAAIAALLIIGVVTIIRMHDLSSIAFEDKAHDGGRFPRDVTADLASARAQGRPDRGSHQHICQGLQAA
GGSVPVAFPVVSIYHKRQ*

>g2052.t1

MRFFNLSKESGVVTVMDFGSSTSACRVAVELRRHRYHSPVEICLNKSSSKAKAEEWKKLASNKRDDIRGTEHFFCNI
KTLPSNFTRATTSQCRVSS*

>g6450.t1

MIPLFVLWGAIVLLQAYATVGQKERYFLTVAPHRALLFTTSASGHSFNYRYQGPNGSIFTEGCLERNRHNLSQSWPR
EHKASARERTTEGNSCVSAKTASSSVSMPT*

>g6843.t1

MQPAFSVSPTSPSPTPTPSADCVVNVFKSPPRQFNCTFYGTTTTSTVYTDCGGCALKTKVLGVGLGCRVSTTVPGTA
TATVTACKKD*

>g7513.t1

MKGLGQRSVDQVYHGCGVVLAPHDCKLNSAIATHLRRALLCCRLMTLLSLSCIERARTSHDTIVNMM*

>g9437.t1

MSFELSQIGYSHSGPTWWGTDRVYTACITKFLIFLVFLIFFLASGALEWGLLLARMASKLSLLLHGLVPAG*

>g10964.t1

MGYATREQMKKTAEALSQGNDSNVNLANQRSSAIITPGVWELDSRTLPPRPPVELDANQIFVASSSSSQTHRSSSD
AHLDQISHTSNESLSAVSVFDSLYADPRDQSSRRTDSVVSPPPPYVSDVQRIPPADIELEKVDAQTHEFAHSPPAEAPS
PAYNNSTRHSEDYTAYRLGPGVVRGDDTRPRSELRVLTEEQVRWQRATRPVVPLRETQAETTSALTSLGGQDTAEVT
SIVPSPAPSPEIVSVERVPSLPHTLPPLDAFETVSFSITMDGSAPTDSTVSPSNHRGTNNQWHWDIQF*

>g9873.t1

MSCRPSCPCSALVWKLVLFFSTRPSAKSLLPFQACLSTIEASRPSWSSYYEYLCLRRNVSGAAKDDQREQNKSQ*

>g11187.t1

MFLTCLNKLRRRFLEKKPKKSASSSPTRAPAAPAPSRMGWEARYMNRGCPPLIFDKSLPPTPHASTSAFDLHPVPKK
KFPAAIHLEHKEKARAVSWASISTIERAQYIAQARDRANGDFVKEEGGEWRVPTVAGPSDLVKEALMWLRYEKESG
KMGKAVFEGSMRDDGL*

>g1480.t1

MADSRKINATNHSHPVHLDYTNQPNQHQHEQYDQSDRGNVDYNLTQNNYQAYASHTMDEEYESQYNTPTTSAP
PGPLRYEIGRYVPSDGYFDCDMYVLVKDAGLNKTNCFGCARPSSICKPGFPNCPPDIRCIFCEERFDTHPGSSPFCHK
MWAFTNFLQKYAGWDMSRGLPEGIPVKPNAKEAKYLIEQGFMKSTQYYDHNMVRKEFYQWLDPVRDPRIPVHN
FYGLTKNQKTKLLNEQNKEVDARGYRFRAHQKQKAAHGPVTLAKGKKRAYSDETHDNAMPPPTAPTKRQRQDDN
TSHYGYHTPARRDYVDSPPLPTTMQENTANRFGQCGMRDVQDDVWAQRDSLPLPTTQQAILVNASDIKDNKRA
QRDALPLPTTQQAMLMNAIDIKDNKQAQRDALPLPTTQQAMFTNTRTEQDYKQAQRDALPLPTTQQAMLTNTRI
ERDNKRALRDALPLPTTQQEPRSAPLNPIPNEPQSHAYSMTQEQCAQTSNNDSPDLQWSGMMIRGASNRVFSND
QPRGSRGRGRGRGRVNSRGHWNNGRGRGS*

>g9871.t1

MLSTIRTVLHAGLILQRSCRIAAQSYQMVGTPDRGYCSFDFDNPTGYFAYNGIDSGGDTQTGSGGTDPFHSFLIICNF
QFIGQADRVTQTAAYSNTAIVPTIIVPFTDVPVTSLVPGSNAQCTAVAGTTSDPFFQIQETYVLNSQTSVAPSTSSTVV
VVPTSVLTTTTTTTTLIQQTETTVTPNTVTSTVVTGTKTVLGKPTLTTKTVRITPWARTVKRVSITYTTTTLSCIPPSKVG
KSSPHQEGRAEVAARAVLARQDNTDIETATCFNPTSSTPTIVNTYILTSTTTSTIVEQALTTTTTISTVTAPAPTAIQNVQ
ATSTLVLTFTRYRWTVLPQSTVTSTISLVNTVTKLPKTPLKPCVRPTPTPPPGSGSGTPQCQNPTTIVGNGCKKVRCKK
*

>g12391.t1

MLRSDLYTTRTSEETASQGSDQHPHAYAGFVGSVAETNVLHLNHYHQIIQYIALELIPQQPESMPELLCL*

>g2853.t1

MTHYCTAFAFQSSQATRPFGIQRALWMTQNDGLSRPKHPSIITIIQRYRQIALSFTSTFSFPLKELFKRDGATYATGFIIS
PDHVAHMADLACAIFADDHFPTDMGALDWDFTSHGRELVIW*

>g9652.t1

MDFFVRLTSKISRQSRERGVPKPVDTADEVIINCGSIVCVPCLDLHMLAAPGETVCNLLRNTKSTIARYTHNISLRKRI
QRKEKENLLPASPLRSMAAFQPHLKSTMIQLKIDIREYSGLGQTRTAPKDSAVPEWGS*

>g10490.t1

MRRTETEVVPPAWGTFDLTTFANQHNEITPPVQHAGQSHLVSSGLSQMLHDSETNTFSIMLQYGIKGAWRKSFNL
HSNTPLNDCIHHKNRHCSVGIEPTRPHHRPLPGTTAHTLLVQRKLYPCTAKVEPEAHRNSTTK*

>g4091.t1

MSFNTQPHDSTIVWHKDALEKGNTYADKCAKREKDMRQAHEAHEAASILYHDSSNRPPTQQQQQPTATTSQHQT
TPSNHQGVQGASSGCP*

>g9878.t1

MNLLVLFAVFASIAYAQDGLCFIPDEGGPGECVPFRPVANRRCRDAPCTGQQNDCWVTGLNSARCS*

>g7303.t1

MLLTVWVVWKNGGPLLYLQGMPVRLRLLLNFLFPIETTPYIVSLGHTFGTENLDQRQMWSRLEPDLFG*

>g1481.t1

MAPNRRQRYTAFEFTRQESLDDTYQSDHQSDYESDSLNDAEQYETNKNELNTREPETRDAGLRAQDKSGKVVFIKK
GAKAIEKQATAEINPPRFMDPGPEHVWKMKIEEQVAIKDSWEKIDEEDAPVPKTWLPKRFQEYPEDAESEIDPEMA
DNPAYLRQIIRSMAKQYARCRAELKEEEAFRIRAEWERNMAQRKAKRYWSYSIHVTETVALPIYNTQQIEDVKLRKR
KKAVAKRERAVKLDRLAQQSEQADYEVAMEHLRNEVIETKKNLNALRDDVLENDAIKKCLRKDVMAELETDMMEE
RTRWQASQNQKRQELNREREEFDLERNASKEKKELATVQAVRSDLIDQGYERGLAVGKRFTAVEQYLRGYHFGERN
ANKKTSKQWLQQRYQEGMKAGQEEMSKKRDLEMEKFMEAFLKEVAHTRDDAIRRTEERLAEKFRVWMQERETA
IRRNSRAYGLYEGTITQIRRQNNVEGSEHEEDALATIIATTMADQVVTASYTNQDQKTCWRNTINEESPLWMNSKV
VEKSKFYEALHPKLTKEFQKATHRFELAKQEQEKQKQAHAARVENEWAARDATAQREEGEFVPVKSGAPAPYRHRL
LFIVEEPLEADEAQEMSDKDMVATFLKNIRDQ*

>g11001.t1

MSDYFGVGGLLLSCSLLGGSSFAAISIWGRKNIHPTPWQLTVTWNMVHILRLHANPDFASATIIKAPSPVTGGLMED
MPQTSILPWHPHLLC*

>g11663.t1

MVAVGTLATRIIAGVTVPDTPLINASIALAREALSDRPYNHVMRAWLNGQTILNKMPPEKRATVDEEAFAIGAILHD
MGWAFDTEYVSDDKRFEVDGANVARELVRKQGANWDKHREQLLWDAVALHTSPDIAAHKELEVALVSGGTFCEL
AGPEIAKQSWGDLITVTQDEWAAIAAEFPRDGMKDYLIDTTVRLCSMKPETTYNNFQGDFGEKYLEGYSREGKKIV
DLMDAFLP*

>g6526.t1

MYQMAPYRRQKRGFGWQYDNYPAFDIAANFGGVGGYGWDSSFDTHYYDLPGTVEQATRSHVEAQNEFEKAKEK
YEEAKKKYEECEKKGMSHPYELHVRRRLIIPHFQSKKQKKC*

>g1258.t1

MGDVTELLASLIDDPGSIRQAGVGLHATSSPSSFTAINRSTIANTAVRRNTSEPRRSNKKPKTASNLNKEHDGPDTAL
NPEPLHQIDAMTTTSSSKTRSAAVIPSSTSNVAPKRKPKTGRNSKGMAAKEDLSLSRESLESTVFETGSAAAAEEKTTI
EIHRSAKEAEWQLFQHGLGWHEGRDPYGLASQPYLWREELVPWWVCRMCKEEDEGTRQGGAWLSQG*

>g5515.t1

MPYMATRIRRRYLIQTLLEKHTWLCEVLFPAPMPEVSVRSNLQQTMAVFWKSPTQTITKLVKLTRMWFFTLLRYTYR
MRFFDGNTSYRSDLYEPNIQFYENRRYISEMTEREVCRLVCIPEEYGATFYRSCYKSAKRCPAWTKFCAGIQTFCKSVG
LLTSLKDPEQCLNTLLDNVSSLVLQQRWLQTASHLDETAILARSTERQVKELEKRVISAQLMNLNFTDDTKQQVEDLA
LECNELRSTVATLSCSQRCDANLSLELQRQRKSWEKELQCLRNRIEENETMNAHCVMTMAEQIEKSCINIGHPTTAS
SGTHNVELLVACRWLLERLPAFDSKQQGFGNRWRLFWQGQWKQYKHNTSRDDHPLRTLVCDEKYNKIGKGLYRTL
SSFLHEYGRLRIDPLDPDVQKVMDVISPIHYDSNGRIDIKAERKRWCR*

>g7568.t1

MNTSKNASTGCTPCEYLMGFNPSQGIDLADARSTLTKQDFELLRLHYREEAEEALNFARVLQKNTYDQSYQAIDLKP
GDYAALVLYHGY*

>g2174.t1

MTKANDQSFGASDNQPSGNSRGTVSLTTQNWGQNTASNVTDRTGAKQTFELKAPAAGDEGPRAAPTLHRLTSRS
VINQTRQIGKQQMTNGNADWDFDTTYQPEIDHSKVDLDWFLEPFGSHHAASSQPASDTPLARFLEDENDRFVSRL
MQLEQLCEECRRKRIVTCDHHAHRKLG*

>g11686.t1

MRHMAAYILMYEDKLLPEEEDALFPQLRAQPRLLRVAFNAAYNVKLELLRDGINTDQQDGTNGRLHETQLQRPTG
DLPLGLDNWFCSVTKGVKSAEDDFESVVCEGYFDPR*

>g4845.t1

MIHVVGDGSQLNYLMCPQMNIALSHCRRTREGATWRSNCRASPPHPLVRNLDNTTSTALFAAPLPASIIHILNSRRT
CCLEMMIGLLVGWKEEGAARQCNEKPRLEG*

>g10729.t1

MEPQNCIALSNRTDFTAYVDLLRNLTNGDFSLVKVCKKDVCEALWGFGNPDISGVGMVIGYLLESGICAFLVMISLW
LERTAKGHGHNAVRLLLANAARTFFDNAIFFTFAIQAASIVTLSRVDFGINAEGMGGFTMEIAWLVSSLTLLPLLPML
LRPDMFKEGRSAGVVVRRLSHQNTARMNDEDKKAEDSAHGLSGALFEAREGQRFLLFVICWAMGFCPFFSRMGG
TFGESRIGDAPNATITLTEWKTVESICFQGIHTLTSNEHNLITAFGILSYLLLSVVIISKIISSALENRGSERNWLTICHNKL
SETCSSYYARLGIEVVVAIASAVQFWAFFRLRQLQSDMTQVVGGNFSDGQWTFGQIVATVVFMPVAAELMFVWR
RRLLYLENQ*

>g6014.t1

MWRYDPRLYRTACFLSTYGGNKRKILDEERDGEYEGPSGKRRRVINISKARGKKTYIAPETKRNNNPYEPGFIPETSE
QADSDNGCCSVCGQRKTPFHINEEIDGPLAYLDLSMPSKPA*

>g84.t1

MWYLVQTPEKRSVVSRDFAAIRASSDARKAEEEALRLQNQEDQAQEARMQKERQAQADKVREEDRKARELRIAE
KARKKAKNKSSIPAPTVTPSKPSASKLPQNVPAMSSAPSVASRPGGVKSRAGGKLAPLYKPSPDQQSVLASPLAVSP
QVPVGSKPSAPSESELEIFERLSKKFGAQSSFSVLPSTLAANDERSPETPGLSAPVAPAEGSPESPKLSASTALRKTCQQ
SHHVQNSFIIW*

>g12340.t1

MKIVSMVSIAATKVGRAFRTHGPCPAKALASKRVRSFQPTYSSLHLFLPRSQKPKAWYELRAVREQFFLRDQYIACAS
TVSTLTAERSTALHSLRHTTITNI*

>g678.t1

MSEQNEKNLKAPEGEFQEWTEKTSESGKTFFTRDVDEKGNTTTKPADGPDHDVDSHGPSFDHEPCYWTPDTEGS
TSTDFANKTGITWYKLKNTIGATYQLTINTNSTYDYTFHTKPDSYGLDVYQTWTTHQVTYTTWEPTVTSVSGR*

>g1234.t1

MQGIQAFQFYSPTGIHVPPQYDRAFIITSTRFLRRAAKTEVLEESSTLLEDVEMQIGETVVDTDTANEDVEAPQVRDL
LEAIVQELEAIHGQYLKDMQFLKSYYGGSHLTADSLEVVMARLAATADLDTARLSIAQMFHQVVEEVAVAGTDFPRR
SRDFSKPLRQCMDSLDDDDDED*

>g7515.t1

MSAASRRRPPGSACVACRRMKMRCTQTTSGSCERCLRLGRRCVATEPYNPNATTEACAIPAVDLDLMQGPGFAQT
IPPDNVSERPAVPQRLLGSAIVSTECPSRSHGPQDEYEPSLRTRPYWSATDYVSAAEAADWILFFKERLVPTAPVLDP
NAYTNHERIVSDQPHLAACIVYVSSAYIAGYSGLRIAMQKGLNDFSRAMLEIQPATSAQQLTDMQVLIILYNFARPEA
ACSLPRQEVTSVFNLWSIKAICESYAFRIELFKTANDVLMRSQSGQLLQRSDQCVQFYLVWLWLFSNSHHVAMITGT
PPTISPDATIRASPMLLQDLKSEVTVQALVADAELCLIWYTLGNIDPGIKEWWCEFGFRGSQTSGDKRLTVDDIDSAL
EACQHRLWINPFQTLGSVMAFYPEFFFRYTRFCLYRFLIPQAETMMDRATYAEAVNRCIYAAVNLLNLPDEVGPYGR
DQLRYIPGFVCVLLSQCASFSLKTLGALPDIIMNTHIVVETIRRLAEFMLNLEQKRSVRSATIEAGRSILRQVEAQYESG
TGASTTATQTIDESEWLQNYFDQQLLDFNTDSSAFRTSFGLEQARDPPTLSFSN*

>g10468.t1

MIGIAEQSGRRAEILLDRDHDRALFSPAALVSKFVAQRPEKRPALIFALIFEYACMFVSVLFQGPGTVTEGLRLNCVGA
PDAAGVDHVRQAGALNVELKLSQTLSPAISCAVLKLVSLNCLYTTVLEYMSPVFCSYHRPQCVGERTENYS*

>g12418.t1

MSRVLHLGKAEGKPGEVYYPLQLRETQIPVPRDDEVLVRLKAAALNHRDLFVRHHQYPAISLENPMLSDGYGTVTQ
LGKAVPDESLLMQNVLLTPMRGWVSDPAGPEDTSKWSITGSSRLNNVGTAQDYVCVHWEEVVPAPKHLSAVEGA
ALPLVGLTAWRALTTKARVQSGQNILITGIGGGVALSALQFGAAMGANIYVTSGSQEKLDRARDLGAQGGAIYKAEK
WESDIRQQLPSSRPFIDAIIDGAGGDVVSKAVKLLKPGGVIVQYGMTVSPKMNWTMAAVLLNAELKGTTMGSRQ
EFGDMVKFVEDKGICPVLSRTVSGLGSLREIDNLFEDMEAGRQMGKLVIDIEDRSSSKI*

>g7608.t1

MKNEGRTQTWVDDNHRVLEHHASTTPQRHLLLAYDYHLRHSRSPTRDEYLPDLKPQRLADADILRIVASRTR*

>g1519.t1

LSQNPQALRLIKLFEQQRDLLASGVQAEEEGDVCPNCKAVLPEGQETCAICPETKEAPPSTWALFRLSRFAKPYKGSL
LLGFLLTLASTA

>g10692.t1

MASPLNDQATFERLMGSLSGQQRELFDKLRIRDMVLQPTVSHEERGRRNLEECQRNLDTSRQILEESQRNLQQLD
RREKNIQNGIQTPSLASTKTSHQQSYEIKFGLRHDAGGVIDRMTNEEIVRCLTRNNAPFDQIQACRVHATALVLMVS
NPEAEAAIHAHQHQIGPMLGIPKDDCHLLRPSFQVQIHNFRYRENGQFNRPNDFIATWSAQNEVHIVDARWTYKK
LVWTLDRLGDAQKLVKNVTVWLSGYQANATAFDKRSTPKQCHTCGKPGHLKSQCPDPSKPFCLRCGRQTKEHNA
WDGGCKGPECCVNCGRSHPAWSPQCQDPRMRRAREESRSYATKQVFWERFPATNSDPTTNAWLKAVQTNSRKR
QRADVDENFPSRATTSAHSDTKPKDRASALPSRAPSGSWNESHMDDSSMHATSSQVASTRSSQFSLEPTQPDIAIS
SMSSTESLAMPTETVSQPSPQPTPPRAMAGTTALTPSPQTPSRPRTRNPYSDRSTARMPQLPGPQIQMPNEFSQR
QMGPRPCIIAETPNAPSAPIKLLPSDTSKNLKIKTTRVRRLGGNNIHRARTEMPRDLHVAFHNFQRQTVNPAYD*

>g5529.t1

MMYSKTALLLLLISSVSFTLAAPVEAEPIDSVTYDPATFDTEMSFTTKLAERADKHLCWGWCQVNISGGPCNWYCPA
GIACKPQNNRGIGW*

>g2947.t1

MTQNTKTADQSLMYTIGRPSSNFLYNDRKDSLTNHSDIQNNVVVTNIIRTDPCNHEHLGKDDEIADRGSEEKNKGS
V*

>g5577.t1

MKTTIRPPLARPAPSVRQLARLTKPMTSETTTTNTSTTATSSHPPQTLTTIPTTTIHRHNTKSTRRQQHQ*

>g294.t1

MRTNGKSQKIHTEGGHETSLEETKSASASSKKDGVKSLHTNLHDLQAENVFSYDCPKCSQSIQATQVLENLDWHTA
LEIQESGE*

>g6586.t1

MATNTHATTKPAKPTPLDWVGRLEVPHTLRKTTLPRSFHLFALPFLRSSTSCSSTSPPPTSCTSHLSHFSQPIVPLPISF
NSENLPGREEREEGT*

>g355.t1

MDNSSHPILNDMAHSHVHKAACDLDYQFKVVDNGTLTSNTHLILVPKCYSRDALSYLWLLNTSPSRL*

>g7566.t1

MAYITLYVPSTTVPGERVFALWKVRAYIASELGPNILLGMDTLVPQGVLLDIAARAMVQLMCEGVTAPLIVEPKDDA
ARKTRKLTTLETVTIPLNSAMKVKVYASHPLSLNYDYMLEVPKDLPYAARPYAMLLNSRLMEAIVRNDTRNPVKIHR
RAPLGLAVPLDVDGVYALDQEDHAFAAMPDLDTMPRLETRFETKSDAGITVYGEPAKVSALLAVAERYPEIWIDRG
GFARIPEDQWMPINLRDGWQEQNMPSKIYPVGPEDRDIIDATFDKLADCNKMERATRMGFFPVPVFVVKRKIVEE
KNGIKAIKKKGRPVLDMRNINHWVIKDCYPLTT*

>g7668.t1

MAPAQSLLLLSGASVVLLWGTMLINGTLDGVSLAAKHGYFPDGRPLRQTFTGYPSVDGNLVVVVAFFDMLITARDV
HAPRWLFFEMCNVLGAINTWVLIESRRRGVRSFLLRHIVFFMFLWNMAGAAVVTPLFFCLLAKSAYTTRDCTIPLNE
ARGFPPTLVVNALFPVIMYAASWLGWSAHTQQSLVAWYHLNPMLMIVTVVLASRPGTSLTQFETPKRKSAPDEDA
PWIVASLVATGVLSAAVHVSVVLSALAASFTRNTDLGILRLYVPSPRSVFAQPRGSMAALVEGAHLFTQFDWIIVAAA
CFIITNHLLEKSAPRLTEKAKKTALLWNLLGTVALGPGAAGSFALAVRENRMRAMVPAVKTM*

>g10037.t1

MNEVSVSNSTVGEPGDFDDSVSPYNGTTVTSGGTRSTSMTHDTNDESHTPSGIEPMKKGRGRGRVTEMPLRRSN
RNKESNGGESAPMQLEEDQHNRVHPSNITPNIGSEGQQAQERTKEEQEEEAVKSQIQRECLETLPKSKRKKNPRVQ
GVEGNGSWF*

>g5930.t1

MDHNGLSETFFPKKRELHIAGFFHTHLPKQGDTGGGVCGEPRSLARQVPDANHHWGEKEDTLPDTSKASESGKAK
GKPEWDGKEQTSPKTQKHEIVAQTDLEANQDSEETLTQKATSGNR*

>g1940.t1

MSKGAKKQLITRNFVFSHRTLFNGFPMNIFAWQTLICMCRSFLSFSFLLSSITLNLQHYHAARRPGVRKTSSIL*

>g6585.t1

MCRLPRRSRAMSAAVSSRRRTLRRKLPPREKPPNGVPPDQSSTLGTGTRRKSPGSRCRCVGLKSHYSSLTIR*

>g4380.t1

MNPIRVFGHSPASPVDCADLCLRSGIPADAWAITAISCSSCALSLPLLLTLRAVDDTRLSFSVLARAVSRFLHLVRIACIN
ELSNVSESVDHHQLITSFLRLTETI*

>g8916.t1

MAEVDAFGYQITAKDTTADVKSFTESVVNEKLPKKKETLRQEIADRAAERSEGMFLWIKLLENEISPGQNAKQLRAA
VQEMPTGISEAYSRELEKIVQLAPNNKNMAVTILRWVLFAARPLLVKELAEALVVSDEDLDEYPDTDELPDAWEDGF
VDEDYVNEMILGRCGSLLQLRSKTPTTPLSDHTVHFVHFSVKEYISKLRNSLTGSAWASSLGLADVKTEEIRISNICLRY
LSLDVFEEIPQDTSVYPFLSYASWAWYFHSFHKRSTPPSQDILERTQKAFDPTASKWKVWTLLLEAELRERESDWEKL
SDVSSEFDEFASNSGEGSEPGQEVSTSMTLVENPIYYASLLGLVDVVKWLADQDLDCDCTRGRFGFPLQAAVARNQ
DGVVRYLLDRNVDVQQEGGLYGSTIIAAAALSSLELVKLLLNAGADATVVDDMKWTALHHAAKRGAAEIVRCLLDH
GAQVNSMTSERITAANLACRLGHKDVLSTLIAEGADIALVDVDDVSPLQQALENGHQDLALELIDRLSFTTTTHLRW
GPLQTAARAGYTSIIKKMIDKKIDVNMLDEYDWTALQLAAALGDTEAVQTLIGAGADITLAHADTSPPLHIAAGNNH
VAIIKLLAENGADVNQLGDGGTTALIVAVSNSCQDALEALLDMGASMKCMYSHEQQSLFDIAIEEGQDNITKALIAR
GCVGPRGLTATAEQGVTTLSDMNQHQYLPILSCQDDTKGLADRISIMTTPFPMCELNEALHVASARGSKSIVQILLA
RGASAKTQDINGRSALHYAVRHLKLDIADLLIEYGADPLAQDDIGSTPLDLAVCHGIRAAAFIRKHMGDLTLGIIRRPS
LLEAINSNQTPRLSSQTIWNLISGPWTGNYEYLSWQEGEKEAFSIEIPSVGMDESRPSTFLSEHEHDEIGDFQYLGFV
DQGGSIWFVKLYEKHGWLYRGRLDAQKRVIRGTWGSNRKLWFGGFELRQ*

>g11287.t1

MFRWDDIRKCLALSRDVERKYTSGDYENIGSVGGCALGMHNNIALCLCCAAGVGSRTHQSWGDMLVVMPLPQH
SWRLAQLSGCLPRSANSPVRK*

>g2074.t1

MEHHKPAVQHSEVDNVEHSSNSKENTTLRKTLDPTLPDPNNPLHWSNGEKYLTFGTICLFSFLSTANTSKFTVAVTAL
AEEFQKTPIETGYLVSFSVLALGLGNFIWVAMLRLCGRRPALLLAILCLGVFNCWSAFAKSYSSLMVATVLAGIAAGGG
EAPIPTVVADLFHVQQRGAMMMTFHVALSCGFFVGPAINAAIAEFVGWRWLCGWIAIAAAVNFAIGIFTIQETFYL
ATAQNDISADAENQMHARQDFSRRLSITSGYNKELQSRTVVWNMVSIAAYPSVLWAGLTIGTFVGWNIVVQVTAA
RVLVQAPYGFDLWDVGVFNFAGLIGALIAMFFGGRLIDIISSHWTARNNTVRLPEYRLPPLLIPSVIGPLGIAAFGLCLA
HKTHWIGPAVGHAMQGFGLTAASNVLVTYNVDLYPMLAGEVLVVVFLVRGVTGCLLSLYAYDWVLAEGLANTFGQ
MAAIQCFFALFAIVFYQYGGRIRTWTSKFGPSRHLSAY*

>g10695.t1

MPWLCSSTPVLLQCLISSARAPSIASSPPPRASGTGPTVFPYHVLSRYRVIALSHDIIVVSRKFQAAKPEASVKTPTFRS
RTTRCHTVALRQDVLIVMAIADTTQRPSSSIIERITATTSSQGQAPVPSRLRLRLRLRLQLPLSSYEYPWLSSGSIVQGN
HISCAPTFLRCHLPSRPSRRIDRPPPFKLTLANVLEHEDGADKEEFDSEQFLIEPALKLFCAQVRQADWPYASLYPVIEV
TLEAVHVKDDL

>g10691.t1

MSALRKSTDREIFRMQRQERIKALLEAQSLSFGDMNEMIHQLKKHQAVMEAGKYDDIDDYPLPLDTVGDEPVTKK
PKRSFDEYQDLERPSTILSTPSKPSTENMREQTELPNHEHARTSFQSVHDDLLPPSSSSQVDSASDYQSSTPGALLGS
DCSLSDPPSPTQTFPSGKPIYGSSPNHFRQPADGGVRGLMRSVHASSSPPPNAAGTLLDPPEGRDWRPLTMNPMI
SKVGKIKTGDFVGEHLATMLPEHCRTRSDWLRHLYHYGTIQLGHENVDAVPCTIIAEYEIAHGVGLGNNGERLCPPE
KFIFGFGQVHRNKTLEQAGPAYVEQMYKAKENRPWLREAVELSRSLKTTSFNPYRSNSQSHSSPSRQMYNHHKGSY
DSSGSRSSTYSSYNRAKSYTPQERSKYPTHFGAFAGDGSSFRKGPDDFVIAVSDLEHVSPDRLGMLKAPGARSTGES
PSHTSTSFAESPSPLERGQRTLPSAFTIAKRESGSTPAPAPVMDLSE*

>g7557.t1

MYQSQATHLLKLRIIHIRIYIKYKGASNDCMSLLNQMRNSTKSVKSPTYHYVASSKEAYSVILKDHPLDIMIMHQKAC
YNISIA*

>g11669.t1

MRVFLPLALTASVANAWRLSWYIGQSCQSQVLYSVSYPYGSGCNTDPDVANSKSLLASQDSPEDSEYEVALYTTDDC
SGLTTGIINEDSVCFPDTFGNVQSYRVQRIEERRKSKPRSPVEISRDFDAWIAKRPTGLAARSQETDLDLAPAKTHPNI
TALAALAANSTRADYFHTFLSVGTSAVTGSLTLTSLVTGCLGVDGAGPIGSLSCATSLAGTAISFIAAIYHGFKGYRDYKA
TLRQGVVNFNSRNDVRRRGIDSTLTMSQEDYMALMLHNVGLNGTHIGYHDRLDTNTTSPAFHFHGEDGQQFVW
TISPLEDGEIHHTIAFHHDAEIEKRQGYEGVRVNGGLDIQACQRRNADWSDLPYSAPAAYNYYVKDLRCLLEASDLY
NADYISSDVFDKSGNSAITIGMSTFRGDDSPNNVRRKRPCTDEDFYFSSSCIF*

>g555.t1

MGTGIDAKQISVKILHASFGLCEAFVDVHGQACWLEVFVPKAGYGLANSSPLTPRNNRIKTNEGFEAQYLLSWK*

>g409.t1

MATASATMSTNPIATTPSLHPPFTSRDRLRFALFLIIPVYAQLCILPSISSTLKHIYIVGTTGICIIQFSGDLVGGELINLGLV
GDIVYGLAISRFPHLSKSYSLTAWIIPEIEGWRGTVIQWAIMLRSLMYLLPENYGSKVEVILANYKDVHSRFGKYVGIFS
LFVTITFLESVLNKNLGPVDENNLKEYLETWHKVLVVWWILCFFIEYLCIVRLNYIHLCRYGIPVRFKLWPHKVVVKAS
QNHQGQIVIEEELQ*

>g5886.t1

MVRILSAVAVVLAIAFGAQASYTCQCNFSDGSHCCAATSAQGANVPCATVCKDAHRNSDNVACNAGGKWSSVSA
WNVQFRAPCAVQNFQ*

>g6620.t1

MEIPVTYLDQKSPYSCHLMPIVTLSSALRNFLINSRNPFANCLATCVINSLRLASRMSNQGSVAFQ*

>g8334.t1

MNALPELPKQLLATSIPAVHSMPFTGTLPKLQSDIISSLSADLRPIVYSDLNVPVGGHVWIGCRSDSTRADTSHVIFFA
DAKWPMRVHTPFEEFQVRRKINNLLSYRGFREILAELDNGDIIDCNSRVFIERNLLRINKAIRTELMDLVFGRNEVEV
SFLMHDEPNRILELYRVSHVRTINHVHRLPSALAHITITIRSSEREMKNGCQSLQFIAVNFANLRRLMYHIQSTSLYLTP
TEEAIALVDAFKAIAVTAKKLQMLHFAWNEEELTTARVLNNRR*

>g8379.t1

MLHLARSCRHWPRFKPVYRRNGSAVVESDVRAQASYHQRPKDLEPKRGWELLQKLDQDSKKIDNSTFARAFYPKD
GIVAAINYVEVPEAGAFVFIGKDPQDDPKYGKPGLDMGDTNFNQYYQKPLADADCEAGKTRFHWWHVDGTYWK
YDPPTFTMLRPIKFPNRRLEHADSGVG*

>g10087.t1

MGPILSNLAPTATRPMSATSFVRCVGSLDSQLAFNREVIMLKDHKRSDDPSQRYAVSSGDARKLRDTAGVQLRSFLLI
WIFSGSQSVQKTVSDAARKYMLPGQPAWC*

>g149.t1

MRTVKDYEASRGVAAVACVNDPKQDKCQFERQAQRTDGNKAPALRNALNIPLCCQKERMNDQWVLEGRRVHHL
LKISAILPAGAVSS*

>g8925.t1

MKSFNPYKDKFTLFFSGESYISLSTPKNFHLASLVLGRDRTYIVSETSFCVYWLAELVPGQRKDSYHQRHKSQIRNTVI
MGAQPSSEAHGAKCVECGVGLPRNNWSFEHLGRDGWVYVFYSIRHDDVQVSEALELAVASSTSDSASLKAGFSTG
RTMREVKVLDYIRNSQKDQGYTCLNCLKECCERGFAGWDRILKETNSLLRSHGIMVETARIQTIGGSFAFLSRG*

>g4271.t1

MLCVCELLTVLNSLHHFRGFIHKIPVSRLLILLHASSAYSPFVQVFGELPSFFLLRLEVTLPGWIH*

>g1410.t1

MLEQSITIHPCYKGLEDKAIRPTDCEISTDVTQSPTLPKGIPHKSKSATLERTLEAESVMGTTSNTPKPTKKTNRVRVSR
RKKNGEEKRLEPGKAPKTFIQVHKLEAQTAGYARFNERANFRRFVMSSRVLTHYLHAFADRTDKKNGKSIQVRQRD
VLEYIRDQSAEMESEEDPRIVHPVQNDGVHAVQRMALSCAQMRRDVENTAAGVRGFDQKEEDDVRAQFSQFTN
KKLPSPDCTKKDGADKEELWKRKHKKSA*

>g8610.t1

MTSVSILGTYKALVHRSKGRYKSISKVVPGLEVEWVASSQKYAWPGFFSFKWEKSNTRLLRARTGKRV*

>g2070.t1

MASENLYIVKRSKFDARNPTQPVYDIALPATFTNLNDAKQYAKRVLPKEGYDIKFFPVYDVKDLSLHWAHEDGVMVY
AEGPSGEVFRVEIDTVPNTAALRSTSSTSQVAELLYHVVQTLVDYNNDRSGSQRYSIVEGTYTSFKAARDRALQVLLD
GGVKKEDFVEYDEYLDTTGGLFGPDVVVRAVHDGGLNVLVSVISSVYP*

>g11661.t1

MSPNGPESIPHAMANFLVTKLTDATIWVQEKSNERREKARRLADKQVDREANDTRRRTLGNIEGEDRSLMGDWQ
AEEGRRCPQEREN*

>g1264.t1

MFTKDMSRILRIVSNFKKLPWTPKSETPKLADFKIQRLIDIMRPTAAEKNCCFGFEKFATVLRCAQLSVSTTVAFLRPV
R*

>g4038.t1

MEAPLGNGITPVFLFAHGSTMMLGEESEPAKVWEQVGNESLRRGVKRIVMMGAHWESVDDTIEVSMNANPKM
LPVGSVKDSRYIPYKVCPDLEGGQKVIQLLARAGFNVKAAPTFDWIHDTFLIIIRMFPHCIPLPTTIVSMNARYEPHFH
LKIGAALRPLRHEDTLIIGSGGSVHNLYRNHWADMMLYRDNFAQPVPPGTWALEFRQAVEDNITGNTGPELRRAIT
RMMKHPRYKEAHGTDDHWMASLFAAGAAGGVEDKGPNTMLAECWELVNMCNTQYQLGSWDAYRRI*

>g5008.t1

MRLRLPRPCERRSKSPSRVGQRSSFAQETTTMMTTTSRVTDTPTKVFLTILDPISMRIHRVVTATLALETTANGVRTT
ADRGVGVLG*

>g4268.t1

MPSISLTLRTARTFSRSTTKTYLNLSRSFSSSLRRNEINKVYPSAAAAIEDMSSDSTLLCGGFGLSGVPDTLIHQVKSTPK
ITGLTAVSNNAGVVGGGLGLLLESKQVKKMIASYVGENKVLEQMYLTGELELELTPQGTLAERCR

>g7306.t1

MARPALLASQNKKTEVGPQQMTVLKSAMPQLIPFFFVNETTVAFILLPSLIYVFSKYILPQRVRLFAARLFISKL*

>g6874.t1

MHAAFGPLQPLQLSTIEPAQHPQVLSRPPSTHHGSIACSRPLELYNVRRIRDRVQDGLQVALRAVDGGLAYWWEV
QDAWLRQ*

>g2662.t1

MALSLQHFTNRTAESDLARQPFRNDSTNIAALQSARAQQKAMEEAALRQEERFTAMTKEKEEDKLALRTALDEART
QLSESKNTARDMKERAEKAEASGRLHQQEMTVLETGMKAELQSLRYTINEANARYSYGGKENCLPRELDGVQVMI
VNMRSGLAIDAGKFYETKAHGHSFNPANENQIFRLNKVNNRDDSSSENFWTITLVKNDRKLYYPSGGTSVELMSGS
TSGRTCHWRIGAGGRHGTWLIKNAEWNTYIQLESSSRQLGVNMMSDNFSDDYNQHWIILPFGWNN*

>g9982.t1

MTCPRSRPEQSKQRRFAFPDCKMGVPTEGLQCRPTPFLPRPALACTERQQPHTGLTSPNLGPNSSILATIRTRKHRD
PEGWLSSIVDCEPLEK*

>g2660.t1

MDVVDIFPENPVEPSIKYAGGFHTAFTVPWATGRVAPCAGLNTQRVAKDTLFVPFSAFKDAHAAPLHYRECQITSIE
DQSNSDIADQSELGTFALSASVGGSFLGASGRGSYEKNVRDSSNKSNISIRAEHTCGQIVLSRAPELDPGAVRLLQFS
VDPINEFRRKYGDFYVAGYQVGAVNNTTIYGELADKSFFEAKRAELEIKALFLKKHKSINEESYSASNEGGLDVSAFDS
LTASYLKFTARTYKESLLAGEIVVNNKQLAMEIARRASKVLETDFLLGREGTVSLNMIDRLCDKGLITELLLVPFASLREY
QTLLARRNMLNRLL*

>g11226.t1

MTNLGFEKYTSQGGDVGSMLGRIPAQSKDACVSVHLNMMSGLDLSDTSNLNPFEKKAINRALDWLDSSNAHALE
LSH*

>g9972.t1

MNRTAVYAYSQASKHLGYNNIGQTITIDDENAFCPSSGEVPLNTNTPPTLKLESSSRFHAKEKTQSHIIIIDHQSRDSS
MYTIIY*

>g9155.t1

MSYQPPAWKHPGHIAHGGRFRGHADLSRQKALPRPMPTALEHAPSIAMLLVIGFWAATQSHVFVAMPLATTRSV
PRGGSNSYRLTC*

>g1579.t1

MLRLDFLATSNLRPRALVTCIQPTIGGIGYAAAIEDFRKGEEDEEDYPRWRANLLSIAYAADEHLVRCAHRIVSMLTKS
DLRYLGTARSV*

>g50.t1

MPRQAYAGLGRGELLLTFRKTIATQGIRRYGLMMKKKKKKKKKKKKKKKKKKKKKKKKKKKKKKKKKKKKKKKKKK
KKKKLSV*

>g7565.t1

MQGAYHNHALEDAQPGDLIDEDIPADRAPDFYEQATRAYHSDPEEPAFSSQDSYDYDNTSDDFDPRVWASQDAH
HEPCEQLPEPIDAFDENDEQDDGEEQQFVGTLAHVQPKAQTQGNYRVACFYCKRSFKSNNLLHKHVRKEHFIAGV
FTTTLLDSDKVEKPKISPQTKPINKEHIARAIRIVKSTAPPPIEGLTTLRNWYELRLNVKSSRTGPDLVVCPDTGAAITLA
DENFVKANFPNAKWEQLPTLI*

>g1184.t1

MPEVIPRLAVAGGRDRFRIVTFVTTKRGTRGTEPKRYVAACKKPPQPHCELDRPRRAVSSPRAYRTARDATHRNARA
TTLPYTVCPQYGSLE*

>g5888.t1

MRWAILAFALAAVAIATPVRPEGETLVPRALGCDSAYCGCWTGCGGNAGCQTSCFNGHCANTFPEGSSVPSC*

>g8885.t1

MPVRAFTRSGSKSARTLLKNKLGHHVHRVVNMDKLRGAHSFLPIFIKDCLELLRFMKVLKQGDDEYVYALMLANQL
AVVCSIAYTLATSNPSSPFRQHLLSRFDFASGNQAVEQMAMAAVVGCLEVVQSLEQIYGTELLWDRSYPFGHPLAA
AAGGNHIEVVDAIVADLEYNYRPDPSPAYRAAFISAIDVSLARQHCDMSLFLIDLYHHHFPGAFKHSLRRWLVKAAV
YPDDRVYLRLTSLAEDGDVRRHMAPFEMACRYGYMPYILRFFDHGILQLNEYFYIYEGLVLVRVDTPLSLAIGVGNW
HPVNILRSMGAF*

>g9049.t1

MSSKFLPIGVDESPRQLFARSLRPRLEAELQQQIVIYAQQFGDVSIYHARHIIREHADAFLQSLQKLPGPSSPSRPHAA
QNMASRAETGASRCRLYLLRPGQPLSYTTTSIFLGNLQSNLRSNPAAIFRQALSSPDLVQLLQRQHGTPSTYLPLRQV
AVCPIVDRIRKLGHAIRVGGAFKKTVRVAIKTLGHLFKDTRFGKQLERPIARFIFGLILERECANFGPTPFFVD*

>g2039.t1

MTRFTNVIIIPSDPQGNSTRYSGRGSSVAKGVNWAQTGQTSLKADTSNASQEFCSRAVGTQLQPCQVLVPLLVRVIH
FSTEDPEVAPAGCCGMYTHPMHTMLGRRQWCLVYHPYREDPTSDDDDLVMD*

>g5486.t1

MSLTSMRIWQPCLNDAPRHDQDHAQTEVASAQTTLQKLQKLSSTHTHFLSLSLQFITYDESLVCIS*

>g4505.t1

MKPQPDRLATRELETSTRSAWRDAKIHRDRLPREIRIHELLEERRAVDSVILATLSDCARISTSQAIEKVIYT*

>g2795.t1

MLAIGSLAEVSVYRGIASYHIVGRALGVVLKYRPSVFVMFSRCRPRVNPDAGVSRGIASLRDIVIGYFGLNTDAGMKL
RRGNMAQALVKHSVQARPGSLRKRKKTPPSRSLQLVAH*

>g2069.t1

MSGSSKADLAAVATSTILTADQLMRAIEASHEGKEDHAKKHIGYAAVGAAVAVGAMELLRRDELHKRRESDSSEEDI
TICEDSHHHHHHGHHCEEVVRVEPRHGRDTTPSGHKRRLAEEIVGAYALGQEIMGHKKHHVAHVVAEVIGAVAAL
KDTSDHVQGVE*

>g5782.t1

MFQVYPALPYPIFQLALNKSPTIAHPPSPQSPSPTAIRSDHSDISMPQSLVQEGKYGDPPRTVRELAAWFDGKTKGRL
KLEVLTWDGKIAKVPMNESGDIW*

>g8554.t1

MLSEEMARHVRATFSTFTTGYLTFALECTDFPYSARQRNVGATHFLRVMAAPVVSRRWLRKSGVGRSGRYTKSKTR
VMYHLYDERDLSSRPFHDFDALANLPNLVTNESLSIILPTTKHTLTWRNDCRM*

>g10730.t1

MSLFHLPLKGEVSLVYLLVLGFITSTGAQSLDAIPSSSTSSTTDLSFLTGTASQVDGQLSAFITITRSSQTTASAPTSRPTE
GQPGSGSDVSSGFPTGYTIGTAVGGAVGGLLIISILTFWFRAHKKKKQRLSIGIGSASSNKSVENEKSKDGENLSKKVS
QEGKLEGSAVGTGTHMKKLDAVRVLKSTNTM*

>g3642.t1

MKIRVAGTPRLPPQPPPPQSPSSQSPTSQSPPSLTCSPKSPSSPNIELSKPKGAMARLAMRKRMEAITIQDSDDKIRR
AGDTGGYASGERDLAHL*

>g2829.t1

MKSIYLTGPVDSGNEALSAEKPNRHHEYLATSGIPNSQTRLILTSEISQRCLNTGPRHSTSGLPSKA*

>g9060.t1

MAGSQIISADDIAHDDAQLEDKVKELTGEALTIESEVTQGSDFQEMDSTVGAVLVKKKGESGAFRVTGGETQNVVT
GPNESGVVIVVIKRGQTDGPKVMC*

>g1672.t1

MRGATPQSRASVQGNYCRSATQSCDQGRAQAHPPPVVWIVEGLPAPKQPKQPSREILDHSRRFIPNTVRPFPSSDS
SRAITHLHHSVCPCVPAKLHKLRVRETTRPVFLLPTQRRAKTKHAAPDLAVRLASRLQFLL*

>g8223.t2

MRSASPSSIISSSTTPLTRNSSPHSHRSSGSSIRQLLAVRRKPSMYEFEMEEEKHLFEQELDVLEPRPSMGRGTSEVHV
VGIFEVLDGKC*

>g5891.t1

MSGNELSDQRLRNRRLHAQDDGPPVPPKTFKTENISKAPLRRVAPRPLDLRNHSKPLPPVPQSERAAASKSEPLSKP
LPLPPKSVTLRSTLLKRRHAWFQQVAS*

>g8579.t1

MVPSKLLVTLTALGYVQLVASFAVPHGDSTSNFTSATISTLYEFPQPTFIENIAVRSNGEILFTTLGGLPELHGFSPNNHR
VVHHFSGYNSALAIHEICPDVFAVVTGNYSLQTFQPTPGSFAVWRVAFKGKRLSPLESKLSTPPEASFINGMTELLGG
DLLLSDSTLGVVFRLNVKSGSLSKVMDDPIMKFTPGASPEIGVNGIRTESILGKVLRILYCDLHKHSVPSPNRPLVRASF
RRLRTPGKW*

>g2631.t1

MSNNNSLIQDTGSRYKCILEYLPAIGHDSTDHENEICQRCKDNNPHSLASLLNNTGRPAGATDMQKCQDAVRKNSE
AQSEATNTCASRKWWFPNVKFSVAQKSLNEELIEVKREINKTVDDENAKKYFNYTPCDPVTTGRPGEICHKCVFFKT
NYMAWTDSREMNTIFEGPLYRVLKADVREKAGG*

>g4843.t1

MTRFRRPRLLTHWRLPGLAVVPDIGRSHESSRVITDDLGSNKRLDCRLDTRYHAADSVISKFLSYGVDVDLAFRLNRE
EADSSYLLVVCKPPTNLPNFPNFELRDPSLSAHLRRSRTRRTRGALPQNAYLTTSLTAVARLDSPTRLCSDPGFFIPQRP
TNKDP*

>g1588.t1

MVLLLILSVWNAITVVCVIVGYYVSWVIYSRTLHPLAKVPGPVWPAVSRTWLMYRMYQGGLETHMRAIHNSGYFH
YRGDCIALRRRLA*

>g220.t1

MPKFQPHRRRVDWLGNGKSAAPAPSSAKLAHFIALLHTIADRSTSSKRVAVVSSFLTLVSNLFPFAL*

>g9061.t1

MSRQYHARLSQHILLAELVSLLQPSTLSTFHTLLSLLALLQHPLISSFPRVFAIFAARLPVSCISIVLDLNDHVDAVEQHEL
CKSAG*

>g11622.t1

MDASTRQFFENIRLEPNWIDQHLMSNFSGLTTTGGWPVSPSGCPLTDIGNYLLRVLHVLATFSESLVMGDVISKHID
EQTGELEAEVDCHTANLLDDKHKALRNIKDNLSNTLQGKSILYPIDQCGSGPHGWRDCQAEWILFSITPSPVLVPSVP
IESPETHIVSPTSEDYFMATMSSEATDYRTYVEAPEGSAEASHDRASSNDWQDIHKLSPRNTEEQTTEVPSIEQLKAR
GKEILLLGHIGSCMSGFARIDQLERHKAKTRHTAPESLSSYTSSV*

>g7944.t1

MKKLKKGSRTHRKLSTRPERRGDHLPETPIQQDNGLLPTTSQQASDRFNALFKSISRSSHTAYQEIDVTKKKTFEETT
QSVQETHPEQNSINQLADRLTITAEGI*

>g12161.t1

MSSSTEKRRNSRQFDDTPTLATSPTTSATTSDKPAPLPISPLPLTSTPASSNLTSASPLSEVQKPGLYHTPTDIITATEKG
MISGSLCATMPSSSSTASSQKHDSEARDVAADAEEETAPVLTRAEEACVGEGRKGSLAGGEREGGRCVMFKGVDV*

>g11487.t1

MVTLIASSSHGACTIHFQDDNIYTPNIARYASGSEADAGEEVYGCENIHSSERKPKVRINPSIMDEARPYARQDKGGE
ACVG*

>g7974.t1

MKIFNLPLSGSPFGKRASFYGRLSATVENMRDKTKLGLRLLLLFVAKHEIEATNWKIIALWAWNREASRARASGFLPC
TVSD*

>g9998.t1

AAVGVVSVVVVVVEGTVKSSITGRSMPAIRSHVMICGDMLKMRYWDVEVGRRIFESMFKYFLRFEFHGSKPWLP
AVNHGSLQRNYTLLSTSSFTTTTQS

>g2062.t1

MPFNFDEYDAECSQMSLEELQREWNHYTRLITGSATSTTISSLAVPFTFGMSGIGVLIGGAGIHNARKKRDIIDRHLS
RYQEQHRTRKRDVMGSAMFSGAVGIATLGVGGWGADVLMAEGIEHGVVTMADQMAVKAVAHVAADGAGLAA
EHMHHNGLRDEEKKHT*

>g9905.t1

MRSLLLRPHNDESLLLYAFGRMREAEDDFMYIGRAFLSLLASAEGEYLDSVRTKILPHELSLESLVPCRNSFDADSGVD
LRPCSNHLDTRSFRSYLASTSDMSDWRDCPDKRFVPEYPASTSPATCYPAHDESDLPNHQSHMRNHMAISNLQVQ
DEQDTDSLLLSSQRRLRSLSPSLLSSNWDGSPVARTKFSFPEETTKTHWLHCYDQLNSLYGGSYEELCYIIRIGDSFFRE
DARDFITHFYRRWNSEDPWSRLPISNPISKRPLTERMLKSLHCAEAIEFDSIIDPVKLRIARMLLHHYFEQLCIKFKKDR
RLCNLSTGKGVASAAKDVVLKMIYGCREKTLALEIRKKQENSFTWHKRIGKRWSYVASHLGVGIALTCSPTLEAHIND
PKKLPDRGLIILLTHISYGYPGGLDVSRKIDPLVNSIMEGVMLPSPSQVEHMASDIEGYKQEDERVLVELEGDWQPLA
QPRCYGQ*

>g10690.t1

MSSPLWRPGVTRRPPKVPFQTATMVTNDGSRNVEALYRSTMELRKRHLETTQSHGISPGKTGNRTTRDTADRSGS
TVFPGVSHYTPEQRFVIVTPAPPIDSKDAM*

>g7226.t1

MDNKSISLLRYVFGVLSKLRCRANHSTQGSFRLLPSPRLENAHRACHLTRLSVHSCFNHVSAPSSGRDFAEYMHTTE
RVRASLAADIRMQWLIPLPKH*

>g8921.t1

MVGNASQRIWTGIQWFTTELLNYRQTKHDVSPEEFLAHCMSLKSKTLERKEAIQTGCWEGNDEAPVPKECTYRPS
DSLISTGGFGGRDEAPIPQPFALI*

>g4274.t1

MTWIWNSAVTICRTLSRILLPKHFNGKDDYDPTYRSFVFYSTLSPNESLSSFDRPSMESGMDEMEDSQRPHA*

>g2288.t1

MQLSPLFASAYAQHLEFHRPSSHIKCCIAALVLLGAFPYVQTGRTRSALLGVTSAARSCACYEKAYTPLIADLRSGYSDP
QRTYPRT*

>g8785.t1

VGKRLATFLRQEEEEEEEEEEEEEEEEEEEEEEEEEEEDSEKEENQPI

>g8260.t1

MAHRRGTRLVLAGILPSETAAGPPFNVVPRRLGPIITAREALSSAESRTLSLGSLLELPSDHCCWPRGASLFTATSNVCS
RLPRHMVLSMKGPLTAAYRCTGAVLIR*

>g5541.t1

MARVTFGGRDATGFPLYFNPNIFFTSDPSDPTPAIPTRDVPVYIFASVTNIGKETTKGASVKFWICSPSTVPVTETPAP
FAVSNVSLSPGETKEVLCVRPWIPEWTNNGHVCIVCEVSDLYDPAPAHPPTRWDLNDRHVAQHNVNVRFNEARRF
GILSSMSAFALPTKANIKVSIRQAPKDTFKPAMAKFGLKGRKNVSDSTQAGLLDRFEPGDTLPKEAPHDVEWKDSKP
GLQRPFHAFLKLPEDHGENSAAIYFVEQHDDKGKLVGGVAVILLGGKQPDVPRIQPAVPAIPVAMPYRPYATDPGSG

IMLPDGLFVSILGQQYINVETRNNGGAILDGLSLYIEGVADPAISAPIRITSPLGGRALGGASFKSIFSADFTNATPGATR
VSFIVQQRSGTTTTSVRVLKKIFVVGVGLDKATKTWQIKVPEGTLHYHARTVIAPKDFDKMGSTSPGCKCPDSGEDK
EITPIPIFPLSGTLFWVPSTPYAGTHGPLPFEDPVNKVLAGIVAGIITLIALIAWLKKLFDDDDDEGGADGSSGGGGGD
STIGGGVSGTFDETTGEITCCTGVQVITDNKFLGAITTLAIAAWVFTFMQDGKDLHEYGRDNTTPAVGELTIGELVEF
DIVPTDKASPGVAWGGDIKWKYERALNSGRTLTYGTTTHSENTHLLKSYRVSIDGKISPDKHYIHMRKKPLVIGAQFV
RLDGSLFKGSDLYVFAWLWSNRGQKISVELRDDGYGQLAGFDEDGAYLSAEEGRDILERVAPGGMLLRSKVTKTVS
DWERGVIPVDKIGGLRAIADEEAIPNTGSYVAVANSVQQYSGPTTWYVFVLAQDVNTVAEGMEPREAAKTVGGCL
LTTGYKLKWDGSPCEMDYDAVVEVF*

>g11772.t1

MATIARRVSRFQIDLDAIKSLISTFVPRHLLRYAHAERRPNCSVALMLRRHTEKAPSVTPFDPSQQSLQTCPTLTSMSA
FYVEMAPQFLAGDHLYQGSPCVAMHGLVPLKCS*

>g2391.t1

MTDNTTAAADKLGANKDTNAIPSQNPSVISSAGAIGKQFNPDGAIGQIGEKVGGPFSKDGVIGSQFDASKNGIAGH
VERAVDGPSNPAGSSK*

>g2253.t1

MQNYAVSGTRADVTPPLPLPKRSIVATLNIISFLRTAYTGDACGNLIISQWPGLTLLRVASRDVGMTAGLDVLGLNFG
RQ*

>g4521.t1

MSKRGFEPEKGALKRRLKRHQPIIQSPCSIPLASAISKPNPEEKKKKKTTTVYMPQIANRTHIHDATQHGLYLYAKYQI
PQQHRIYRHV*

>g7581.t1

MKNSFTDTGIESPSVSVAEWLWRVTQAKACLLWKTHSQEVSHRATCVGSNPTADSILFACFCVFSNSRTRCDTVCLK
QSSLFAHEDFRNVPSYLA*

>g7615.t1

MKLSIYSFAFLNSAIAYASGNPRLHEPRAARCALVTPLIPKLLLPTAQAFCSDLLKIKASTVTTTATTTRTITASTTTISVTA
SAVSTLVSTSLLTLTLPNTVPQVVTVTVSTQVEATAASTETLTFTEYTDAAAVARAAPTKPAQWAAFAGRELSSACSCIV
SAARTTITLLQTSTMKTTRTVQSTITVSSVAMATATKIVTITGLPSTDTVATTVTSTRTSTVVSNTVGYAVRQAPNRT*

>g4726.t1

MVAITFAAAASTGGAAFLPSLGAGLVACGTEATTAAVAVTAATTKAAVGVGAAVGQTAVTGAVSGAVNAAVSGAVG
TALGTAATAGTAQVLAGAAGTAASNAAVAAGAVAPGALSGAASALAVVGPAGWVMLGADGYSWDCWKPIVLDE
SLEPSQGISLRELCCHPNLRRITAFGDGFLAENIRGEKFRLSPTDVNGALAFHGTAM*

>g4672.t1

MRSSLIFLTLFALPIFGATLPRKDADGIPGYGVEDLSWEITLPDDSNVTLSGTIEQVAEQLDTLSPTWRAGLDNANAET
LEKRTYFDPQDVRCGNFPYLRNPFRITNGIKYLRGLSGRPANGPGPGNCGRVSCANTGAIWWCNDTPERKELASFG
SIADGAQVIVDRCTVRGIDVGVPPSPSVQVAGQAFHPTGWNVIVRKDSC*

>g2901.t1

MPRNLFRVQRARSATDVTWSDPQVSLFPPDVKVCCGRVLVGETLNMLHVPKFKENQREEIRWEDMQSWTSIPSG
SSKVETHPTY*

>g9870.t1

MSLRLAELQDGPSQWPDIADAARILWLASISPDIIAFRHELPTYYPNTSYSATELWTGLNASRRAVFVAIDENITIAFE
GSDRNELVKNTWANAKGPNWWDIPYAVYDGGNRVHSFFRDMWYGMRDETFRALSEAIRNIKEKGSTPRKITIAG
FSQGGGVSTMAFTDILEHIRCTYGSESAPSQAWAEDNNIGNLVQHLTFAAMAAGDQGFHTVLNNLYERHAIRAW
DFMHYQDWTRHAHDLAFRSWRGHRYILPDAVVRHHQAEFGQHGHSILGYLGVAEWMAMNGTDQVKSEYAY*

>g661.t1

MVRPGELVVRPAGNDADVQLLDAFRIEDRAQRIRAEHIGLDLHDAVGTDGLGAQFVGQLLRTVGSDVRQGQLRAL
GREQPGETAAD

>g7472.t1

MRLDTYNLRLDADGLQKAVVPVYAAMSLVSTTIHIDLTSEVENCAVCSTVLGEGGGKAGLVYVLKDCRCVICGLCVQ
FPGTPPFLPCQHTDHLDLGWQRPLRLYNLKCAICLDSDDATRIALICGTFTCYAEALANN*

>g6981.t1

MTVIAIIEADYNQCAILEKFLGSTFPRGEASVKHSRGLFRCKMPRRLEPHELENLMNSGVFEHYREE*

>g9956.t1

MLPIHIKGIVSVVFQIHHSSYRNTFALSLSTPPIVSIKFRLSPLSVQTHFRISIRSTWSNSTLSCSPPLASLFPQ*

>g9938.t1

MASAQPRHACHSPRLGAVCASDVSTPVTFPSTILLIIAYRTIAPYRFSTAAVLTTAALPSRLPPPPLPRVPAVTASSRKIG
TMSEQTTYK*

>g7708.t1

MDERISVRFDISRQTCSDWNRYHGYHTVYGNRLVRITYSLDRPPTNSPSSLRPQIYQCPNGQYYRHI*

>g11668.t1

MRILKKLTRIASNILNIVPSVIRNMQIPAYCPELCGVKFFDQRGVEVETKKVMAIESAPIVIEDAVEGDEGMVIISVAVA
DIAMVIDMSIPGSGGACQSVGKGDWACT*

>g10160.t1

MSIFRRSGNHGHVGIQGTTGHRRTRRHVGRIVRHHPIMRQYNMQSTRCRVRQKLEVSKTVIHRPLCDGSFHVACT
RTQRREHRLQ*

>g12249.t1

MWSDATISGSVTFSTLSDCILAGYISNTPGSLQQKHRVRSRYRGDSPEQSRGWAGERLRLRSCSILRSSASALA*

>g1738.t1

MLTKALDPTKFRTACQNLGLTHQNNTQLAISAVTTLPISDEQDLSDNKKDIAIQTVYLQDAYEDADRIAALTSAWCTT
QTGITPECKEA*

>g694.t1

MLSLQCFGTISRASTSSDLEPFENLHPYVIEKYRIPLEKFGFSKDTCMLPCAERLMVRSLHISLHQNFNYAITISKYAVAS
MESW*

>g2492.t1

MIQFVTTSTFHRPERCCTSVLSHTASYKCLMHGMLQGRGAFPSKPCAPHCSLQPGDMNSAPHRTPPCSSPPLPFHF
TMHTATRYTSRICREPSIMATVVQRQSCKPLCINGYSFAPYPRTDLL*

>g7313.t1

MPWKTPKSSQPQLQAAQFHIDPYSPHSSFQTSPSPALLPRSQYIHPPQPTATQYLPKAHLCVLPSRNHKGRTLREMP
ETHFAFLCVMEMWRLICLCLENAHRLMDQGFIIFIWAKPE*

>g5143.t1

MDGTGIGAKSFSKAELEKQLKLEEQRKAHGKTDRIKHEEELKEGEDRRKKGRVNRWINYALSSLAS*

>g12053.t1

MSHELSERSALMWHGYEVHPSSRHSDNPSNGRLERLFIDACQRNFDNRSARLQYRALETPKADLEHSHTKSCARLL
ALDTITASHTIPKHEH*

>g10384.t1

MPDQKDLSDNERSKTTVQASDTPADRQPEYERDDPVKMMVTVSGNTHHKTFHVARSQFVGSRWMYELKDPVT
NSSWEEGKLFPGNDLENDQ*

>g12051.t1

MERLYQWHYNRLFGPKASGDRKSVPEFWRDTILRAFAARQLRWTRKHRLQLNEDGTIENLPLKASWRWGPHREG
PEFNGLFQPQHLTLSVVVPRDEVHITVQSIIDNNRDNSPYILTPKDVNPAQISYAKLQQIIAATDNSPYSPSTHELSYLH
PDLATHFVEITNNETLVDALRHCQEYMNEMVAVFMIPRGTESKTKTFTTSTENIAVLPEPQKIEKKKETMLNNVQS*

>g8214.t1

MHHSAPLSILRHRHPSPPSRRSICHNGRNDTSLDTEALYISPSAVNSACHHSQSPERSPGNRPCCRWHRHRKIIAAV
APSAFVPQHTSPPNFSAADISSFTSTSIDHTIRRRLHPSLCVLTGRKTAPWRLQSFR*

>g7176.t1

MFSMHIFMESEEASSCDTIHAQYTYSRYLQATLSVCTTLQSILLHLSAKGYKKRRTLPYHWRGSDSIGKCGRRHKEIHI
LSAHHGPLRALCQFLRPLGSFSTPGRAHITKHHLQYIHHSNDRISQP*

>g7843.t1

MLLEASGVVQDYFGSLWRLYDKQADLLRAMVILYDDTMNTFERNLHELEKTNDSKYEVTNETIIQLCTGRVKDLKGR
LEALVPVMDGIHSTITRYIPTTPATITLAPPEQLVPAKTEQNVNISLKRRADSLHERTGSTLVERNAKVLSKRSMSAPPE
RRTNALDEQSASGPSEHQLHRGTPMCT*

>g4007.t1

MEICVPLRLWLRAVPTAVTRRLAGERAARSRASIELSLLELCLTISPVNSEDVGRRSMTCKQRFQTTSILQNIGQSNLY
DGMYGMHAQSSLA*

>g11454.t1

MYMRLYTRPSAAHTSSLIIQRTAFWRLLSLCHHHPIQIQIQIQITSSQFFLCFLLQSTDFLYKYGFHSTSDFGSKLLSFRYI
SPVYSGQRVVTAFTTTLLLSV*

>g3546.t1

MRSLFDKRPTMRNIIMVLNMKFADLVTAPAPHNRRNFLCQPPNHNSARWMKATLRISSLSAALQHRYRG*

>g10699.t1

MLVAQSDQGNPLLLCLSTELCNKFTSMFVLGQPVGVFASQIQRASRKCKHTYRRHDLDVSALLLIRRKISSEASALLCT
LTRFHLSSFEIRRGLQNCIASAPFVMSISNYTLQRSA*

>g947.t1

MRLRGRIQPPNTKHAADSALYRRLHVAHTKTRAHRHQRRNYPANDVRQRPLSWAEGGSIHFLCSTTADKKQSHAA
SVDIPKVYWCSLGVADLIAECDDYSWCLYYEELSSRWLAGMGILTEPHEKQCATATNSIRYA*

>g4166.t1

MLSLPEESDMAACGLKKGRNLMLHNHTILDTYLSPSSLLNSPYRHLLSSSSSSDPCNSQPYPTRTPATVHPENLQIMH
TRQPSTPLTIPQQTRKPPHVNRPSQNYSSNLLNLLRQHLPRASMA*

>g4459.t1

MLLSTSGQARVQRAAALAFDIGIDYYTTQQSLTQGYTCCTGSCQRHSRPDEQRGKGVLLTVPHRQLTRNEDPRRD*

>g3836.t1

MPTDQSLTALFLHGSVGPRQVDTRPRYSSESGLPGPDAQPDYIQEYTDGSGQSSTSNKHLNIPGWLEEDRSDPNLR
VGALRAIPPLC*

>g99.t1

MFAITQTLQSNRSTKCKSHSAFIRTSIPYIRPENKLRDMALDTQSVIAVIALLITCPPTIWLLYRIYTRRQPRHNHEISTLL
PTHASHPSIPMSRQNILHHHQRYHTWSLHTTIMLEDLATAFNHDSRGTRA*

>g4103.t1

MEYVLYLYSASRFSAHQSGVGPGQFTSARVDPLSEQPALAPSQPCQPSAINLTLLSAISISTGIFPPGSHFSTSYPLRLPL
EHPPPPSAYSRRDCASERANITRIVLRLASSHSFRNT*

>g1803.t1

MQDGDRREQGSSGGGGRFDYGYFSDAVLRDYAAKAVH

>g8861.t1

MQEYFGKPVLNISIARAAPPYDPALMLRTLYLSESDDSCMGVVTSQTCLIRAALVEIPVNLEWSKIAADSMPTPLSTF
NSEGDYANAPNGTRAGPLSTLGVMVGAFFTTYATLEDLNTVSSLNVMARLFWTRRQDSERCGHEYISPTQYVIYAM
QEFMAHAAIVAAGASGGRYDRHFTMDRTTPTTIYRADYWYLAGALVVMFAGLMLVVTLLWGWWSLDRWNITLS
PLETGKAFGAPLLKSERGSDEVDCILKDRGHVKVRYNDMGIVLNADGEEADAELQLLRMRSP*

>g1883.t1

MDYPPDTPDRPLPLSPRSPSSTLGSPPLALEFPALHSAAGDTGASDDFSMLESYRGSSDSSLNGTSDWDFRIADIYEQ
VVSLEDVSQIDQVIADLRDLKALYSSRNIEK*

>g11538.t1

MDSLSHVKSEKDKENACYHADRNTQVLSRRSQRPATNWRWVTGVFAVGTATTSSLVINAWHDASMVHGSAVEGY
SFGSVSTAMPRLSLLSYSWSDGSRQRQWARLSPLSHDPKPIRLIVACLVYSIAMNVFYHIHKSDKYQLGILACGAMSA
WGLGLASGENTPSVLTGLGPWAVMVSLVLSAAFHRARGHGEMVKTEL*

>g10133.t1

MLRCDFTSLSDQNPASTALHRDAPLCAPHAHRFFAGMRRVGQRLLRPPHVLFTTFSLTIFKRATSPRLQVKSRKRISS
RILSKPPNRSTRNWPRVPPTKAFLQSKNDMHSPLHSDSCYGFFPVDAALLRASLSTIFTTIGYQLIHTRRYLNTHSGSL
SAVNSNPSLSARYWLAPPPL*

>g234.t1

MTFRSSRPYGRSSCGVVFCPISNSAGSLHSLHRVRRVSLENNSSFPGTSRGEGYAVTLQYSCLLLFSTFEAKSATSTSSI
VIEATPIPIPARAPMLSPCGGFDAFDLEDGGPAVGEVEMALEDVRGVSVDDAVSVGVDVGVDEDVDKGVDEEAGS

DDSTVGANCNDDVEVEDTGSPATATSANKFAVSLQQLASLQHQLSSPHSLTGILSACHFLTSISPIIVRNNTTYYAFIAD
VSQTFHTLPCRIYTPLSPPQLHTFICSVKSFVILA*

>g11011.t1

MVAERRHKQRASHEKRNTSEDARPRLREIDNGGKDSSKIDDTFEWDECTIFYIVEDTLCYLNLLGGIRRTTIQPGVVA
NVLDIRRLLPHVEWRDPRELRN*

>g2352.t1

MGCGMSKPPSEYPPPMRQTKRNKRRHGGGGTYGGGHGAYAAGYGGGGGFFSGGDGGGGFSGGDGGGGGCG
GGDGGGGGGGGC*

>g3108.t1

MADDNATICLRRRSHLKPRTHFYEEDTKCGVRPRSVPRSFKSQPHNRHLDQMSCPARLYPLRKPQNHRVPIPRWKL
SLPDHIEHVFISYVGVEQHEDSDIATRAREEAVSTIQHWFEQSGAPEAVECFTTIDNYSRTGATVWVYYWSDRRRLD
RVVGSLNLKDIHQGVSTAGRPLIGLWHEFFIASVSRLETNYSGLDYLPGLARLPEASTEEHNLSTYWGAARDRIPDSA
HDLFPRAADLPRPDPIPNGVGQHLRGTNHANMVHIRSGQYWENCGKEEADSYEQKLEPTLKQGLRYLHENSAETG
ALSIQYLQNEDLTSNDSGSNKEPRKESCGAGFFANLEDLEHWAKSHASHLKIYGGALAHYKAFGDERRFRTWHEVC
VINEGDAMFEYVNCLPDTGVIASVPLTSVKI*

>g3109.t1

MSCPARLYPLRKPQNHRVPIPRWKLSLPDHIEHVFISYVGVEQHEDSDIATRAREEAVSTIQHWFEQSGAPEAVECFT
TIDNYSRTGATVWVYYWSDRRRLDRVVGSLNLKDIHQGVSTAGRPLIGLWHEFFIASVSRLETNYSGLDYLPGLARLP
EASTEEHNLSTYWGAARDRIPDSAHDLFPRAADLPRPDPIPNGVGQHLRGTNHANMVHIRSGQYWENCGKEEAD
SYEQKLEPTLKQGLRYLHENSAETGALSIQYLQNEDLTSNDSGSNKEPRKESCGAGFFANLEDLEHWAKSHASHLKIY
GGALAHYKAFGDERRFRTWHEVCVINEGDAMFEYVNCLPDTGVIASVPLTSVKI*

>g3140.t1

MSHSENSSEKSEPHQRTVSATKRFGGDSSHGGRNPFNAISPLTPGGLASPTTGGSSAFGLGSGAFASFGSAAKTPKT
PGTAFDFKAAAMSSPSPATPSDKKEKPVSKVVNSIRKESISTASIPENPSTPSAPPDFNAPWPLKYTWAVWYRPPTAK
NVDYEKSIVPLCKFSTAQEFWKVFSHLKRPSSLPSVSDYHVFKQGIRPVWEDDENKRGGKWIMRLKKGVADRYWE
DLLLALVGDQFLDAGEEFCGFVLSVRSGEDVFSIWTKSDGVKNVKIRDTIRRVLKLPEGTNIVWRSHDDSIAQRSAID
QARHEKSQHEKRRVVSTDESREKSTGGS*

>g3141.t1

MDNRENLWTRRSNTSKLSLSMSHSENSSEKSEPHQRTVSATKRFGGDSSHGGRNPFNAISPLTPGGLASPTTGGSS
AFGLGSGAFASFGSAAKTPKTPGTAFDFKAAAMSSPSPATPSDKKEKPVSKVVNSIRKESISTASIPENPSTPSAPPDF
NAPWPLKYTWAVWYRPPTAKNVDYEKSIVPLCKFSTAQEFWKVFSHLKRPSSLPSVSDYHVFKQGIRPVWEDDENK
RGGKWIMRLKKGVADRYWEDLLLALVGDQFLDAGEEFCGFVLSVRSGEDVFSIWTKSDGVKNVKIRDTIRRVLKLPE
GTNIVWRSHDDSIAQRSAIDQARHEKSQHEKRRVVSTDESREKSTGGS*

>g3146.t1

MVGSIPNDIFQFSTYSALSAGFNQGQPRTADLTSHGTDGIGVYEDGSLMILKDRQAYALTKDGKANSAPMNARLPL
ALVTVYQPSFRLKIPSISLEGFDDLVSSDDVGPAKGVNTLMPFKIAGRFDSIDFENGPSRSAIDGTIFGFVVPAWMKAI
SGPRIHAHFLDASEEVGGKVTDFTMAEEAVLSFAKCGRFHLGFPQGGEWEEMKL*

>g3147.t1

MDRLSDFSSLPNMVGSIPNDIFQFSTYSALSAGFNQGQPRTADLTSHGTDGIGVYEDGSLMILKDRQAYALTKDGKA
NSAPMNARLPLALVTVYQPSFRLKIPSISLEGFDDLVSSDDVGPAKGVNTLMPFKIAGRFDSIDFENGPSRSAIDGTIF
GFVVPAWMKAISGPRIHAHFLDASEEVGGKVTDFTMAEEAVLSFAKCGRFHLGFPQGGEWEEMKL*

>g3543.t1

MSTCVRKQAAIWKGKLKKGDVLVSNHPMFGGTHLPDITVITPAFSGDNIIFYVASRAHHADIGGILPGSMPPASKELY
QEGAAIKSEKLVSEGHFNEKRITELLLDEPGQYPGCSGTRCLSDNINDLKAQVAANQKGINLISTLMSDYGERVVKFY
MTNIQANAEQSVRALLKDVYKRFEGQDLSAEDFMDDGSPIRLKVSIDPEKGEAVFDFSGTGPEVYGNINAPEAVTYS
AIIYCLRCLISEDIPLNQGCLKPIHVLIPKKSFLSPSDNAAVVGGNVLTSQRVTDVVLKAFKACAASQGDCNNLTFGFG
GTTYGEGDKSGERKETKGFGYYETIAGGSGAGSTWNGTNGVHTHMTNTRITDSEVFERRYPVLLREFSLRQDSAGK
GMHVGGEGVIRDIEFRIPVQVSILSERRVYHPYGMEGGGDAACGLNIWVKKVDKKAIDQNSDSKRPTEPLDTTHAP
RTSDAVEAAQKKDAARAKEDVEYRYINMGAKNTAAMRPGERIIIHTPGGGGWGKEGEESRVQNKVDPRGSWKG
GSIAERQSTAEASA*

>g3544.t1

MATPKISERGIRIAIDRGGTFTDCVGNPGTGKMEDDILIKLLSVDPSNYDDAPLEGIRRLLSKFTGTEIPRGQPLDTSKI
ESIRMGTTVATNALLERKGEDIAMVVTKGFKDCLEIGNQSRPNIFALDIRKPEVLYKKVVEIDERVTLEDYAEDPERNQ
TQAKSIEEAGDKADLVKGLSGETVRILQRPQEEAIRRQLQEVYDGGLKSIAVCLMHGYTYPKHEALVGKIAKEIGFEHV
SLSHELMPMIKLVPRATSACADAYLTPAIRKYIDGFSKGFEGGLGTKSVKREEGSRGARCEFMQSDGGLVDVDIFSGL
KAILSGPAGGVVGYALTSYDPKTRIPVIGFDMGGTSTDVSRYGAGRYDHVFETTTAGVTIQSPQLDINTVAAGGGSRL
FWRNGLFVVGPESASAHPGPACYRKGGPLTITDANLFLGRLLPDFFPKIFGRNEDEGLDAQASEKLFKELAEQINKEV
AGGNKEKEMSLDDIANGFIKIANETMTRPIRSLTEARGHDTSKHRLATFGGAGGQHAVAIAEALGISQILIHRYSSVLS
AYGMALADVVDERQEPESTVWSDKKETREYLQNKMADLKSKSTSTLRDQGFDEDHVHFEEYLNLRYRGTESALMII
KPTKEEAQQEYDGDEWAFGKAFVKQHEQEFGFTLPDRDIIVDDVRARGIGKTFEGLEKSVDQQLKEIKPKDISGDTK
RYDTRKVFFEGGRQDTSVYKLEDLEVGDRLKGPAIIADGTQTIVVTPGATALVINTHVVINIGETEDQEKEVGTKEVDP
ILLSIFAHRFMAIAEQMGRALQKTSVSTNVKERLDYSCALFDENGGLVANAPHLPVHLGSMSTCVRKQAAIWKGKL
KKGDVLVSNHPMFGGTHLPDITVITPAFSGDNIIFYVASRAHHADIGGILPGSMPPASKELYQEGAAIKSEKLVSEGHF
NEKRITELLLDEPGQYPGCSGTRCLSDNINDLKAQVAANQKGINLISTLMSDYGERVVKFYMTNIQANAEQSVRALL
KDVYKRFEGQDLSAEDFMDDGSPIRLKVSIDPEKGEAVFDFSGTGPEVYGNINAPEAVTYSAIIYCLRCLISEDIPLNQ
GCLKPIHVLIPKKSFLSPSDNAAVVGGNVLTSQRVTDVVLKAFKACAASQGDCNNLTFGFGGTTYGEGDKSGERKET
KGFGYYETIAGGSGAGSTWNGTNGVHTHMTNTRITDSEVFERRYPVLLREFSLRQDSAGKGMHVGGEGVIRDIEF
RIPVQVSILSERRVYHPYGMEGGGDAACGLNIWVKKVDKKAIDQNSDSKRPTEPLDTTHAPRTSDAVEAAQKKDA
ARAKEDVEYRYINMGAKNTAAMRPGERIIIHTPGGGGWGKEGEESRVQNKVDPRGSWKGGSIAERQSTAEASA*

>g6150.t1

MIPLDAQQLKLYFHTEFHWMKVKVKVSRDERGELFITTLILAIEPSTLVISSTTVSRERKVSTLATSRKRSYEDYIAMSS
AGGNAEIIATKGDVILQLGKVGVGDGVRNVLVSSVVLSLASPVFATMFHGRFPEGQSLSPASPRTVPLSDDDPECIT
MICKIGHMQTSQLPTTLTAMDLFKLALVCNKYDCVGVVRAWAMIWIAALLETPAAQDFEKLFLATHLLDLSGEFSRV
SQSLIRDQSTTFSIVKAMAGQEFLPWTVYQCILLGQMAYRQEINKAFGSIVTGKERCSASQQSTCMFLHSLKKCGIW
PTHDYSVPAIKSKLENVGEAAFPADSKRCASYQNLCICGAVSCTKRNLVYELDKIYNAVQGLCLKCVRHKAFGESRAQ
CGAHLISGAMVYL*

>g6151.t1

MSSAGGNAEIIATKGDVILQLGKVGVGDGVRNVLVSSVVLSLASPVFATMFHGRFPEGQSLSPASPRTVPLSDDDPE
CITMICKIGHMQTSQLPTTLTAMDLFKLALVCNKYDCVGVVRAWAMIWIAALLETPAAQDFEKLFLATHLLDLSGEF
SRVSQSLIRDQSTTFSIVKAMAGQEFLPWTVYQCILLGQMAYRQEINKAFGSIVTGKERCSASQQSTCMFLHSLKKC
GIWPTHDYSVPAIKSKLENVGEAAFPADSKRCASYQNLCICGAVSCTKRNLVYELDKIYNAVQGLCLKCVRHKAFGES
RAQCGAHLISGAMVYL*

>g6251.t1

MAVIGMTCKLTDDSVVVNGDLKISFRRTVRVPETKESNWLPPDLGAFPLKPVSQHSKSLPPGMAVKGGVFFPMYQ
YEAMWINFSTVHEDMLPYMIKIYVGGVNVVSAQTAMESTASRSRRQVRLQTTPSNPKPASPLQDYVVVPGQKWI
DGIANGDGTVRQFVAAPLHSGLSVETQMTGKDAIGGIQIEITP*

>g6252.t1

MQIFLKQLDGSAKTFNVTNLEPIENFKLRIREATGIPTSIIRLIYAGKQLEDRRTFEDYRISKEATIHMVLRLRGAGSLVHI
STPEMNTAAGGLIKQVIHRDIHPPNWDTTKTTVFNAQILNSMLYQAVTGARPLVPSILHERYIHHGLPYYKMYEELS
NIYSNFDMVKSVGQITGKIECKVVTKAPRIVKIGKDHENVSPPGPPQVSHTMFKLEDALESYHIPQF*

>g6508.t1

MTSRRSLGGGRVLGSGRNLSPAAPLPQPAATAPAHRRNASLLSPSESSVSLSSQTSISPASTHETREDISSKVLVGPTEN
AAASASSRLVCPICNEEMVTLLQLNRHLDDNHQNLVVEEQDEVKNWFEAQMHKAKKFQPLAVLNQKLKGLDVFE
SNDAPMPPPISHSASSSTAHAAYEPAPVRRDPEEEVTRAHWQRPGYRDVCSDPVCARPITASQMALGGNQPAVNC
RQCGRLFCEEHTMYQMKLSRQAKHDPVRGIWCRVCETCYKSRDGYNDHHGVERDHFDDFTNIRRKRVDRERME
VSRLEKRLTKLTQLLADPPPIEETPTTGGWFSSLAGVKNQRKALEQSIITWEEDAKVSNCPFCQQEFATYTFRRSHCRL
CGRVVCSDPKTGCSSEVGLNVDASAQKTEKDATQMSIDVRMCKDCQHTLFARGDFERELADKPKDQRAYENLAQF
ERGIRLLLPRFHKLLQALQDPDKSPTPQQLTDATKVRKRLMDGFAQFDVAAKRIRDLPTDSPTQQKLQRAVYQQAYS
FLSLHMLPLRSLPKILKHAAPQGRSNGGSALASIKYNDRSAGSVVSSSEVSAMESEEKELRERLIVLEEQKFMVSEMV
ADATKHRRFDEVSSLAQNLEDLNKEIDQINGQLGQLDFASAYNGGMASPQPG*

>g6509.t1

MLFLTSVIRHLDDNHQNLVVEEQDEVKNWFEAQMHKAKKFQPLAVLNQKLKGLDVFESNDAPMPPPISHSASSST
AHAAYEPAPVRRDPEEEVTRAHWQRPGYRDVCSDPVCARPITASQMALGGNQPAVNCRQCGRLFCEEHTMYQM
KLSRQAKHDPVRGIWCRVCETCYKSRDGYNDHHGVERDHFDDFTNIRRKRVDRERMEVSRLEKRLTKLTQLLADPP
PIEETPTTGGWFSSLAGVKNQRKALEQSIITWEEDAKVSNCPFCQQEFATYTFRRSHCRLCGRVVCSDPKTGCSSEV
GLNVDASAQKTEKDATQMSIDVRMCKDCQHTLFARGDFERELADKPKDQRAYENLAQFERGIRLLLPRFHKLLQAL
QDPDKSPTPQQLTDATKVRKRLMDGFAQFDVAAKRIRDLPTDSPTQQKLQRAVYQQAYSFLSLHMLPLRSLPKILKH
AAPQGRSNGGSALASIKYNDRSAGSVVSSSEVSAMESEEKELRERLIVLEEQKFMVSEMVADATKHRRFDEVSSLAQ
NLEDLNKEIDQINGQLGQLDFASAYNGGMASPQPG*

>g7994.t1

MLTSRARIAARRIASPSIPRSLPIRTSPLCLNASASRLPQQQKLLRPLVSSRTYASGGRPQPPGGTHRMNLGGEPEKPA
LEQYGVDLTERARDGKLDPVIGRDGEIQRTIQILSRRTKNNPVLIGSAGTGKTAILEGLAQRIVKGDVPESIKDKRVISL
DLGSLIAGAKFRGDFEERLKSVLKEVEEANKGVILFVDELHTLLGLGKAEGSIDASNLLKPALSRGELQLCGATTLNEYR
QIEKDAALARRFQPILVGEPTVQDTISILRGIKERYEVHHGVRITDNALVAAASYSNRYITDRFLPDKAIDLVDEAASAL
RLQQESKPDAIQELDRQIMTIQIELESLRKETDIASKERRERLEQTLKQKQDEVKVLTEKWEKERAELDEIKNAQTNLE
RAKLELEQARREGNFAKAGELQYSRIPELEQKLPKDEEVTAGANRQGSLLHDSVTADDIASVVSRTTGIPLSKLNSGES
EKLIHMEDTLRQYVKGQDEALKSVANAIRLQRAGLSGENRPIASFMMLGPTGVGKTEVCKRLAEYLFSTPQAVIRFD
MSEFSEKHTVSRLIGSPAGYVGYEDAGQLTEAVRRKPYAVLLFDEWEKAHKDISTLLLQVLDEGFLTDAQGHKVDFR
NTIIVMTSNLGADIIVGDDVLHSVDKDSSEISPAVRSAVMDIVSATYPPEFLNRLDEFIVFRRLSREALRDIVDIRLKELQ
QRLDDRRIVLECPDEAKQWLCDRGYDPKFGARPLNRLIAREIGNSLADRIIRGEIRSGDTAKVAINEEGTGLTVAAA*

>g7995.t1

MHTTARRIASPSIPRSLPIRTSPLCLNASASRLPQQQKLLRPLVSSRTYASGGRPQPPGGTHRMNLGGEPEKPALEQY
GVDLTERARDGKLDPVIGRDGEIQRTIQILSRRTKNNPVLIGSAGTGKTAILEGLAQRIVKGDVPESIKDKRVISLDLGS
LIAGAKFRGDFEERLKSVLKEVEEANKGVILFVDELHTLLGLGKAEGSIDASNLLKPALSRGELQLCGATTLNEYRQIEK
DAALARRFQPILVGEPTVQDTISILRGIKERYEVHHGVRITDNALVAAASYSNRYITDRFLPDKAIDLVDEAASALRLQQ
ESKPDAIQELDRQIMTIQIELESLRKETDIASKERRERLEQTLKQKQDEVKVLTEKWEKERAELDEIKNAQTNLERAKLE
LEQARREGNFAKAGELQYSRIPELEQKLPKDEEVTAGANRQGSLLHDSVTADDIASVVSRTTGIPLSKLNSGESEKLIH
MEDTLRQYVKGQDEALKSVANAIRLQRAGLSGENRPIASFMMLGPTGVGKTEVCKRLAEYLFSTPQAVIRFDMSEF
SEKHTVSRLIGSPAGYVGYEDAGQLTEAVRRKPYAVLLFDEWEKAHKDISTLLLQVLDEGFLTDAQGHKVDFRNTIIV
MTSNLGADIIVGDDVLHSVDKDSSEISPAVRSAVMDIVSATYPPEFLNRLDEFIVFRRLSREALRDIVDIRLKELQQRLD
DRRIVLECPDEAKQWLCDRGYDPKFGARPLNRLIAREIGNSLADRIIRGEIRSGDTAKVAINEEGTGLTVAAA*

>g8019.t1

MRNGLYIVSELRRRHGVDTLSRESRTKIIHNGISGPLEMIEKTLVPIFTQGLISALLHGVDVDMEFAFLESTLLNHLHLQ
TFNNLREARFSYCEMRDASIILARDFAIKLFQGLEPSPSDVERQTHVLACHQTWFRALLAFEESSALISEEDRLAMITL
KIGYYTTYTASACVHDASQMSFDGYLDSFKTIIYHAKFLVNKTADTASPAQEHRMHSGAAANFTFDTCLVPALYYVAL
RCRHPPTRRAAIALLSRDLPREGLWDPDLYRIVAERIVEIEEKEVDGRGWPVERTRLWSASVTADVGEESGLRSDFLF
ARHVGSGVGNTWSEKEVPSVAELYVEVCNAT*

>g8020.t1

MSLNTVAGASQHSRKNNPKVKTGCKTCKKRRVKCDEDRPYCLKCTSTGRKCDGYSRMKHAYQTQVISFALGPSRM
PQHPVSSFSGSGNAQYLEFYHYHIGPMLSRRFDGDFWCGIVLQMAQAESSVRNAMIALTYLNQTQRDSLADTRH
DTSKRDVETSRQFGLHYNKAIRCLVARMSEASYAPETGLVTCLLFACIEFLRADKQNALLHMRNGLYIVSELRRRHGV
DTLSRESRTKIIHNGISGPLEMIEKTLVPIFTQGLISALLHGVDVDMEFAFLESTLLNHLHLQTFNNLREARFSYCEMRD
ASIILARDFAIKLFQGLEPSPSDVERQTHVLACHQTWFRALLAFEESSALISEEDRLAMITLKIGYYTTYTASACVHDAS
QMSFDGYLDSFKTIIYHAKFLVNKTADTASPAQEHRMHSGAAANFTFDTCLVPALYYVALRCRHPPTRRAAIALLSRD
LPREGLWDPDLYRIVAERIVEIEEKEVDGRGWPVERTRLWSASVTADVGEESGLRSDFLFARHVGSGVGNTWSEKE
VPSVAELYVEVCNAT*

>g8071.t1

MVLAQAIPSDSVAYALASSSAFAAEMASSLSAGNTPTWYQALPTDVQSLLPQIYPAAVQATPTPTPTPSSSAYVVKSS
SVVQSSSAKITPYPTGMNSTMVKPTMSATGGASISVTLSPSDTASSIPPPFEGAASRLSVGAGLGAALVLGMLAL*

>g8072.t1

MKANTMMMALGALAMGARAQDAVAPEVQQSVLMVLAQAIPSDSVAYALASSSAFAAEMASSLSAGNTPTWYQ
ALPTDVQSLLPQIYPAAVQATPTPTPTPSSSAYVVKSSSVVQSSSAKITPYPTGMNSTMVKPTMSATGGASISVTLSPS
DTASSIPPPFEGAASRLSVGAGLGAALVLGMLAL*

>g8174.t1

MRSRLSIASRPHLTMPPRIRIRPRALGACASHPTHPISLPAIATYASVAAASATTPAPPIQQTHKAAPPVLRYPPTQPPS
HKPPEVRKTQLHRQYQSLLKSSPLILIFQHNNVKAVEWMSIRRELAIALRKLDAERAKNGQETLADEVKMQVIQTNI
FASALRVVEFFHPEQSMLDHAQHPTDPRTPTSAQIPQTSNDAEDERFTHGLSRRAHEIANNRKLKLELEPLLSGPLA
VVAFPDVAPQYLKAVLSILSPSKPDFPAPSRKASPDYYEPAVQAGLQKLMLLGARVEGKVFDVDGTKWVGSIPGGRD
GLRAQLVQMLQGIGGSLTSALEGASKSLYFTVEGRRMDMEDKEKGPAGNDNQS*

>g8175.t1

MPPRIRIRPRALGACASHPTHPISLPAIATYASVAAASATTPAPPIQQTHKAAPPVLRYPPTQPPSHKPPEVRKTQLHR
QYQSLLKSSPLILIFQHNNVKAVEWMSIRRELAIALRKLDAERAKNGQETLADEVKMQVIQTNIFASALRVVEFFHPE
QSMLDHAQHPTDPRTPTSAQIPQTSNDAEDERFTHGLSRRAHEIANNRKLKLELEPLLSGPLAVVAFPDVAPQYLKA
VLSILSPSKPDFPAPSRKASPDYYEPAVQAGLQKLMLLGARVEGKVFDVDGTKWVGSIPGGRDGLRAQLVQMLQGI
GGSLTSALEGASKSLYFTVEGRRMDMEDKEKGPAGNDNQS*

>g8515.t1

MYYETILTVPQADALFTPPSSYSPQKPIARLDLGFDSPSDDVLDADIFNMTGTGSPQPYIKEEVDEMSFNNRFLGHN
GLDMNHQYANAQFSGHENANGINPSELNMSGSMMGNHFGTNSYTQGGAGIADDELAESLGTFDQQPGFNNFA
QEQVGQQDYYHNHTNNTSMNQIYSNTPDDLPIHSPFNHPGNFDFNQYQSVPQRMNFPGSMPSGSMRQRITMS
RTGSDSRTPMSPKTPAMAGLHLGTPDSGNFTQPIMTNMHRHQKSVSGQWDGTPGSAHSWIDSPTASPHTGGLH
HQQITDVLHSGKHNSLPAKVDIGQNADAKKRRRRESHNLVERRRRDNINERIHDLSRLVPQHRLEDEKIRKHINNNG
PLSPTMGASGMSPPQATSLLAGGQGRRAAGNITQGLPIEEKDKGPNKGDILNGAVSWTRDLMWMLSKKIQECDE
LAARLQQATGEEWVTEQTEDEKRMKTEILEALEKNGSGTFRYSRGPGSGLRVPKHTNLAGEPLSGVSPQSLSPGMQ
STGSGSGSNQPQYWTSLKEEDEYGMDMG*

>g8516.t1

MPKPTDALFTPPSSYSPQKPIARLDLGFDSPSDDVLDADIFNMTGTGSPQPYIKEEVDEMSFNNRFLGHNGLDMN
HQYANAQFSGHENANGINPSELNMSGSMMGNHFGTNSYTQGGAGIADDELAESLGTFDQQPGFNNFAQEQVG
QQDYYHNHTNNTSMNQIYSNTPDDLPIHSPFNHPGNFDFNQYQSVPQRMNFPGSMPSGSMRQRITMSRTGSD
SRTPMSPKTPAMAGLHLGTPDSGNFTQPIMTNMHRHQKSVSGQWDGTPGSAHSWIDSPTASPHTGGLHHQQIT
DVLHSGKHNSLPAKVDIGQNADAKKRRRRESHNLVERRRRDNINERIHDLSRLVPQHRLEDEKIRKHINNNGPLSPT
MGASGMSPPQATSLLAGGQGRRAAGNITQGLPIEEKDKGPNKGDILNGAVSWTRDLMWMLSKKIQECDELAARL
QQATGEEWVTEQTEDEKRMKTEILEALEKNGSGTFRYSRGPGSGLRVPKHTNLAGEPLSGVSPQSLSPGMQSTGS
GSGSNQPQYWTSLKEEDEYGMDMG*

>g9210.t1

MVKSLSWMHAHSSHMRRQWPRVGGGVCAVSVSEPLCQLQDNSYYTDLFRAGQTATTTMSSSFFTTPASQRKRK
RGEVAASAPKRRNTDAHARRSQREESISGSDESDGDGAPTFDDDQDDVDGGSTTSEDENETAAEKRLRLAERYLEN
IRNEVEDEVGFDAEQIDKDLIAERLKEDVAEGKGKIYRSIATELDFDHASHATGYSGLQKATTGCAVKLPYAWTISKDL
VVEKWEIADPKSYAPDPSRPSNLTPRTTPKRLLWRKGNKNKKGDRAFLGHTGEIVSIAVSDSGKYLATGDKHARLIIW
DADTLTPRHLFTRHRDAVLSLSFRRGTEQLFSGGADRAVIVWSAPESAYIETLVGHQDAVIGVAGGLELNQETCVSV
GARDRTARLWRVVEENQLVFHGGGTARQKGLDKLRKGRFGKNVDGDEKQDEQANGGTSDNDDDPPIAYAEGSID
CVGLLDAGLFVTASDNGALSLWSVNRKKPLFTYPLTHGRDPPLSPEQMSANDDAPTSVKPGPRLPRYVTALATVPFA
DLILTASWDGWIRAWKITADKKSIEPVGKVGRVPTVEDESMVNGDAKDGAILIRGIVNGLAVQERGDRGKDGLCIV
AAVGKEPRLARWMSGKVKNGIYVFEVPRKGLTNGMADEDEEDEEEEA*

>g9211.t1

MSSSFFTTPASQRKRKRGEVAASAPKRRNTDAHARRSQREESISGSDESDGDGAPTFDDDQDDVDGGSTTSEDEN
ETAAEKRLRLAERYLENIRNEVEDEVGFDAEQIDKDLIAERLKEDVAEGKGKIYRSIATELDFDHASHATGYSGLQKATT
GCAVKLPYAWTISKDLVVEKWEIADPKSYAPDPSRPSNLTPRTTPKRLLWRKGNKNKKGDRAFLGHTGEIVSIAVSDS
GKYLATGDKHARLIIWDADTLTPRHLFTRHRDAVLSLSFRRGTEQLFSGGADRAVIVWSAPESAYIETLVGHQDAVIG
VAGGLELNQETCVSVGARDRTARLWRVVEENQLVFHGGGTARQKGLDKLRKGRFGKNVDGDEKQDEQANGGTS
DNDDDPPIAYAEGSIDCVGLLDAGLFVTASDNGALSLWSVNRKKPLFTYPLTHGRDPPLSPEQMSANDDAPTSVKP
GPRLPRYVTALATVPFADLILTASWDGWIRAWKITADKKSIEPVGKVGRVPTVEDESMVNGDAKDGAILIRGIVNGL
AVQERGDRGKDGLCIVAAVGKEPRLARWMSGKVKNGIYVFEVPRKGLTNGMADEDEEDEEEEA*

>g9257.t1

MSSVGAWQFAPLVTEFPTNVLPNLGFWSATNGSWEYQIQVAWPLNWTSQEESSTVETMYVLDGNALGMTATEA
FRRRRPVEFNMPDCIVVSIGYPETIEDSPYSTQRSYDFQPPVCDGCPLPTVPGVQSNANEFIEFIDTALRPWVQQTAF
PKAEFDRDALYGHSFGGLFVLYALTVRPDLFDTFLSASPALFFNNDYVFNNTDFLAPLTSPSANNTSNATKPAFQISYG
ALEQNLVQRRTETDEEFAFRQSILVPQRMTDLSQKLYSTLEDSPALRDISIHEYPFSDHAAVGAAALADGIDYFLDW*

>g9258.t1

MTARDITAAGIQHYQFQSSYKASLSFTSTLDSTLHSPVQEVVDTICERCRAYCSMITERVLMLGGIGSVVSRGRKECR
GGAEKCIEEVGAYGESVQDEEAAKGVTVERIAVEFSLWESCLLDPGAESGVDELDKLIGIGLHARHCWQWAAVADR
RLEVVASLGASGMLNSTGRRRRKASVAVMPSALPSRTYLVGVELSSCDVQFNGHAT*

>g9293.t1

MKHLFRNPRLFPHLEAWSGQLPGITSGFFLWNSGTEPQKSSVGLLRAIIYETLQDLIFGPLEEDGIIVQNLFSDRWNA
FLSYAGGLHEFNFPELKAAFEGLVSDVKKKFFFMIDGLDETEDYPKELMDLVFSTARRDNVKFLLSARSSPAFQSAFEN
KPRMMLEEYTKDDIHSYLATTFNMETKLQALRGKMDGEEESTIVRILAEKSSGVFLWAQLATKFLLENLEPDSDFLILK
DRADALPYILDDLLAHILSKLAPEDIEVINKLHNLLSHQNTCPAILPFSFAYTAETPATLAADVRPLTAVEISKRVEDMRIL
TQQRCNGLLSIFDTTPSDQHASFESLRISYAHRAIRDYFLAYPGLLQTSLSNVSEENNWNSAQQWANAHLWLLKTLS
PPTTKRSSNGNDNSRASTLRVWTPLSSALESSLELHLATNKFPLTYIDAALTTAVFLALRSETGHDLPQYASSSHAASTS

TTPSASTTPTTLTTPLDLAVLLNLTAYIALKAKTTDRREVRHALDYSRGMRKRMGVGGEEVWLSGWGREGLRKEFEK
GRQDGEALLEYYAKAVRFGVSKPELEAPEWV*

>g9505.t1

MAPFFANQSCDPFQPQDRPCELGNYVRYAVDASGPTDVQKAIAFAASQNIRLVIRNTGHDYLGRSTGAGSLAVWT
HHLKDITHVPAYKGSGYNGPAFKIGAGVQGFELMAASRDKGLVVVGGECPTVGVAGGYTQGGGHSAISTSFGLAA
DNVLNWEVVTANGKLVNANPKENSDLYWALNGGGGSTYGVVVGMTVKAHKEAVFGGASLSFFTTDNAQDVFYD
AIQAFHEELPAMVDAGAMVVHYFTSSFFMISPLNAYNKTEVEVKAMLAPFVARLDSKGVNYTAEYSEFDTYYEHYD
KYFGPLPLGNIQVGIAQYGTRLIPRSVVSNITETWKAVVEKGVTWIGVGTDVKSFGSQQTTSVHPAWRKAIVHATLTL
PWNFTAPWSEAFAVQEKMTNEIVPLVEAATPGSGSYVNEADFRQPNFADTFWGENYGKLLNIKKKWDPSGLFYAT
VGVGSEAWTVQTDGRMCRA*

>g9506.t1

MRSFVFILTLLSVVSATNPTSNRCKAFPGDKSWPSQSDWNSLNKTVGGRLVATVPLGAPCHGSTFDNATCESLKSQ
WQYEKIHYESSSSVMAPFFANQSCDPFQPQDRPCELGNYVRYAVDASGPTDVQKAIAFAASQNIRLVIRNTGHDYL
GRSTGAGSLAVWTHHLKDITHVPAYKGSGYNGPAFKIGAGVQGFELMAASRDKGLVVVGGECPTVGVAGGYTQG
GGHSAISTSFGLAADNVLNWEVVTANGKLVNANPKENSDLYWALNGGGGSTYGVVVGMTVKAHKEAVFGGASLS
FFTTDNAQDVFYDAIQAFHEELPAMVDAGAMVVHYFTSSFFMISPLNAYNKTEVEVKAMLAPFVARLDSKGVNYT
AEYSEFDTYYEHYDKYFGPLPLGNIQVGIAQYGTRLIPRSVVSNITETWKAVVEKGVTWIGVGTDVKSFGSQQTTSV
HPAWRKAIVHATLTLPWNFTAPWSEAFAVQEKMTNEIVPLVEAATPGSGSYVNEADFRQPNFADTFWGENYGKLL
NIKKKWDPSGLFYATVGVGSEAWTVQTDGRMCRA*

>g9569.t1

MTELQTPLKSILNFRDVSEFVNQTTGTRHLRNGLLFRSARPDEASFRDRQRLLKEFGVKSIIDLRTKTEHIEQAQKHDT
RIKASAAIPQSNDDVAEPLKIPGITYHEINFNGSAFSRMLLSKLTWLEFFRLAGLMIVGYRKDAIKILAPHMEDMGLV
GLAEQSLDVCTREVRQVFDVLGEEQNWPVLVHCTQGKDRTGLIVMLVLLLLGVDQKTIDDDYRLSEPELEPEKEGRL
KEMASIGLTEQFAICPPDLVSSIHSYLLEKYGSVEGYLEKAGVAKEQVDFVRGKLLVNVS*

>g9570.t1

LASFRDRQRLLKEFGVKSIIDLRTKYHIEQAQKHDTRIKASAAIPQSNDDVAEPLKIPGITYHEINFNGSAFSRMLLSKLT
WLEFFRLAGLMIVGYRKDAIKILAPHMEDMGLVGLAEQSLDVCTREVRQVFDVLGEEQNWPVLVHCTQGKDRTGL
IVMLVLLLLGVDQKTIDDDYRLSEPELEPEKEGRLKEMASIGLTEQFAICPPDLVSSIHSYLLEKYGSVEGYLEKAGVAKE
QVDFVRGKLLVNVS

>g9619.t1

MSLVNFHSLHQPFSYTTLARAATLPTKSEVSIAKDRLQQLTGHLMRSHTTWSKIVEKFVGKDEKTGSEPTKANMPEE
ATKEPAADMIKSPGMTTIDKVVEDKKNQLPPWPSPVPTPHKDFVHSNPPEPPEYRKFTVFTAGSIEMGDAVNWQP
LMATMLNHLPITVCNPRKGSWDQSIKQQAKDELFKQQVVWELGALEQADVICFFFDTETKSPVSLLELGVWAASD
KVVVCCGDAYWKSGNVHLTCERYGVPCVKNFTELVPLVEEMLKKKGMELDNKGDLIGENVHVPKEKPKKKTQLEAE
KAELQKQVDDLLAKLAAQPKM*

>g9620.t1

MRSHTTWSKIVEKFVGKDEKTGSEPTKANMPEEATKEPAADMIKSPGMTTIDKVVEDKKNQLPPWPSPVPTPHKD
FVHSNPPEPPEYRKFTVFTAGSIEMGDAVNWQPLMATMLNHLPITVCNPRKGSWDQSIKQQAKDELFKQQVVWE
LGALEQADVICFFFDTETKSPVSLLELGVWAASDKVVVCCGDAYWKSGNVHLTCERYGVPCVKNFTELVPLVEEMLK
KKGMELDNKGDLIGENVHVPKEKPKKKTQLEAEKAELQKQVDDLLAKLAAQPKM*

>g9623.t1

MVLREDQEKATYTQKVDFAKLRSELMTADSTESSLTRASHERLTNELAKLNSRLRDEIQRTQASVRLDLNLEKGRIREE
ANVQELKLKETETRIEQETAQLRERLEAVKFSTLQWLMGVCTGTAALMLGVWRLLM*

>g9624.t1

MAAPARQATAIPRFLLPQISWPARVSRPAALAALEQGRHRSCIASESRAAFPKSLPLQARTRRTLWPKTSPQYASGH
ALALEFRRNFSATAQHSKDHHFDTLKFVQRLKEEGFTEEQAEGMMRVLGDVIEESIQNLTRTMVLREDQEKATYTQ
KVDFAKLRSELMTADSTESSLTRASHERLTNELAKLNSRLRDEIQRTQASVRLDLNLEKGRIREEANVQELKLKETETRI
EQETAQLRERLEAVKFSTLQWLMGVCTGTAALMLGVWRLLM*

>g9708.t1

MTDESSDEYVRKTFASRPDIVDNFLALPVPIWKADILRYLLLWDQGGIWFDLDVSCEGIPIDDWIPAEYKGNTSLVVG
WEFDHGWPGNYLHQMEIWAIMAKPRSPHLMQCINDILQELADKTAEHGITVENTTMDIMGDTVEFTGPRRLTSA
VYKSLGGMTNRNLVGSDTEELLQPKLVGDVLFMPGRSFAPMTNTYSPEEEAILSPQLVTHHYAGTWKNSHGGED*

>g9709.t1

MLNPIHRLTLDSLPPAFIEQPTPSLFHIQGATTIPKYEGIPKLIWYKLGPRGLSEEARNWTDSCIKPNPEYEARFMTDES
SDEYVRKTFASRPDIVDNFLALPVPIWKADILRYLLLWDQGGIWFDLDVSCEGIPIDDWIPAEYKGNTSLVVGWEFD
HGWPGNYLHQMEIWAIMAKPRSPHLMQCINDILQELADKTAEHGITVENTTMDIMGDTVEFTGPRRLTSAVYKSL
GGMTNRNLVGSDTEELLQPKLVGDVLFMPGRSFAPMTNTYSPEEEAILSPQLVTHHYAGTWKNSHGGED*

>g9733.t1

MKDLLVSSGEKPYYNLGEYGREIHTTSQEARKWFNRGLVWCYLFNHEAATQCFETAAAHDPNCAMAYWGIAFALG
PNYNKSWRMFTSVDRQNSLRKILSALARAEKVQADVSPIERALISALATRFLRSSVAIPEDLSRFDYAYAEAMRSVYEA
YGEDLDITTLFADAVMCTRPRQLWNLNTGKTTGADIDEARIALEKGLAQVEGRNHPGLCHLYIHMMEMSPFPELAL
NAADNLRRLVPDGSHMQHMATHIDTACGDYRRSVDSNFDAIRADDKYFSRRDVKSLI*

>g9734.t1

MMSGRSADALYAARRLPEVLSLEFMSIKTPRMVDWTEWQLVTLPHALIRFGQWEKILQLRLPANRDLLSVMTATV
HYAQGIAFAVLGRITEACHARDAFEDARKAVPDNRMYSPSSMAAPVLAVAPAMLEGELEYRKGNHSKAFSILRHAI
NLEDNLAYADPPLWMQAVRHAFSALLLEQGYSEEAEKLYLEDLGLSDSHPRRKARINNVWGLHGLYESFMRNGKEE
KAKSIRIQRDIAMAASDVPIKASCFCRLSAIKRDYDCHS*

>g9797.t1

MVKAVPTPVALPVTQGSSSSLSFTQGFLLGQLSIAILMFCFIKFFIFGEPPSADDRALHLNSLRRARTLAHQQSYKQLR
TRANSTSLSLRHKPSRSIIRKGEESRGGPSITTILAKTYYNVKGHQPESLDWFNVLIAQTIAQLRADARQDDAILTSLTE
VLNTGSKPDWIGEIRVTEIALGDEFPIFSNCRVMPAEDGFWYGPGNTGNEKERLQARMDVDLSDVITIGVETTLNL
NWPKPMSAVLPVALAVSIVRFSGTLAVSFIPSSSPPPTAAPMTSPTSETHRGSSPSRPASSSGAPPHRPTTLAFTFLDD
YRLDLSVRSLVGSRSRLQDVPKIAQLIESRVHAWFDERAVEPRFQQIVLPSLWPRKHNTRGGSATEDADAAAGENE
NDEDDEIAVMEGNGTAPGSTSFIPSPIAENATLEERIEAEGQKMREAEIRAGVRKPSASDSRSRSRNNREDGMRWR
GEQRDRDRPKGPHGRPKLQSRTTTGIAGAIPGALPRG*

>g9798.t1

LSLRHKPSRSIIRKGEESRGGPSITTILAKTYYNVKGHQPESLDWFNVLIAQTIAQLRADARQDDAILTSLTEVLNTGSK
PDWIGEIRVTEIALGDEFPIFSNCRVMPAEDGFWYGPGNTGNEKERLQARMDVDLSDVITIGVETTLNLNWPKPM
SAVLPVALAVSIVRFSGTLAVSFIPSSSPPPTAAPMTSPTSETHRGSSPSRPASSSGAPPHRPTTLAFTFLDDYRLDLSVR
SLVGSRSRLQDVPKIAQLIESRVHAWFDERAVEPRFQQIVLPSLWPRKHNTRGGSATEDADAAAGENENDEDDEIA
VMEGNGTAPGSTSFIPSPIAENATLEERIEAEGQKMREAEIRAGVRKPSASDSRSRSRNNREDGMRWRGEQRDRD
RPKGPHGRPKLQSRTTTGIAGAIPGALPRG

>g9847.t1

MSFEMVTVRVGEADVAVDIPVYRDQLSAVSLYFRGAFEGPFKEATDRILPLTDVSEQTFRIFLQWTHFQANSQSSAA
SMRTHDAVLQKLMTKLRDETTTIPGIDEKGYHDERKKNDSRWTNPESVVDCQLMRASFLRLYVFADKYDIPQFRDD

ILTALIAQPHIWKWPSSPDQDLIEHAYADLPQSSKFIRFLVLSVATWYIRDSSYEDATRLLCDLKETHKDFALEVAILQVE
IYRDKVFPNKHGASSQWELTLPHSCILHEHRVQDKKQCRERIRNHPYIFNILIDACLQDALGMEERCNKA*

>g9848.t1

MLPNFLTMSFEMVTVRVGEADVAVDIPVYRDQLSAVSLYFRGAFEGPFKEATDRILPLTDVSEQTFRIFLQWTHFQA
NSQSSAASMRTHDAVLQKLMTKLRDETTTIPGIDEKGYHDERKKNDSRWTNPESVVDCQLMRASFLRLYVFADKY
DIPQFRDDILTALIAQPHIWKWPSSPDQDLIEHAYADLPQSSKFIRFLVLSVATWYIRDSSYEDATRLLCDLKETHKDFA
LEVAILQVEIYRDKVFPNKHGASSQWELTLPHSCILHEHRVQDKKQCRERIRNHPYIFNILIDACLQDALGMEERCNK
A*

>g10182.t1

MPILRNTNWNYGTPKARYRKNELQLRQPKGFPIKLLKITLDRAREEAELEAATKRYADPAFEIYEDWKENDELEEEDD
YDGKVEPESEVEDSIAEDTLDDFAYGNADVSDASAEFPTSRVIPEKPTALVDSLGRTRAATSRRKAQEVSDSIISLGVSR
QGSGSTTVDNKKRKHTNTQKAKQPRKTLPKKSV*

>g10183.t1

MVPGKAVTEKTREEPRVDSGAALSDPAALSGGSANNKLTMPILRNTNWNYGTPKARYRKNELQLRQPKGFPIKLLK
ITLDRAREEAELEAATKRYADPAFEIYEDWKENDELEEEDDYDGKVEPESEVEDSIAEDTLDDFAYGNADVSDASAEF
PTSRVIPEKPTALVDSLGRTRAATSRRKAQEVSDSIISLGVSRQGSGSTTVDNKKRKHTNTQKAKQPRKTLPKKSV*

>g10437.t1

MARQAGGPTQWFGIFHDAASLPHTNHTVAHLSCLMPGRWTCRPKLCLSPIRLSLRAQCRQSSAHVPGRILRGRDL
GPTREDAHQTFRLRRDETAKELPLSPLLDPLVVDERSRFEQTKERPKFAEFTPFQKKLWMNPFAHALASPIRHCRAV
QIVLPTAFLVSLNARPHPTTKDPWLLPVSLTTDKKHLGPPFRFLGRHLTAAYMGKRRAWERALYARMNEKYGGHNL
RKMVWREDMPDFILDVMRKRVVSKLSWNFGFRGRLIPVASPRTEDIEGVEDVSCVLIFRSLRTRADDLQNQADRIT
AELEKWSNYFTKSFEAKLDPHAALEVTHKAPNWYSGPVVSHFKPRVRYPELEFHTTFWRGKKVAVYSLTDLLGENK
AQELIEGSQYAGERSVVIKAARHNVPVEILLMQLQAYIAQPGP*

>g10438.t1

MGKRRAWERALYARMNEKYGGHNLRKMVWREDMPDFILDVMRKRVVSKLSWNFGFRGRLIPVASPRTEDIEGV
EDVSCVLIFRSLRTRADDLQNQADRITAELEKWSNYFTKSFEAKLDPHAALEVTHKAPNWYSGPVVSHFKPRVRYPE
LEFHTTFWRGKKVAVYSLTDLLGENKAQELIEGSQYAGERSVVIKAARHNVPVEILLMQLQAYIAQPGP*

>g10458.t1

MAPTAPTHDYYVTLEVLPTALHDEIKASYRRLARLHHPDKNIGSQHATAKTQLINAAWDILGDVDKRVEYDRSRPQP
KAPSSGSRSGPQPTSTQQPKYQASHRPDTTAQDEARAQAESDRKRREWLEFERYHEEQIRWCRNTIKPLDAEVDRL
NATIDANRTRLANDVPYA*

>g10459.t1

MLDAKSRQDNRLSGTRLPKKRQRGFDNSKRSKQENLQNARERNVQPRRQDGKPTKKKQSVLHRKELESKKQKKSE
AENLTNVTQVSTTATTGVGGTKLKVITPVRIAPDHYSSLQSSVPAATP*

>g10634.t1

MAEVLAVFAAIQATTQLVEQAFRIVDRLRRAYSRQKALAEVLTRHRSELESIKAIIGIIDDEEDLQIPTVATELVRLQGVQ
VRLAELLENLDPKMTSKMNQIARQLTSGSADEKKLGDIMDELVQVKTMLVLRIQVAHVGVIRTIGKEAVVNAEAIQR
IDEYLREHVHNCEGLKIAKLLKGRRRSNDGTVPLTLADLKALSNEANGEGEDSGDETLVDDIEVVSRDLPVKTERIITR
NIARNQALQINAAIIEDLWKDVNHLKIHDNVAEDESTQVNYAMNREAFSVVLEQRNKAVSAPRQKTAVRAKRAS*

>g10635.t1

MVVATSVSRNISSGNAVQINGPVADKVYNLNFKTLVVSPDCKCGWGIEIVSSGNESALPSRCAAKVPNKSKVLKTPS
SSKVSGKFTVQLPKKTSDRCVLQGQNLTLNSSTSDTRHVVRAIFRPTILNNFIAHKIVRRLSLVVYSGSSFTMETTSDRI
HLPPTNDYVDLARYSQEGKEEFAICRFYVVKDSRFDLLFGADSVNT*

>g10706.t1

MVTAFAYSGRLDFDPTTDYHPIEGSAEGFRFTAPIDVELPSQFAVGEDFYQVSVHDADDVVVDIDPQSDRLKLLEPFE
SWSPGCTKGMSILAKVKAGLIHEVARDLRDRGIKWCIVGDENYGEGSSREHAALEPRFFGGVAIIARSFARIHETNLK
KQGMLPLTFANPRDYDRIQLGDRITLQGVKDGELRPGRQASMRVECAHAGEWVAPLNHSYSEGQLRWLRAGSAL
NHIKQTVLGR*

>g10707.t1

MPFVQPGDHDSKGSSSFRRSDFLADGELRRVEIQSSRVSGFLLCRPTKLRLKLDTTVFRSRPLTSTSVRCHCPTHGPH
VLLSTTAPASRNASHCLLQHSSNMKEDAKGLISTVAIAVAPNVSVPGPKNLRTLPFPPETERIPASLQMISLVEAHPCR
RPVSRTPTTCGHMSSQGIPAIASTASEPPTPMPGIPMPPALGV*

>g10759.t1

MPVLHDILISMSTDDTLPWTTSRCNRLLRPLSSKLTKLRKELERPRTASAETRSVSSAFAIKGSPQKTTNFTRPANKPR
GFEKARDPDWRPGAKPGTGKKTTYGGRGRGRVSDLHKSGSHNGNANTSRPGEIAFTPLISRMGGQLHSSPQIPISP
LRRYSKVKGPLLAPLNPPAIHVSDQIRKLVQGLSEAYANILQSTTTGGEARWKGTRSLMGACLRKLPAYIELEEHFAKL
DRLEEDGEEEDRDIVSEIYEHLETRFEQEHGQGWRPFKCVVRAHATALICDAIMDKVFGLDNVLVLVTHCLNSSSWD
EAERLLLAYAPLVDSVPIPINTRADLFDPQRSSYLHSVGNFVRHTGRYRLLYDLLDHMVALELLPLEWLATDSMRPVW
DRLVRTITENDHRTLRSAHQFFETVTIASMGLPDARLLEDEPSGFISRRFVPSSRKELRQALETTFSSLLTVLCSIALVNA
NRDDLAGQEITQRIVWVLDAVVITISARGNIQDELRLFGADVDDVQIMAQRAIWIHFASFLVHLEGCVCSGISPLDC
STSIELVNWISDQYSSSNINLALTFATLPSLFSSIARGTGRIWKDNGFDQLQRLVTALMTTSARRLPHKLWTLKRLALES
AREFAQGTGDGEHMTYVRNIEKTMQTRGRLVIMHSPQKNESPTTSGGFRWEEGIGEWVTCTPFVRQNISHSARK
PIRTLELLPTPMQSEDDMTEAPDVEDHLPGGTPDTPIYEITSLEYYGDQVLQSSPIKNRPRTSTPSLGKRIRAPSPKVVI
PMKRTNVTPPDTPPFRYYSELPNEEQDSPRRSRRPRKELKTLPPQRYSERFRTSLQSGLRDLKRPIYIEPALEVSHSDA
EVQSISQDDSDSSLVSVEEVHPEIFQTKALPSQRACTRGSHPINSTVNNDGDTSERDELGRSPAHPRSQMRKRTSVR
QGIQPRRQMWKAGQGMMADVNHDSEDELSFC*

>g10760.t1

MSTDDTLPWTTSRCNRLLRPLSSKLTKLRKELERPRTASAETRSVSSAFAIKGSPQKTTNFTRPANKPRGFEKARDPD
WRPGAKPGTGKKTTYGGRGRGRVSDLHKSGSHNGNANTSRPGEIAFTPLISRMGGQLHSSPQIPISPLRRYSKVKG
PLLAPLNPPAIHVSDQIRKLVQGLSEAYANILQSTTTGGEARWKGTRSLMGACLRKLPAYIELEEHFAKLDRLEEDGEE
EDRDIVSEIYEHLETRFEQEHGQGWRPFKCVVRAHATALICDAIMDKVFGLDNVLVLVTHCLNSSSWDEAERLLLAY
APLVDSVPIPINTRADLFDPQRSSYLHSVGNFVRHTGRYRLLYDLLDHMVALELLPLEWLATDSMRPVWDRLVRTITE
NDHRTLRSAHQFFETVTIASMGLPDARLLEDEPSGFISRRFVPSSRKELRQALETTFSSLLTVLCSIALVNANRDDLAG
QEITQRIVWVLDAVVITISARGNIQDELRLFGADVDDVQIMAQRAIWIHFASFLVHLEGCVCSGISPLDCSTSIELVN
WISDQYSSSNINLALTFATLPSLFSSIARGTGRIWKDNGFDQLQRLVTALMTTSARRLPHKLWTLKRLALESAREFAQG
TGDGEHMTYVRNIEKTMQTRGRLVIMHSPQKNESPTTSGGFRWEEGIGEWVTCTPFVRQNISHSARKPIRTLELLP
TPMQSEDDMTEAPDVEDHLPGGTPDTPIYEITSLEYYGDQVLQSSPIKNRPRTSTPSLGKRIRAPSPKVVIPMKRTNV
TPPDTPPFRYYSELPNEEQDSPRRSRRPRKELKTLPPQRYSERFRTSLQSGLRDLKRPIYIEPALEVSHSDAEVQSISQD
DSDSSLVSVEEVHPEIFQTKALPSQRACTRGSHPINSTVNNDGDTSERDELGRSPAHPRSQMRKRTSVRQGIQPRR
QMWKAGQGMMADVNHDSEDELSFC*

>g10765.t1

MKCLPANKLPVAHGNILHDWHAEVIAIRAFNRYLLDECTLISTPPYPTSGLLRKVSPDEQTRDLQQPFTIREDVSLHM
YCSEAPCGDASMELTMAAQEDATPWTSTPPTLSSTPHSGDEGEASALRGRSNFSLLGVVRAKPSRPDAPPTLSKSCT
DKLAVKQATSLLSSTSSLFVSPQNAYLETLVLPDSQYIPQACERAFSSSGRLRCLDSNEWKGGYRFQPFTILPTDREFT

WSRRALCSTEKAVPSNISAVWTPTWQETLIGGVMQGRKQLDPRGASKICRRGLWLEGLRLAGVLGGTVVTSGALR
KRKYAEMKGSEELIERTRAKEDIRKALKGWVRNTGDDDFSIKPSLSTP*

>g10766.t1

MPPDANAIADCVLRAFAQLPEKRKPRPRHDGAREWVPLAGIVLSKGQSLSCVSLGTGMKCLPANKLPVAHGNILH
DWHAEVIAIRAFNRYLLDECTLISTPPYPTSGLLRKVSPDEQTRDLQQPFTIREDVSLHMYCSEAPCGDASMELTMA
AQEDATPWTSTPPTLSSTPHSGDEGEASALRGRSNFSLLGVVRAKPSRPDAPPTLSKSCTDKLAVKQATSLLSSTSSLF
VSPQNAYLETLVLPDSQYIPQACERAFSSSGRLRCLDSNEWKGGYRFQPFTILPTDREFTWSRRALCSTEKAVPSNISA
VWTPTWQETLIGGVMQGRKQLDPRGASKICRRGLWLEGLRLAGVLGGTVVTSGALRKRKYAEMKGSEELIERTRA
KEDIRKALKGWVRNTGDDDFSIKPSLSTP*

>g10813.t1

LANRVDRNLISFVSHLNSAPGLGDLAKKAGLKYFGTAVDNVVLDNDEYTSIAFERSEFNQVTASNGQKWVYTEPQR
DFFNYTLGDQIVDASKEADQIRRCHTFLWHNQLPTWLTTGTWTKASLLTVLENHIKNVAEYYWNDCYAWDVVNE
AFNDDGTLRKTIWLDTIGPEYIEHAFRLARQYASPGTKLYYNDYGIERVNNKSLAVARMIREYQGVSVPIDGVGLQA
HFTVGRAPLYTDIRASQQLFSNLGLETALTELDVRMTLPDNANKTAAQATIYADSVRACADEESCVGVTVWDFWDT
VSWVPETFVGEGNACLWNQNLQRKPAYYAVADVLRKAGEKRG

>g10814.t1

MKLSIICSVFPLVLASPTPAPIHPRSSFVSHLNSAPGLGDLAKKAGLKYFGTAVDNVVLDNDEYTSIAFERSEFNQVTAS
NGQKWVYTEPQRDFFNYTLGDQIVDASKEADQIRRCHTFLWHNQLPTWLTTGTWTKASLLTVLENHIKNVAEYY
WNDCYAWDVVNEAFNDDGTLRKTIWLDTIGPEYIEHAFRLARQYASPGTKLYYNDYGIERVNNKSLAVARMIREYQ
GVSVPIDGVGLQAHFTVGRAPLYTDIRASQQLFSNLGLETALTELDVRMTLPDNANKTAAQATIYADSVRACADEES
CVGVTVWDFWDTVSWVPETFVGEGNACLWNQNLQRKPAYYAVADVLRKAGEKRG*

>g10912.t1

MPPSLNGHGPASPASFGFSDADLTAFTRRFEHFQQFEEEKTQFLAEIVTRYEYLYQQYQALAIEHERDRVWITAWKN
EKMQYENKHNHMLREMSDNPFITVLIDGDGMIFCDKYLGDGEQGGRRAALDLSAAVQEYVDNECNDIPYGARIV
CRIYANVRGLGDVLVRKGVYQDPSEFEKFVRGFTRGRALFDFIDVGAGKDRADEKIIESCKLYSQDYHCRRVLFGCSH
DNGYARMLEECSDRPGLVNKIILLEGVPFEKELVNLPYDTKKFPGIFRDKKIVFWGAPIYSGGLPPAFIPARRVDSNDS
SKASVSSTGLPSRFPPPAKSPVMDSPLPSRATMMGLPRTPSTSTLASDGMITATKPVLPMNWAAKVSAPPPPVNHS
SPTYKPANREEVIARNRAG*

>g10913.t1

MCNVHFLRNECPYEKNCTHLHAYKPTESEVATLRLVARMAPCSNGSGCQDIRCIYGHCCPAPHKTHHVKGTKNCIF
GDSCKFPIELHDIDTNVVKTLVIR*

>g11026.t1

MAAETPISPTIQPLLRISLDLEASSDPKSSHVSLRQIEDGDISIFGSDDEELGARPGFWRRLKMSIRRRNGKSEIDEDFG
RVRLPDEKTQSKKYRLKKKHAARACVIIPLLVLIFFGVLHIVNVFLGYVPTFLDDHIVSTFDWSQHDKSDLLDVTRDVT
PVQCHSHNDYWRRVPLYDALRWGCTGVEADVWLFDEELYVGHNTNALTRDGTFTSMYVDPLVKMLDHRNEFGD
FATTSSVKNGVFITEPAQTLVLLVDFKNNGHDIFPVVSQQLTALREKGYLTYFDGNTTIEGAITVVATGNAPFDLITANS
TYRDIFFDAPLDKLEEGTSVGQRGGQGTVGTTPSSHFDSTNSYYASVNFGESIGTLWFGRITEAQVLKIRAQIKSAKER
GLQARYWSAPKFPIGLRNRVWKLLVEEGVGYLNGDDLQGMTRLDWGVKRHWGLIV*

>g11027.t1

MFSRKKHRYADCLSGVLHIVNVFLGYVPTFLDDHIVSTFDWSQHDKSDLLDVTRDVTPVQCHSHNDYWRRVPLYD
ALRWGCTGVEADVWLFDEELYVGHNTNALTRDGTFTSMYVDPLVKMLDHRNEFGDFATTSSVKNGVFITEPAQTL
VLLVDFKNNGHDIFPVVSQQLTALREKGYLTYFDGNTTIEGAITVVATGNAPFDLITANSTYRDIFFDAPLDKLEEGTSV

GQRGGQGTVGTTPSSHFDSTNSYYASVNFGESIGTLWFGRITEAQVLKIRAQIKSAKERGLQARYWSAPKFPIGLRN
RVWKLLVEEGVGYLNGDDLQGMTRLDWGVKRHWGLIV*

>g11210.t1

MALSASSAALHETAYRLRWWNPPFRGTAFVTSLFPCPRRAFASCKVVQDEHQDNNELFEYTSGRWIPNESLRTAER
RMPFNVSELKKVAAKALNKPVEDVDMLRKVAEGGFNRILEVTMNDGASILARLPYPLTVPRRLAVASEVATPDFLRA
NRIPVPQVSAYSTGKNAVSAEYMLMEKMPGKPLSSVWVHLTDDERFKILHQIVTMEAKLFAMELPASGSIYYSYDLP
PTMPRIDIPGFDNGLCIGPYAHSVWWYGERIDLDIDRGPHTNPRLVLRSFANKELTWIHAHGRPRYPFDREYMDSF
DHKKQDPQEHASSLRDYLSLIPYLVPADSTAHTPTLRHPDLSPNNILITDDLIIAGFIDWQHAMVLPAWLTAGIPPAFE
NYADEGSRGLQIPKLPDDLRSMDKTLRAKAEE*

>g11211.t1

MRISRSTFHRLFAIVVIEEQEMQRRENLRISLKEGDVFRKNMSEMMGVLPDGSISHENFYVAKERESMIREGVGTKL
KDDLLEAERVLLAWPFHDFDEDE*

>g11227.t1

MLQIFKRHKKRRSQATEDPQTTNRRSSSLASIETDTTSSPPSTRYSSSPPSLPVSKGRPSPAQISYSTSAVPELQLHTVP
EATVRPAPVIRNSLPLSASASSTSAVSTPTRVIPTTPNSTGRNTQLVFELSPQPDSPTSPSPVRSQPYNPPSPSPIRPSVS
ASVSTESVDTIKPLQPSTPIMQTTRPTSSRQQTNGASAVLSPPASPQALFRTKGRLPSFTSSRATSGNFSTMMSYNKR
GNSNSHQTSLVQDSAASLRKRSAAELGAGVYQSIGNVSYVAFLEWIRSERLTTLPHKGSRWDKVLIRALYFAEQLHKF
ELAVRPFAQDSSSAAIIGYGHAQLLLELSHHNSEALDKAFSVFYKFAMSFQSVLHRSELLNASSEITEQICLLYTDLITLV
VDVAIKFYKTVKGMTPGSTTLDIFELFGETIETFRTRQNNVVELIWQSQIEKGHFEEEEAIDVQHLSRWLSSQDRVIA
AITRDHETYVDNQAEFTCLWFLKHMTRFFQSDNRVFLVTGQSGAGKTTLAGSIVERLQRPMSKKQYDTLFCSLSPDI
PTAATSLAVVKSLLFQLLNLRIGNMGVYKALFDAYHQCRSSDDLKAYEECLWHALGEALRNPVSGGNELVMIVDGLD
EIAASNSASIQAAGVVSPADLLERLAKVTQQGKGVRLITLSSSMKLPSSAKGIQHQIINEDVRDDLHAVALRALIHNH
HFHGQRPFEQEQILDRIIRMANGSFLHITLTCELLNTQKSPDALMKTLEGFESSKPSVQDLILKLFTTLNPTNDAKTLLS
WVLAAERPLTIDEIQTLFSVDVQRGTISEKGVNTNEIFKTLAPLFTLHQRIVRFKHPVIHSTLHDFSANGKIPIPLKDSET
DMLLRVLAYSKHTLKDHTAEPTFDTFDPSTPDRLFRKHPFLEYAVRYWVLHLQQSPLAPKSSGEFKPTSELQKVFPES
TMFPILESYCWDTQLPIPEALELQRLVSILRKNMLPENHPGLLQTYLSIVTSYLLISNVTEATKYLYLCTIVSRKVLSDMH
PLTLDCASQFLKITETQTTTTRTEIMTRREEILKVLITTYERQYGSTSELVIETRKVLAQLYETINEQERSIEIYRLIQDATIQ
LYGRDSHQAHDIHDHLQVTLGKGTGDREIEIYDDFSFEVDQDDEEHVEILTISTINAMLIQIQKLLTMKRFAEVERMY
VELWMEVSSKCRTVQSVEWHEKNIEMATSYSQFLKSQKRTNESVAILTSVWQQYESHHLSFAKSIVQRLTSVAKEM
RSMNAHSQALSIFKFAHSYYHKSSSEESSESREISQQLSQTSNELVQHSLNSSSSTTETTTTISESVYQDVFFTTISSSKTI
ESSTMALAKKLVVQHMAKKNYTAAISVVHATLQRTWSSFLANSIHDVTMATVFTQESIELVERLAECYLQTRQLERV
DDVYSRFFRAVLVTENVDKAIFEKSKNLLINFYDKHGYADKAISTYQEILVVYRNRLGHTHEHTIQTLYILAQRCRSHPR
NHGYWLDYYLQILTALNKDSDVCHKDALDAMLVVTVTYWEDRRYAEAVTFYRVLWNTFVRQTKQHKVFTDVKFVE
TLYERYYQCLEETKASWSELYKVTKEYRETSIAVFGAESTIAVNATLALAQVAQSSEQHMSEAISLYEAVSQSGKQVTTT
TSISEIKHALSSLYSKQMHSSSSSSMKAETVQRALSMSESQFEQSTRENGYSHESSLTSLRELSSLYSRQQKSDVAVKQI
SRTVSEIISKETSSQKQIESARSIAATFQAIEQVNTAHAMVHELHRQICAKDARNASKWSFDLTKSSRTAIAFLASLQYN
LRQDLSISFSEIYADLTMEYIYWVQFHQTLDNNESLTNILLAAAPLRYFLRRNDQKDMIAVVEEQAVGLFVKRDAQDL
KDFSKESPRIFIIGILDHLGNGRNKNFNRAVILSSNESVAKCTKAKMFHEAYDIANLGFLYASKHDGYNGPRAISMGFK
LASLLVGRDGEKCPDPTLRKKMLVLSNRIVKKIFEICRNLKINFAQVQLYELSHLSVLLGEQEDYESLEWLLTTLWQTRD
AQKSWPAEVLLNLGRRLICARYLANEEVKAIRLAEDIAYNMRRAHGPRAPVTIETYELLAQLYTSTGNKYQAQAAKG
EKTAGLAADYFKKAIGIHEDILRVLVNDPANSANEDEDDDDDTTAELLAREGVNVNNAPNSPPSAAVDSDALDKSAT
ALKHLHLLKLAYQRYGGWPKSYDEYEHLNAQLFRVFGSEGKWKGVEGTEKWDAKAFGAGKAESQEGGFQGIADW
SLGSDELVSIGHGHGQGQHTAMVNGNAMLHNKRIN*

>g11228.t1

MMSYNKRGNSNSHQTSLVQDSAASLRKRSAAELGAGVYQSIGNVSYVAFLEWIRSERLTTLPHKGSRWDKVLIRALY
FAEQLHKFELAVRPFAQDSSSAAIIGYGHAQLLLELSHHNSEALDKAFSVFYKFAMSFQSVLHRSELLNASSEITEQICL
LYTDLITLVVDVAIKFYKTVKGMTPGSTTLDIFELFGETIETFRTRQNNVVELIWQSQIEKGHFEEEEAIDVQHLSRWLS

SQDRVIAAITRDHETYVDNQAEFTCLWFLKHMTRFFQSDNRVFLVTGQSGAGKTTLAGSIVERLQRPMSKKQYDTL
FCSLSPDIPTAATSLAVVKSLLFQLLNLRIGNMGVYKALFDAYHQCRSSDDLKAYEECLWHALGEALRNPVSGGNELV
MIVDGLDEIAASNSASIQAAGVVSPADLLERLAKVTQQGKGVRLITLSSSMKLPSSAKGIQHQIINEDVRDDLHAVAL
RALIHNHHFHGQRPFEQEQILDRIIRMANGSFLHITLTCELLNTQKSPDALMKTLEGFESSKPSVQDLILKLFTTLNPT
NDAKTLLSWVLAAERPLTIDEIQTLFSVDVQRGTISEKGVNTNEIFKTLAPLFTLHQRIVRFKHPVIHSTLHDFSANGKI
PIPLKDSETDMLLRVLAYSKHTLKDHTAEPTFDTFDPSTPDRLFRKHPFLEYAVRYWVLHLQQSPLAPKSSGEFKPTSE
LQKVFPESTMFPILESYCWDTQLPIPEALELQRLVSILRKNMLPENHPGLLQTYLSIVTSYLLISNVTEATKYLYLCTIVSR
KVLSDMHPLTLDCASQFLKITETQTTTTRTEIMTRREEILKVLITTYERQYGSTSELVIETRKVLAQLYETINEQERSIEIYR
LIQDATIQLYGRDSHQAHDIHDHLQVTLGKGTGDREIEIYDDFSFEVDQDDEEHVEILTISTINAMLIQIQKLLTMKRFA
EVERMYVELWMEVSSKCRTVQSVEWHEKNIEMATSYSQFLKSQKRTNESVAILTSVWQQYESHHLSFAKSIVQRLTS
VAKEMRSMNAHSQALSIFKFAHSYYHKSSSEESSESREISQQLSQTSNELVQHSLNSSSSTTETTTTISESVYQDVFFTT
ISSSKTIESSTMALAKKLVVQHMAKKNYTAAISVVHATLQRTWSSFLANSIHDVTMATVFTQESIELVERLAECYLQTR
QLERVDDVYSRFFRAVLVTENVDKAIFEKSKNLLINFYDKHGYADKAISTYQEILVVYRNRLGHTHEHTIQTLYILAQRC
RSHPRNHGYWLDYYLQILTALNKDSDVCHKDALDAMLVVTVTYWEDRRYAEAVTFYRVLWNTFVRQTKQHKVFTD
VKFVETLYERYYQCLEETKASWSELYKVTKEYRETSIAVFGAESTIAVNATLALAQVAQSSEQHMSEAISLYEAVSQSGK
QVTTTTSISEIKHALSSLYSKQMHSSSSSSMKAETVQRALSMSESQFEQSTRENGYSHESSLTSLRELSSLYSRQQKSD
VAVKQISRTVSEIISKETSSQKQIESARSIAATFQAIEQVNTAHAMVHELHRQICAKDARNASKWSFDLTKSSRTAIAFL
ASLQYNLRQDLSISFSEIYADLTMEYIYWVQFHQTLDNNESLTNILLAAAPLRYFLRRNDQKDMIAVVEEQAVGLFVK
RDAQDLKDFSKESPRIFIIGILDHLGNGRNKNFNRAVILSSNESVAKCTKAKMFHEAYDIANLGFLYASKHDGYNGPR
AISMGFKLASLLVGRDGEKCPDPTLRKKMLVLSNRIVKKIFEICRNLKINFAQVQLYELSHLSVLLGEQEDYESLEWLLT
TLWQTRDAQKSWPAEVLLNLGRRLICARYLANEEVKAIRLAEDIAYNMRRAHGPRAPVTIETYELLAQLYTSTGNKY
QAQAAKGEKTAGLAADYFKKAIGIHEDILRVLVNDPANSANEDEDDDDDTTAELLAREGVNVNNAPNSPPSAAVDS
DALDKSATALKHLHLLKLAYQRYGGWPKSYDEYEHLNAQLFRVFGSEGKWKGVEGTEKWDAKAFGAGKAESQEGG
FQGIADWSLGSDELVSIGHGHGQGQHTAMVNGNAMLHNKRIN*

>g11303.t1

MDPIRYTRFDFEPCTPFPTYQRKFPNIHDNHQPNTPEKSRTIRGIGITRAIEAGKRLKRFVGTAEPAPFVTPPAIPKDLE
REFVAGMKEYGSDHRMQSMGLDVLSQAASMQERLPELGEEHESPVKATWTSTQSGQQTFGSSPPRMTSQGPAA
NSVSSDGPQPARASYDTPSLAAFKKQMEAFEEQHVPAYSNANPNVGSDGIEASGSRHAYKYNTGSESLSSQKGTSRI
KPLFPSRVGAQSWDEGKGISQQHMVPCQPTPAASSSRDSACFENLFRQPPTFSVPYIVVQSPSPTKLAPSKKRLAPQ
AGDELFRPDVKDIRSNNKKATGLPTLPASSSALWEVNFPGYTPALLTILLTWSHAMHSFYRRLPDPKTFSIHAAFPRRI
TPPAYNRLISVGFYDTSFIPHKEIRFLGPGDMAEIGYAEIDIFRSKEEVVAFNAQAQEHPSKAQAMKRRLGIGSKHGER
KRTLGVYRDQIHLADSGEGR*

>g11304.t1

MLAWHLSAVTDTSTCLHTVFPDNHEPILSKPPATPAPPEQPLRRLASLQNLISPSHGQKHLHREIRSVSSSAGSTEVEE
SSILPQEGAQTLKRTVVKLEKAGSIPLIEGYRVDLKEFRGWLDAVSKGEGKVIVWRESPPLISLA*

>g11329.t1

MKYTSVLSASALTSLASSSALVPREARVVPPNALDTWKRASPQAPDGYAPAEVDCPSTRPSIRSADKLSQRETDWLQ
KRRPNTVQPMREFLERVNIEGFDVGQYISNHQDNTTALPNIAISFSGGGYRALLNGAGALAAFDSRTNNSTGSGHL
GGILQAATYVSALSGGGWMLGSIYANNFTSVESIINKGEDSSIWQFQNSLFKGPPTEGIQLLSTAQYFENLVSTTRAK
ADSTAGDFNTSITDVYGRGLSFQLINASDGAPAYTFSSIADDQDFSTGNAPMPILVADERSPGDLIVSLNATNFEFNPF
EMGSFDPTTYGFAPLKYIGSNFSEGSLAQDQGCVAGFDNLGFVMGTSSSLFNQIFLQLNSFQNIPEILLNFVSDILQGI
GEDGDDIADYSPNPFYHYHNETNPSADRERLTLVDGGEDGQNIPFNPVIQPIRQVDVIFAVDSSADTVDADDPSQN
WPNGTSLVATYERASGSLMNRTSFPYIPGQDTFVALGMNSRPAFFGCNSSNVTTGDNIPPLVVYLPNAPYVFWSN
QSTFGKLDYTIDERNGMIENGYDVITQGNSTREGASNWPTCVGCAILSRSLERNGEAIPQVCQQCFTDYCWNGTT
VDRAQNYTPSMILSQLAATDKDSGVGKFVPNVLGLAMVAAVSGFLMM*

>g11330.t1

MTASALVPREARVVPPNALDTWKRASPQAPDGYAPAEVDCPSTRPSIRSADKLSQRETDWLQKRRPNTVQPMREF
LERVNIEGFDVGQYISNHQDNTTALPNIAISFSGGGYRALLNGAGALAAFDSRTNNSTGSGHLGGILQAATYVSALS
GGGWMLGSIYANNFTSVESIINKGEDSSIWQFQNSLFKGPPTEGIQLLSTAQYFENLVSTTRAKADSTAGDFNTSITD
VYGRGLSFQLINASDGAPAYTFSSIADDQDFSTGNAPMPILVADERSPGDLIVSLNATNFEFNPFEMGSFDPTTYGFA
PLKYIGSNFSEGSLAQDQGCVAGFDNLGFVMGTSSSLFNQIFLQLNSFQNIPEILLNFVSDILQGIGEDGDDIADYSP
NPFYHYHNETNPSADRERLTLVDGGEDGQNIPFNPVIQPIRQVDVIFAVDSSADTVDADDPSQNWPNGTSLVATYE
RASGSLMNRTSFPYIPGQDTFVALGMNSRPAFFGCNSSNVTTGDNIPPLVVYLPNAPYVFWSNQSTFGKLDYTIDE
RNGMIENGYDVITQGNSTREGASNWPTCVGCAILSRSLERNGEAIPQVCQQCFTDYCWNGTTVDRAQNYTPSMI
LSQLAATDKDSGVGKFVPNVLGLAMVAAVSGFLMM*

>g11816.t1

MLLPRPSVPENRCTPIGVEVVSYEQRPHDIYSRRQLAIATPDRGIEPFHYRTQDRAKTSHLPTQPSCPVLAPPLASSSS
RCDT*

>g11817.t1

MSGAGPSPRFFQQPLRYMKWASINKPAYFYSIMVGCAGPALVVTVPPIRRYMGEEPIPKIPMTYPVPKGPRPRPTG
FDDE*

>g11870.t1

MGGFLQFIYPIVSLLSIPLVSTATLSNTPVPRTYGIVLFRMFDMLDVYGPSEILQFIGGSYPTNIVYIAETLEPVTTRPVM
AAMNPLNSSVYPSLTPTHVFETAPDLDVLIIPGGPGWRNPTSLNATMAYIREITPKVRQVLTICTGSALAARAGILNG
KRATANKSSWPATVAANPNTTWVPSARWVEDYSSSPPIWSSSGVTAGIDMMLHWVEKSYSAENATNIARFIEHVR
ITDPSIDPFARNETVVEQAY*

>g11871.t1

MLDVYGPSEILQFIGGSYPTNIVYIAETLEPVTTRPVMAAMNPLNSSVYPSLTPTHVFETAPDLDVLIIPGGPGWRNP
TSLNATMAYIREITPKVRQVLTICTGSALAARAGILNGKRATANKSSWPATVAANPNTTWVPSARWVEDYSSSPPIW
SSSGVTAGIDMMLHWVEKSYSAENATNIARFIEHVRITDPSIDPFARNETVVEQAY*

>g11934.t1

MSLKFLSLALAAVATASPFDSMSKATSDDPYEACQPQGATGTTPPAVGTELSSLYTDILSSIQGITFDKRSVHGRADGF
GCRQSLDCVNVQNINIPMCYDKFTTNFQFPDGSFGNVAGGTYSSGGTEVNLISGDYTKDGQSANIYSNNEAEKPN
TSTLSIPAQYTGTGVGGAIPVTELGSIIVYTTTIPAVTYSAPTTVAETVEVATVSGVKVSTTVPATTITQATTIAAKTNVVT
QTNTASAAPQSTGAAGQMSVDTTRSFGMSVVGALLYALL*

>g11935.t1

MVQHPSRVSAAEKFGTAACAWPQVLWVCLSGTNQGHAFSVCRGGFGAAGMTLSPHFLYLLMALRLEYADLSPVE
SRDFDLVYWLAVNIGPRDLTGRMSRDSGIAADLPRTAKRDGVVVGGGHMYGSRVVDSQSFATLHTRAAPGQGVG
DEHMIAQRITDPSQLVKVLSYCGAPSVKRMVPVGQFGMVETNLIKPLAYFKMSLKFLSLALAAVATASPFDSMSKAT
SDDPYEACQPQGATGTTPPAVGTELSSLYTDILSSIQGITFDKRSVHGRADGFGCRQSLDCVNVQNINIPMCYDKFTT
NFQFPDGSFGNVAGGTYSSGGTEVNLISGDYTKDGQSANIYSNNEAEKPNTSTLSIPAQYTGTGVGGAIPVTELGSII
VYTTTIPAVTYSAPTTVAETVEVATVSGVKVSTTVPATTITQATTIAAKTNVVTQTNTASAAPQSTGAAGQMSVDTTR
SFGMSVVGALLYALL*

>g12058.t1

MLPPARSPGLGHSVSPSMGSSTRDNYTTGTPGVPGYSNALSPRSHLPRYNTQPDTPRSVHTTTMPLTYNGAYATTY
TTTAGSPETPPFNTQETLANITCEGNAVTPSIDAKIEKGFFYSGDRVWTCYRRNYFSVNVSFQLSPWIANARLYLEQP
NKAAQQIQSMAVSLAAAVDGATGKTIELIQHTPKRDKGPQLPMKKELLAPTPPGKSHEHGGYGLSNFHQTSTVAGP
QLPLQSDSDSSQQYSPTSHASSNYQHSFERIQFKSATANNGKRRAQQQYYHLIVELWANVQAPREAEPKWIKVAAR

MSSPVVVRGRSPSHYQNEGPHNAGTSRGAPGSGLGGGGHHGLGSTGRPSYLPWTTGLSGGSATGMGSSMYRG
NTYSLDPSPVGSHSVSSASSLSGAPVEGITGENHMTEDDDAKIMDAPQDYSYYPASIYEPIPPKLESTLPPPDRRIKDE
YPIAGWHLGGCGRFQGMESSRGYYPDVQAHTY*

>g12059.t1

MNQVLQSPLPLAPENPFSSEQDCLDHIGAHAHHNSFAGVPDQLRRDGVLAHTATGIHQASITGRYGSHRPDSAYQS
SAALSDSGSGSISTSPSSNASSALAYTSYPSYSSTLATNYPTHYSVPGTYFSAACGYGSMSSTRPLNDASSSAMLPPAR
SPGLGHSVSPSMGSSTRDNYTTGTPGVPGYSNALSPRSHLPRYNTQPDTPRSVHTTTMPLTYNGAYATTYTTTAGSP
ETPPFNTQETLANITCEGNAVTPSIDAKIEKGFFYSGDRVWTCYRRNYFSVNVSFQLSPWIANARLYLEQPNKAAQQI
QSMAVSLAAAVDGATGKTIELIQHTPKRDKGPQLPMKKELLAPTPPGKSHEHGGYGLSNFHQTSTVAGPQLPLQSD
SDSSQQYSPTSHASSNYQHSFERIQFKSATANNGKRRAQQQYYHLIVELWANVQAPREAEPKWIKVAARMSSPVV
VRGRSPSHYQNEGPHNAGTSRGAPGSGLGGGGHHGLGSTGRPSYLPWTTGLSGGSATGMGSSMYRGNTYSLDP
SPVGSHSVSSASSLSGAPVEGITGENHMTEDDDAKIMDAPQDYSYYPASIYEPIPPKLESTLPPPDRRIKDEYPIAGW
HLGGCGRFQGMESSRGYYPDVQAHTY*

>g12065.t1

MSADQQPLRIAIGCDDAGVSYKKAIIKDLEADSRIASVTDVGVPENSDKTAYPHIAVDAAKLVAEGKADRAVLICGTG
LGVAISANKVPGIRAVTAHDSFSVERAILSNDAQVLCMGERVVGIELARRLVKEWVGYRFDKSSASAKKVEAIMEYE
RENYKGLVEDQGKSKGC*

>g12066.t1

MTPPKRALFSLNLDTDADHVYQKVEDHNKDGVDRVKVLESKGPTRGHTDLSYVTRPFEKTVHRAVNEVIDGRKVK
SAEGEISYSTTRTERRSGGTSRRNQDTDDEDREKSRLNIVSKTKRRNIERKYDTSEL*

>g12117.t1

MANATQGLFKLPNELLLELAALLGPAYLRRLSQVNHRLLDFALDYSRTYVSRLAALPDDVVLRIVAYLDNVGYLGYMA
NIYLSRLAQASHRFYPLVMDAILREEVRTRSTRLMEFALAKRRIGMMKRLLQRGADVEARTFHYPCFSFCYTHFSMH
SVCYTMNCWMVSPLSLATWHGDIGMAKLLLHFGADVHAFFVHHDDCSFSGTTLPTLLSTPLLFAAYHGHEGLVQL
LLQAGASPIAQGSLYILSVAITERHGQIVLRLLQVLDSINEPSDSVIKELFRVASEGNLRSLPSELVRRYEHVILRRLPSVTS
TEWHQEGYIAATMELAVAAKFVKVVEYLAPLKTRLRTEQTSYLDSALYNIISCDTCRGKVRKRELHQEVYQIVETLLAQ
GANPDVSGQGESARDIASTHPDPRVRNRLLQAARKQPPKDLSSRVGRSWVNPSQASPPHVPRRSKRSRLPPPVNL
WDFVKPHNSQG*

>g12118.t1

MPSILPPKSVWAEIRSTTTPASAQAKVSQDHAFPALSRPHKTSMANIESSSIATAAFKGKTEASKTGAKESDTKQTYAS
GVGKTPSTQQRSRKSVKGKKKWEKLVLS*

>g12145.t1

MEDNLDIPEESMRHIEALLARHPPHLIQRMFSQAIDSRRNSTASSFMSTSTASSTTTSSSGSSWRSRLSMASTFSSST
GRSDTRGSIASSSSSRSKRRAEPRSYPAGLDPTRPVPATMPHVDTESPLDLKMEPSYTPTDDISVASPSLIPDDDKSQS
QSGEPFMFCTYCAEQKSQKTFKAKSDWKKHEMRMHETGEDWPCVVNGCNRIFDRQKDFIKHHQRYHAGRPLPS
LTDIGITLLPRRVFGCGFDKCKEVSIGWDERCDHVAKHMKNGATMDQWKYSNVIRNLIRQEALHDTWKELIGCLDE
RLRESRSQISWCPDNSRILRQKLQCCDLRPSREEVLITALSLRADIQLDSVHQQLPPGFVTPSRDSVPHVEKLSREQR
MHILIGNSNPQHSRARLASINAALLRICSSLAHVPDYDCGSSPFVEPQTPAVDTNNRRISYMDVDPGNYLDVAQPAI
PDLPPAMTTSTMHTPHAHTPIMDPEQHTHMYHDEYMEQVKVPPNPLETLYPNYFGGAPQFEESQYYDRPSFGQII
SKPLSKIGNRLSSSRSNTHPRPSSQMSQASSDIHPDYTVAAMDMRHPHLHQHQHEQPQHPIQQHHQHPQQHM
MAQQPQPYRDSTSQLHEHIHMYTTQS*

>g12146.t1

MASTFSSSTGRSDTRGSIASSSSSRSKRRAEPRSYPAGLDPTRPVPATMPHVDTESPLDLKMEPSYTPTDDISVASPSLI
PDDDKSQSQSGEPFMFCTYCAEQKSQKTFKAKSDWKKHEMRMHETGEDWPCVVNGCNRIFDRQKDFIKHHQRY
HAGRPLPSLTDIGITLLPRRVFGCGFDKCKEVSIGWDERCDHVAKHMKNGATMDQWKYSNVIRNLIRQEALHDTW
KELIGCLDERLRESRSQISWCPDNSRILRQKLQCCDLRPSREEVLITALSLRADIQLDSVHQQLPPGFVTPSRDSVPHV
EKLSREQRMHILIGNSNPQHSRARLASINAALLRICSSLAHVPDYDCGSSPFVEPQTPAVDTNNRRISYMDVDPGNY
LDVAQPAIPDLPPAMTTSTMHTPHAHTPIMDPEQHTHMYHDEYMEQVKVPPNPLETLYPNYFGGAPQFEESQYY
DRPSFGQIISKPLSKIGNRLSSSRSNTHPRPSSQMSQASSDIHPDYTVAAMDMRHPHLHQHQHEQPQHPIQQHH
QHPQQHMMAQQPQPYRDSTSQLHEHIHMYTTQS*

>g12212.t1

LTLALDHSSPRQSRGKQRKSYRVGEEYSFLGEDEDGSVPSGTQTPAFQGDDEDEDDAFMPDAQDGELEYDEDEEV
AEEDEDISEEEEDDDSDEDSTGPPRRNARGPKAVRAVPTTPAVRRAKKLADNIPSPVTFAKGSGVKVRPVDADAQL
RTRGIPDFDKIGGHEPRLKNLFGPDAAHLKPVLASRDYWFPQETFPIRSFAKVSSDDADVGSLRRSFFEVSGAREKES
EALRTWYADTGKPAFARAQRTKTLTKDEAKEYMITPGPETVNVLAGPVNTPQVNTLAEGSYMNIAKPFPASQDRRG
WLFNLGSWIQDAQWASNEESNTQYLAVAVEQRLTSEDQPKPMEQPKAPAFNPTQPFPACIQIWAFQAEEEGFLNA
KTEPRLELVICADWGAPKQLRWCPIAASDDNRNSDGRRNIRIGILACLWSDGRVRVLDVSAPVSEPDLHAPTYLHVT
HAAFEVSFPQTVPSCLHWLSGTTLAVATAAGTVGIWSLTHAGTLAAPEANNYSPRPWFYQQVADTYILTISSGWPS
NPQFLSISTADGFARLIDIRSPTADTVVSIRGRTLCLAQAWHEHTQSFVMPDEHYILRHTPIRRYYHNLYSMRLENSIT
RVATSPVHPGVLVGGTDGDVQTSNPIMRITNYKIMPWQQKWFVHEWRGPMERMLVKPTDANVVAAEGGPVEQ
SNNDATSEPQAGMSADEAKKVPQEILSQPLVRITEGYKAGQTGIAHTSAATKRGNPEVGRSISIFEEQTAITALAWNP
NLKFGTWAVAGMGSGLLRVEDVGI

>g12213.t1

MDHSSPRQSRGKQRKSYRVGEEYSFLGEDEDGSVPSGTQTPAFQGDDEDEDDAFMPDAQDGELEYDEDEEVAEE
DEDISEEEEDDDSDEDSTGPPRRNARGPKAVRAVPTTPAVRRAKKLADNIPSPVTFAKGSGVKVRPVDADAQLRTR
GIPDFDKIGGHEPRLKNLFGPDAAHLKPVLASRDYWFPQETFPIRSFAKVSSDDADVGSLRRSFFEVSGAREKESEAL
RTWYADTGKPAFARAQRTKTLTKDEAKEYMITPGPETVNVLAGPVNTPQVNTLAEGSYMNIAKPFPASQDRRGWL
FNLGSWIQDAQWASNEESNTQYLAVAVEQRLTSEDQPKPMEQPKAPAFNPTQPFPACIQIWAFQAEEEGFLNAKT
EPRLELVICADWGAPKQLRWCPIAASDDNRNSDGRRNIRIGILACLWSDGRVRVLDVSAPVSEPDLHAPTYLHVTH
AAFEVSFPQTVPSCLHWLSGTTLAVATAAGTVGIWSLTHAGTLAAPEANNYSPRPWFYQQVADTYILTISSGWPSNP
QFLSISTADGFARLIDIRSPTADTVVSIRGRTLCLAQAWHEHTQSFVMPDEHYILRHTPIRRYYHNLYSMRLENSITRV
ATSPVHPGVLVGGTDGDVQTSNPIMRITNYKIMPWQQKWFVHEWRGPMERMLVKPTDANVVAAEGGPVEQSN
NDATSEPQAGMSADEAKKVPQEILSQPLVRITEGYKAGQTGIAHTSAATKRGNPEVGRSISIFEEQTAITALAWNPNL
KFGTWAVAGMGSGLLRVEDVGI*

>g12.t1

MLCDDVALDDADQKYEAWKRGLDTAQGAIIRFVTKEAKSRVAGKFGGYLEGSHNISLIVMVSAKPTYVIRFPKLGRT
AVCFLEEKVRNEVQIMHLLRERTTIPLPTIYTWGTTDDSPSQIGPFIIMEYIGGRRLMEELKQPTETKVGEVDLWKDL
GNPRVLHAYTQIADYLLQIYQLDFNAAGAISRSHTRDWVVAERPLTDHMNSLATDVANFPTELFPTTYCSSPRMYFQ
QLADQHLNHLHNHRNAVDSLEDARRYFIARHRLKQSIDRFLYTNNNTSPFKIYCYDF*

>g13.t1

MIVDENLDIKAVVDFEFWNALPAQFAHGPPWWLTSLRPDEWIDSGFDFGALRSRLEPHVEQFLPVMEKVEKEKAT
DGSVALLSVPMRDSWISGRFWFNLAMDDSWTIDAVYWAALHKPGDEVLDEAMEDELKAIYDMKMKQLAAFNA
ECKERGIGDAGHVRNWIMIV*

>g25.t1

MGCKAPRRWAAKWPEGLDLLLRVGQHARAQTILQFFLDVVESSGPTHEQQLLGARGINTVEPRNIEEVLSAQFEDF
SLGLRPKHFAPLMGSGIFTQDGAAWRHSRALLRPQFTSNRYQNFEEMKKSVESLTDQISPNSVVDLQPLFFRLTFDT

TTFLLFGKTLSSLQSSDIAGKESEFAAAFNLGQDYLSHRGRLGDLYWLANTPEFWRACKTSHRFVDDAIQDALDNAD
KSEPEKTEGEDKKSYVFIDALIQETRNKKELRDQCLNVLLAGRDTTACCLSWTLRLLARHPQVLERLRKEIDEVVGLGE
NAPQPTRVDLKKMRYLDLVLKEVLRLYPSVPVNSRAALKTTTLPVGGGADGQSPILVRKGEAVGYCVYAMHRRTDIY
GEDALEFRPERWEDGTLLRDVGYGYLPFNGGPRVCLGQEFALLEAGYTVARLVQKFPFLTVPQDDPVVAVGKEKQIL
TLVVASGDGCRVHMRS*

>g26.t1

MYVLSLCHREINLVSGEAAFANTMLSMGPWISALSDGLPAPSISLLAALGLLALLAYKGWSSHEQENSLYRLEIEKGCE
RPRLWAAKWPWGLDLLCKAFWHGQNRTVCEFFYQISELSGPTHEQRLLGARNIGTTSPVNLEAILDTQRKDFNLGF
RIPQFRDLMGTGVFTQEGKGWSHSRQLLRPLFASNRFQAFEDIRRCVEDMLDNIAPNTVVDLHPRIFQLTLATTLF
MLFGDSAHRMISAADKEEQNLASAFNDAQEYLAYRTRVGPFHWLINGPPMWRACKTIHSFLDRAIEEALAVSDER
LIQQSEYKRYVFIDELIQQTRDPVVLRDQCMSLLLAGRDSTAACLSWTMRLLGRHQRVLTKVRDEIASIVGLGPDAR
RPTQDELKEMTYLNLAIKESLRLYPPVPVNQRAASYDTTIPEGGGPDGLSPVLVRQGESVGYSVYAMHRRTDIYGPD
ALEYRPERWQNDGLQGIGLGYLPFGAGARKCLGQEFAMLETRYTIARMIQRFPFITASLDSLKRASWNCITIFDLSPS
EDSRASMEPTINPYHTQPDIVVLKNLVQDVKCDIKGDPRSRNQQISPETMEKVADDQDQTVDHDIQAVKLLKSYND
ASSAEFAPTVFTTWDEASIPKVLNEYLVKPYARIAMKVVRHPTDVVFLTHIILYLTVNLGSAVWLFRNFTYLHAIVHLAY
TGWCIGSFTLLMHNHIHNNGVLKKSWKWLDMTFPYIVEPLMGHTWDSYYYHHVKHHHVESNGPGDLSSTIRYQ
RDDPFHFLQYYARFLFLIWAELPLYFVRKGQSNLAARAFISEASSYIFLFTMYKLNPRAATWVFLLPFAVLRLALMVGN
WGQHALVDEVDPNSDFRTSITMIDVMSNRVCFNDGYHTAHHLNPLRHWRDQPVHFVKSKDAYRAGRALIFYDVD
WFMMTVKLLMKDYLFLADHLVPIGDQIGMSRNELADMLRTKTRRFTEEDIKNKFKH*

>g124.t1

MKLILSTSNTMSGGPSVIRRPFFEKSNTELTSSLRANFAAHQPSPTPSKTNYATWTTKTDSALYVPTHTPSPLAEPRES
YDITVKLFYLPNIPTDRRCAQTREAIELVLKELGTSSIDLLIVSFPGITFDADDEDSDLDDDDPPSPPSATQSTSENTAGD
CPEAGAPPEDIETMVTTWRSLEKLHAEGLVSKLGIAEFGVARLTKFLEQTKIKPSVNQINVRDCCVVPKPLILYAKQQQ
IELLTHNDCTNILPRGTLRQILGSGEEGSGVLAGNGNDEGLRGDVEPQWVVKYTAVVKDRGVVESKGYFAVAELRDS
*

>g125.t1

MSGGPSVIRRPFFEKSNTELTSSLRANFAAHQPSPTPSKTNYATWTTKTDSALYVPTHTPSPLAEPRESYDITVKLFYLP
NIPTDRRCAQTREAIELVLKELGTSSIDLLIVSFPGITFDADDEDSDLDDDDPPSPPSATQSTSENTAGDCPEAGAPPED
IETMVTTWRSLEKLHAEGLVSKLGIAEFGVARLTKFLEQTKIKPSVNQINVRDCCVVPKPLILYAKQQQIELLTHNDCT
NILPRGTLRQILGSGEEGSGVLAGNGNDEGLRGDVEPQWVVKYTAVVKDRGVVESKGYFAVAELRDS*

>g286.t1

MGSKVSKVKEQASNGKLQPSQERLDKEEMKEPDNYYTHRSKSMRRKAGKTGASSGAGGAQGGTAGGAGGGGIA
*

>g287.t1

MLFFAKSTVYSGSPSGPSSAQVKCLIEERYAQTPQDQAPETTLVNVVLQTQNRRQHHQSSSFPNLAKHKRPTSRYQT
LPISKSSRNHLRHPCRQKPHQTSLKLPKLRAHPINPSPNRIQHRL*

>g310.t1

MISDYNLNAMITLLTPQRIPEMSAYLQSLHVLREVRTDVKSPQALFILGTPNHCDNTGMPAISYELNANGQVERKHY
CVNKGSGIARLLFVASIREKLLSLLMGSNATRFIYNIENRTISPRLPEPFSINRQFRELALRKFSRVHAVGRIVAYNAQAT
FDDLRLMYDWCMASRLFIRSEHAARHEKYPPTILMRSELSERDNLSNLRIPITNMVVATTKLPFNTEIRVELVHIDLPV
SVKVAVKTVTLHRLQQQFLVFLTDILDIYPGQSTQVCPCVIMDGNFAIKEAECKREDGRNFLVKNKSTALDSDEMEEE
VDLCMNKLLERDDPVTAAYLPHREVDDDGHQTAYILDSPHDHSLLGVAVWLACCL*

>g311.t1

MRRVQDVRGLMSVIVDFAMGEVCGRNGIISFQQFVHAQINFLFHLVRVKSGTLILDKEIPSILSLTLCFLDCKVTIHYD
TWTDLSTLAWVDVQDISKKDKELLLKPVKGYCFDRNFNANRQVDMN*

>g704.t1

MAMQRPVTPSASYDPVPHIDDPAHISRGYGSSSVNDFANSPAGAQHDRSRLGRFEEDFDARTRGSSVLDGDMHQ
RSSSRSSTLNQGATPSRSGTLKKKSSVKRTGSMKRSGSRKSMHAGSIRGVTIDDQERGYDREDSVFYTPVPTKGTPT
EILADRFQTWRKFLKDLITYFREVAASYEHRAKSLLKVSNVINNTNAPAALLVEGGLNDANRILRDFHKQAILEANKAR
DVEADVINQLSGLRADLAQKIKEIKSLSGDFKNNVEKEKETTRKCVTALEEALALVDSDPTAVAGKGDPYVVRLGVER
QVERQIDEENYLHRAYLNLENSGRELESIVVGEVQKAYNALASILKRDADEQYNTVEKLRNGPIAMPRDLEWSRFVR
SDPHFVNPDMPLRRLEDIQYPGKYHPATTEVRAGMLERKSKYLKSYTPGWYVLSPTHIHEFKSADRIYTQPPVMSLY
LPDQKLGSRSQPGSSSHKFVIKGRQAGSMHRGHTWVFRAETYETMVAWFEDIKALTEKSGEERNAFVRRHASVRS
TSAGSARSASSDGGLEEDEADAVPFSANQSMKEQAIRDQSPRPSRPSPGGRFPSDLMVNRNLQAPLSPSSGSSEVG
NDLTTMSGGPPQEVHPMYLAQTAPYQPEPVQPVQYTQPIQHQSQNNFVNPYPPTQQPYDPLTTDPSSYAPLPQTY
PVQPASVNQPSLEHAQPIQRHDSNNYGNWMAPAVGGAATGVLANEAYRKKQLAQQHLDAQHQQDQTQQYPF
DQSRAAGFPDATPTQVPERHPDHILPDQIDDTGYVAPSAAPAVQPTSAPIVAPISSGQTQPHHTSSPDTLATTSSFLG
ESEVGAAPAFGKTVNGGPVPVDLVDAADEIVHPGMGKRINTDISVSDLHVPGEYPKTTVGDATSAARMPTGTTTS
QQPTTFLKYN*

>g705.t1

MYLAQTAPYQPEPVQPVQYTQPIQHQSQNNFVNPYPPTQQPYDPLTTDPSSYAPLPQTYPVQPASVNQPSLEHAQ
PIQRHDSNNYGNWMAPAVGGAATGVLANEAYRKKQLAQQHLDAQHQQDQTQQYPFDQSRAAGFPDATPTQVP
ERHPDHILPDQIDDTGYVAPSAAPAVQPTSAPIVAPISSGQTQPHHTSSPDTLATTSSFLGESEVGAAPAFGKTVNGG
PVPVDLVDAADEIVHPGMGKRINTDISVSDLHVPGEYPKTTVGDATSAARMPTGTTTSQQPTTFLKYN*

>g722.t1

MRPKTSRKDSSSSKKSSSVKSPSNSSRSSSPLASPEPKKIQMAPQLPLYVPVEPFEIETSGVIRGTLGGGPTKTSGAVK
QRVDSFDDPREPWPQEAAYTTSKSPSPKHLNEPTPKLPTFAPTTRRSSSPAHSKPSVSSKPPVPNKPSYLSPPPLLDQ
APATLHFRRRLDKATPSSYTFASDSTRLGEIPQRQWTKPFDYEEAERLNAEAALTGYPNAPMADSKDAKKKKRFGFM
RRGD*

>g723.t1

MAVKLDHAHVARKAEEAEVSTCVSGIIEAFTNGLDIFKKLRERRRKRRSKRDTEVPDSTSTAEHQLSKSLRRGPEEVA
GKYTECYYSGIGPRFAKGDSIAHASLAEILIKLNTGLVAIIAAFLNHDGHKSGKGHLDLDYKSLTHLSDASRREALQSLA
QLYQRLSQSQLQLHSIGAPPCPRCGTSKHHDCSSRSTSPEKEKRKHSSSSRQRSSGPVVTRMSIKSRSSNEPRLVVMR
PKTSRKDSSSSKKSSSVKSPSNSSRSSSPLASPEPKKIQMAPQLPLYVPVEPFEIETSGVIRGTLGGGPTKTSGAVKQRV
DSFDDPREPWPQEAAYTTSKSPSPKHLNEPTPKLPTFAPTTRRSSSPAHSKPSVSSKPPVPNKPSYLSPPPLLDQAPAT
LHFRRRLDKATPSSYTFASDSTRLGEIPQRQWTKPFDYEEAERLNAEAALTGYPNAPMADSKDAKKKKRFGFMRRG
D*

>g732.t1

METTIDEVSASAIASPLPVRSAPKSPAPVPTSDSPKKSHILKLSTRKTPKKKNFPEGSTTESGAVVEEDPTQATPSDTVN
VAEAGIIVNTNVTPKVNGKIAQETDSTAASPDPTAASAGTTRRGLRTRKPAQQRPYYHDSQLFEDVEPTNGDAQDS
SNTSPAAQGRRVSVASISKNIDDALLASLDEEAMALLQEETEPESAKPKHFKGKGRAWKKEGSDEDEEFSIAASMKA
ASKKAAKAARMKAKGQIPKKRGRPRKSGRSEELIDEETDEDKDAVKRKRPPPRKSALSEEVIIDSSDEEEAKEMEVEE
STTPKHTPNKSYTPQGLPKYISEADNGANGGPELGAGNELEVASPSKETV*

>g733.t1

MAPNGHQASAKKGIHKSSDSSSVFWEDKHGHREFMDWKEANQNGPFPSYEEWVESQKKSGGLSLYYTALRMFE
DSDSTDEEAPPVKAKPQAAKARGQVGRPKKSASTNGNGLWRTVSPATSTAVANSEDFSPSGKKRRKARKKPLEVVA
SASDSEVVKDVGASAPVAEIATPTAPMITFNGQRKSSTRKARKKPIVRDHLAR*

>g777.t1

MGNSGQVCTATSRLFVQDTIYDKFLEAFKKQTKENTKIGSQFDADTNHGPQISKAAQQKILSYVDTARLDGAELIYG
GKQDGVPEKGYFVEPAVFANCSNDMRVVREEIFGPFVVIQSFKTEDEAVEKANDTEFGLGAAVFTKDIMRGHRVAG
AIEAGMVWINSSQDSHFGIPFGGYKQSGIGRELGAYALSSYTQVKAVHVNLGTFL*

>g778.t1

MAQSIELTAPNGTKWTQPTGLFINNEFVSAGSEDDIDAAVTAARTAFKSSAWRDLSAAERGQLLWKLGDLCEKNSHI
LATIDAWDNGKPYQQAMDEDVAETISVFRYYAGWADKVYGQTIETSNAKLAYTKHEPLGVCGQIIPWNFPVMMA
AWKLGPALACGNTVVLKPAEQTPLSALFLASLIKEAGFPSGVVNIVNGYGKGAGSRLSEHPHVDKIAFTGSTITGRSI
MKAAANNLKNITLETGGKSPLLVFGDCDLDQAVK*

>g831.t1

MQKPDIWALFQHPPCDTYTKGRVCLLGDAAHATTPHQGAGAGQCIEDSYILANLVKDANNVDELQRAFSAFDQV
RRERTQKIVKTSYEAGKLYDFELFGDDLDKIEDNYMHRMRWIWDVDLKAQLEQAQKIMKQQT*

>g832.t1

MAPTSSSSKPYSLAIVGGGISGLCLAIALVQFDVPVTIYEAAPHFGEIGAGVAFGPNAGRAMEFLSPKIYQAFLKCKTG
NADNSKMDSWFTIRVGDARREDKEGYVREGKKVGDALFEVPMHSSGGRGGVYRAHFLNELVKDVPDGVAKFDK
RLVEMNEAQDGSGDIVLKFADGSTAQHGAVIGCDGIKSLTRKWLVGRDNPASKAVFSGKYAYRGLIPMDKAVELLG
DDVARNSQMYLGYHGHLLTFPIEHGKTMNGKLSRYLGR*

>g1031.t1

MELTSTVFDLPEELLATLTLKDQPEHPIEETPPQIPTTANDSSNTAEEESNSPAKATSCNLCGLSFASLADQRNHVRSDL
HGYNLKQKIKGAKPVGEAEFEKLIGDLDESISGSESSESDEDEDADGTKAKESTLSALLKKQAKISDPEFDEFSSRKKQR
GPGKPPLMWFTSPSIPDNMSLGVYRAILSNTEQEEESHVLDALRKKQLSPKQPPKIKANEAGVPPPGIDIGPHYFLC
MIGGGHFAAMIVALAPKIGKNHTGFDERSATVVAHKTFHRYTTRRKQGGSQSANDNAKGNAHSAGSSIRRYNETA
LIAEVRELLSSWKNMIDTADLIFVRATGATNRRTLFGPYEGQVLRHNDPRNRGFPFSTRRATQKELMRAFIELTRVKQ
SAIDEAALAALDSSRKEAAAPTPIPEKPKPKKPTKEEEAATLHTSQIIPMIKRSKVPALLNYLKTNNIPPSFNFLPTNHHT
PTPLHLAASLNSAPIVFALLTKAGVDPALMSGEARTPFTLTGDRATRDAFRVARSELGESAWDWEKAGVPTAITKAEA
DKRDAQEKTEKAAESKAEADRRKAETERVRKESEAEEVRRKQQRLGKGKSLGALPVKTGADLREEEMRGLAPEARA
RMERERRARAAEERLKRMER*

>g1032.t1

MAQKSDHLLQRPLYVFDLPEELLATLTLKDQPEHPIEETPPQIPTTANDSSNTAEEESNSPAKATSCNLCGLSFASLAD
QRNHVRSDLHGYNLKQKIKGAKPVGEAEFEKLIGDLDESISGSESSESDEDEDADGTKAKESTLSALLKKQAKISDPEF
DEFSSRKKQRGPGKPPLMWFTSPSIPDNMSLGVYRAILSNTEQEEESHVLDALRKKQLSPKQPPKIKANEAGVPPPG
IDIGPHYFLCMIGGGHFAAMIVALAPKIGKNHTGFDERSATVVAHKTFHRYTTRRKQGGSQSANDNAKGNAHSAG
SSIRRYNETALIAEVRELLSSWKNMIDTADLIFVRATGATNRRTLFGPYEGQVLRHNDPRNRGFPFSTRRATQKELMR
AFIELTRVKQSAIDEAALAALDSSRKEAAAPTPIPEKPKPKKPTKEEEAATLHTSQIIPMIKRSKVPALLNYLKTNNIPPSF
NFLPTNHHTPTPLHLAASLNSAPIVFALLTKAGVDPALMSGEARTPFTLTGDRATRDAFRVARSELGESAWDWEKAG
VPTAITKAEADKRDAQEKTEKAAESKAEADRRKAETERVRKESEAEEVRRKQQRLGKGKSLGALPVKTGADLREEEM
RGLAPEARARMERERRARAAEERLKRMER*

>g1148.t1

MGGVMSSLASAKTKASLDPSSATKDVPNLSNFIACDQWAPAEQNVRKALGHYNFPLQTLTNQNLKGVQEADIVLL
SCKPHGFKTILGEEGVREALKGKVLVSILAGVTREQIEAFLYPDGPEKTENACRVVRVMPNTASFVGESMSVIQTSDP
PLSEAQYKLVEFVFSSIGRVTNLPPANMDAATALCGSGPAFFALILEAAADGAVAMGLPRAEAQMMAAQTMRGTT
GLVLNGEHPAVLKDKVSTPGGCTIGGLMSLEEDGVRGAVAKAIREATVVASRLGGEQKEFVNHRR*

>g1149.t1

MASAQESKPLTLTVLGSGTMGIAIMGGVMSSLASAKTKASLDPSSATKDVPNLSNFIACDQWAPAEQNVRKALGH
YNFPLQTLTNQNLKGVQEADIVLLSCKPHGFKTILGEEGVREALKGKVLVSILAGVTREQIEAFLYPDGPEKTENACRV
VRVMPNTASFVGESMSVIQTSDPPLSEAQYKLVEFVFSSIGRVTNLPPANMDAATALCGSGPAFFALILEAAADGAVA
MGLPRAEAQMMAAQTMRGTTGLVLNGEHPAVLKDKVSTPGGCTIGGLMSLEEDGVRGAVAKAIREATVVASRLG
GEQKEFVNHRR*

>g1574.t1

MIFGGLEIVAGGYLIHRHYRKKNDKEQLEAEAQKRRHNTFPGANPAKQNGWNTQAHHRPQHHQQQQQQQQQ
QQQQQQQQPAVPQQKYACYAPAAQPRPPIYQPQCQTQAQLQYQSQPQCRPHALPHTHSFNIPRRPVPQRKPQIII
QPSLQRTDSFATISRMPIANGSRPQDIPEQTSPAAGLSPVPQHGIYGNAGFSVSTPAFGATPTSPGLTYELATEPQGG
ASQTIDDNWETYGHGHGGGGLHYAPTVSTELGERDPPPPYTP*

>g1575.t1

MQPSASGSDIGDIMIFGGLEIVAGGYLIHRHYRKKNDKEQLEAEAQKRRHNTFPGANPAKQNGWNTQAHHRPQH
HQQQQQQQQQQQQQQQQQPAVPQQKYACYAPAAQPRPPIYQPQCQTQAQLQYQSQPQCRPHALPHTHSFNI
PRRPVPQRKPQIIIQPSLQRTDSFATISRMPIANGSRPQDIPEQTSPAAGLSPVPQHGIYGNAGFSVSTPAFGATPTSP
GLTYELATEPQGGASQTIDDNWETYGHGHGGGGLHYAPTVSTELGERDPPPPYTP*

>g1593.t1

MSEPTETGGLPLDLKTHQQQQPQDSIIHQPTLPKVENCTTSQSPNNSSKGTKAASPRRRPGTIKQAPSGPHTGSSRS
EQQPALSPTDMSSHNPSIHYTRTGRISKAKKGLKVHNCENCGRSYTRAEHLRRHQKNHAQEDALVCQVSGCGKTFF
RIDLLHRHQERHNEPGRDTPQQSPEGSPEPASIAISASIPALASPIMEVTSNPPTTSYYQPVSPMLETVPFTSQLKHPR
SQYNRSSAAVTVPMDAMTPGLWHEPYSPTPGYSSSSGYASPIASTDFPTFGSAPYHRARTPSNASIIDPSWSYQSRS
PASTTSTMAFTWGSNDKGSTASNLAYMNACSYPMTAMSIPTSMDTMAGYGHFGPRTMIQRDEEEGVILFGDEQY
GSFIPLSPSSIDLPL*

>g1594.t1

MIAIGSQYSTGSSDKKKGRDLHDRCLKLLERRDHEASTEPDRLCDFQTMFLIEILSQYRARRAAKVLSSRFDKVYHKAI
ENCRSMTPKMADLVASPSPNHWARWVELATWQRLLLSCFVLESQQRLLLARESLPSLIHDCHLDIPLPADRSLWDA
TTSTEWATAAQQHLYTPSYVDEITPRSLTGPLDTFQSFVILAVHYNRNETPAPYISSSTASNLESLLDSAPATTRMLLVA
KLVQVTPIRALVAVASESWIFSEKVATPQAFAALKTTLRTWVAQLWSTSEPEGVPVKEALKLSIKILQQAVEEQRDSVP
LEVGTDMSIFFAALVLWIITVAANTRTKGLHQIAKQQSRRQSHSESSTIFNSAWAPTTPSAPSIGHASSSSVRHTLAPS
NSQPQSPIIEATAENPLLSHAQIIINTISFLTDTLALSDNTIALRQSPADHARRQAGCVSLLLWVKLRLRGVPLEDQSGP
ADPWTNKSGDGLGELLEGVVGSIERILKKGWSDWGI*

>g1598.t1

MAESSMDTKLYVSGEDARSFVTAVLVGNGVAPENANVVAKCLVAADLRGVDTHGMNRIPSYMERIRQGVLNSAAT
PTVTQVTPVVAQIDANNGFGFLAADAGMAACIESAKTYGIGMASIKHSNHFGMSAWIVQKALDADMMSLVFTNS
SPALPAWGGKSKLLGVSPIACGAPGKDQPFVLDMAPSIAARGKIYKAKRRGEKIPLDWALDKDGKPTDDPEAALDG
GVMLPMGGPKGSGLAVMMDVFSGVLSGSAFAGDVTNPYDPSRPSDVGHFLVAVKPDLFMSLDDFRERMQILYDR
VTGAEKAAGVERIYFPGELEQIQQREREQSGIPLVQAEVDALNAEAARVGAKPLVVKLRERV*

>g1599.t1

MTPTYHLDISHPNKRICNWESFRESGEDARSFVTAVLVGNGVAPENANVVAKCLVAADLRGVDTHGMNRIPSYME
RIRQGVLNSAATPTVTQVTPVVAQIDANNGFGFLAADAGMAACIESAKTYGIGMASIKHSNHFGMSAWIVQKALD
ADMMSLVFTNSSPALPAWGGKSKLLGVSPIACGAPGKDQPFVLDMAPSIAARGKIYKAKRRGEKIPLDWALDKDG
KPTDDPEAALDGGVMLPMGGPKGSGLAVMMDVFSGVLSGSAFAGDVTNPYDPSRPSDVGHFLVAVKPDLFMSL
DDFRERMQILYDRVTGAEKAAGVERIYFPGELEQIQQREREQSGIPLVQAEVDALNAEAARVGAKPLVVKLRERV*

>g1616.t1

MSISSTEIKFENNGDLVTRTQQCRILASTCLMAFTIIGLNQSFGVFQAHYGRQASALEGVLLQAELSQRSLISAVGSLG
NGGLVAAFGLFYYPHLPRLGGHVKYLCGLGTAFITIGFAAAAGSHNLTTLVACQGLLVGIGAGILNYVLAPILPEYFPQR
SGLAQGAMFACGGLGGMVWVFVLTALLESIGIRWTLGLLSILSFAILSISSALALPPRKFERRSTEIVSWKVFRDPLFAS
LAIVNLIMALTLAIPTAFGSEFAQSIGASITHGSYLLAINSGIGIPGRICTGWLSDKIGHLNMLIVATAVYAIATWALFLSS
AMTSNLGSYVGMTVCYGLSSGVFNTVMNSAQKMLFGAEMYYPKSGATITIRGIGFVIGTPIAGALVSRIAGEDLVGR
DFLKLIVYTAALLTLSLLCLLNVRRLDARGNGWKLVR*

>g1617.t1

MSVHNSEQAIGQVQSSLDGKSHARMSISSTEIKFENNGDLVTRTQQCRILASTCLMAFTIIGLNQSFGVFQAHYGRQ
ASALEGVLLQAELSQRSLISAVGSLGNGGLVAAFGLFYYPHLPRLGGHVKYLCGLGTAFITIGFAAAAGSHNVSKVLPV
PAPQLLTMIQLTTLVACQGLLVGIGAGILNYVLAPILPEYFPQRSGLAQGAMFACGGLGGMVWVFVLTALLESIGIRW
TLGLLSILSFAILSISSALALPPRKFERRSTEIVSWKVFRDPLFASLAIVNLIMALTLAIPTAFGSEFAQSIGASITHGSYLLAI
NSGIGIPGRICTGWLSDKIGHLNMLIVATAVYAIATWALFLSSAMTSNLGSYVGMTVCYGLSSGVFNTVMNSAQKM
LFGAEMYYPKSGATITIRGIGFVIGTPIAGALVSRIAGEDLVGRDFLKLIVYTAALLTLSLLCLLNVRRLDARGNGWKLVR
*

>g1631.t1

MNTPSTILDVVTRRLHLQPDHPLTITRKLIESRFPGYKTHNDLFPIVTTGQNFDSLGFPLDHIGRSRTDTYYLNKNTVL
RTHTSAHQADTFRNNESEGYLISADVYRRDAIDRSHYPVFHQMEGARTWDRQQAEREGKTLAKVIWDDVEKIPKH
NVAVEDPNPPFHAERNPLQTGHSAEEAEAMAAHLKRSLEDMVVTVFNAAKSATDTESPNEPLKVRWVEAYFPFTS
PSWELEVFWQGDWLEVLGCGIVSQPILNNASVPTRIGWAFGIGLERIAMLLYSIPDIRLFWSKDDRFLSQFSEQKTM
CRFVPFSKHPACFKDVSFWLRSSSNAAGGAVAVHPPAGGMSNNTPTTTSGAPIPPAAPAPSSSSFHENDVMEIARE
VCGDLVEDVRLTDEFVHPKTGRKSLCYRINYRSLERTLTNEETNELHERLRGLLVERLGIELR*

>g1632.t1

MRIPLSSTALCRQLCQNASLRIPRSSVSPWPSRHAARHNSSGAFPKEIKIEGRTYQTDEWMNTPSTILDVVTRRLHL
QPDHPLTITRKLIESRFPGYKTHNDLFPIVTTGQNFDSLGFPLDHIGRSRTDTYYLNKNTVLRTHTSAHQADTFRNNE
SEGYLISADVYRRDAIDRSHYPVFHQMEGARTWDRQQAEREGKTLAKVIWDDVEKIPKHNVAVEDPNPPFHAERN
PLQTGHSAEEAEAMAAHLKRSLEDMVVTVFNAAKSATDTESPNEPLKVRWVEAYFPFTSPSWELEVFWQGDWLE
VLGCGIVSQPILNNASVPTRIGWAFGIGLERIAMLLYSIPDIRLFWSKDDRFLSQFSEQKTMCRFVPFSKHPACFKDVS
FWLRSSSNAAGGAVAVHPPAGGMSNNTPTTTSGAPIPPAAPAPSSSSFHENDVMEIAREVCGDLVEDVRLTDEFVH
PKTGRKSLCYRINYRSLERTLTNEETNELHERLRGLLVERLGIELR*

>g1690.t1

MAKELLVHGVEQSAIDAALRSVLGSDYDSKYTDSWVDLLLQNHADVNTVEGSCFIFAAQKHSHTIFDKLMHHNPN
FNIVVPALLSSKLQDEVVVAAIQSCFDHGCTLEEVGIGYNKPPILITAMTVYPRNSALLTLLLTNGLDAEITIPMVLHSSS
GPEPVPALLWALAQPQKRISDAVISALLEAGVSVTRVSPVSETTALMLAACEGRHEIVNALVVRGTDADLRDAWNKS
ALYYASSSLSGEATVKALVPRVLGNDGSLHEAVRQLNIEVVRVLLEHAHDPNYPSRLHGGRNALGELCLKATVTTSAH
RSKLRQLLRLLLRNRANPKFRARNEKSTVILALDNAYSALPVAEALLETEIWQDLNDEAHIYCDPASGLRYSPYSYVELV
VTPARAPVKQALLDLLRDKACVPVYYSSSALQPVGATGIPPSIAKLVDKQKEHELDIRHEKEKFEHSRTMEETNHKDV
LRRKHESQDVELALQTKATQHYTALEQQKHEFEVQRVREAERMKRSERVAWHNLLQEQERDAAAARQSAEERKV
REAMAAESKMIEQRKLELEHRATVERRMLKEKEEHYERNVKRQKEVRQIEGGPPQWGTVD*

>g1691.t1

MERRVSLKDLIRNAGVIPVDQRQAVLPAPTQPAVSVSEEDISQARNILLDRRAKNPESKNVLKSIFKSSKEKDKGQDA
GQFSQEELDQALSAVIRSPTTGPGLIQAFLTLGAKVNIIETTDKKRRSSNQANTALRRRSTVLQQAASLRKADGVNILA
SSGADQQTLDEGLKAALTANDQESIGLLLRHGADLNNFPNALANAVRSNDLNFVKLLLRAPKPLRPQIISSCLPAAVQ
QSSDAIISLLIAYGADPNFDSASALNMAIGRQAYKIAIALVAGPIPLNEPNLQHALDTTMRLPTRQATLQFLQLLFCCG
LPPDSRGLPDFLIFLARSNDSSGAKMMVSYGVPTTTNDAECLREAIENSNWGLVDVILQTPIAPQHATVALSVLPTN

TPQSDRLRVVDALLKKGATGPALSPLLTQATKERDAQLIDLLLSAGAPVDVSDNSALHFAVVNRDIRSLRSLLNARPPP
ESLANLFPLLFCTDAISISERREIARLLLEHGARGPGVDQALIDAIADTSAGRDGALITDLVRGGANVDKRALSLAVTQ
VDMSLLRLLCNTRPNSSSTSAALPLAFDSQSSRHSKTLEVIDLLLSHGVEEEPAGQALQIAINGGPDNIDIIERLLTASPR
LLSTAFKYTSALQDPQKKAPILSALLKLGVPQDSLDQALVTETRQAITTNDTTSTRLLLGHGASWTSPTITI*

>g1698.t1

MTSMRSITEYLGGANLCPTTPHPITGETVPLTFGHEFSGTVEEVGDGVTDYKPGDRVVIQPIIYDDTCGACEEGLQN
CCWSNGFIGLSGWGGGLADHIVVPTSTLYHLPDNVPLEIGALVEPLAVGWHAVKISPFKKGEVALVLGGGPIGISTIL
ALKANGCDRIIVSEVSRKRQEFARKFGAHYIIDPTKEDLAKRCRELTGGKGVHVVYDCAGVQAALNQAVHATRARGC
IVNIAIWEKPCTIFTNDFNFKERTYMGIATYEIGDFQEVIDALSRGAMDPKEMITRRIGLTEVEEKGFKSLINDKDNQV
KILVEVGGGS*

>g1699.t1

MGRNLRIRYEPRCEGRSTETDVNLYLHEYLGGANLCPTTPHPITGETVPLTFGHEFSGTVEEVGDGVTDYKPGDRVVI
QPIIYDDTCGACEEGLQNCCWSNGFIGLSGWGGGLADHIVVPTSTLYHLPDNVPLEIGALVEPLAVGWHAVKISPFK
KGEVALVLGGGPIGISTILALKANGCDRIIVSEVSRKRQEFARKFGAHYIIDPTKEDLAKRCRELTGGKGVHVVYDCAG
VQAALNQAVHATRARGCIVNIAIWEKPCTIFTNDFNFKERTYMGIATYEIGDFQEVIDALSRGAMDPKEMITRRIGLT
EVEEKGFKSLINDKDNQVKILVEVGGGS*

>g1700.t1

MPYPETTDAFAVTDIKNWSTFKRQELPLKKFEEYDVDIAIDACGVCASDVHTITGGWGEELPLPLCVGHEVIGKVVK
VGDKVTRVKVGDRAGIGAQIGADLTCDQCKNDQENYCPNQVDTYGAKHTDGTVAQGGYSSHIRGHEYFVFKIPD
NLDTALAAPMLCAGLTTYSPLKRLGAGPGKKVGIIGLGGLGHFGVLWSVAMGADTYVISHSPNKKEDALKMGAKDF
IVTKEKGWAEPWKFQFDFVINTADATDKFDLSEYFSILKVNGTFHMVGFPDNPLPAMPAQVFAPNGCYMGASHIG
NRPEMEEMFELASKQNIKSWVQEIQLSEEGCKEAVERVYKNDNVKYRLTLTGFDKVFGKRA*

>g1701.t1

MARVSTLCDCAVGVLGTVCVNLVGAVILLVVLALVAGEIGTDLSTNTGAVSDLDASDLVTNLDDLADNLVSYAKRKRK
LLSPSSSDGVDI*

>g1720.t1

MRSSFDAGRPVPEIPSMMSSPAVLDMPSSQHQSTMSHASLTSGQNSGYSLTALSMAAEYQALQGNMANRLSHD
AMQTPMPSHTGTIAASVEVPPSAMLTEPSGQSYESTFGESLDSLTSFLDSEPLNSYHFASWINTEQPMPFFSPDSFG
YGQQALTEPDRTTAPTPGWRPTQLEEPPSLSRFGSRFPSLQPEDQPSRRRSFADISHQDRQSIIEMLHQFSTVIPSDF
VLPTRLALCRYIAAYINGFHEHMPFLHIPTMSVETCSIELLLAIAAIGAQYTFEGEKGVELFNVSKAIATQRIRKRDSRLV
QLQHHSESDCSSPRNQQNDAQSPRRKGSVSGPLGLPSDVNGPSLGEDLMQTAQALLLLMALCTWAKHKEILREAL
AIQSILATLIRDDGLETESLQENISWPDWVRRETTKRTKFIVYGFSNLHCIVYNIPPFMLTSEVKLPLPCSAAEFKAPTE
ALWREARKKGAPEVLFQDALKRLFTKDGRDVTECNSSSGNYALIHALIQHIFFLRQVARCRFEGPGDLSTEDVASLEN
ALRNWQIGWRQNPESSLDPMSQNGPVAFNSTALLRLAYIRLYVDTAGRALETRDPLLIANAFRAGPAIRRSPKITRAV
LHSAHALSIPIKIGYRLVAKTQSFIWSIQHSLCTLECAYLMSKWLEALSVQEADPPVTEDEHKIIAIVKSMLDETEFAIPA
DVTPGSPNFTRHLSAGVLRVWAMIFKGSQTWAIVDVIGSSLNLYADMLESG*

>g1721.t1

MEAATSKEQVPQAQPGKKEKRFQCPHCQRAFARLEHLQRHERIHSGVKPFSCSECNYSFTRSDLLVRHERLTHRKV
QTTQQNHQHTPAESTYSSVETRPHKRMRSSFDAGRPVPEIPSMMSSPAVLDMPSSQHQSTMSHASLTSGQNSGY
SLTALSMAAEYQALQGNMANRLSHDAMQTPMPSHTGTIAASVEVPPSAMLTEPSGQSYESTFGESLDSLTSFLDSE
PLNSYHFASWINTEQPMPFFSPDSFGYGQQALTEPDRTTAPTPGWRPTQLEEPPSLSRFGSRFPSLQPEDQPSRRRS
FADISHQDRQSIIEMLHQFSTVIPSDFVLPTRLALCRYIAAYINGFHEHMPFLHIPTMSVETCSIELLLAIAAIGAQYTFE
GEKGVELFNVSKAIATQRIRKRDSRLVQLQHHSESDCSSPRNQQNDAQSPRRKGSVSGPLGLPSDVNGPSLGEDLM
QTAQALLLLMALCTWAKHKEILREALAIQSILATLIRDDGLETESLQENISWPDWVRRETTKRTKFIVYGFSNLHCIVY

NIPPFMLTSEVKLPLPCSAAEFKAPTEALWREARKKGAPEVLFQDALKRLFTKDGRDVTECNSSSGNYALIHALIQHIF
FLRQVARCRFEGPGDLSTEDVASLENALRNWQIGWRQNPESSLDPMSQNGPVAFNSTALLRLAYIRLYVDTAGRAL
ETRDPLLIANAFRAGPAIRRSPKITRAVLHSAHALSIPIKIGYRLVAKTQSFIWSIQHSLCTLECAYLMSKWLEALSVQEA
DPPVTEDEHKIIAIVKSMLDETEFAIPADVTPGSPNFTRHLSAGVLRVWAMIFKGSQTWAIVDVIGSSLNLYADMLES
G*

>g1740.t1

MVACAQCRQRKVRCEGGPVSCDACETPSCRRCRELNLPCNYTLSVRQYRAGERHKRSLVSVEQVVNQSGGDDVG
RATHGTPTTGSEVTASPDTSSTEHIDKILGCDRAVIRQHIDAYYDYIYPVPIFSFLHRAEFLGQYTAGIVSPALLLAVCGV
SSRFLPSARERAGMIKSWIEQAETIIFQNMGKMNIADVQALMLLELHCMYNHQNGKSFTYISLAVRMAYLLKLHKE
NSQLPFVEQESRRRLLWCMFALDRLHAGGVPEYILLPATSVQVQLPCPEHFFQIDTPVATPHIHETQADNSKITPAAF
LLRIFNARNHVLQYTKQLLDASLSPETSLAQYQMLEGELREIHESLPAELVFSTRAYQLRTFSPERTTFTILHLYFHHCHC
ELYRLLNPGYREALPQSVIYSSSPELVAYAQSKCLEHAISIGEIVASTYGLVEVSPYVSDTSCFVILYQASCAILYACHRDSP
AYVMKAETARRYLVAFITTLQSLLSYFPRYAIYVKDIRNMLRSIDEPNVPLPAQKASNEVDFRARIVPSEENSDEDNLVS
EIAVSTSNVRDDTAHSTSQTAEVPVGPLPELVEMSLHTPSTLEGQDHSFADLETYQYNLDLTMDLDQGLLWDWADA
LGSGFSI*

>g1741.t1

MKAETARRYLVAFITTLQSLLSYFPRYAIYVKDIRNMLRSIDEPNVPLPAQKASNEVDFRARIVPSEENSDEDNLVSEIA
VSTSNVRDDTAHSTSQTAEVPVGPLPELVEMSLHTPSTLEGQDHSFADLETYQYNLDLTMDLDQGLLWDWADALG
SGFSI*

>g1767.t1

MGHLATSGQKILRVWGFNDVKTIPGSGTVYFQSFSGSSATINTGTNGLQRLDAVVKAAEKHGIKLIINFVNNWTDY
GGMAAYFSACGVSSNAQWYTAARCQGMYQSYIKAVIARYRNSNAVFAWELANEPRCNGCQTSVLTEWIRKTSNYI
RSLDSDHMITVGDEGFGLPGDGSYPYQFGEGIDWEANLKVGNISFGTFHLYPDSWGVSNAFGDGWIKAHAQICKK
LNKPCLFEEYGVPKKEDHCPVEGGWQKTSLSLKDSGMAADLFWQLGDTVKSTGQLTHDDGHTVYYGSADWKCLV
DDHVKAIG*

>g1768.t1

MKLLSILSLCAAAAALPATFSLSPRASISKADGLKFNIDGVTKYYAGTNSYWIPFLTNDNDVDVIMGHLATSGQKILRV
WGFNDVKTIPGSGTVYFQSFSGSSATINTGTNGLQRLDAVVKAAEKHGIKLIINFVNNWTDYGGMAAYFSACGVSS
NAQWYTAARCQGMYQSYIKAVIARYRNSNAVFAWELANEPRCNGCQTSVLTEWIRKTSNYIRSLDSDHMITVGDE
GFGLPGDGSYPYQFGEGIDWEANLKVGNISFGTFHLYPDSWGVSNAFGDGWIKAHAQICKKLNKPCLFEEYGVPKK
EDHCPVEGGWQKTSLSLKDSGMAADLFWQLGDTVKSTGQLTHDDGHTVYYGSADWKCLVDDHVKAIG*

>g1811.t1

MQYCHEKQMVVFGRFRVTCTVDTWTVTTTTNGHFYHSILHFYTHTAFYPALCTWLVTTHALSFLLDTTASSHANLTS
SKGGAKIANFILLEEKAIPPSPAKPKLRRRNVSICRRIPIPRLFHQIPERRAIRNDQMRRWVPAAVFVLGAETQRCLQE
MNSWIRRSAASASAGGGYWAVCKWRVVRRNTRFDDQTSFRRLSINDSINTVCRILV*

>g1812.t1

MYDHDDLNSPSHQPSVVSPCSDRPPQFNWRNCGYDCGEGCEKPSEDEGGAGVSDEETAVDADEDQSSMTESQD
EDVGMAHDTNDSTSGFSDQDSTDCINTVVYGQPPEGSLIVEPCVTADHSPFAHSPVTSTGGSTRSRSPDPGILYWIS
SMSFATPLPIGIEYHFLQTPLRLSPQYEHCRGDPASHLVVPDRSPLWYLVEQSWNGDAPAYGYISPP*

>g1848.t1

MRRGLDSRYMISQLRPTDFKILSLTTIATPHRGSAFADYMFQTIGPRRIKRVYSVMEYFGFETGAFSQLTQEYMQNSF
NPKTPDIPDVRYFSYGASLEPTRWSVFAPSHAIIKQKEGVNDGLVSVQSSRWGDYKGTLIGVSHLDLINWTNRLKWF
FWELTGSKRNFNAIAFYLDICDMLAKENL*

>g1849.t1

MIRICARVTTRSSAYTATPFRSSPTYTARQFSSSQWRKEDPRISDFGREIVDEFAVMREKYDTPKHTIVLAHGLLGFDE
LRIAGQFIPGIQYWRGITDALAHKGIEVIVAAVPPSGSIEARSAKLAESIAVKAKGKQVNIIAHSMG*

>g2480.t1

MEQPGDLGSLSFFSSMMYARAFMGQNDIRSSNLNIFSDNSLLRKTIHPDRLSIPDFATSMFPVISRMFPRAASSPHL
VILGNSSAFDSALKNISHPSTLQLVRQQSIDHQIIYTIFHRLINDSDAFLVLDEPKTELDQVFRTGILYCFSLGKRVLSGLI
DSTPSPFDLALKQNIFRAALVLDNGTVLSDVLEMSQLSLTNRRLTVEGDDYYPLEYAGLKGRIQATQALLDHGADPNL
QTNPEDFLRKILGMPWEARNPRIGVQILRLLIDHGLESSPRAFVNRIHISNGDELSVLATHCLDKSFETFFHHRGLPKV
LLQRHWDDPSSKTLKAILDKAFMESNGQQRLWNSMLTDALSAAVLRNHRSAVELLLSMGATPNIHCLISAAQSND
VQALKTFLRHGLNPNTEGKEATLPHIYDARNNWDREEDCTALSESIINPSRGVFQVLQEQGFVLDLSRHPAGFASAF
VAACQVGDSALIEQLLSLPNFPRTQGKLVRAVEFAMKGDQYHIIRRLLSVGMKATPKSLELAIQKKQLPIVILLARHVD
VAKDLVWAQPNEQANSIIWEAIRWGDQTAIEHVVRAGYPLNVCDEMTYGNLRDWELLPGVKAPSSDGLWHYTPL
GAAILGKDTTTTRILMAYGVRAVLFNSHSTSYQAAGHYATNDFLSVWVITPLAAAAVADDLSLIREVLRMGADPFDN
SALFICAVIGSEEEVVKLLLFAFKTRYPNGAHSFGSDALYRTITRGNVQLLKLLARDVDITGPVLTEHYHEIPRGAPIETIF
TSPLGEAVRQHAKNKGTSGAFDHLLPLVKDLNAVVHRTHKHGNMTSLLYAIFLGSLATVQKLHQAGADISLPAEWQI
PRTPLQAAAQAGSKDIVEYLLHHGVNPNEAPAERSGATALQLAAISGNIGVAAVLLEAGAKVNAPPAFCDGRTAFEG
ATEHGRIEMMIFLVRHGADLLSNNGAQYRRAVDLAEDNLQPVAKKLATDLYEQLLASQAINSIGMGGDAWAGLDIS
SFGGLRA*

>g2481.t1

MADNRYSQRIPRRTWDTHKHNILRLYLLENRPLNVVKDMMHSEYGFSATKSQYETRLKKWSCYKYAAAHDRRAKA
SVLDDPRTEGLVTERYMQGPGSDASTEIHSFPRNIPIRTSSPHHSTVQSCPLSHLSPRISSSAAVDDRLIFPSFSVNSVD
RQPSTSQHEFESLVSQDPTWASNTGDAADDLFHSALHTSIDVDTTSASSVPSHDMEQPGDLGSLSFFSSMMYARA
FMGQNDIRSSNLNIFSDNSLLRKTIHPDRLSIPDFATSMFPVISRMFPRAASSPHLVILGNSSAFDSALKNISHPSTLQL
VRQQSIDHQIIYTIFHRLINDSDAFLVLDEPKTELDQVFRTGILYCFSLGKRVLSGLIDSTPSPFDLALKQNIFRAALVLDN
GTVLSDVLEMSQLSLTNRRLTVEGDDYYPLEYAGLKGRIQATQALLDHGADPNLQTNPEDFLRKILGMPWEARNPRI
GVQILRLLIDHGLESSPRAFVNRIHISNGDELSVLATHCLDKSFETFFHHRGLPKVLLQRHWDDPSSKTLKAILDKAFM
ESNGQQRLWNSMLTDALSAAVLRNHRSAVELLLSMGATPNIHCLISAAQSNDVQALKTFLRHGLNPNTEGKEATLP
HIYDARNNWDREEDCTALSESIINPSRGVFQVLQEQGFVLDLSRHPAGFASAFVAACQVGDSALIEQLLSLPNFPRTQ
GKLVRAVEFAMKGDQYHIIRRLLSVGMKATPKSLELAIQKKQLPIVILLARHVDVAKDLVWAQPNEQANSIIWEAIR
WGDQTAIEHVVRAGYPLNVCDEMTYGNLRDWELLPGVKAPSSDGLWHYTPLGAAILGKDTTTTRILMAYGVRAVL
FNSHSTSYQAAGHYATNDFLSVWVITPLAAAAVADDLSLIREVLRMGADPFDNSALFICAVIGSEEEVVKLLLFAFKTR
YPNGAHSFGSDALYRTITRGNVQLLKLLARDVDITGPVLTEHYHEIPRGAPIETIFTSPLGEAVRQHAKNKGTSGAFDH
LLPLVKDLNAVVHRTHKHGNMTSLLYAIFLGSLATVQKLHQAGADISLPAEWQIPRTPLQAAAQAGSKDIVEYLLHH
GVNPNEAPAERSGATALQLAAISGNIGVAAVLLEAGAKVNAPPAFCDGRTAFEGATEHGRIEMMIFLVRHGADLLSN
NGAQYRRAVDLAEDNLQPVAKKLATDLYEQLLASQAINSIGMGGDAWAGLDISSFGGLRA*

>g2670.t1

MTTSSLITEWAKACHADQLANYPPDNVPTQLIVILDRLLSSRTSPTASAAATANLIQSEHDITHGLSCLIGLFLFAAGQ
NTSLDSLELLVSYLVELAKQPDAINEGPEPKVWDEGGGVMHEIPSGAPIIVEGK*

>g2671.t1

MLSWNITEWFQGPELWTHSHYNPTTPEVAAAKWMDMNKLIAVIARNLDAQVLPSLAGHINLSRITLSMALEHSP
NTRLGKNTAMHLPAAALWFSVAGGELERMCEAGEQKMAAGDVWAERVRDNGSGDSEVVDATRLQFWKERLAE
LQQM*

>g4436.t1

MTANLDTRRIRSATPAELRDYYNQILKDAPIGEVTTQLLQVVESGSIPPITFAPWLGVAKSPSVIREALTQDVSVLIRKF
AIKQLRKSLCSSRWKETWEGIGGTAGVLDIFAGLSVLEVRSACTAIGSCGKGTDLDEKRELFTQLYKALHPDRFPDSPH

KTKDKRALGRFYASLIPACSADLVAEAIMSGFKGTWKNTRGKYVIKYYPEYMQKEILQSLGAEESSPVDISTFRGLLHH
YPPVKSTTTGFSESMEFSLMVLETLSNSPRRVVEDRVFVDQLVNPLLKRAIKNRTDWSITQRIVNLTMHYLDMHPIK
GGQRSGLQVDFLQQVAFCWSQQPSLFEQQLRRICSHPVFGTVSKTAIGDWDNVLTGIQAKQSYPLLRLCYQASTSL
DLDSDEDLAKTKGSIHPEFFSNMSPEDSLLLFTRLRRVRGDTDLIYLGYGGGSIFSLASTFDGSSGDPNIIHIWLLNLNN
KEEEAKTMAKEYIDMRKKKAATASQPDQRAFFAKSAIFATIACGSLWMLQHTMKWTNRFLGDPTVIREINLQAPTN
EGVRLLSGVPELINKRLSLTELQQRVYAANSILQDMFDTACEALRQPSFQAYNWYGVFELFYQVIKLRIDLTPMVKKQ
LDASDEDVLTNLWADTISMVIAVEEKAQKDGYERLQANLFRGIIAYNKPFTYELESCDVSTYAFLDSLARARDELWCR
LRPLTYPATVTLPHPFPRGLPLQYLTAPWTLNVEDLSSVAPYIASRTRRTVFPDSEAALQPVPTDKDSQQAIGMFVDS
YQHALQLYIPKDRDRVETQTRVKRAWEYATGPLSSKRMAEDEAIRFWKDKKPRLMSEWPPQGAMSTVEKPWPLIP
DDDGSGEPCEWNPFSSRPDFPKRELDELTYVDFSVGISENSTSRPPVNLPVISSAEVPAYKDDPRQIWSDTRGMGEG
GVLSALLYLDTKYVGDNRLLATPFPSTTDVRYPSLFLDDDFLNADEPSQYTAARAIRGHLDTIPPLLLAQSSHNLDKAL
NKFDADSEKEINANPHEVSMELITRLGESDRPGLAMQSVIRTILDRPESSSWHRKLLKPSLLRRLPAAQAEYLIEAFTD
ELIHMLRTKDKNKKSSDHQKVASSDKFHVKVTTRKMLAQLLQDLDLVGTDYALSILRKLSNVDAHVDVRLNVIKSLL
ALLEPGSQEQSREVFTILESFIPLAGALDEREALNEARWVKCEETLSLPELQAGIMGTSSSDSPILVALVTYLRSGKIPTT
EIQPFVDRIMMPILQTLRDQTARWVALFLRKHGFEDILLGDFNIPSIPRDPSVNRTLLSMGTERLGHIPRTILEEYVVTY
MSFRAAIPAPLQAMNERLVADPALRSLPEVEAWMRLYGKSLDVQSEGTDIVGLLDHVMEDSEDPAITPKSIQEQFLK
LFTALLWNDTPAYTKLTRLCDRLMLGGYLGRTWWKPYGKPILEAMVSYIDTLRTREWGRDPARNPSVLPDTFSWRL
RLLDYPWPSLDEEKASREERCKVFASQLVAIADELSSLSIYHKQLDQLSTYIATDASTGSQRAEISNNRVLTATYVGDIS
NTRLSWLTAPELLRVEIAAMLVTRVGNQLWEANTTVDKDIRGRLKAVVETWKACENEGVRRKGWEIEENYLK*

>g4437.t1

MATSWLAKTLHLSSRDAFSSSRDGHGIASPLSRAQSVYVGHHRFQDWLSVWLPPRPSEIAAKHKSITQPF*

>g4473.t1

MTTTEDSGRTQRFAHIFARLQELQNYGWDPAIEPFHSSYDNYHFFGYEKTAPKQERPQSRGRSTVASPTLSHSPDSA
TRSRPTVTRTHRSSGSDASVQTHRTNTTVKGVAREGRPVVCRVSAQTLRLEREFQLAKLVVKESDPECRHFVRPIEFV
RLSTKAGEEPLVASIFEAPGPNYLHDLVNFGPNFWKVTSNAQSHQPNPSIPNRGVPLLTFLDFAVGSVECLEILHHGH
EIVHGEIRGDAFHFAENGIVKMINFGSGARSFENGLTSAGWSTLSKETGIELKLAFISPEQTGRMPAEPDTRTDIYSLGI
LFYTMLGGQAPFEGATALDVMQNVISKRIPPISSRRLDVPEALSEVIQRMTQRNIEDRYHSTSGLKHDLARIRELLCE
GDVEGLKAFRVGSKDISCFFNLPLKLIGREKEKKTIVDVLERVSRHRRSSIKTLQSLSSGSSFSDQRRDLSFDDVVSESTS
SRGSDSRLNSVSADTPVFMGDARSIQHDSQDSVAQSEVSTAEGSVDGRPQLQSANRGRSNNSIESSALHSRSYQSN
DGLRTLASTRKLRRKARCEVIVIAGTTGLGKSRLVQSVQSTARSTGYFAMAKFDPAKKAPFDPILRLMSSFFRQICMST
YSETQLFPSEQPLTTNFYSSQRGRCYDRVSSEAAPLS*

>g4474.t1

MDRRASSPAVHCGSAGHTAEAWLRSGGASKSSRFMNVFIDVLRLLAVHKLCTWSLEDVQYADPESAELIHHIVQAR
IPLVLMLTYTNEETLPRELIPLLRHATKVQLLPFSEAQTADYVAETLHRDHQYILPLVAVVQEKSRGNLFYIREILDTCYRK
RCVYYEWRENNWVFDLDKVFAVFESPEYGSSVTTDFISKRLTELPSASRKLLAWASLLGGTFSFELLQKLLDPANKPA
DTGRLPLFDENECAVTALDAALKAYVLMPADQDARFRFSHDRYLTAAVNSLDSDWDTQLMHFTIAKMITAGEEYND
DSTIGSKALYMQSRHICLAAELIKAKETYRAPFRDMLYQAGETACESGARSTGIYYFAHSLMLLQDDPWDDNQPDVS
YQETLQLFVRSAECYWHQGMFDEALSLIRTTFQHARDPCDMASSFILQSRVFAIRGDSFGAFQALKDCLSLLGSPIPP
TTWEACDTEFQQIYGKLQNIDKEELLKRRASPDDRVLMTIGPVFIELLSAAFWSNSLLFYQSTLKLISIHLEKGTMSQV
ALAYVHLGTVAGGRFNMMQFAVDTGAIAKRIFQMFPDDYYTLGRGQTLQPMFLGHLEAPVGDLIPSLEGALQASLT
AGDRILTLLNLGVQAHFRVMASHDAVELEAWVEETPLDMRNWQKDMRGGVFLMAARQYAKALQGKTDTGSPSL
LLSDDDHNEAEYIDFLEQTASNPKRPKSIYLATKLPLLVLYGHNREAVALGEMLLPMLSSLWSERLNYSVRYYLSLAYM
VLLRDEPEHSRRGGMIQHVQGTLKLLEACCTVTDANYRGWIHLLTAVLADINGDPQSTLENYEAAMDHNERYDFIL
DEAFALELYTGWLVNKKAYRAARHALKDCLSTYRRMSAYGKANHFASRYEWLVHSGRSFTTADQAVQTSVVDTGNT
AFRLEQNDHDHFLGPETAVDRTQDWIVPETRRQESAPALHNGLSAVGLDMLDLSSILESSQVLSSELVVDKLMAKM
SSILLESTGGTLCGLVIEDSQIEWSIACVATNEPDNDSGFSSGVTSFPASQPLDTVDDVVARQVTLYTLRFRETVFVQN
LYEDDRFSNVSDAYLQRNPEGKAVICIPILHSDHLLGSIYVEGPPNSFTERNTQVLRLLVNQISISLANALLFKEVERVSA
SNEAMLEMQKRALSQARAAEIKAKEAEAIAVRNMKLKEEAAKAKSLFLANVSHELRTPLNGVIGMSELLKASPLNSE

QSGYADSIRVCADTLLSIINDLLDYSKLEAGKMNVLEMPISLSETIAEVVRALAYTNAERGLKTIEQLHLDPEMLVMGD
PVRLHQILMNLLSNSYKFTPRGSVTVRAVVDQETDDWVDVTCSVIDTGIGIPDEQKQKLFLPFSQIESSSARSYGGTG
LGLSICKALIENVMHGTVRLDSQPGQGTTVTFSLRFKKVPKAQAGNQPQRTREPDPMARFSSQGNNGHEQSSGTC
IDLSTIPRRDLRVCIAEDNLINQRIAISFVQKLGFKCDAYLDGFKTIDALERASENGRPFHLVLMDVQMPHCDGYEATK
LIRKHPNPEIRNVLIIAMTASAIQGDREKCLDSGMNNYLAKPVRAQTLKALLDSYLNKNNEVEEIPNLAIEAKKLVKEA
LNEAEALPDVTSGNRDLEKQEDRNGGASVVGETKNGITAAEREGKKIERPSSVRMSTTQHILPNGRTEPVPPSE*

>g4556.t1

MRACLSAVTLLRLGALTISATQSVHPPSSRRLPASSSAEPSRLSQQLLGGQVPLIDKSDYSGNHDPILRTGLALETLQN
WYNTSTGLWESTGWWNGANIMTMIGDFAKAAPTHIPLQDLARDVFVTALLKAPAKNPQPGIEDPRSTNTTLVAID
VTSLETGYTKYLDLNTSNLHVIYPPNWHDNGGQVIFPRTSSNYGSNSFYHPDEISDYRDWLDGYYDDDLWWALAW
INAYDVTFEAAYLVLAEDVFIALSRTWGTYCFNGGIYWSWKQDYVNAIANELFFSTAAHLANRAQDRRKRATYLAW
AEESLQWFLESGMITDDGIINDGLTEDCKNNNKTAWSYNQGVILGGLAELHRAGNIPKATPLRLATHLARAALIALS
DEDGIVHDECEPDCGDDGAQFKGIFMRNLVKLHSVVPDKMFANAIRRNAESIWTRDRKVTSDGLPVFSVNWAGP
WISPANASTQSSAMDALVAAIVTRSTPQ*

>g4557.t1

MLSTFFVRYLLFVLTFIPNISSRSDYSGNHDPILRTGLALETLQNWYNTSTGLWESTGWWNGANIMTMIGDFAKAA
PTHIPLQDLARDVFVTALLKAPAKNPQPGIEDPRSTNTTLVAIDVTSLETGYTKYLDLNTSNLHVIYPPNWHDNGGQV
IFPRTSSNYGSNSFYHPDEISDYRDWLDGYYDDDLWWALAWINAYDVTFEAAYLVLAEDVFIALSRTWGTYCFNGGI
YWSWKQDYVNAIANELFFSTAAHLANRAQDRRKRATYLAWAEESLQWFLESGMITDDGIINDGLTEDCKNNNKTA
WSYNQGVILGGLAELHRAGNIPKATPLRLATHLARAALIALSDEDGIVHDECEPDCGDDGAQFKGIFMRNLVKLHSV
VPDKMFANAIRRNAESIWTRDRKVTSDGLPVFSVNWAGPWISPANASTQSSAMDALVAAIVTRSTPQ*

>g4568.t1

MAQLAIAPARTDPLTGRCTHLSATMAGYGQIVLGFRRTCKVPDNDEKFYAPDLGPYLLYDVKKYRGLPRSILDAGGW
FIAMRDDEAMYVDFGSSRKFNVTVGSHTSRKAHCIVGWRHYLDGLPDSDSPGCVRQFVAAPASGRSQINSSDELL
VDDLSLAIRTPKDVLNQAGIIVTTRDMEEVNIPINDVTTVDQLKSRIEHEMGVPKNVQRIVFRGKQLEDDTTLSSNGI
AEGTAVHLVERMTGGGSGCLPAFFASHKMILKPGDRIWQPEAFKEEETQTSWAEIPTVIRVHILNAENVRDWTGID
PPQSTVDPERYKSAGLSSAGDKQEYERRALENPDLYDQIEEYYYGKSTIMPLREHNPFRVILPLIGTTWLYLYPVFHGC
AFPKPPSTHAQAVAPFRLLSLGDPQLEGDSSLPDPNALVFPSIENLVPNLRDAANYTTKRHVLDQAARGVVKDAGK
WLEGKRKAVDLWGNDWYLAHIVRSLRWWTEPTHISVLGDLLGSQWVTDGEFTKRAGRYWNVVMRGLEKVPDVI
FGAIESENTELPPAETSQEQEGEGEDVVEEKEKKKERRPLWGGTTEVLGADKDWQKRVINIVGNHDVGYAGDIDES
RIERFEKAFGSSNWDIWFTLPDELRSNETDPTDPTNPSPSRKDATRPPTLRLVIFNSMNLDTPAWSSDLQTETYSFTN
HIITSSLPVTDKTHATILLTHIPLEKEAGICVDSPYFDFFEGGQGLKEQNMLSDHASKIVLEGMFGMSPNKDAEGGGK
GRKGIIVNGHDHAGCDVMHWIRQPGVQDTCQVANVKREEAYWPTSTANDTMIATSMDIDGVDINFVAANDTDT
DTAIQTETDSAQEESTPESSPEPTLEPASEPPPPPKWRAQRFPNRPYDIHADDECTSINDAPHIREITLRSMMGEYSG
YAGFLSAWFEEEKGEKGEWVFEFSSCGVGVQHWWWGIHIVDLILMVCVVIGGLVWGYERVVDDARNTEVKKDV
KKGKTDDARDRNEARRIVQTTRGRTSSVMEVQERKI*

>g4569.t1

MQLSRFFFRVALLLLPIALIGTTWLYLYPVFHGCAFPKPPSTHAQAVAPFRLLSLGDPQLEGDSSLPDPNALVFPSIENL
VPNLRDAANYTTKRHVLDQAARGVVKDAGKWLEGKRKAVDLWGNDWYLAHIVRSLRWWTEPTHISVLGDLLGS
QWVTDGEFTKRAGRYWNVVMRGLEKVPDVIFGAIESENTELPPAETSQEQEGEGEDVVEEKEKKKERRPLWGGTT
EVLGADKDWQKRVINIVGNHDVGYAGDIDESRIERFEKAFGSSNWDIWFTLPDELRSNETDPTDPTNPSPSRKDAT
RPPTLRLVIFNSMNLDTPAWSSDLQTETYSFTNHIITSSLPVTDKTHATILLTHIPLEKEAGICVDSPYFDFFEGGQGLKE
QNMLSDHASKIVLEGMFGMSPNKDAEGGGKGRKGIIVNGHDHAGCDVMHWIRQPGVQDTCQVANVKREEAY
WPTSTANDTMIATSMDIDGVDINFVAANDTDTDTAIQTETDSAQEESTPESSPEPTLEPASEPPPPPKWRAQRFPN
RPYDIHADDECTSINDAPHIREITLRSMMGEYSGYAGFLSAWFEEEKGEKGEWVFEFSSCGVGVQHWWWGIHIVD
LILMVCVVIGGLVWGYERVVDDARNTEVKKDVKKGKTDDARDRNEARRIVQTTRGRTSSVMEVQERKI*

>g4654.t1

MEKLQIAEGVASQSLEEEGHPCLPLTASEGGFTPSSACMMGARTLAEQSKQGSRQLSTASLVAMACGLGGLQILWS
TIFSHGSSYLFSLGISTTQSSLIWAVAPLCGSTIQPIMGVVSDRSRIVWGRRRPFILGGVLSTIFAATTLAWSEPISVAVC
TLLGISNVDGWWGTVTRTVAILAIIILNISIQPLQLGLRSLHVDICPREQQAIASAWAGRFAGMGNIIGYVLGSLPLPW
ISSDYDAMRFRYMVYWTTFALIASSLITCYYTEEEDPSMSTYDPGLERPVYRVFRNLLDGFSQAPNRVRHVYLVQFFS
WLGWFGFLFYSTSFIGQLYVDEQARDNVTISSSIKDHGMRLGARANLLFATVALATNIALPRLSVMLRSMLARAKIYS
KGARSSSILTIIWSFSQALYATSLLSTTLVSSSATATFMIAIAGISWGVTQWVPFAIIGEETAKFSIDEESAQREDEHVWS
VVQGGTIVGLHNTAISLPQIIAALISSAIFWVAQNLNQKHAIVWVVGWTGVPGAVAAWLAFIM*

>g4655.t1

MGVVSDRSRIVWGRRRPFILGGVLSTIFAATTLAWSEPISVAVCTLLGISNVDGWWGTVTRTVAILAIIILNISIQPLQL
GLRSLHVDICPREQQAIASAWAGRFAGMGNIIGYVLGSLPLPWISSDYDAMRFRYMVYWTTFALIASSLITCYYTEEE
DPSMSTYDPGLERPVYRVFRNLLDGFSQAPNRVRHVYLVQFFSWLGWFGFLFYSTSFIGQLYVDEQARDNVTISSSI
KDHGMRLGARANLLFATVALATNIALPRLSVMLRSMLARAKIYSKGARSSSILTIIWSFSQALYATSLLSTTLVSSSATAT
FMIAIAGISWGVTQWVPFAIIGEETAKFSIDEESAQREDEHVWSVVQGGTIVGLHNTAISLPQIIAALISSAIFWVAQN
LNQKHAIVWVVGWTGVPGAVAAWLAFIM*

>g4734.t1

MAVSSTTAQPAEVVQKADTTATNTSNAAAAPRKRRRRAPATGATDDCFACKKRQVKCDRRRPYCGPCVDIGKECSG
YRTTLTWGVGVASRGKLRGMSLPIAKSPSATATQEPKSHRMNSIASTSTTSSTGAHQDYQHNFNSRASIDFGAHSPR
SPTSPMFSAPQEYQYFSPTSPIPIPTSSTPVGYPMQHHGEQFEFVHPQASKFRHHQPHRGPLQRLQTTLHVPFEDN
GMSASSASLGTYSDSDFPSPSEFPNTPVDEFPFADPSIPRYQPFDQAPMGSLDHYFIHEAPRSLPTGDDMSSSVSSD
QSIHDYPEVSAASQSMGPVVFQDVFVEGEMSSSFNGFQPGFSFLPSEEPYHRFAPFPADAIPSGLLRQIYMPGAGG
F*

>g4735.t1

MRNEELQNAIGALATNNMRMRGGIEGRRLSVLHDTRLIPDHAYSQMTTQELREMHGEATDEEKHYKANSIALLNK
KLVDYDGSQDDSVLATLLILCLFHVCDSGFTKFKTQLAGVQKLLSLRDRTVRSEFVGWIETFFTWFDVMTAAVNDREI
EVRGDSLDMMNLSANLGALEHHSGCEGRLFKLIARLGRLNLLSQNRPVRDNDDTPTSSPRPKKVKDFYSFSFDHM
DGNGWGTPINEHVDPFTSARDDPRSEFWSEWNDLRIRLQEWEFPSPGDMHSTSCEATSSMLAMSPEQAALFHIS
ESFRYAALLYTERLAQPTLPASALNFQNLVSQGLYHISQIGMTSCVNKFLLWPLFIIGTECVDPEHRAVVRQRCVEIQR
ESGFFNNLSGLEVLERVWREDDEWSDGDVQVPRQNGPFSTVQHPFRWRRAMDRVDGEYIVI*

>g5061.t1

MLVATAASAQYGPGGRYGPDGVGGGSGNDDNDFGSSFGGGRGGFSLESRHKIIIAHGVLAALAFVLFFPVGSILIRL
GSFRGVWLVHGLFQLFAYVVYIAAFGIGVWMINNIPVNMLDNYHPIIGIIVFVLLFFQPILGFVHHLKFKKHNRRTIW
SYGHLWLGRILITMGMINGGLGLLLASDAPAFTGFAPSRGQTIAYGVIAGIMWLFWVSASIYGERKRKISRKAALNKE
VDEGSPRPYDDGKEYYAYVGRCSTSLCSVANRMTGNELMLERINVVSEESSSH*

>g5062.t1

MTSTTRAMLLSLATAASAQYGPGGRYGPDGVGGGSGNDDNDFGSSFGGGRGGFSLESRHKIIIAHGVLAALAFVLF
FPVGSILIRLGSFRGVWLVHGLFQLFAYVVYIAAFGIGVWMINNIPVNMLDNYHPIIGIIVFVLLFFQPILGFVHHLKFK
KHNRRTIWSYGHLWLGRILITMGMINGGLGLLLASDAPAFTGFAPSRGQTIAYGVIAGIMWLFWVSASIYGERKRKI
SRKAALNKEVDEGSPRPYDDGKEYYAQRTYA*

>g5214.t1

MVSRGGRSKGCTNCRRRRVKCDENRPVCGQCTKRGLECEGPKDLTWIDQSRTFEAKPTNTAQNAVVKIASPTELSL
KAFEDDICLAYTRKTLLRGGPVEIACNMVESYGASIESDNPGLDLLRKGILTLSVTFFGSQHRQDYITKKGYSQYSEVLR
QLNSHLSHPELQLTNETLLTALSCMLLEIFLPTGPKNFLKHQRGLDAIAMLRGPPKESDGTTATIFRGLRILSIVGSLAES
RTSLYSREEWKQVPPVQASEAGYFQHHVFTILADCTRLIGRRNALLASKAIPASFEPLLVEVEAVLSALEALYPHWEVL

NRNQLAGVTEQSDMAKTLGVANHVSATAYMLYNTAYICVLQIKDSLSPSPINVAWRNAAATSIAKCLELKEYEQREG
APQSNTIAYVAIRVAWQALGGFDSPEGRRLAHVVDSATNSVFQQPSLPSDESLFSRFVERMPVVLPDCTSYPNEQY
DTWSEGPPGIIEFSMQQHLA*

>g5215.t1

MCPFAYSTFTESKPGACADKKDENRPVCGQCTKRGLECEGPKDLTWIDQSRTFEAKPTNTAQNAVVKIASPTELSLK
AFEDDICLAYTRKTLLRGGPVEIACNMVESYGASIESDNPGLDLLRKGILTLSVTFFGSQHRQDYITKKGYSQYSEVLRQ
LNSHLSHPELQLTNETLLTALSCMLLEIFLPTGPKNFLKHQRGLDAIAMLRGPPKESDGTTATIFRGLRILSIVGSLAESR
TSLYSREEWKQVPPVQASEAGYFQHHVFTILADCTRLIGRRNALLASKAIPASFEPLLVEVEAVLSALEALYPHWEVLN
RNQLAGVTEQSDMAKTLGVANHVSATAYMLYNTAYICVLQIKDSLSPSPINVAWRNAAATSIAKCLELKEYEQREGA
PQSNTIAYVAIRVAWQALGGFDSPEGRRLAHVVDSATNSVFQQPSLPSDESLFSRFVERMPVVLPDCTSYPNEQYDT
WSEGPPGIIEFSMQQLVIHPRF*

>g5216.t1

MGEPDDPSIRSFKSKAASALGLGRKKDTQTDVLPTHNPHTTAHPGLGGSNPPRSAPGTTNSDHSDHHEKPVAPDD
NSTDVQHGSVNTAADGETPSAIGSEKRPFYSPTRVKNGTLRFITHTKNALTHSWINVLLVFVPLGIVVKLVDLKPEIVF
SMNAIAIIPLAGLLAHATEVVAARVGDALGALLNVSFGNAVELILFIILLASDQIEVVQASLLGSILANLLLILGMAFLLG
GLKYQEQVYNNTVTQMSGVMLALAVMSLLLPTAFHAAFEDNSIADHETLSVSRGTSVILLLVYGLYLLFQLKSHRYLY
ASTPQHIIDEESHPGVLAGAFDSSSSDSSSSSSSSDSDSSSNNSDGTMKKKAKRMVKRLRRKSSASSKDDGALSAMS
SPSAELNQSPFETERTSSVVTANNAGPITSRRHSFDVMSGDEADNDQYVPVVRDFEAASHTSRSLTKKEKRKNKKSK
KDRRDRNKIDTIPEKEVEPNDPTPKVTFAEEVQHDSAAPMNARKYNRPALPSLLSNNVFSNPQNLAPLGGPAPNIR
MAAPRDNAALRAPLRRSKSLPEQIGRADSTGSAVKHPAVSPQAAMALNEDDEDEEAPDMSIKAAIFMLLISTGLVA
VCADFMSDAIEPMVETSGISQAFIGLIILPIVGNAAEHVTAVTVAMKNKMDLSIGIAVGSSIQIAIFITPVIVILGWIMG
KEMTLYFNIFETVALFVTVLVVNFLVLDGRSNYLEGSLLIAAYIIIALASFFYPDGCDASAIGGNEQRCNNVQVARLAQ
GMVKRMIGM*

>g5217.t1

MTVRGKADAGGFLVVASRFLFPAPGRDAETTRRSAPSLQLPRQTAQDTCLMHKPDDPSIRSFKSKAASALGLGRKK
DTQTDVLPTHNPHTTAHPGLGGSNPPRSAPGTTNSDHSDHHEKPVAPDDNSTDVQHGSVNTAADGETPSAIGSEK
RPFYSPTRVKNGTLRFITHTKNALTHSWINVLLVFVPLGIVVKLVDLKPEIVFSMNAIAIIPLAGLLAHATEVVAARVGD
ALGALLNVSFGNAVELILFIILLASDQIEVVQASLLGSILANLLLILGMAFLLGGLKYQEQVYNNTVTQMSGVMLALAV
MSLLLPTAFHAAFEDNSIADHETLSVSRGTSVILLLVYGLYLLFQLKSHRYLYASTPQHIIDEESHPGVLAGAFDSSSSDS
SSSSSSSDSDSSSNNSDGTMKKKAKRMVKRLRRKSSASSKDDGALSAMSSPSAELNQSPFETERTSSVVTANNAGPI
TSRRHSFDVMSGDEADNDQYVPVVRDFEAASHTSRSLTKKEKRKNKKSKKDRRDRNKIDTIPEKEVEPNDPTPKVTF
AEEVQHDSAAPMNARKYNRPALPSLLSNNVFSNPQNLAPLGGPAPNIRMAAPRDNAALRAPLRRSKSLPEQIGRA
DSTGSAVKHPAVSPQAAMALNEDDEDEEAPDMSIKAAIFMLLISTGLVAVCADFMSDAIEPMVETSGISQAFIGLIIL
PIVGNAAEHVTAVTVAMKNKMDLSIGIAVGSSIQIAIFITPVIVILGWIMGKEMTLYFNIFETVALFVTVLVVNFLVLDG
RSNYLEGSLLIAAYIIIALASFFYPDGCDASAIGGNEQRCNNVQVARLAQGMVKRMIGM*

>g5461.t1

MCADQVRDAARPHDEPHMDTQVTSEAKCQGNAARPSTNSVEPQSTVSSEYGSPQTGSTENTSCKAHDASEVGT
QGLLPDLLGLQPTPTNHESFKTASTESPSQPLGAHRDACEASMQRNDFSRNDTGLAAQTKPAPLEEILAIETTGQEPS
AEQEHLVKDLNTLPSPDDEPQEQGHYNAATASVAIPQDPDRHAATRTSTEETQILAGA*

>g5462.t1

MAAKYESVKCKSCARVLAVVSETGLTPTTSPDTPHAGCSTCQQFDSLYDAIRAADADWAVFENKRDSITKSFARENH
MRAHMEFDNWLMTVEAPGSPRDGQDAQLENRDSNDGDDTNQDEEQIPGTKRRRSHSPTTQRFGNTSGLAPHE
REGIPLSLRPSPKRSRSAAASTGRKRLKFSDSVQFYDDYRSSEQFHRPSEMYVRGRNAPPEGSEYMDTSGSGLTFLKF
TGMKKVGAKWVELSEDELAGQSEKAKSWAKLKITCSC*

>g5490.t1

MDSCNTDFSLAANIARFAPLASDCTGIITITYISTDTSATVGDITRTSLVPFTVTGYSTTVTVHVTPLRNGTTLTSTESRD
VTTSVSFEKRATSDVAASFTYVLEGQTRNLLIYLRTSPYQNSSLAITPTLTKLPTFCSTVTSAITVTNIVTLPGTTGVTITLT
STMLTASTLFITDYPDPAVFEPTKTSTVNAIPTRPTEPGTTGAPIGEPSIPALPQDTSHGLPEPTASNPVQRSSRGPGLP
PDALPTSPAKPPNTKMTSIAGEIRTTTLPGIGPVVIDPSLSAIFISGVGSAITPGQQTTINDIPISFDTSATFIVVDGTRTV
GVPPSTPAPRPAPVVIGGTTIDVGQIATELDPGEVTTVNGVPISLDDAASNVVIGETKTIPLLEVTQMPAASMVVVGE
TTLDVGELASELVPGEVTVIGTVTVSRDSSAIVAGTTTISLPSSPSDVVVGGMTITAESLPSGYSFIPATAGGGLLLPNG
QTLMPGAMTTISGVEISLTPAETPVVVVEGSVSITSTRTVGSTFEGFNNGGTLGIGQPTSTSTGGLTQLTGDAVAATR
DPRNMLWVSAIVGMLFGIACSLI*

>g5491.t1

MYIQYPMRVTLLAAVLFTDFSLAANIARFAPLASDCTGIITITYISTDTSATVGDITRTSLVPFTVTGYSTTVTVHVTPLR
NGTTLTSTESRDVTTSVSFEKRATSDVAASFTTSPYQNSSLAITPTLTKLPTFCSTVTSAITVTNIVTLPGTTGVTITLTST
MLTASTLFITDYPDPAVFEPTKTSTVNAIPTRPTEPGTTGAPIGEPSIPALPQDTSHGLPEPTASNPVQRSSRGPGLPPD
ALPTSPAKPPNTKMTSIAGEIRTTTLPGIGPVVIDPSLSAIFISGVGSAITPGQQTTINDIPISFDTSATFIVVDGTRTVGV
PPSTPAPRPAPVVIGGTTIDVGQIATELDPGEVTTVNGVPISLDDAASNVVIGETKTIPLLEVTQMPAASMVVVGETT
LDVGELASELVPGEVTVIGTVTVSRDSSAIVAGTTTISLPSSPSDVVVGGMTITAESLPSGYSFIPATAGGGLLLPNGQT
LMPGAMTTISGVEISLTPAETPVVVVEGSVSITSTRTVGSTFEGFNNGGTLGIGQPTSTSTGGLTQLTGDAVAATRDP
RNMLWVSAIVGMLFGIACSLI*

>g5492.t1

MAKALSLAASGGWLGNDGSWSTFLVEVGTPAQTFAVLPAIQAQNVWLPVDDECTRLQGASASCGRSRGASPFQQ
RYSPGFQANMSSTWKAIGLYELGLGRNYGINGNGLSGFDSVGVDGFTMEKLAITAYASPGFWVGQMGLLPLPLNF
SETINSPSLISALKDEGHIPSLSYGYQAGAQYRGRKVPASLVLGGYDFSRSSEPLTIDINIVDMAKALTVGLQDIIVANTL
NGTLSMVDNERILAPIDSSIPEIWLPKSVCDRFESAFGLEYHDGTGRYVLTDQARDRLRELKPTLTFTIGADAVTGGN
TTLIQIPYEAFDLQIGYPIFANATNYFPIRRADNESQYAIGRAFLQEAYIGVNFESGIFNVSAAKWDNLEPNIITISNKDS
DVGAARGDLAGGTIAGIVVGCVTAILLCIACTWFFVLKPRRRRRQTTALDEPDEKIARDALHSAPELIGTSVHELPAKH
GHHELGEVKMVPELGNEELVHEVPSEGGR*

>g5493.t1

MSSTWKAIGLYELGLGRNYGINGNGLSGFDSVGVDGFTMEKLAITAYASPGFWVGQMGLLPLPLNFSETINSPSLIS
ALKDEGHIPSLSYGYQAGAQYRGRKVPASLVLGGYDFSRSSEPLTIDINIVDMAKALTVGLQDIIVANTLNGTLSMVD
NERILAPIDSSIPEIWLPKSVCDRFESAFGLEYHDGTGRYVLTDQARDRLRELKPTLTFTIGADAVTGGNTTLIQIPYEAF
DLQIGYPIFANATNYFPIRRADNESQYAIGRAFLQEAYIGVNFESGIFNVSAAKWDNLEPNIITISNKDSDVGAARGDL
AGGTIAGIVVGCVTAILLCIACTWFFVLKPRRRRRQTTALDEPDEKIARDALHSAPELIGTSVHELPAKHGHHELGEVK
MVPELGNEELVHEVPSEGGR*

>g5510.t1

MLFAFALLPLISLLSLGHAYQALEDETLKHLPGPGDDFDIKKGSILAPILIPRVSGTDGNALVRQHFLNFFKSQLPEWRI
EMHNSTSTTPVSKGKEVPFVNVIATRDPPGSFEGDVSRLALVAHYDSKYTPKGFIGAIDSAAPCAMILHIARSIDAALT
KKWAAADPDDFDVEHKGVQILLLDGEEAFKTWTDTDSLYGARALAEDWESTFHAASSIYRTPLDSIDLFLLLDLLGSK
GPNVPSYFKTTHWAYKHMATAEARLQKLGLMKSSPNHESKMAKRKDKKPRAERPFLPDANKQNDAFMGGFVQD
DHVPFMARGVEILHMIPTPFPRVWHTIEDDGEHLDMDTVEDWTKLVMAFTAEWMELEGFFDFKAEKREVGAEK
SELEIP*

>g5511.t1

MGGFVQDDHVPFMARGVEILHMIPTPFPRVWHTIEDDGEHLDMDTVEDWTKLVMAFTAEWMELEGFFDFKAE
KREVGAEKSELARL*

>g6794.t1

MDDLLPPYESVMRRDPWVLVAPYLPSQDLCSAALVCRQWHQTFTPNLWGSPASHFGVENDTVYVALTRFKRTLPY
ARASVRELTHTLRFPPAHAELYDGPHAEWLRDCLEHLPRLQCLIVDGLPFFDHASLLSLRHSSLRWRSARPTAYPVFG
LRLLDASSCMNATSMGLAEALPHFPDLVSLDLSKTAAARDKAVLSTLKYLRNLRVLNLRSTGIKDEEFSIVAHAIGSRV
RSLDISNNFLTDTSVRLLLELCLKETTIAAHTSRAPLPPVQNAVPTDEPGAFESENLVGHLRKKLTEGFVGSLAIEETSDV
GVSHLFLSSNAVTVEGISALLRSGRLQVLDVGILPAVVRNPTAVSPGSPDNDLELPSVSKLAPVLNEYASGNLRYLRINY
ELVTEDAPLEVPRSPRAELDGDLGIYRPTGAHELEAVQLPIAELDSQSAAVLEAPGDASYPAELPGSPTASLSSALIKSTL
GSADNGKQSATIVPEISVTVQPLQINRGPAYAPEPMPVDPPLTPLSPLRIDENYQPGSAKGVPNTRGTIAPPTTTCDS
NMPSQLTGNTAIRSRASSFYYIEDRKARLDFRQASQPRLHPGMLPKLHTLVLTGVPTMTIEKKIIHRIIQYIQDAAEEAS
IARKRARHTYVLPPGRRRIVAEEEYACNQFALRRVVLEMAPPQPTPKKITSSWRAYPTKSSTEDRDSEAFWDAAAHD
FSFFDDEECGQPGREPDRTLPLAAMDGLELAPSHPAVPTEPSKNIIETGPLLDVTGEIGKFRRQRKAAYNNLVAMGE
AEPDVEGYWPGDITVLRKPVNSEAGELDCYGNRYESGWYYR*

>g6795.t1

MGLAEALPHFPDLVSLDLSKTAAARDKAVLSTLKYLRNLRVLNLRSTGIKDEEFSIVAHAIGSRVRSLDISNNFLTDTSV
RLLLELCLKETTIAAHTSRAPLPPVQNAVPTDEPGAFESENLVGHLRKKLTEGFVGSLAIEETSDVGVSHLFLSSNAVTV
EGISALLRSGRLQVLDVGILPAVVRNPTAVSPGSPDNDLELPSVSKLAPVLNEYASGNLRYLRINYELVTEDAPLEVPRS
PRAELDGDLGIYRPTGAHELEAVQLPIAELDSQSAAVLEAPGDASYPAELPGSPTASLSSALIKSTLGSADNGKQSATIV
PEISVTVQPLQINRGPAYAPEPMPVDPPLTPLSPLRIDENYQPGSAKGVPNTRGTIAPPTTTCDSNMPSQLTGNTAIR
SRASSFYYIEDRKARLDFRQASQPRLHPGMLPKLHTLVLTGVPTMTIEKKIIHRIIQYIQDAAEEASIARKRARHTYVLP
PGRRRIVAEEEYACNQFALRRVVLEMAPPQPTPKKITSSWRAYPTKSSTEDRDSEAFWDAAAHDFSFFDDEECGQP
GREPDRTLPLAAMDGLELAPSHPAVPTEPSKNIIETGPLLDVTGEIGKFRRQRKAAYNNLVAMGEAEPDVEGYWPG
DITVLRKPVNSEAGELDCYGNRYESGWYYR*

>g6820.t1

MVVRITHTVPATAGLTGPSRLCKWRREGRRAADEMTGRLSGLSPAGDPTNLVILATQTLPAIRAPLPHYPRRFQPSLT
PPYHNDPKPDGPWSSQGGQGGSGDNHSMSVLSAGEPRSERNTPQMSPIQSYPPMQPPYRSPREAPVNPPVSLP
PLRHLSKTPPTPSDRRFGNPFGVHSILNPQAELVEQQRALRRSNSQMDSPSPVDTQNSQSLPSISRPNSVDSTQSTQ
EGQQHARPFQPPERPPIRSLYPEKGIRTHSLSRLNPPTGTIDAQQSPFLTASTRPSEIMTTQPALPTPPAGGRAHYFPA
TAPTPPPNMIRTEIRRPSVNFPQSGSASPIAQYSPYSQPASVASSQYDNNSTQGQYGPMRRHTPTHDSRQGSVPM
ESERNSMIAMAPSNQSSVQLMTIQSQDGSLHNIQVETQTASKGADEKRRRNAGASARFRARRKEKEREASISISRLE
QNVRDSNEDAEYYRSERDYWRSIAMQAQPERHITRPPSPRLRRVSVAPSRAPSSTTGHGSEASYDGYEEEMREEER
NVRRRTSSYHPAIGSHHTDVSAPSHESRGYPASTFGSVNHAPAQGHQYPQHQQAQSAGPQPTAKRPAYRDSFPPE
ANRYLRPRGRFGCHRCHHYHRTRIEELRFANVQYFVIRTTPWAALLFYSSARGHQRE*

>g6821.t1

MSPIQSYPPMQPPYRSPREAPVNPPVSLPPLRHLSKTPPTPSDRRFGNPFGVHSILNPQAELVEQQRALRRSNSQM
DSPSPVDTQNSQSLPSISRPNSVDSTQSTQEGQQHARPFQPPERPPIRSLYPEKGIRTHSLSRLNPPTGTIDAQQSPFL
TASTRPSEIMTTQPALPTPPAGGRAHYFPATAPTPPPNMIRTEIRRPSVNFPQSGSASPIAQYSPYSQPASVASSQYD
NNSTQGQYGPMRRHTPTHDSRQGSVPMESERNSMIAMAPSNQSSVQLMTIQSQDGSLHNIQVETQTASKGADE
KRRRNAGASARFRARRKEKEREASISISRLEQNVRDSNEDAEYYRSERDYWRSIAMQAQPERHITRPPSPRLRRVSV
APSRAPSSTTGHGSEASYDGYEEEMREEERNVRRRTSSYHPAIGSHHTDVSAPSHESRGYPASTFGSVNHAPAQGH
QYPQHQQAQSAGPQPTAKRPAYRDSFPPEANRYEHRT*

>g6940.t1

MAVERQIATLPPAQPTAILRGHAAHIHSTLFVRQNSRLLTGDADGWVVLWNVQTKRPLAVWPAHDGPILGFAQW
GEANIITHGRDNRIRIWRLESDDEAGKLSSTLPEDGNRQVTPKPWLLHSLPVNTLNFCSFSMCYQYPTSIAEDDSILV
ALPSTDDKAIDVYSFPCELLKVRVPRIQTVETGMAMAVKLVRSQAMQRILLLAGYEGGVTAAFMLAEEHSNTGIETA
QLVYLSQPHSQPILSLDASPDGDLYYTSSADAVVAKHRIPELPHNPGGVDSSSDVSTSKPTPSLFAKKNTNQDKPGTS

APSGLSSLLASSAPLPKFRPAPPVQPLVTPQPPCRTVDTKHAGQQSLRVRSDGRLFVTGGWDSRIRIYSTKTLKEVAV
LKWHKEGVYSTAFAEILTPEEAGHSSKAVNKRQMERDEQTRLKHWVAAGAKDGKVSLWEIF*

>g6941.t1

MQRILLLAGYEGGVTAAFMLAEEHSNTGIETAQLVYLSQPHSQPILSLDASPDGDLYYTSSADAVVAKHRIPELPHNP
GGVDSSSDVSTSKPTPSLFAKKNTNQDKPGTSAPSGLSSLLASSAPLPKFRPAPPVQPLVTPQPPCRTVDTKHAGQQ
SLRVRSDGRLFVTGGWDSRIRIYSTKTLKEVAVLKWHKEGVYSTAFAEILTPEEAGHSSKAVNKRQMERDEQTRLKH
WVAAGAKDGKVSLWEIF*

>g6975.t1

MNATHTDVLLDRYQRLMAPHMPFVVIPSGINASSLASSKPFLSKAIETIAFFHDTVTQKHMVKDLMQQVSERMLIK
GEKSLDLLQGKHTHTLHMTHILTLYIDILGMLVFGNWYNPHLFAPPSHTVLLHLTMALTHDLDIDRAPGFCEKVALM
AASQAHGVPQPAKVVTNDERRAVLGTFYLTSQILTSFRKIDTLHWSPWLTKCAEALTQAKEYKSDILLVQLAQSQRIM
QEAMGTECDHAPVSFYAKSFLSDLDNIELASANGMSAIVLRLQQACTRTAVWERSFGNLASHTVKETDLRQRLDG
MWHCMDAAKRYTDIYLELPVEEYPLVPFGVFAQFAYIFVVLVRASLIEMDGWDVKALRSFIDFSSLLEKASQRYDAVS
TSHPDGLILNNEAFAKWSAKTKWAKSFYDTKFLPVDPNTSTVNQPPPGCTTIEADETQESRRAFPTPLQVPAEFSTD
TFAIEDSMWHSLGNPAIFLGDIDSAFTDIP*

>g6976.t1

MDSSNEELEDATVTSQNSTHTHHGSCTVCAKAKSKCIRRPGNGDCERCIRLHKTCQIREPVKRKRKARKTTRSAQLQ
NLETKLADIAQALAIKQAQEVSAGSNSRSPISQQMDALYQESDDTYPTPLVITPSITRSNDFLASAHDPTLETVNPEHV
STGMNATHTDVLLDRYQRLMAPHMPFVVIPSGINASSLASSKPFLSKAIETIAFFHDTVTQKHMVKDLMQQVSER
MLIKGEKSLDLLQGMLVFGNWYNPHLFAPPSHTVLLHLTMALTHDLDIDRAPGFCEKVALMAASQAHGVPQPAKV
VTNDERRAVLGTFYLTSQILTSFRKIDTLHWSPWLTKCAEALTQAKEYKSDILLVQLAQSQRIMQEAMGTECDHAPVS
FYAKSFLSDLDNIELASANGMSAIVLRLQQACTRTAVWERSFGNLASHTVKETDLRQRLDGMWHCMDAAKRYTDI
YLELPVEEYPLVPFGVFAQFAYIFVVLVRASLIEMDGWDVKALRSFIDFSSLLEKASQRYDAVSTSHPDGLILNNEAFAK
WSAKTKWAKSFYDTKFLPVDPNTSTVNQPPPGCTTIEADETQESRRAFPTPLQVPAEFSTDTFAIEDSMWHSLGNP
AIFLGDIDSAFTDIP*

>g7027.t1

MRGAVADITSLPSIGTLLTALANHFSGKKLSILVFNAAFNTRPRIGSASEADISQSLTGNLHWPIVLMENLVRQDLFTP
HSRVVVISSDRVRDPSPGSGLFNATRAAMESLVRSWAIELPHSFPGTTVNAVSVGLTDTPGLRSFPPEAVQALKDQR
LPKVKVAEGGRMGQAEDVADVVGFLVSEQSRWVSGSVMAANGGAEWVGGSS*

>g7028.t1

MANYVPPAPQLREQDLKGKVAIVTGASKGIGRAISLSLATRGCSILGTYSSPQSAHNFDTLSSTVRDLYASSNNEVPT
MRGAVADITSLPSIGTLLTALANHFSGKKLSILVFNAAFNTRPRIGSASEADISQSLTGNLHWPIVLMENLVRQDLFTP
HSRVVVISSDRVRDPSPGSGLFNATRAAMESLVRSWAIELPHSFPGTTVNAVSVGLTDTPGLRSFPPEAVQALKDQR
LPKVKVAEGGRMGQAEDVADVVGFLVSEQSRWVSGSVMAANGGAEWVGGSS*

>g7402.t1

MQQSTSSRPPLSDHAKPRLNLVHRYPTLHNASSPAGCRNLSPHQYEPCHGGPLGTPFLMMSHIAGRKLSEFAPYLT
ASERSSIDQTLGIYVRALTTLSATQFGMTHRVFAKKGSNSWREAFLALLEATIRDAEDMLVNAHSESIRFWVGKHAQ
CLEDVIEPRLVALNVCDPENVLIDERTKQVAGLVGFSNVIWGDPLMSGGMADGNEAFFAGLGDYPVKTASARTRM
LMYKTYRATVRIVAHHYRPHDGIDELGARRELSYALNDLAAI*

>g7403.t1

MLKCPPPYNVQLLRHEKHFLETERKTLETLHEYTQIPVPQLIKYDSHGGPLGTPFLMMSHIAGRKLSEFAPYLTASERS
SIDQTLGIYVRALTTLSATQFGMTHRVFAKKGSNSWREAFLALLEATIRDAEDMLVNAHSESIRFWVGKHAQCLEDVI
EPRLVALNVCDPENVLIDERTKQVAGLVGFSNVIWGDPLMSGGMADGNEAFFAGLGDYPVKTASARTRMLM*

>g7467.t1

MPLYAMRMAFSVNIMRSAKDTKRSGHARNESGLAVALPPERLCGGHLGQFDRSQPNNFNMAFEWLAPILTPAAL
RSVQADGRPVITLFLVIFGLSVFGVLIWYVHFVTSKMYPKKKDPNAKKGPLLGFIKR*

>g7468.t1

MAFEWLAPILTPAALRSVQADGRPVITLFLVIFGLSVFGVLIWILPMEDDMTIKRIIATFMLYRQAQSRSSLRRNHSDR
HWSPFDSPRRVVCLAGGLKDGSDLLILVANGVDRPPRSTRVSYL*

>g7688.t1

MGDKVTSCIDEVEESDLSAEGKDKAKALITKELKKLLKTCAKATDRVYVALDLLRSSLHIEESSQNLDISIVKDLIAESTP
RKSWLRSVTEVVLKTVMGAAASIAAVQFYSTSGFDLALQTYPFGTHSQVISLVQRAQAVTNLTGEIYSIKLEDLDQRY
KELEAASESHSLRIDNLVEALGVPNAEGTYYSSILKSDCEALSNVQKISADADRQIQQLRSELATMRKNIHRMDIRLM
KRLDKVDRHGL*

>g7689.t1

MSIGSEASMTEITDKDDTSDGLWQHSNTQTANDDEQTKKDTSGGDGAIAATPTQPRAPNVTDATNADGEPKKAE
FVNQFQENLRLEVEAEKEVRGATEEETKAAGIENDTPEKSPENSDMVNSAIVTHKRRVDYADGRAKKSTSAVKIQR
NRADEVLGSMSDIEKAMFQLYEYIKAQNDSKKPLYMWATFGGYSSVNVALLDLHDKAIEFEEAHRLPVANARILLSR
AT*

>g7814.t1

MDPRKKQELLTGLQVFLGEGTEDYYWQNGTLYRRGYMFYGPAGTCKTNLSTAIASHYNFPLCVIDLAGMDDTVLQD
KVKNLPPRCVSFSENIDAAGIVRERTMVPKTDVGDEGDETSSESESTDTWDTEPSKVKVKKADPKKMNTSLSQPSR
TEVTLSGLFNTLDGPGSREGHIVIVTTNAPDSLNEALHRPGLIDTQVYLGFADRIIDGITFTRIFGSDKQVKESQKAVGK
LGREFGRLLPSDFFTPAEIQKFCMNRRGRPQKAVDEFPGYIDEKRTGKTKFQYDIHRRAPKPTFHKEGVIHDDDSSDE
VEYSPRNTRDSSEGAHNAFPAFSRISSTETSVKRPAKEQLFVAHGHEHDDDALAYGDYYPRRQGVWEDRLAGDVV
NTCQGLRDMLTFSLVEPFRHQNISHKEAGYRTW*

>g7815.t1

MSDYSSHDDDRRVDSWRQTKGRIAKRQSNFLETVLDTSIASSMEKPLFKVIRMWIDSNTGLDLKFLVWAIALCVPT
HKYGSATCKWLSSNLTNSVELDHQDALVTDLLIWINK*

>g7832.t1

METGYIRRVHEIAEYLVPSINFGNDFGSSRLPLIQAVGQGDFHWVRQILLKTDIDPDIRDCQGWTALQNACSIKTDSP
ESRNQEAIIRLLISQGVDVNAAGDESFGRTALQAACERGNERIVDLLLENGADVNEDVATANGHSSLVCAADAGDF
GIVKKLLDLGADINQPRSKESGHTALSAAAYRGNLNMFDFLIRNGANAQGSAGLFALKTAVSRNQLAIARRLLELGLD
VNSCANGPAPLHQVTHVDMLNLLVTYGARFDLPGSEGSGQTALQRAAQWHAFDLVIELVRLGSDVHLPGPATGGS
SALQAAATRCRYGTEESVRIMRFLVEEHGVDVNEPRSQEEEYTSLETACQETAQQDEHRWSIEGVRFLVEQGAVITP
FTLHVATAWNHTKLLDFLLQNGVRIEHISSPANIRLYPYFLSERRKFGATVIETARINGHLALAEALENWSPSTQLGIGL
NE*

>g7833.t1

MAVGFGFSVGDLIAVGKLAHDIANALNDCRGASAEYRSLIELLESLKTSLNLIGNFISTLPITTSSRVDQAFMNGILFHA
GCCYKLLNEFAADSRRYTQSLLNGQGSKAKVAFRKIKWSLYSAEDSRRLERRLSTHMEAFDRYLLAINIQTETSFAEES
RGQANRISVTVAAIYDTVQTTQNMMPKMLGYPWEGDVMSSHVHVEDALGRPLVLPSTLCRTRETLKDTMRIMFS
DHPGLREVVDGKFELVDKQSQLTVFDGRDVIPRMRWLHYQPSDAIQPGAKLLMNVLRVRTFRAQESSLINSVQTLE
TCPRCGFATPGRQFRKCGNCDMGFRRSVRELFERVPRSAAHNLRWPNNPLPRTIPNATPPVDCGPKKLPNADMET
GYIRRVHEIAEYLVPSINFGNDFGSSRLPLIQAVGQGDFHWVRQILLKTDIDPDIRDCQGWTALQNACSIKTDSPESR
NQEAIIRLLISQGVDVNAAGDESFGRTALQAACERGNERIVDLLLENGADVNEDVATANGHSSLVCAADAGDFGIVK
KLLDLGADINQPRSKESGHTALSAAAYRGNLNMFDFLIRNGANAQGSAGLFALKTAVSRNQLAIARRLLELGLDVNS

CANGPAPLHQVTHVDMLNLLVTYGARFDLPGSEGSGQTALQRAAQWHAFDLVIELVRLGSDVHLPGPATGGSSAL
QAAATRCRYGTEESVRIMRFLVEEHGVDVNEPRSQEEEYTSLETACQETAQQDEHRWSIEGVRFLVEQGAVITPFTL
HVATAWNHTKLLDFLLQNGVRIEHISSPANIRLYPYFLSERRKFGATVIETARINGHLALAEALENWSPSTQLGIGLNE*

>g8520.t1

DYWGDGGGGGEGRGTVRSDGGRRTWEKEERVLENLFFVRFLGCDVPHYHFWTRLGEWGDPCFPNPAWEAAGQ
MDEDEQDAVEIGGDAKGGKRRYRLGSKNRVESLSPAKTRLIRVRRPSLRLSSSTASSPSSPSSPVSIASAITRAEYHFHT
HGLTSTLSAYRARWLAKGKTAHWKTFAANLPTLWPSGNLPDVPPVESRPEACLALKIAAIDVSSAGGKTMFSPLMG
GEAWTRNDDKLWEVVEVVEGVENNKDEGNETGDETDAFGGDDVDAEEADGVLEEIGSGNDDEALYRRGSMVFA
HDDIDTDPIMAWLDKVSPTFALSTTLMNASYISDMLEVGIGLDEWEERYLQDCKNENGEA

>g8521.t1

DYWGDGGGGGEGRGTVRSDGGRRTWEKEERVLENLFFVRFLGCDVPHYHFWTRLGEWGDPCFPNPAWEAAGQ
MDEDEQDAVEIGGDAKGGKRRYRLGSKNRVESLSPAKTRLIRVRRPSLRLSSSTASSPSSPSSPVSIASAITRAEYHFHT
HGLTSTLSAYRARWLAKGKTAHWKTFAANLPTLWPSGNLPDVPPVESRPEACLALKIAAIDVSSAGGKTMFSPLMG
GEAWTRNDDKLWEVVEVVEGVENNKDEGNETGDETDAFGGDDVDAEEADGVLEEIGSGNDDEALYRRGSMVFA
HDDIDTDPIMAWLDKVSPTFALSTTLMNASYISDMLEVGIGLDEWEERYLQDCKNENGEA

>g8571.t1

MLGKQLDPLIDELHELEYALQDLFADFHEFDGFSNQEIASEVQNVFGSTALLDEVLEEPEEWSCCQDDTGLKIECAEI
QSNWLHRTIDWFQRGLSSLGDSGRVEREEDTADVYTAVKNTSEVVHDRAEVEEFADSEGVMDSDYGDEDAMDFD
VLDSP*

>g8572.t1

MPMPTASRPNGSCTKIPEHTPLPVETPPTDTKRGTCRLHFLESEEQDSLDYPRYKIAIQFELFTAVRSCPLCDAELIHYG
MIARVEHLLAQACKAEGNVTALRKRYDHAYEPFERLEELEEDVAGVMEDFRNETNNLSQPTNDYLFNQLHKVLNK
NKI*

>g8657.t1

MSGVRAHIASRLRSSSLSAVPPPQHSSNYSPDLAKIAASVLYRSPLPSHEGRPVFILNAAALPDSHEADYDSLLPYVLA
RLPEEDELLKGFEYEVVFFAGDGDGSVTSKKHRPGWGWFLQAYHVLSRAMRKRLQRLYIVHEKAWVRILTEIFSTIVS
PKFRRKIYHLSNLTQLAHEIQIQNLLIPPSTYLADRRVSEHISALNGSGKRAFGARTPFPTATNGKTRFPRVLRETTSFVL
MEQNITSEGLFRVPPHSRLRDALKEAYDRGQKYIIWKDNDATLPLPPYPHAEHQDEILAEVVPTDAYSVFMAAAMIK
AWYASLRQPIFPTDSYRDLKRLYGDSQEILELEKLTDLFSPTSEWSLLPGISREILCRHLLPLLSAIAAREEENKMTAENL
AVCFAPGLLCGPDQLEDAKMSSIIRRIFTQAIDMWTEGLREACGQTEEAFYQELKLPGDENDWEDPADAKRDSADS
KGSLEDQMSGITLLDNEKLPTYHEPQQTQSEELPPPLPPRSRAPSAKSSADSVQRKPAPPLSVPPRYSTVISDAPENV
AESPVTYAATTDGFAPPRNDDNRIGQPPKVPPRWNGQSDEKKSD*

>g8658.t1

MNRSPSINSLARPVYPVTPQVPIISVPTKASTLPVPAAPPRLRTPSPSLMQRMPSFENFAKDQNKGGVEAEDEAARG
RTLKPKKMNLKKQSVEDLRRLYEERAGTASVLVQAGKQKGV*

>g8729.t1

MIFLGRSRARRKEYMHELEAKLRSYEQIGIEASSEIQTAARRVLHENQKLRSLLHGRGVSESEILAALEDMSDRRYEH
DTAASRLHAMLERRVNTNVVSSTSSPIPSYTRAASIPRQKPPVPPVTIPPTRPTALSYNDSPSPGSMVSTMSTPPPAS
YPATLYTTPMTPSAAQIKSEHVQYDYPYDQPYSNAWAYSSDGNYTAQPVSYYNSSSCVDAANIIRTMRAGVGSGLE
ADLGCHASDRHCYVNNATVFSAMDRYSQHNATI*

>g8730.t1

MSDSRVSKAQNLARIRDNQRRSRARRKEYMHELEAKLRSYEQIGIEASSEIQTAARRVLHENQKLRSLLHGRGVSES
EILAALEDMSDRRYEHDTAASRLHAMLERRVNTNVVSSTSSPIPSYTRAASIPRQKPPVPPVTIPPTRPTALSYNDSPS
PGSMVSTMSTPPPASYPATLYTTPMTPSAAQIKSEHVQYDYPYDQPYSNAWAYSSDGNYTAQPVSYYNSSSCVDAA
NIIRTMRAGVGSGLEADLGCHASDRHCYVNNATVFSAMDRYSQHNATI*

>g8810.t1

MRSSYVLSAAAAFAVVGAQDIDFAGVDATPDPVINIIPGLKEQVVLFKEAEAIADVISQVTSDPLDVKPVSEPSAAPD
VKRHLKRAACDPEPSNPNTYGIDLSSASKFRADTKLASLARGASTPSGYFNTFTNLQGASSAYGYMGYKVVESYDPS
LCASECNSKSGCLSFNIYVERDPSANPGPDCQDPEAVANIKCSFWGGPVYTDTATNTGQWREKFEVAIAGNNGYSS
LTTQSAEGYK*

>g8811.t1

MGYKLFVQNAFDAKLCATACKETNKWALEHPPTEGKPQLCRYFNTYILLKNGVSQGQYCSMYTQEWDASFATNDG
QYRGDDHYTIQYSFGFTDESDDGVPVCPNDISYLKSSGQEFCSSYISYEQPTTTATATATTTPAPLTNTITRTSTSYTTAIT
TVTAAKFAGRHARRVDNGTEVSHDIIVDPYEISIVAVQTISPEDAGMPKLNDTMAAELNGGAIEKRDLATPASIAAW
PTSKISAAYSQVGTGTATRTATITATAPTPLATDIKDFQTVVIVQSTCTAPAPRPTAPSYTKIVGGEPSEGIPMVPGPADG
RHWQEQFSLALPFPVKIFETSSTQISIAVRGYITAGPYTFNAFADELYVYGPGWNQGLFYRVDGAIGSRKIHFSWFTG
SYYWGHERFHITATFDEAKPGILISKIFDTWGQVNPRRTISVTGNGKNIPFAQYGSTNVHEGQEITFDTTGAGSVSAK
DFDCIDCCKKTGWEWHVCSEV*

>g8835.t1

MRISHHLRSGSLLSWDQLADAPDLPVPPPMYRERTVSDQSRVSVQNRQLARHHRQTSSSGFASTKIPSKWGNVLP
QDVDARPDVASSIYSSRPQSPPDSFGGSMVTLSHTGTENQVFNISSSDLRRPRRSASFPTDNEETPKPAKRHGLSGLT
TVIGHSIESSLLAVPVPLARENSVADTKTSKFREEFSPSPPKKKTSQSVSIIKLLNPKRLSGRSQSEANLHPEASVAAMD
GPSDILAIPADRERGHSRSLTSFHTEQNAMGRNKGANHVWDRALQAHQEEKASLFLPRNKDLAVHASPFRERSGS
VSVRRASIEGLDPTSRADDGSSTRVSTLCPTLRTTGEEPSEGLYTLASRRSALLGRDDVSVGQEVATTFERQGDRTDIV
GAWGRYPSHTRHDRTASAGKNDNVQPRDFALEAAIKFASATDDEDLIDPTERRPSTPLLPGEKKKKKQKKVGSGRIA
KSNSMTFGRTLMKNYAKMFRSQSTEFQRHGRGHRSSIASGGILEHPELELLPHVWAGEVHKVHADVQTLKHEEQP
LNNGFLMRITQSKGKLRADDSMATLRPRRNSSAPNLNEVSFRDGAAGVGHASDRAHVWSAYYENCVDAYPRLSM
DTNRLLEDFSHPKRLSFDSRRASIHSRAMPIRIPKHERRESQKSNASRVKSFVYQDHNDSASEERSLVSVRRSTMDLIS
KFKEQEATEHERILSFSRMDSVRAEEGVAAP*

>g8836.t1

MATKLCCTAEQDHCSESPELPNARLSQLTPANRPLLPKSTGSPGSRFTSARSEDLHELREIFHNAQASDQDRAMPM
RALRARFSRPSMHSIRSLHKMTSMRSLIRRKFSKELPEKASGVSSAHTQPKQNSITSIPDTVVKQLKDSNQQLRITKH
DLRKDLLSDKKPDEGGYDSDAKVLDNIAKNIGKKPLSKRPSIRSVDWTTTTGSKSTPESSTKGRISAELKRDLYPYSVD
KPQAASVSNRVTQVFSTPDLRSSTSGERNRKVRRSHSATSMGLPKPSPISPLRLPSLTNHDSFVMPWSEVMHKSLRL
SQFPIPPQHINTVTSPTLLGKNDIQKHAHDHHQNQNSDASASSPRTNVSLALPAARIVEIRVQQPTSIASPRPSTSVR
GCSRENSPSRKGTRTEGEDDGDNNPRHSVHLHSMRISHHLRSGSLLSWDQLADAPDLPVPPPMYRERTVSDQSR
VSVQNRQLARHHRQTSSSGFASTKIPSKWGNVLPQDVDARPDVASSIYSSRPQSPPDSFGGSMVTLSHTGTENQVF
NISSSDLRRPRRSASFPTDNEETPKPAKRHGLSGLTTVIGHSIESSLLAVPVPLARENSVADTKTSKFREEFSPSPPKKKT
SQSVSIIKLLNPKRLSGRSQSEANLHPEASVAAMDGPSDILAIPADRERGHSRSLTSFHTEQNAMGRNKGANHVWD
RALQAHQEEKASLFLPRNKDLAVHASPFRERSGSVSVRRASIEGLDPTSRADDGSSTRVSTLCPTLRTTGEEPSEGLYT
LASRRSALLGRDDVSVGQEVATTFERQGDRTDIVGAWGRYPSHTRHDRTASAGKNDNVQPRDFALEAAIKFASATD
DEDLIDPTERRPSTPLLPGEKKKKKQKKVGSGRIAKSNSMTFGRTLMKNYAKMFRSQSTEFQRHGRGHRSSIASGGI
LEHPELELLPHVWAGEVHKVHADVQTLKHEEQPLNNGFLMRITQSKGKLRADDSMATLRPRRNSSAPNLNEVSFR
DGAAGVGHASDRAHVWSAYYENCVDAYPRLSMDTNRLLEDFSHPKRLSFDSRRASIHSRAMPIRIPKHERRESQKS
NASRVKSFVYQDHNDSASEERSLVSVRRSTMDLISKFKEQEATEHERILSFSRMDSVRAEEGVAAP*

>g8878.t1

MRGRGGGGSRGRGRGGGGMARGRGGMRGRGRGASRGGFVAARTRDEVEDSNVHMREPSVSDDSESDASDD
AASDEEEEEAQPISNAYASLLQSFSSRNTGSDEHRKKRRKLGHDEPEPDVQVDEESPALEAEDAVAEEEEEEELDNDEV
EDEDDQEEQKLEQAYERHFANPDENELAMRLKRIAANQWTSQKLDRKPGLGAGVLQVPSDEVKTPVRKLRSVRDV
DLKARLVENAQKRIGTFDELEQAITPSIFGYQDLLFGARTVQNANRLRDITCLHALNHILMTRDRVLKNNAKLAQAK
DDDTDTEYRDQGFTRPKILFLLETKQACVRALDSITAVHDFEQQENKKRFLDSFSLPEDKFSEDRPADFRDLFEGNDE
NEFRIGVKLTRKTLKYYSTFYNSDIIFASTLGLRRAIESGDPKKRDSDFLSSIEMVIMEQADAGLMQNWEHAEFVFEH
LNLQPKDSHGCDFSRVRNWYLDGHAGFVRQTIVLSAFLTPKINTLYNRHMRNFAGRLKYTADHTAGLIETLSYGIKQT
FLRFDSPSHLTDPDARFKYFSTTVLPSITRLPKPVEGGLGVLVFIPSYLDFVRVRNSLVDTDIAYASISEYTDATDVRKARS
HFMNGKHSVLLYTGRAHHFHRYNLRGVKRVVFYGVPENPIFYDEVVGFVGKSVERAEISRAEASVKVCFSRWERLEL
ERIVGSKRVGKMVGDRGDVFDFV*

>g8879.t1

MREPSVSDDSESDASDDAASDEEEEEAQPISNAYASLLQSFSSRNTGSDEHRKKRRKLGHDEPEPDVQVDEESPALE
AEDAVAEEEEEEELDNDEVEDEDDQEEQKLEQAYERHFANPDENELAMRLKRIAANQWTSQKLDRKPGLGAGVLQ
VPSDEVKTPVRKLRSVRDVDLKARLVENAQKRIGTFDELEQAITPSIFGYQDLLFGARTVQNANRLRDITCLHALNHIL
MTRDRVLKNNAKLAQAKDDDTDTEYRDQGFTRPKILFLLETKQACVRALDSITAVHDFEQQENKKRFLDSFSLPEDK
FSEDRPADFRDLFEGNDENEFRIGVKLTRKTLKYYSTFYNSDIIFASTLGLRRAIESGDPKKRDSDFLSSIEMVIMEQAD
AGLMQNWEHAEFVFEHLNLQPKDSHGCDFSRVRNWYLDGHAGFVRQTIVLSAFLTPKINTLYNRHMRNFAGRLK
YTADHTAGLIETLSYGIKQTFLRFDSPSHLTDPDARFKYFSTTVLPSITRLPKPVEGGLGVLVFIPSYLDFVRVRNSLVDTD
IAYASISEYTDATDVRKARSHFMNGKHSVLLYTGRAHHFHRYNLRGVKRVVFYGVPENPIFYDEVVGFVGKSVERAEI
SRAEASVKVCFSRWERLELERIVGSKRVGKMVGDRGDVFDFV*

>g8897.t1

MVLRLSDLGSMEPRVLPPEQKDDRDDDLRCVDDVFHINCERPARCQDPQLEVKDTKEITQRNQVDSPLLRLPAELR
NMIYWNVFAGAVIAINKPHTKKDNILTTYPSVRALMYACSQLYREAEIYMFGLTLFVPSYFALRFLWLSNMCAGMPG
VGTASIMGCKITAGNARAIIEGKIKKEDLAYYPCLRQIFVQLWDHRVSVKRSEFYRRVFREAFGIPNLDVKFIK*

>g8898.t1

MTLISRSTQRNQVDSPLLRLPAELRNMIYWNVFAGAVIAINKPHTKKDNILTTYPSVRALMYACSQLYREAEIYMFGL
TLFVPSYFALRFLWLSNMCAGMPGVGTASIMGCKITAGNARAIIEGKIKKEDLAYYPCLRQIFVQLWDHRVSVKRSEF
YRRVFREAFGIPNLDVKFIK*

>g8954.t1

MTPSRLRSLSPMPKNHVTRSISTPQLAKTARSQSQSESESDGMNTIPDPRSATPQPPLSRHDSTDSHPDLSQEVSTL
STKLINAINHSTMLDDSLQQTRHELEAAREKLAQLEAQVREHEEKVSKGLLMDKIVYDKMEKQLSSELHEERKRRAQ
AEAAKRKTDSEVEALTAALFEEANVMVAAARKETEASEKRGEQLKQQLGDAEVLQHSLQDQLQDLKGVVEKMSSH
GDDNESHILTTNTAPSTPGITPADKMSKLFEAMNLTPSTPGTDEITPDHPLHFSHLIHPVLRSDLTAFKEFQDMLKTSA
RSSAPASRASSGNYSSLSVLGLGSLTNSSTTSLPSKSSASVTNSPRESVATAGMPNLKDEKFYKRALVEDIEPTLRLDIAP
GLSWMARRTVLNSITSGSLVVEPNPAPSSRFRVPVFPCSLCGEARNGDQYARKFRFKPSDTEDSQRYPLCDWCLGR
VRATCDYIGFLRMVAAGHWRAETEEEKKSTWEESVRLRERMFWTRVGGGVVPSFVPLRDGSPHSPTFTSDDIKQQ
REERMSEEESIISFEPRDITASGSAASTMGNESPRKSEDDPFQSKSGDKAKRISIGNTVISSDNAAPELGSTTPPPAEQ
EGGEAAKGNDDQKIESPLPIPSLTIEEEKKIEQDAEAQLQDEVKKSLEKPALQRQRSSSNPPPVAATLRKKEESSNRLSL
RIPGSMGAGMSMPGAFD*

>g8955.t1

MAEYAFIAHSAPWVHGSGTMTPSRLRSLSPMPKNHVTRSISTPQLAKTARSQSQSESESDGMNTIPDPRSATPQPP
LSRHDSTDSHPDLSQEVSTLSTKLINAINHSTMLDDSLQQTRHELEAAREKLAQLEAQVREHEEKVSKGLLMDKIVY
DKMEKQLSSELHEERKRRAQAEAAKRKTDSEVEALTAALFEEANVMVAAARKETEASEKRGEQLKQQLGDAEVLQ
HSLQDQLQDLKGVVEKMSSHGDDNESHILTTNTAPSTPGITPADKMSKLFEAMNLTPSTPGTDEITPDHPLHFSHLI

HPVLRSDLTAFKEFQDMLKTSARSSAPASRASSGNYSSLSVLGLGSLTNSSTTSLPSKSSASVTNSPRESVATAGMPNL
KDEKFYKRALVEDIEPTLRLDIAPGLSWMARRTVLNSITSGSLVVEPNPAPSSRFRVPVFPCSLCGEARNGDQYARKF
RFKPSDTEDSQRYPLCDWCLGRVRATCDYIGFLRMVAAGHWRAETEEEKKSTWEESVRLRERMFWTRVGGGVVP
SFVPLRDGSPHSPTFTSDDIKQQREERMSEEESIISFEPRDITASGSAASTMGNESPRKSEDDPFQSKSGDKAKRISIG
NTVISSDNAAPELGSTTPPPAEQEGGEAAKGNDDQKIESPLPIPSLTIEEEKKIEQDAEAQLQDEVKKSLEKPALQRQR
SSSNPPPVAATLRKKEESSNRLSLRIPGSMGAGMSMPGAFD*

>g9143.t1

MPKALHFKGDKKVKKRRKAADPYDADEKPSKQLTTSAPAEAESDDSWVSADAPTDISGPIVIVLPTDSPNTCLACDA
NGKVFASELENIVEGDVATAEPHDVRQVFVANRIAGTEQLSLKGHHGRYLSCDKLGVLSATATAISPEETFVVVSVPD
NPSAFSLQTARDKFLTIDGSSGKTPEPRGDAEHIDFSTTFRIRMQARFKPRLKASKEDKANVKISRKELQDQIGRRLN
DDEVKKLRKSRVQGNFHETALDMKVKSSHDKYASM*

>g9144.t1

LGVLSATATAISPEETFVVVSVPDNPSAFSLQTARDKFLTIDGSSGKTPEPRGDAEHIDFSTTFRIRMQARFKPRLKASK
EDKANVKISRKELQDQIGRRLNDDEVKKLRKSRVQGNFHETALDMKVKSSHDKYASM

>g9886.t1

MCGYNRVNDSYACANPEILNHILKDELAFPGYIVSDWEATHSMVGTVNAGLDMERPGITSPTGIFYFGDSLADAIEA
GNISSARLDDMATRVMTPYFRLGQDEDFPVTDPASGPVFLTYTYGHQSPLAASFPETPARDVRGDHAKGIRELGAA
GTVLLKNLNKTLPFSNETSFDVFGNDLPDPTIGSVFLDYGNAAMGYPMGTLDIGGGSGTVRHTNLVSPLEAIRNKIR
SLGGRVQWLFDNDEIADGRFRSIYPVPQVCLIFLKAFATEGSDRSDLDFHWNATLAVESTAKLCPNTVVVTHGPGVV
LMPWADNENVTAILAAHYPGEETGNSITDVLWGDVEPSGRLPYSIPKAAADYGPPIVELPRNVTDPDAWQADYVE
GQLIDYRRFDANDDLEPHYEFGFGLSYTSFVMSHELNVELRGVPLTPVPDKSKGIAPGGLRDLWTVVAVATVNITNN
GGRAGSAVPQLYLSLPQETTPAGTPVKVLRGFEKVHLLPGETQAVTFNLMRRDVSFWDVDSKTWTIPAGSIRFTAG
FSAKDLHAVRDVEVLV*

>g9887.t1

MRVAKIVTLLSTSLPIRSVWAEDLPVRVPLSWTEATVKAIHFVSQLNTTEKIGLVTGSYGSSPALPCVGTLVAIERLNYT
GLCMSDGPAGLSRSDGVSVFASGITVAATWDRRLMYERGLAIGQEFRAKGAHVHLGPAAGPMGRHPQSGRNWE
AFGPDPYLAGVAMNESIFGIQGAGVQTSSKHYIGNEQETQRTRSTREDGTVIEAVSANIDDRTLHELYI*

>g9970.t1

MGSLRTLFLSVYASNGIIRTLFLPLCYLTLFSPFILVLAWYFGLGQTLEKAIPEDFSWFDLLPTFAISAVFVLLPTRLLSSSG
KVVKSQDGKRRVQSLPYWIPGVRHLWSIIFGGEEWLNGVRDSATASILAYQAAGTKHNILLSDSLLGQIYKNLHSLEL
PDSSQLAILRNTFNLPKVMESHYLEIQNEINQVVETELYKGVAEEGLIKASLRILTESLPDFVTFNSSIVDQMQWERVS
NVELTDGTSEAECDLFTLLNEFCCNAILPPIVGAQFTESYQLLATDLASCNERFWALALGLPRLTPVQGLPGAALARKR
LLKNFANLFTELTHPPVRRVPDDDESVSGEEDTDADVVTPLMKLNELFTKHELPAELRASIGLHLVHGIYSDVIPLVFW
TVLHIYSSSKAQDVKAPEGSPLENIKAESKAWAQAVQPPSIHPSFPAPPEIRFGSVKEALTSSSLPYLRSCINEARRLHT
CSASTYRVKKSLHLQEDGPGGKEQWELETGTYIDMGLSQSLINSSSAFHASPETFRPNRFLNTPPSSSVLSPAYSSQSY
KTDLIISIVSGVTQLWEIAPAPKKSFFDHIQEAGNEASFGAEALTAEQKAAKEVANREKEDAKRKEARWVFPKAVDGA
VTRVPKGDIRVRIRRREGLPTSKVVRRIG*

>g9971.t1

MESHYLEIQNEINQVVETELYKGVAEEGLIKASLRILTESLPDFVTFNSSIVDQMQWERVSNVELTDGTSEAECDLFTL
LNEFCCNAILPPIVGAQFTESYQLLATDLASCNERFWALALGLPRLTPVQGLPGAALARKRLLKNFANLFTELTHPPVR
RVPDDDESVSGEEDTDADVVTPLMKLNELFTKHELPAELRASIGLHLVHGIYSDVIPLVFWTVLHIYSSSKAQDVKAP
EGSPLENIKAESKAWAQAVQPPSIHPSFPAPPEIRFGSVKEALTSSSLPYLRSCINEARRLHTCSASTYRVKKSLHLQED
GPGGKEQWELETGTYIDMGLSQSLINSSSAFHASPETFRPNRFLNTPPSSSVLSPAYSSQSYKTDLIISIVSGVTQLWEI

APAPKKSFFDHIQEAGNEASFGAEALTAEQKAAKEVANREKEDAKRKEARWVFPKAVDGAVTRVPKGDIRVRIRRR
EGLPTSKVVRRIG*

>g10091.t1

MRGGGHMPISDAANINNTGVLISSTNLNTLELSQDGETMSIGPGPRWGDVFNYLEFTNKTVIGGRLAPVGVPGLLL
GGGISWYSAKHGLASSEGKIKAYEAVLADGTIATITANSTYSDLYWALGGGANSFALITRFDLQTFPSITPLIADAHYGS
SNVTRDAYFAAILNMALTNEQDLASTIIPVCRWGPGSTAPSYESTLFHNGTSVPTSGPLAEFIHGNTTGLTALNGTAT
MRPISLAQYGRAMRSAFREGGQSHGLRQKFRVVSMKATAENLAIAHDTFFNSLAASGLANRVPDFFAGLDFNIVN
REMVKRSAGLPQNIPLEPAFWVEEAISWGSGGEFDNEVAEWVKNVNVEIERKLQEVNGLNMYVYLNDADKGQK
VFEGYGGESVGRLKTVRAKYDPERVFTDLMPGGWKVEHVEL*

>g10092.t1

LDPSTACQILKNTYPNITYLPEDAGHADENQVSWDSSAWLGPACVFAPTCATSLSFAVKTFVATHTKFAMRGGGHM
PISDAANINNTGVLISSTNLNTLELSQDGETMSIGPGPRWGDVFNYLEFTNKTVIGGRLAPVGVPGLLLGGGISWYS
AKHGLASSEGKIKAYEAVLADGTIATITANSTYSDLYWALGGGANSFALITRFDLQTFPSITPLIADAHYGSSNVTRDAY
FAAILNMALTNEQDLASTIIPVCRWGPGSTAPSYESTLFHNGTSVPTSGPLAEFIHGNTTGLTALNGTATMRPISLAQY
GRAMRSAFREGGQSHGLRQKFRVVSMKATAENLAIAHDTFFNSLAASGLANRVPDFFAGLDFNIVNREMVKRSAG
LPQNIPLEPAFWVEEAISWGSGGEFDNEVAEWVKNVNVEIERKLQEVNGLNMYVYLNDADKGQKVFEGYGGESV
GRLKTVRAKYDPERVFTDLMPGGWKVEHVEL

>g3659.t2

MLQTEVEYQTPVMMATAVSTPRKLWEHPDPKSTAMWKFMQDANARRGLHMQTFRDLYNWSVGPSRTDFWVD
MWKASNLIYSGTYTTVVDTSLPMEINPHWFQGTYLNFAENVLFTASPTDSTVRTTTHKEESKIALTEIREGNTEVRHL
TSSSTDMGTKGILERLLQIRPKYVLVDDWAVYNGKTIDLRPKIEGILEGMESGGVVEFEGVIVQPRFPGKPADVGGLS
KTVRLDEFLEIAGGNDTLVFERVAFRDPFLIVYSSGTTGVPKCIVHSTGGVLMNVCKESILHKEMTPESVILQYTTTGW
IMYVVSVQVCLFGTRSILYDGSPFQPSPEAFLSILEEQRVTDFGTSPRFLHELQKRNISPRHLFQLSHLKSVGTTGMVLT
EAQFEWFYDAGFPASVHLRNQSGGTDIAGRFGLENPLQPVYAGGCQGPALGTKIEVYDSLIESGPGCAIPDGEPGEL
VATASFPNQPVYFWGDDAQNTRYQSAYFTRFPGVWTHGDFIQMHPITGQILFLGRADGVLNPSGVRFGSADIYSVI
ETHFPEVADSICVGQRRPGDVDESVILFLKMNKGFGYTESLVNKIKSKIAEERSRRHVPKYVFQTWDIPTTVNLKKVE
LPVKQIVSGKKIKPSGTLANPESLKFYEQFADVENVVRRMTNEKKSYRRSTL*

>g10937.t1

MNGTKGTNGSPTGPKLLWEHPSPKTTPMYQFLKLVNETHNLRLSNYSELHAWSINNVNNFWQRAWDFVGVRHQ
GTPTSAVDDDAPMFPRPDFFPRATLNFAENLLFPTATVDASSPAVIAATETTRETVTWQELRDRVKLCQAGMIHLGL
KEGDRVAGYVANHTSALVAMLAATSLGAVWTAVSPDTGVHAVLDRLRQIEPALLFADNAAFYNGRSHPVLPKVADI
ARDLPSLQAVVIFPTVASVQFDVTSIPVESGKTYDYATFTALKSMSELVFKQLPPDHPVYILYSSGTTGAPKCIVHGAIG
TLLQHKKEHIIHCSMTPRSRLFYFTTCTWMMWHWLVSGLASGATLVLFDGSPFRYVNKSDPSTSVADDLAMHRLID
EFGITHFGTSAKYLSILEQKSVDPSAAGLVMDKLEAIYSTGSPLAPSTFSYVYSAFPSTINLGSITGGTDIISLFGAPNPLL
PVYEGEIQGAGLGMAIAAYDYTGADVSASGEPGDLVCTKPFICQPVGFWGSEGDSKYWKSYFDKFTNEKKQPIWH
HGDFVRFNPATGGLWMLGRSDGILKPAGVRFGSAEIYNVVLQHFPEDVADALCIGRRRESDTDETVVLFLKMAEGR
QISDDLVGKIKTTIKQALSARHVPAVVEECPEIPVTTNGKKVEGAVKQILCGLNVKTSASVANAECLDFYRDWARTH*

>g3659.t1

MLQTEVEYQTPVMMATAVSTPRKLWEHPDPKSTAMWKFMQDANARRGLHMQTFRDLYNWSVGPSRTDFWVD
MWKASNLIYSGTYTTVVDTSLPMEINPHWFQGTYLNFAENVLFTASPTDSTVRTTTHKEESKIALTEIREGNTEVRHL
TWGDLRRRVGVLANALRARGVGKGDRVAVIASTSFDTFIAFMATVTVGALFSSSSTDMGTKGILERLLQIRPKYVLV
DDWAVYNGKTIDLRPKIEGILEGMESGGVVEFEGVIVQPRFPGKPADVGGLSKTVRLDEFLEIAGGNDTLVFERVAF
RDPFLIVYSSGTTGVPKCIVHSTGGVLMNVCKESILHKEMTPESVILQYTTTGWIMYVVSVQVCLFGTRSILYDGSPF
QPSPEAFLSILEEQRVTDFGTSPRFLHELQKRNISPRHLFQLSHLKSVGTTGMVLTEAQFEWFYDAGFPASVHLRNQS
GGTDIAGRFGLENPLQPVYAGGCQGPALGTKIEVYDSLIESGPGCAIPDGEPGELVATASFPNQPVYFWGDDAQNT
RYQSAYFTRFPGVWTHGDFIQMHPITGQILFLGRADGVLNPSGVRFGSADIYSVIETHFPEVADSICVGQRRPGDVD

ESVILFLKMNKGFGYTESLVNKIKSKIAEERSRRHVPKYVFQTWDIPTTVNLKKVELPVKQIVSGKKIKPSGTLANPESL
KFYEQFADVENVVRRMTNEKKSYRRSTL*

>g616.t1

MSSLPPVYIVSAARTPTGAFLGSLSSLSAVQLGSHAIKAAVERAGLKPTDVDEVFVGNVLSANLGQNPARQCALGAG
LPESTVATTINKVCASSIKALIVGAQTIITGNADIVVAGGTESMSNTPHYLPNMRTGAKFGDQSLVDGVLKDGLTDAY
KKEHMGLQGEECADDHGFSREQQDDYCIRSYKKAIAAHDAGLFKEEIAPIEVPVGRGKPPVVVDRDDEPKNFNEAK
TRTLRPSFKPGNGTVTAANASPLNDGAAALVLASEEAVKKHGLKPIAKILGWGDAEHNPSKFTTAPALAMPKALKH
AKVELSAVDAFEINEAFSVVALANMKILGIEEDKVNLHGGAVALGHALGASGARITTTLLGVLKEKKGKIGCAGICNG
GGGASAIVVESLQ*

>g319.t1

MLGYLTSRWQSTGEQKKTSPIWFDRHFSPILLSVAKKACTHPIHTIVTIAFLASYSYLGVLDRGLLESGIEDVSSRVDFH
SLLAGSKTLRVGEETSWQWEAEEPKTPLYKDKTAQELALVTLVFPKSSTINSAPSQQSVPNNVSAKLLPSSSSSFSTLS
HDTSLAFSIPYTEAADFLQAMQEISAPEDAAQSHGSEKEGTREEKKWMMKASKNGSAPGGVRNWIIESWTSFVDL
LKNADTGDIVIMALGYLAMHLTFVSLFLAMRRLGSNFWLATAVLLQSAFAFLFGLAVTTYLGVPINLILLSEGLPFLVVII
GFEKPIVLTKAVLSASLDGRRAAEEKRSEPLTIQSSVQTAIKRTGFEIVRDYFFEILILVAGAMSGIQGGLRQFCFLGAWI
LFFDAIMLLTFYTAILTIKLEINRIKRHVALRRALEDDGVDGKVAENVARNNDWPNARDVQVVSHTTTVFGKKITVPK
FKILMVAGFILVNVLNVVTLRFGLSPSKVSHVSGVGATPPLDPFKVAGNGLDHIYEHAKSTTTSTIVTILMPIKYELEFP
SIHYAEPSITDNEFAFGSNISTHIVDGVLKSLEDPFLSKWIVLALVMSVVLNGYLFNAARWTIKEPHQPLAPPSPSEVQ
DGAPTVPGTPRVPSMHMPTPPRTPGPEDQNGRLQPLTQIATRTPEIPVGPSEEQQRQPNRPFEVLDLMIKEKQAP
KMTDEELIEMSLKGKIPGYALEKTLGDKTRAVKIRRGLVSRTHATRETSTLLERSLLPYKDYNYDLVHGACCENVVGYL
PLPLGVAGPILVDGQNYFLPMATTEGVLVASTSRGAKAINAGGGAVTVVTGDGMTRGPCIGFDSLVRAGAAKNWL
DSEEGQRTMKDAFNSTSRFARLQSMKSAIAGTNIYVRFRATTGDAMGMNMISKGVEHALTVMATDCGFEDMRV
VTVSGNYCTDKKSAAINWIDGRGKGVVAEAMIPGSVVKSVLKCEVEDLVQMNISKNFIGSAMAGAMGGFNAHAA
NIVAAIFLATGQDPAQVVESANCITIMHNVNGNLQISVSMPSIEVGTIGGGTILEPQSAMLDLLGVRGAHPTSPGDN
ARQLARVIAAGVLAGELSLNSALCAGHLVKAHMAHNRSNVPSRAPTPAPATPVTGAQTPVTGGGPLSALSSGPAPR
R*

>g9135.t1

MELLEKLHPDEVNSPIIQKLGRFNGRSRILSRPTTIDFNRCTGEDAIPPEIDCISRIFRTSGVRLATSACKKAMQEAQLG
PRDITHTVAVTCTDQNNPGYDHFVCQELGLGSSVQRSLLHGVGCAGGLSALRTAADIAAAASLRGRPARILVMASLF
SDGAAALVVCNYLALEEKQTPVFELVEWSSMVVPDTSGHMSYVIKKNGMIATISKFVPEAAIKAIIPMFRHLRAAFDI
THKAASLVRSQASSFDWAIHPGGASILQGAKQALCLTDDHIRASLDVYQHHGNSSSPTVLIVLDKLRMMGEGRENV
VATSFGPGLIIEMCIMRRCRRIVVPPPFMIDKHTSCKLHVEQTEAPPAVQGNDDVTEAGERLATPPSASFKNKLKRYQ
YTETRTMQDTPLLSASPTKSRKRSRTTATTAVKSFPESSARPKKKRMPSRYADPSKYAHLSPLVDILQPNLICVFVGTN
PGVQTAAAGHAYAHPSNLFWKLLHSSGLTDRRLKPEEDRSLPDLYCMGNTNIVERPSKDAAELSKEETAAGTAKLDA
KFLKYKPEAKDFKYGWQDEEHNMGKSRDDKEERNTNGNVWKGSRVFVTTSTSGLAASLKPAEKEAIWKPFGEWV
QKRRAERNFLPRPLEAQEASDETLEP*

>g10614.t1

MAARPSNIGIKAIELYFPSQCVDQAELEKFDGVSQGKYTIGLGQTKMSFCDDREDIYSLALTTVTSLFKKYNVDPKSIG
RLEVGTETLLDKSKSVKSVLMQLFEESGNYNIEGVDTVNACYGGTNALFNSVNWMESSAWDGRDAVVVAGDIALY
KKGNARPTGGAGCVAMLIGPNAPIVMDAGQRGSYIRHAYDFYKPDFTSEYPVVDGHFSVRCYTEAVDACYKAYNE
REKTLKSQQNGNGVNGSQELETALDRFDYMAFHAPTCKLVAKSYARLLYNDFLANPSNPIFAEVPAELRDLDYATSVS
DKTVEKVFMGLAKKRYASRVQPSIQVPTQCGNMYCGSVYGSLCSLLANISSQDLEGKRIGMFSYGSGLASSLFSFTVK
GSTENIAKQLDIQNRLEKRRVVAPEVYDEMCNLREQAHLQKDFTPKGSVDTIVPGTYYLTGIDSMFRRSYEVKQ*

>g12302.t1

MEVPTATSLRDIYPEDALPVETKRWESLLAKFKDLYGKQADFVARSPGRVNIIGEHIDYSLYEVLPMAITADFIMAVAV
RPTEEKPRIRIANLNSEKFPTREFDIPEGEIPIDASEHEWTNYFKSGLKGVSQLLQKKRSKFTSVGMDIVCDGTVPSGG

GLSSSASVVCTSSLAVLAANGEEKIDKTELCELAIVSERAVGVNSGGMDQAASVFSLRGSALYVSFKPSLNYTNIEFPK
TDPELVFVTAQSFVAADKHVTAPVCYNLRVVECTLAAVFLAKAFGLKKDLPTDSSPLGVSLRGFHDTYFEDKEGVSDN
TKISVSEFETQLTKLIQHTEDYLPQEDGYSRQQICGLLGISEDELNQRYMSKFPVRAEKFMLRQRALHVFTEALRVIKF
RSLLASPPSNDRAYLQSLGDLMNTTQDSCREIYDCSCPELDELCDLARAAGSCGSRLTGAGWGGCSVHLVPKDKVE
AVKKAWVDKYYKKKFPDITEEKLKQAVVVSEPGSGSMLFKVTGNKLA*

>g7246.t1

MSDIHRASTTAPVNIAVIKYWGKRDPKLNLPTNSSLSVTLAQSDLRTHTTASCSSTYPKEDTLLLNGQSQDVSGART
QACFRELRALRKQLEEKDSSLAKLADLPLRIVSENNFPTAAGLASSAAGFAALVRAIANLYELPSSPTDLSRIARQGSGS
ACRSLFGGYVGWEQGSAPDGSDSVAFQVAPASHWPNMRAVILVVSAAKKGVSSTTGMQTTVATSSLFQSRATETV
PRRMKEMQEAIQNKDFEAFGKVTMMDSNSFHATCLDTFPPIFYLNDVSRAAIKVVESINAAAGKIIAAYTFDAGPN
AVVYYLEENEKEVAGLFKQILNEKDGWQGERGQAIQANADSLEKVKFDAGPAIAFLEEGVSRVILTGVGEGPVKTEH
SLIDEKGEPINSA*

>g4553.t1

MATLANLKRTLVQAAITDTAILSSRRVPLTDAQYAAGFSAFTQGPSWSSYCDFIIPELSYLLAPLEKSNASISVLEIGPGP
KSVLGDLPESLRARIRRYTAFEPNALFARKLEEWLDGKEEMSKNLGSPFAGHKGDFIIHQAPFNVDNDTTIEVSSVEK
YDIILFCHSMYGMKPQHKFIEQAIKMLVDGGIVAVFHREGLHIDGLACHHSASYPIGVTGVQDNDDALDCFASFVAG
FTIHDVDEKKAIQPIWRKTCRALARRENAYPNTLLFSAPEVMVAFNQHANTVPELTIDVLPVLEHKPVKNKEACSQR
PATIVRPAYVNDVQTCVWWALEHGLGLTIVGGGHSAHCLRSNVVAIDMSAFDCVSVLAGEDREEGLGNDSSPLVVV
GAGCKADKIINNMQSEGLTIPLGSRPSVGAGLWLQGGIGHLTRLHGLTCDSIVGAVMISVASGEIIHVGCVPSKHVPA
ASSRPENEADILWALKGAGTNFGVVVSVTFKAYPAPTYVVRNWVLPLTNHADAKRKLKEFEEKVANAYDNTYSADA
YLYWENERLNLGITTFDTTASYMTTRSKLNGDIWGTENDFRTMNGFEVFEAEMYMSAMHGGHGGGKTSSFKRCV
FLKAVGGPRVAHFLVTAVESRPSSFCYLHLLHGGGRDGEVTPDTTAFGCREWDFACVITGVWHRDQDGTKIAQASV
QWVYEVVESLLLLSNGIYGADLGPDPRDAVLAQRAFGPNAVRLGRLKRVMDPHHVLAWTCPLPKPPLPKLITLVTG
GHGAGKDYCASVWCSSFSDMQENETFSTHCTHARVVRISDATKREYATETHADFRRLVEDREYKEEHRAKLTAFYKE
QVRRRPRLPEEHFLKLVHDAADADVDVLFITGMRDDAPVAFFSHLVPESRVVDVRVQTSENSRVARGVVSDSKANV
TALSYRPNRIFENNLPGPRAAHEFAQSCLHPFIHSDIQRLADMVRAVPDFPKPDIQFRHVLGISQQPGGLTLCASLLQ
SHLTESWAGVDAIVCCEVGGLVFAPALASLVGVPLVPIREAGKLPPPTISVAKDTSYISSLTRGQTNVGKRIEIERGAIVG
CESAVVVDDVLSTGETLCAVLRLLKKADVEKIVVMVVAEFPLHRGRRLLQKRGFGMVRVQSLLVFGGQ*

>g9540.t1

MAANSSMRRNTVVAPPSLHPRTSSHSTQTPEPSPIDSMPAQLGSNMPSISTTKPNGAPALLRKQSSPMMPPFMVS
APGKVIVYGEHAVVHGKAAIAAAISLRSYLHVSFLSKSNRTVKLRFPDIQMEHTWNIDDLPWDSFSKSGKKKYYYDL
VTSLDPDLMAAIQPFIDEVSPKAPESIRKIHHASACSFLYLFLSLASRKVPPCVYTLRSTIPIGAGLGSSASISVCLSTAML
LQIRALSGPHQDQPPQECELNIERINRWAFVGEMCIHGNPSGVDNTVSSGGKAVLFQRRDYDKPPLVIPLHSFPELP
LLLVNTRQSRSTATEVAKVANLRNVHPALTENILNAIGLVTESAHKLLTSPDFDATSPDALKHLGELVTINHGLLVSLGVS
HPKLERIREIIDHTGIGWTKLTGAGGGGCAITILKPQPPALTNGHANGLHHDISSDESDTSDEADIDCGASIISNGTKLK
YKILDSLEVKLENEGFEKFETTLAGDGVGVLWPAVLHNGNEEEGGEEIDQEKFLRAEGTIGIERLVGVSSSRRKGVREV
REGWKFWRPWEVPER*

>g8301.t1

MSSSDAPLSPKAVSCPAKVLVAGGYLVLDREYTGLVFGLDARIHTVVEPIKTRSGVTINEILVTSPQFREAIWEYGYRSQ
SEDGGITITQLSVGHEQSIARSRNPFIETALTYALTYIHALLPKTLIQPSSIRILADQAYYSNPGVTRSSNQISQPHKISRFQ
DFNVSLKEAHKTGLGSSAALVTSFTAAVLGFYLPRELFDVRTEKGQTILHNLAQASHSHAQGKVGSGFDIASAVFGSC
LYKRFSPSLLGNLPQPSSPGFATQLRSLVEGPTWDTEIQKAAIKMPEGLRLVMCDVDCGSETPGMVKKVLAWRSQK
PEEAEKIWKELQSGNEALAAELTRLATEVKGDNASKHDTLRKIIDGNRALIRDMGEKSGVPIEPPQQTRLLDYCSKLD
GVVGGVVPGAGGFDAIVLLVEDKEAIIGSLKTSLAEYKDPEAIGRVGVIGVREEMVGVKEEELSLYKEWEAAS*

>g6628.t1

METLDNTEWDVLIVGTGLQQSLLALALSRSDKKILHVDENDFYGGAEAAFSLQEAEEWAQRMKDDTVDAVFSDVT
ITKPEVADAAPALSFSRAYSLSLSPQVIYARSSILGCLVSSRVYRQLEFLAVGTWWVYSTGAQSESSLSHARLLKVPNGR
EDVFQDHDLDFKAKRALMKFLRFISEYEEQIEVWEEHRQRPFSDFLSEQFKVPASLQGPLLALTLSPAGPDRTTTEYAL
PRIARHLRSIGVFGAGFGAVIPKWGGLSEISQVSCRAGAVGGGVYVLGKGIAPVTEGIAQTTENGTKLRLKDGEVVT
AKWIVGGNSSIASQDTYCRSMTIVSSSLSHLFPPIAEEAPAPAAAVVVFPSGSLTLNSQAEELPPVHVFVHSSDTGECP
SGQCVLYASTSMHNQDGFALLRKAIESLLSAQDIAPSPTILWSVEYQQRASSGSEALPSDNDHVVRFPPTSMDLAFD
DAVLENVKDMWQKIVGDDAGEFLVFQDREAYDDDE*

>g11037.t1

MPATSPNHPFASPNAIPPRTSSTGILNLNANTGAAKQPSRTSVLRPLSEIDWLGQSKKSKTSHSADPLNAPFQPQSL
QHPWPQTMASQLNSPPRTEHTDMDSPMDNAQAAVSLEATTNYPTPLSPPSEHAKDIGEELIYGNGVAWTEAKER
ILLGPYDYLYGHPGKDIRSQCIAAFNLWLKVPSERLEIITKVVGMLHTASLLVDDVEDSSLLRRGIPVAHSIFGTPQTINS
ANYVYFRALSLLLSMNNPKLIEIFTEELLNLHRGQGMDLYWRDSLTCPSEADYLEMVGNKTGGLFRLAIKLMQAESK
TDIDCTPLVSTIGLLFQILDDHLNLSPTSGYTTLKGLCEDLTEGKFSFPVIHAIRADPSNQILINILKQKTTDEEVKRYALRY
MESKGSFEYSKSVIEELRSKTDEHVRVIERELGQEGREGAEALRVMLARLVLK*

>g4503.t1

MASNGLQDYSDFSLPIPPNRYIDDPEYFCRFHPRVHKDTQASDDATVQCQIDVFGRANVGFVKGCLSLRSGGWTSL
TYPYCLPERLALVSYINEVAFFQNVGGRKKAVDPGWQNRHTQVFAKIAIQLVDTDSQLGSLVIQGLQTWKRGEAM
MEEEIRYQNLTQYLSDRLVDNAFEFLFAVARFGSNFSLTKEEEKLIEPVLKPYYEMMILTNDYFSWETEYTTFLRSNEKV
VVRNAVPLFMEWYTMSSQEAKVALKEKIQSLENDYCSLKAEFFSQYQIPGSSPAIMRWFEILEGVLAGNIFWSKTCP
RYNTAFGSEYKQYLAQRINEGAYFFNSSTESSAIISNDILKISKLNLSGQVVTDSEGGIGSMGRMKASLFTCERLPPLD
QSVIEYPSRYIASLPSNEIWHTFIDALNTWYQVPQHHLTIIRTVITQLHSSSLMLDDIQDHSPLRGGNPAAHRVYGIG
QTVNSAYFQCADALRQIQKISSDAVLVFTEELMALQIGQGADMYWTYHSITPTEEEYLQTVDSKTTALFRMASRLLQ
GQATMNRCMDMEGFLTLFGRYIQIRKDYQNLESSKNTKNQGFCSDFDGGKYSLPLIHASKHGSPEINAILQQRKRT
ESLTTDLKIVLLSELKAKGSLAYTLQVLQNLERAIKDELQSLESEAEIKNWLLWRILQQMSLDNHLG*

>g9594.t1

MAPAAFDANTQLNYARHIKYWRRNLKTLLPHFYTSNDSNRMLLALFTVSALDILGDLDAALSAEERQGHIDWVYSC
QLPEGGFRPWPGSNYGPLRSEENKNWDPAHIPGTFFALLTLVVLGDDLEKVKRKEILTWLVKMQRPEGSFGETLGD
GDFVHGGNDSRFGYMATAIRWILRGDLEGPCEGVPDIDVDKFVNCVRQAECYDGGISEAPFHEAHAGFTCCAIAAL
HFVGRLPLPPSQKPDSLIRGVTDVPKTLHWLVSRQTLTLDEDDGLDTLNDETDTSETCHDAHTFVKLSSRPSAQAKS
NLKGRPHIHFELEWVGVNGRCNKVADTCYAYWTSTPLQLLGRLDIIDRQPIRKWLLDKTQHLVGGFGKVTGDPPD
MYHSFLGLMVLAMFGETGLQDVDSALCITHKAKRHLESLSWRRKILGMDSSHSGTQAPSQESIGLTGDKTQIDA*

>g12046.t1

MTSRTPRLATSVARQQTSFSLSRRSACPNRSIHTHNATTPTLHHPSRRRNQSSWAAAVNVAQNVMSSPPPDAPIK
MDPFQTVAREMKFLTGNIRQLLGSGHPTLDTVAKYYTQSEGKYVRPMLVLLMSRATALTPRGSRGGMGIGAQSADI
SITNPRILADENPDQSPISSIRHDSAYTAEDSDILPSQRRLAEITELIHTASLLHDDVIDHSVSRRSAPSANIEFGNKMAV
LAGDFLLGRASVALARLRDPEVTELLATVIANLVEGEFMQLKNTARDEKNPSWTEDTVTYYLQKTYLKSASLISKSCRA
AAILGGSSPEVVEAAYLYGKNLGLAFQLVDDMLDYTVSEAELGKPAGADLELGLATAPLLFAWKDDQSLGKLVGRKFS
QQGDVQRAREIVSQSTGLEQTRALAQDYVDKAIDAISFFPESEAKTGLIEMCTKVMKRRK*

>g8650.t1

MAIKDGQSNEFQKLAYKSLSNYFQEHADEVIEIEILPPAIQPPDGITMQDGSSLGIPKKVLALAYVEARHLFFMKNQG
TQDVLASSWALQATKIILLFDPEHLTAANYRKRTLSLLRNEHGPHVGTPYHRALRQELCFLNSILTSPLHRQSKSPTLW
YHRSNIVESLRLIELNDVPQDRIAEFWHSELAAVCKSGEQHPKNYHAWQYARRLVHKIENHGLADDVARRVKDWC
CRHPSDISGWSFMLYLMPITAASLQQELVGDVIKYAISLDCTQESLWIFIRTSLALSSTTFLTSYQSLQAYKKDLEETDRH
AVAMERVCNAITWIDTHRQSGT*

>g11334.t1

MAMPLRPSAQAPSSRLQEIIPGSERIEELSDEELGSDYEDMGTATEEEQANIAYLESIRVPIKDSLVTETSETDYETAKNI
LPYLEGNPNDFSLNTFGIPNLQRPKHDDFLRGQLGDYPARAAGLDAARPWLLYWSLQGLTVMGSDITSYDKTVPHT
FSLAQHPDGGFGGGYGQYAHLACTYAAVLSLATVGGAQSYDTINRKALWHFLGRMKQADGGFTMCQGGEEDIRG
AFCAMVVLSLTNLPLELPPDAPVRKQGFTSFTDGLGDWISKCQSWDGGISATPGNEAHGAYAFCGLGCLAVLGPPK
ETLHKYLDVNLLIYWLSARQCTPEGGYNGRTNKLVDGCYSQWVGGCWSIVEAATTTGLWNRGALGRYILAACQEK
KGGLRDKPGKGPDAYHTCYNLAGLSAAQHKYVYDENVNKTLGTGNYGAPFHWKTEGRYDGEDVVWDEADALRA
VHPVFVIPFMAVYETRKYFEDKEGF*

>g1641.t1

MASHGISRGSGPIVRSEEARQKELQQIADYKDLADLVNAKVAEKQYTIEVLGLVTKLLNENPEYYTIWNHRRRVLIALL
TSDALEQSPEDLLQGDLHLTFALLRKFPKCYWIWNHRNWLLREGEALMGVEASHKLWSGELQLINKMLHADSRNF
HAWGYRRFVVSQIERLAASEDILTTGTTPKSLTESEFEYTTKMIKTNLSNFSAWHNRSQLIPKMLCERDADAQARRA
FLNSELALICEAINTDPFDQSIWFYHQYLLSILSPSCPSGSLVVQDLTNGERQKYYEHEMEYITEILEDEEDCKWIYEALL
GLAEAYLAVDAGTGSFTTKDMKSWLDELKRLDPLRQGRWSDLERRLDL*

>g9304.t1

MASNFIRAWLKVILLSPYRLLRALLATTANSVNPKARGIKGKMPKYYHDDEAWADIVPLPQDDGGLHPLAAIAYSEE
YSEAMGYLRAVMAKNEFSERVLGLTEHIISMNPAHYTVWLYRAKTISEIGRSLKDEIAWLNPTALKHLKNYQIWHHR
HTIIDELGSCEGEPEFINSMLELDSKNYHVWSYRQWLVKRFDLFDKPEELEWTHSMIEEDVRNNSAWNHRYYLVVG
GREGKPSADLVQREIEYTKAAIRKAPQNQSTWNYILGIIRAAELPKSTLKDFAGEFADVWKPDNVHSSHALDLLADIY
AEEEDSKENAEKALELLATKYDPIRANYWNFRKGLLDHPKVAA*

>g9658.t1

MASMPDEMHLFVDKHVRYIQSLDTRKDELEYWLTEHLRLNGLYWGLTALHLLGHPDALPRTTILDFVFSCMHDNG
GLGAAPGHDAHMLYTVSGVQILATLDAFGDLEDRIPGGRLKIGKFIADLQHRETGTFAGDEWGEQDTRFLYGALNA
LSLMGLLELVDVEKAAQYVDSCANFDGGYGTSPGAESHSGQVFTCVGALTIAGRLDLVNQEKLGAWLSERQLKNG
GLNGRPEKKEDVCYSWWVMSSMAMLDKLHWIDGQKLTNFILQCQDPELGGLADRPGDMVDVFHTVFGIAGLSL
LKYPGLEEVDPVYCMPRSVTRRCLGHLSNYYKVKVRSTHEEASRRLVETYDQWYAEWAVEELMSERMNQKDKRRT
RARTTSRPVPTPPLKIEAPVPRPVRRAAASNLLDPRLSGQQHAMIKLEHSPSPTPQANGPVLRNRNTFAPHMQYW
PTESRASSRSYTNQDYDTSSEYSDPSERRRPYRYTAAAKYAIEDESAEVLDPMAVSESTKLKGVYWPGMDIFDSATPE
MRRKRNQKKDSSVVEQLELNSQEVEATELIFTPLGTFKRQRRISCSESDEDDETEIKAESPQPVRRRPALANLDANAT
RRSTRQSKRPVFPFLSRNQYDEDRGPSGYDHSHNNYAPKRKRFEVFQDNDDVPFSQPSSMNYLTSGFTHQASPSP
APVFTSYKSFNDPFQYENKENILPPFHQTGYSNFDGQQSNGYQYPTYSYGIGPDQQAFQYTSHLYNTTSAYHQHDQ
DDDDQRTITAPPSPSTS*

>g12046.t2

MSSPPPDAPIKMDPFQTVAREMKFLTGNIRQLLGSGHPTLDTVAKYYTQSEGKYVRPMLVLLMSRATALTPRGSRG
GMGIGAQSADISITNPRILADENPDQSPISSIRHDSAYTAEDSDILPSQRRLAEITELIHTASLLHDDVIDHSVSRRSAPS
ANIEFGNKMAVLAGDFLLGRASVALARLRDPEVTELLATVIANLVEGEFMQLKNTARDEKNPSWTEDTVTYYLQKTY
LKSASLISKSCRAAAILGGSSPEVVEAAYLYGKNLGLAFQLVDDMLDYTVSEAELGKPAGADLELGLATAPLLFAWKDD
QSLGKLVGRKFSQQGDVQRAREIVSQSTGLEQTRALAQDYVDKAIDAISFFPESEAKTGLIEMCTKVMKRRK*

>g4729.t1

MAPQLGVQPSLDTIREVLAAAVKSPNPPNLVPVFSSIPAEFLTASSIYLKISAKSKSKLSFLFESAATTETIGRYSFIGADP
RKVIKTGPGHGEETDPLPLLEKELAKSRVATVPSIQLPPMTGGAVGYVGYDCVKYFEPKTRRDDMKDVLGVPESFFM
LYDTLVALDHFAQVVKVITYVKVPDSMDDLEQAYEEAKSTLNKYVTILKGKDIPLPEQGPIQLGNQYTSNIGQDGYEN
HVKELKKHISVGDIIQAVPSQRFARPTSLHPFNVYRNLRNVNPSPYLFYVDCDDFQIVGASPELLVKEEQGRIITHPIA
GTVKRGKTLQEDAALAEELSNSLKDRAEHVMLVDLARNDVNRVCDPLSTRVDKLMVVQKFSHVQHLVSEVSGVLR
PGKTRFDAFRSIFPAGSERCAQGARHGAHCRA*

>g6099.t1

MAPAILTQRPRILCLDAYDSFSNNIVALVEQNVDAEVVKIFIDDPVLAKPSASGDYSAFTDYLKGFDGIIAGPGPGWAK
CDEDVGLMKELWRLRDEQIVPVLGICLGFQSLCLAFGADIERLNEPKHGIITAILHKSQSIFRGVESLLATQYHSLQVKL
DHPIQNKRAVRYPAQLWEPTETCPQLEPLAWDFDSDLNGAVLMGVKHIQKPFWGVQFHPESICTNDEGKRIIRN
WWKDAQSWNRKRMFRNVPKHGLMPNLPSSPMTADDFRRELGVYDARKQSKRSYTDFVAKLDSDTITPFELASLI
AHGDGQSLPDLPPSTVHCATTGSGRLTVADACELFELTRGEAIVLESGLQSNLVPMAVGTGRYSIIGVVIPEETLRLHY
YAGTRRMELRDGKGQVHTDWTVSDPWPYVREVMKSLQPSTPPKGSTWAPFWGGLMGYASYEAGLETIDVHGH
KEASYPDICFAYITRSIVFDHQFKKIYVQSIRGPFDQDWVIDTTERLYEHAGFKSRETTPNSAAMRQADPFEAHGTM
HQYIDSCVQLTVGETEYCKKVSACQDAIADGQSYELCLTHRNEIRARKPTACKISHAEDNEHSWNLYKRLTGQNPAPF
SAYMRMHNVHILCSSPERYISWDRSQTAQCRPIKGTVQKKSGVTAEMAHAILSSSKERAENLMIVDLTRHQLHGVY
GSENVRVSQLMEVEEYETLWQLVSVVDAVPSGIYKPTTPEDWEDPVEYASKKPAKQSVPYLGFDAFVESLPPGSMT
GAPKKRSCEILQDVEDGARRGIYSGVLGYLDVGGGGDFNVVIRTAIKIDDEQSEEKGDVWRIGAGGAVTSQSTPQG
EFEEMIAKFGSTKRAFMPLPPPKPKTKNRRVEIIEPDDPEFAELLASMRGGEELTLDDAQIMLRAVERELRRRNAAGE
GDDITEVE*

>g1062.t1

MLSQMIKPSTMRSAGFLGRMTKNRVQARSLATVEGNTQRAIPTPSMRRATAVSNEPATFTIKNGPIFEGKSFGAKT
NISGEAVFTTSLVGYPESMTDPSYRGQILVFTQPLIGNYGVPSNARDEHGLLRYFESPHIQASGIVVQDYALKHSHWT
AVESLAAWCAREGVPAISGVDTREVVTYLREQGSSLARISIGEEYDADEDEAYIDPEAINLVRRVSTKAPFHVSSSLGD
MHVALIDCGVKENILRSLVSRGASVTCFPFDYPIHKVAHHFDGVFISNGPGDPTHCTTTVHNLRKLFETSQVPVMGI
CMGHQLIALAAGAKTIKLKYGNRAHNIPALDLTTGKCHITSQNHGYAVDPTTLSSEWREYFTNLNDQSNEGLIHNSR
PIFSAQFHPEAKGGPLDSAYLFDKYMENVQQYKSHQNSFSEKNNKPSPLLVDLLSKERVGCHPDAPDFEGHAAGM
ANEEITVGGPVAPSYQPITQKPVASAA*

>g2874.t1

MATDNSDPIPPHKTFDTILVLDFGSQYTHLITRRLRELNVYSEMLPCTTKIADLPFTPKGVILSGGPYSVYEEGAPHVD
HAVFDLGVPILGICYGLQEMAWHFGKNAGVAAGEKREYGHANLKVESHGGHMDQLFKDLGNDLEVWMSHGDK
LSHMPQDFMTVATTTNAPFAGIAHSTKKYYGIQFHPEVTHTKKGKVLLKNFAIDICEANTNWTMSKFVDQEITRIRK
LVGDKGQVIGAVSGGVDSTVAAKLMKEAIGDRFHAVMVDNGVLRLNEAVQVKKTLDEGLGINLTVIDASDLFLDRL
KGVTDDPEKKRKIIGNTFIEVFQKQAEKIKAEAHESADAGDIEWLLQGTLYPDVIESLSFKGPSQTIKTHHNVGGLPKD
MKLKVIEPLRELFKDEVRELGKELGIPEDLVWRHPFPGPGIAIRILGEVTREQVRIAREADNIFIEEIKAAGLYKKISQAF
AALLPVKAVGVMGDKRVHDQVIALRAVETSDFMTADWYPFDGEFLKRVSRRIVNEVNGVCRVVYDITSKPPGTIE
ME*

>g4728.t1

MDTCIALRTMLVKDGIAYLQAGGGIVFDSDPYDEWMETINKLGANTHCITSAEEKHLAEQQDSADDGEKDASTLEG
DAKTRISLAA*

>g118.t1

MPALALIDHSPNNPTPSPPIPTASNVILIDNYDSFTWNVYQYLVFEGATVTVYRNDEITVEELIAKNPTQLVISPGPGH
PERDAGISNAAIKHYSGKIPIFGVCMGEQCIFYSYGGTVDVTGQVLHGKTSPLKHDGKGVFAGVSQNVPVTRYHSLA
GTHGTLPDCLEVTATIPANEDAEVKEVIMGVRHKEYVMEGVQFHPESILTEDGRLMVRNFLKMQGGTWTENERLQ
KEAHAQAVGAAANGTNGAKKDKQTSILEKIYDHRRASVAEQKKIPSQRPSDLQASYDLNLAPPQINFPERLRQSPFR
LSLMAEIKRASPSKGIISLSACAPAQARLYAKAGASTISVLTEPEWFKGSIDDLKAVRQSLEGMPNRPAVLRKEFIFEEY
QILEARLAGADTVLLIVKMLDEVVLKKLYDYSRSLGMEPLVEVQNAEETEIAVKLGAQVIGVNNRNLVNFEVDMETT
NRLINMVPKETILCALSGIAGPKDVEPYIKSGVGAVLVGEALMRASDTAQFIAELLGGSSTKKAQTASSPMVKICGTRS
AEAAKAAVEAGADLIGMILAPGTKRTVTAETALAISEMVHKTKKPIVSKSGLLADSKAASDFFEHGASRLVSNDSRAL
LVGVFRNQSLGYVLEQRRLLSLDVVQFHGQEPLEWAKLVPVPVLRAFNPQNLGIGSRGYHALPLLDAGSGGSGQQL
DLSDVKAVFAKDDGIKVILAGGLNPDNVQSMLAGLDEYRDRVHAVDVSSGVEEDGQQSLDKIRAFIKAAKNQQ*

>g8333.t1

MATQAGDEDKKISITPLLKRLWHESPTTKPTADEIAAALALIFTNSLSEVQTGALLTCLHFTDQDRQAEVLAKCSKAM
RDYATGIDVKGLQELINQRGRKEGGYHGGLCDIVGTGGDSHNTFNISTTSSILASALLMIAKHGNKASTSRSGSADLL
SCAPPKPPVITAIAPNTIHEIYSKTNYAFLFAPIFHPGARHAASIRRQLGWRTIFNLLGPLANPLHDLIEARVLGVARKEI
GPDFAESLKQSGCKKGMIICGDEELDELSCAGPTHCWRIVEDASTGEANINYFTVTPADFGLPEHPLSEVSPGQSPE
QNAQILMKILMGEVPPDDPILHFVYINTAALFVVSGICDADTSNMGEGDDGNVIKEVGPGGGRWKEGVRRAKWA
ISSGAAYSEWQKFVEVTNSVA*

>g5361.t1

MSHQENTKNVPEEHQVHRTGSWLPADHRVHKKWLSGIIERVEGNPKELHPVLREFKDLIENNTRIYILVNSMFEEIP
TKKPYKNDPVGHKQVRDYHHMLELFNHILTTAPEWSDHEYSIGMVGTPFNAILDWPMGTPSGFAFFLDPEVNKMI
KKVLNAWAEFLDSPASAYVLGTDKIGWLSEHGTHDLALTANIGQTSHSFEEMFQCDPSKEHHGYKSWDDFFTRHFY
EDKRPVASPEDDSVIANACESKPYKVGRNVSKRDRFWLKGQPYSLKDMLAMDPLHEQFIGGTIYQAFLSAMSYHR
WHAPVSGKVVKSYLVEGTYFSEPLFEGLGDPSAKGGIDEEGEKTGQGYLTATATRAIIFLEADNKDLGLVCFMGIGMT
EVSTCDTTVKVGQHVKKGDEMGMFHFGGSTHCILFRKGVELEGFPDTQNVEHNMPVRAKLATVKKAT*

>g2633.t1

MDGIKKTFAQCKKEGRSALVTYVTAGFPTAEETPDIMMAMEAGGADIIELGMPFTDPIADGPAIQTANTQALKNGV
NIGSVLQMIRDARKRGLKAPVLLMGYYNPLLSYGEEKMLQDAKEAGANGFIMVDLPPEEALRFRNFCRSYGLSYVPL
IAPATSEHRMRVLCKIADSFIYVVSRMGVTGASGTMNAALPQLLERVHKYSGNVPAAVGFGVSTRDHYLSVGKIAE
GVVIGSQIINTLLKAEPGTGAKAVEKYCDEICGKSTRGAPREVGIIETLNEAKEPTNVHVDKVITDKDTPDGPGLADQL
EMLNTDDANGTNGTHEQNGFDEKHKFPARFGEFGGQYVPESLMDCLSELEEGFNAAIEDPKFWEEYRSYYDWM
GRPGHLHLAERLTEHAGGANIWLKREDLNHTGSHKINNALGQVLIARRLGKTEIIAETGAGQHGVATATVCAKFNM
KCTIYMGAEDVRRQALNVFRIKLLGAQVVAVEAGAQTLRDAVNEAMRAWVVHLDTTHYIIGSAIGPHPFPTIVRTF
QSIIGNETKEQMQAKRGKLPDAVVACVGGGSNAAGMFYPFSKDLSVKLLGVEAGGDGVDTDRHSATLSAGSKGVL
HGVRTYVIQNKHGQISETHSVSAGLDYPGVGPELASWKDSDRAKFIACTDAEAFIGFRLLSQLEGIIPALETSHAVFG
AIELAKTMNKDQDVVICVSGRGDKDVQSVAEELPKLGPKIGWDLRF*

>g10837.t1

MSSPAPPASLTRASVEAAHALIKPHIHDTPVLTNTTLTNIANTRQAPEALQGTEWEGHEPAHPRVKLFFKCENLQRIG
AFKVRGAFHAVTRLIEKEGLEQVQRKGVVTHSSGNHAQALALAAKTFSIPAHIVMPSISTPSKIAGTRAQNANIHFSG
STSTEREAVVADVIKDTGATLIPPYDHPHIILGQGTMALEIQDQVDKLLASGEKLDAVIAPCGGGGMLSGIAVALHGT
GVRVFGAEPSFQGGDDARRGVEAGERVTSVKTLTIADGLRTPLGHHTWNIISNKDYVQALYAVTEQNIKDAMKLVL
ERMKCFVEPSAVVGLATILYNEDFRNMVQREAADKTWNIGVVFSGGNTTIEAITKIFAEVPENRAERQEGVLGRDG
RRVAENVAG*

>g3669.t1

MASRNYLNAYSGPDALRNYFDPDHQPMLPLVEIPPSLNPYYQDGVRIHAKMMSMHPSNNVKIMPALNMLTKEV
HPEKSKTVVEYSSGSTVISLALVSRINHGIQDVRAFLSNKTSAPKLRLMQFFGLDITLFGGPSQPEPHDERGGIHQAR
MMAEQDEGILNVNQYENDANWQSHVKWTGPQIHQQLPGISLVCAGMGTSGTMTGLGQYFKLAKSSVIRLGVCT
AAGDRVPGPRSLALLDPVEFPWRDSVDAIEEVGSKDAFGLSLQLCRSGLICGPSSGFNLQGLFNYLEKRKSLGTLSEL
AGANGLIDCAFVACDGPYQYMDEYFDKLGSTAFRPIHNENLAAVDLYRYDEAWELTPTRALSQFADNVEEYTGAVLL
DLRKPEDFVTSHIPGSYNLPLQSLNASTPSPFLDAVVLEKQWRELEATFTPDRINAHDLAGKNVYIVCYGGDTARVAT
SVLRAKAISASSVKGGITALRQELPNLQMNERGRGLVQQDWLKMPDVATKELRADSLSPQVRSLNGIVV*

>g18.t1

MNGITTEKTNDHVEKGYLKHQIPNPTVVWQSLQKVIPLQDRNIRFWWHHTGYHVACMVDASGYSIEKQYEVLLFH
LHFICPRLGPAPESDGKSRWHSLMAHDGSPLEYSWKWNTSTGKPDIRYSWEPFNPGSGRTTDPHNHALSLDYMST
VKNVLPGVDFSWITSLLEEIEKGDQKASHFLHAVEYSQTKPFGLKSYFLPRDYKILQAGSATTMNEWDEIILKLNPNN
KGRDTLMGFLSNNLQGKLLQPCVLAVDNVKPEKSRLKLYFMTPHTSFSSLREIVTLGGSRDVPEPSFQDLKSFIWTLL

GLPDDFPEDASVPAHPPVAKTWLDEENLVECFVYFFDIAPHNSDVDVKFYLPTRRYGPDDRQIATRLVEWMESRGR
GAWCGRYLQMLEKLAEHRGLENGKGLHSYISYQVGKGPEPDIKSYLTPETYHPARYMLSA*

>g3581.t1

MAQTPRKPLITIVGATGTGKSDLAVEIARKYNGEIINGDAMQLYRGLPIITNKITQDETKGVPHHLLGCISLEEETWTV
GKFVGEALRTIDEIRSRGKLPVLVGGTHYYTQSLLFQDALADEPELNLNENSQALPILEEPTDVLHEKLREVDPIMADR
WHPNERRKIQRSLEIYLRTGKPASQLYKEQKLQRDVLAAQTDGAASDSLRFETLIFWVHANKDVLHRRLDGRVDKM
IARGLLSEVEELSSFRERHESSTGATIDQTRGIWVSIGYKEFLEYQSALSDDAKMAPELEKLKSLAVEKTQAATRQYAN
RQIKWIRIKLLNALFGAGQKDNTFLVDGSDISQWEDKVVKPATAITEQFLSGQPLPPPSSLSPVAAEMLTPKREYDLG
QRPDLWQKKVCETCGTTSITENDWNLHRQSRAHRRAVGMRKKQENASKMRKGGAEDPKAEVVDVLEHYLETFP
MEQELK*

>g8776.t1

MSTIETVTQHPEITAENVLRLFPEVNTTLIGGSHNSATSDNALQGYDEEQIRLMDEVCIVLDNNDIPIGSATKKLCHL
MENIDRGLLHRAFSVFLFDSQNRLLLQQRATEKITFPDMWTNTCCSHPLGIPGETGVGLQESIQGVRRAAVRKLDH
ELGIKAEQVPIDDFKFLTRIHYKSPSDGKWGEHEIDYILFMKADVDLNVNPNEARDSRWVSQEDLKTMFQDKSLKFT
PWFKLICESMLFEWWDHLDSGLDKYMGETEIRRM*
