## Supplementary Tables S8-S12 for "Genomic and transcriptomic analysis of camptothecin producing novel fungal endophyte - *Alternaria burnsii* NCIM 1409"

**Table S8:** MSA results to explore the presence of plant CPT-resistance conferring mutations

| **CPT Resistance conferring mutations in CPT producing plants →** | **N421K** | **L530I** | **N722S** |
|---|---|---|---|
| ***Homo sapiens*** | Q | L | N |
| ***Alternaria alternata ATCC11680*** | N | L | N |
| ***Alternaria alternata ATCC66891*** | N | L | N |
| ***Alternaria alternata BMP0270*** | N | L | N |
| ***Alternaria alternata SRC*** | N | L | N |
| ***Alternaria sp. MG1*** | N | L | N |
| ***Alternaria SPS2*** | N | L | N |
| ***Alternaria burnsii NCIM 1409**** | N | L | N |
| ***Cladosporium cladosporioides MD2 transcript 1*** | N | L | N |
| ***Cladosporium cladosporioides MD2 transcript 2*** | N | L | - |
| ***Cladosporium cladosporioides MD2 transcript 3*** | N | L | N |
| ***Penicillium aurantiogriseum NRRL 62431*** | N | L | N |
| ***Xylaria sp M71 transcript 1**** | N | L | N |
| ***Xylaria sp M71 transcript 2**** | N | L | N |
| ***Xylaria sp M71 transcript 3**** | N | L | N |
| ***Naematelia aurantialba*** | N | L | N |
| ***Catharanthus roseus*** | N | L | N |
| ***Ophiorrhiza pumila*** | N | I | S |
| ***Camptotheca acuminata*** | K | L | S |
| ***Nothapodytes nimmoniana*** | K | I | N |

*** Fungal endophyte producing CPT, its derivative; ** Plants producing CPT, its derivatives

**Table S9:** MSA results indicating the residues involved in the catalytic function of DNA topoisomerase I

| **Residues involved in catalytic function →** | **R488** | **K532** | **R590** | **H632** | **Y723** |
|---|---|---|---|---|---|
| ***Homo sapiens*** | R | K | R | H | Y |
| ***Alternaria alternata* ATCC11680** | R | K | R | H | Y |
| ***Alternaria alternata* ATCC66891** | R | K | R | H | Y |
| ***Alternaria alternata* BMP0270** | R | K | R | H | Y |
| ***Alternaria alternata* SRC** | R | K | R | H | Y |
| ***Alternaria sp.* MG1** | R | K | R | H | Y |
| ***Alternaria* SPS2** | R | K | R | H | Y |
| ***Alternaria burnsii* NCIM 1409*** | R | K | R | H | Y |
| ***Cladosporium cladosporioides* MD2 transcript 1** | R | K | R | H | Y |
| ***Cladosporium cladosporioides* MD2 transcript 2** | R | K | R | H | Y |
| ***Cladosporium cladosporioides* MD2 transcript 3** | R | K | R | H | Y |
| ***Penicillium aurantiogriseum* NRRL 62431** | R | K | R | H | Y |
| ***Xylaria sp* M71 transcript 1*** | R | K | R | H | Y |
| ***Xylaria sp* M71 transcript 2*** | R | K | R | H | Y |
| ***Xylaria sp* M71 transcript 3*** | R | K | R | H | Y |
| ***Naematelia aurantialba*** | R | K | R | H | Y |
| ***Catharanthus roseus*** | R | K | R | H | Y |
| ***Ophiorrhiza pumila*** | R | K | R | H | Y |
| ***Camptotheca acuminata*** | R | K | R | H | Y |
| ***Nothapodytes nimmoniana*** | R | K | R | H | Y |

*** Fungal endophyte producing CPT, its derivative; ** Plants producing CPT, its derivatives

**Table S10:** MSA results to investigate the mutations involved in modulating drug binding to DNA topoisomerase I

| **Mutations involved in drug binding** → | **N352A** | **F361S** | **G363S /V/C** | **G365S** | **R364H** | **M370 T** | **E418K** | **I420V** | **G503S** | **D533G** | **A653P** | **G717V/ F** | **T729A** |
|---|---|---|---|---|---|---|---|---|---|---|---|---|---|
| ***Homo sapiens*** | N | F | G | G | R | M | E | I | G | D | A | G | T |
| ***Alternaria alternata ATCC11680*** | N | F | G | G | R | T | E | I | G | D | G | G | T |
| ***Alternaria alternata ATCC66891*** | N | F | G | G | R | T | E | I | G | D | G | G | T |
| ***Alternaria alternata BMP0270*** | N | F | G | G | R | T | E | I | G | D | G | G | T |
| ***Alternaria alternata SRC*** | N | F | G | G | R | T | E | I | G | D | G | G | T |
| ***Alternaria sp. MG1*** | N | F | G | G | R | T | E | I | G | D | G | G | T |
| ***Alternaria SPS2*** | N | F | G | G | R | T | E | I | G | D | G | G | T |
| ***Alternaria burnsii NCIM 1409**** | N | F | G | G | R | T | E | I | G | D | G | G | T |
| ***Cladosporium cladosporioides MD2 transcript 1*** | N | F | G | G | R | T | E | I | G | D | G | G | T |
| ***Cladosporium cladosporioides MD2 transcript 2*** | N | F | G | G | R | T | E | I | G | D | G | G | - |
| ***Cladosporium cladosporioides MD2 transcript 3*** | N | F | G | G | R | T | E | I | G | D | G | G | T |
| ***Penicillium aurantiogriseum NRRL 62431*** | N | F | G | G | R | T | E | I | G | D | G | G | T |
| ***Xylaria sp M71 transcript 1**** | N | F | G | G | R | T | E | I | G | D | G | G | T |
| ***Xylaria sp M71 transcript 2**** | N | F | G | G | R | T | E | I | G | D | G | G | T |
| ***Xylaria sp M71 transcript 3**** | N | F | G | G | R | T | E | I | G | D | G | G | T |
| ***Naematelia aurantialba*** | N | F | G | G | R | K | E | V | G | D | A | T | T |
| ***Catharanthus roseus*** | N | F | G | G | R | M | D | I | G | D | E | G | T |
| ***Ophiorrhiza pumila**** | N | F | G | G | R | V | D | V | G | D | E | S | T |
| ***Camptotheca acuminata**** | N | F | G | G | R | M | D | I | G | D | E | G | T |
| ***Nothapodytes nimmoniana**** | N | F | G | G | R | T | D | I | G | D | E | G | T |

*** Fungal endophyte producing CPT, its derivative; ** Plants producing CPT, its derivatives

**Table S11:** MSA results indicating the residues involved in drug binding to DNA topoisomerase I

| **Residues involved in drug binding** | **E356** | **H367** | **V502** | **Y619** | **D725** |
|---|---|---|---|---|---|
| *Homo sapiens* | E | H | V | Y | D |
| *Alternaria alternata ATCC11680* | E | H | V | Y | D |
| *Alternaria alternata ATCC66891* | E | H | V | Y | D |
| *Alternaria alternata BMP0270* | E | H | V | Y | D |
| *Alternaria alternata SRC* | E | H | V | Y | D |
| *Alternaria sp. MG1* | E | H | V | Y | D |
| *Alternaria SPS2* | E | H | V | Y | D |
| *Alternaria burnsii NCIM 1409**** | E | H | V | Y | D |
| *Cladosporium cladosporioides MD2 transcript 1* | E | H | V | Y | D |
| *Cladosporium cladosporioides MD2 transcript 2* | E | H | V | Y | - |
| *Cladosporium cladosporioides MD2 transcript 3* | E | H | V | Y | D |
| *Penicillium aurantiogriseum NRRL 62431* | E | H | V | Y | D |
| *Xylaria sp M71 transcript 1**** | E | H | V | Y | D |
| *Xylaria sp M71 transcript 2**** | E | H | V | Y | D |
| *Xylaria sp M71 transcript 3**** | E | H | V | Y | D |
| *Naematelia aurantialba* | E | H | V | Y | D |
| *Catharanthus roseus* | E | H | V | Y | D |
| *Ophiorrhiza pumila*** | E | H | V | Y | D |
| *Camptotheca acuminata*** | E | H | V | Y | D |
| *Nothapodytes nimmoniana*** | E | H | V | Y | D |

*** Fungal endophyte producing CPT, its derivative; ** Plants producing CPT, its derivatives

**Table S12:** Analysis of important residues and mutations in DNA Topoisomerase I of all organisms used to obtain the MAFFT alignment file

| **Mutation/Residue in DNA Top I (Number according to Human DNA Top I)** | **Significance of the residue and (or) its mutation** | **Reference(s)** | **Inference from the current study** |
|---|---|---|---|
| N421K | N to K mutation confers CPT resistance | [45] | Present only in DNA Top I sequences of *C. acuminata* and *N. nimmoniana;* Fungi have N421 like *O. pumila* and *C. roseus* |
| L530I | L to I mutation confers CPT resistance | [45] | Present only in DNA Top I sequences of *O. pumila* and *N. nimmoniana.* All other sequences show L530 |
| N722S | N to S mutation confers CPT resistance | [45] | Present only in DNA Top I sequences of *C. acuminata* and *O. pumila* with a gap in one of the DNA Top I sequence from *Cladosporium cladosporioides MD2;* All other sequences show N722 |
| R488 | Residue involved in catalytic function; contributes to CPT resistance by hampering the water enabled contact of CPT E-ring to itself | [45] | Highly conserved in the DNA Top I sequence from all the organisms included in this study |
| K532 | Residue involved in catalytic function | [13, 45] | Highly conserved in the DNA Top I sequence from all the organisms included in this study |
| R590 | Residue involved in catalytic function | [13, 45] | Highly conserved in the DNA Top I sequence from all the organisms included in this study |
| H632 | Residue involved in catalytic function | [13, 45] | Highly conserved in the DNA Top I sequence from all the organisms included in this study |
| Y723 | Residue involved in catalytic function | [13, 45] | Highly conserved in the DNA Top I sequence from all the organisms included in this study |
| N352A | N352 shows a dynamic mobile behaviour, plays a role in CPT resistance; N352A mutation | [13, 46] | N352 is present in all the sequences involved. No mutation found. |

| | | | |
|---|---|---|---|
| | renders a CPT sensitive DNA Top I enzyme | | |
| F361S | F361 to S mutation gives a CPT sensitive enzyme | [13, 45] | F361 present in all sequences involved. No mutation found. |
| G363S/V/C | CPT resistance conferring mutation | [13] | G363 found in all sequences. No mutation |
| G365S | CPT resistance conferring mutation | [13] | G365 found in all sequences. No mutation |
| R364H | R364 to H mutation confers resistance to CPT | [13, 46] | R364 present in all sequences involved. No mutation found |
| M370T | M370T mutation confers CPT resistance in CPT resistance human lung cancer cell lines | [13, 47] | M370T mutation found in all the fungal sequences used, except for that of *N. aurantialba* which had a M370K; *N. nimmoniana* showed the M470T mutation as well. While *O. pumila* had a V, *C. roseus, C. acuminata* had M; M370T mutation is present in other non-CPT producing closely as well as distantly related fungi. |
| E418K | E418K mutation confers CPT resistance in CPT resistant and part revertant human nasopharyngeal carcinoma cell lines (HONE-1) | [58] | E418 found in human and fungal sequences. E418D mutation found in all plant sequences. |
| I420V | I420 present in drug binding site of DNA Top I. I to V mutation does not give any altered sensitivity to CPT. But this could be an important residue to look at, as it is present in the drug binding site. | [45] | I420 found in all sequences, except in *N. aurantialba* which has a V420, and *O. pumila* which again has a V420 |
| G503S | Mutation of G503 confers CPT resistance due to the mutation making the E-ring contact of CPT and DNA Top I impossible | [13, 46] | G503 highly conserved across all sequences. No mutation |
| D533G | D533 is a residue that is needed for enzyme sensitivity to CPT; | [13, 46] | D533 highly conserved across all sequences. No mutation. |

| | | | |
|---|---|---|---|
| | D533G mutation potentially confers CPT resistance | | |
| A653P | A653P mutation increases the rate of enzyme-catalyzed DNA relegation, and thereby confers CPT resistance | [13] | A653 in Human sequence; E653 in CPT producing and non-CPT producing plants; G653 in all fungi except *N. aurantialba* which had A653 like human DNA Top I; |
| G717V/F | G717 in the active site helps in conformational flexibility that modulates CPT binding. Mutation of G717/V/F can confer CPT resistance | [13] | G717 found in all sequences, except in *N. aurantialba* which has a T717, and *O. pumila* which again has a S717 |
| T729/A/K/E | T729 is located in the hydrophobic cavity and its integrity is necessary for facilitating the CPT-DNA binding and subsequent sensitivity to CPT. T729/A/K/E mutation causes CPT resistance. | [59] | T729 found in all sequences with a gap in one of the DNA Top I sequence from *Cladosporium cladosporioides MD2* |
| E356 | Residue present in drug (CPT) binding site of DNA Top I | [13] | Highly conserved in all sequences involved. No mutation |
| H367 | Residue present in drug (CPT) binding site of DNA Top I | [45] | Highly conserved in all sequences involved. No mutation |
| V502 | Residue present in drug (CPT) binding site of DNA Top I | [13] | Highly conserved in all sequences involved. No mutation |
| Y619 | Residue present in drug (CPT) binding site of DNA Top I | [13] | Highly conserved in all sequences involved. No mutation |
| D725 | Residue present in drug (CPT) binding site of DNA Top I | [45] | Highly conserved in all sequences involved with a gap in one of the DNA Top I sequence from *Cladosporium cladosporioides MD2*. No mutation |
